# Development of a High-throughput *in vivo* Assay for the Determination of Adenylation Domain Specificities

**DOI:** 10.64898/2026.07.14.738513

**Authors:** Leonard Präve, Jing Liu, Yuanyuan Zuo, Londoño S. O. Nicolas, Anna Wacker, Helge B. Bode

## Abstract

Natural product synthesis by non-ribosomal peptide synthetases (NRPS) is greatly defined by the substrate selectivity of the adenylation (A) domains. Previous assays for specificity determination were mainly performed *in vitro* and were requiring protein purification. In this work, we developed - based on NRPS engineering - a novel *in vivo* assay suitable for high-throughput application named ASCR (<u>A</u> domain <u>scr</u>eening). Using the recently described XUT fusion sites, A domains and their upstream condensation domains were assembled as di-domains to characterized NRPS model system, which allowed detection of defined tripeptide products via mass spectrometry directly after cell culture extraction. We evaluated the assay by screening in total 54 A domains from five known and seven uncharacterized NRPS, covering a broad range organism taxonomy and GC content of the investigated NRPS-encoding genes. Additionally, we applied the assay to elucidate and confirm the structures of novel cyclic pentapeptides derived from three novel NRPS from *Photorhabdus temperata* K122.

## Introduction

Non-ribosomal peptide synthetases (NRPS) represent a major enzyme class involved in the synthesis of complex peptides that cover a broad variety of biological activities, including the production of antibiotics, antifungal agents or anticancer drugs.^1^ NRPS-based biosynthesis is defined by its modular domain architecture with each module consisting of a condensation (C), adenylation (A) and thiolation (T) domain. Thereby, the A domains play a central role in determining the peptides structure by selecting and activating the individual amino acids as building blocks, which are subsequently covalently bound as thioesters to the T domain. Next, the bound amino acids are connected via the N-terminal amine of the amino acid bound to the downstream-located module to elongate peptide formation along the NRPS. In a canonical NRPS, the synthesis is initiated by a starter module, which can harbor a condensation starter (C_start_) domain to introduce defined acetylation to the N-terminus of peptides.^2,3^ After peptide chain extension by variable number of elongation modules, the synthesis usually ends through termination modules that consists of an additional thioesterase (TE) domain to release the peptide either via hydrolysis or by head-to-tail and sidechain-to-tail cyclization to obtain linear or cyclic peptides, respectively.^1^ Even though, NRPS can consist of a single protein encoded by one gene, multiple proteins that interact through so-called communication (COM) or docking domains (DD) frequently facilitate the synthesis.^4^

Initially, the specificity of the A domain defines the structure of the final non-ribosomal peptide (NRP). Not limited to the 20 proteinogenic amino acids, selectivity to various non-proteinogenic amino acids like D-, hydroxy-, β-, N-methyl-, homo-amino acids or even combinations thereof have been reported^1^, leading to over 300 different A domain substrates.^5^ While a lot of non-proteinogenic amino acids are directly selected and incorporated by the A domain, several modifications of the amino acid are introduced after thioester bond formation at the T domain by in *cis* acting enzymes like condensation/epimerization (CE)^6^ domains or epimerization (E)^7^ domains to introduce D-configuration. Other in *cis* acting enzymes are heterocyclization (C_Cy_) domains^8^, N-methyltransferases (Nmet) domains^9,10^, monooxidase (Ox) domains^11^ and polyketide synthetases (PKS)^12^. In addition, *trans* acting enzymes like hydroxylases can further modify either the precursor amino acid or the bound amino acid after the A domain selection.^1^

Different A domain specificity prediction algorithms have been developed, which often showed low accuracy for predictions.^13–17^ A significant improvement to high-accuracy substrate prediction was recently reported with PARAS and PARASECT^18^, two predictors based on machine learning, which are already linked to the widely used secondary metabolite genome annotation framework antiSMASH.^19^ The major driver for the improvement was a 3-fold extension compared to previous datasets and further curation of the available data.^18^

Conclusively, further extension of the dataset with less or not represented A domain clades should further improve the accuracy of the prediction framework.

In general, the substrate specificities of uncharacterized A domains can be clarified by different methods. By determining the structure of a given NRPS-derived peptide, amino acids within the structure can be assigned to A domains encoded by the associated gene cluster. Alternatively, diverse assays have been established to characterize A domain specificities *in vitro* that are mainly based on the domain’s ATP-dependency and consumption to activate amino acids for peptide formation. For decades, radioactive ATP-PP_i_ exchange assays were used to clarify the enzyme activity in the presence of different substrates^20,21^ and was further developed into a non-radioactive method by usage of γ^18^-PP_i_ to determine ATP-PP_i_ exchange via mass spectrometry (MS).^22^ In other assays, the A domain release of PP_i_ in the presence of various substrates was determined through the presence of a pyrophosphatase to form orthophosphate which can be colorimetrically determined through malachite green.^23^ Another way to monitor the ATP consumption is the usage of the indigoidine synthetase, producing in dependency of ATP the colorimetric detectable indigoidine.^24^ More recent assays, were based on hydroxylamine to generate amino acid hydroxamates, which were initially colorimetrically verified.^25^ The assay was significantly improved by development of the multiplexed hydroxamate assay (HAMA), where A domains can be simultaneously exposed to multiple substrates in a 96-well format and amino acid hydroxamate formation is determined by MS.^26,27^ Terlouw et al. describes an alternative approach were *in vitro* and *in vivo* amino acid activation and loading to the T domain is detected via MS after purification of the expressed A-T didomain.^18^

In this work, we developed a high-throughput work frame called ASCR (<u>A</u> domain <u>scr</u>eening) to determine the substrate specificities of A domains fully *in vivo* with a minimal workload. The assay is based on the described eXchange Unit Thiolation domain (XUT) NRPS engineering strategy, which enables the assembly of different NRPS before or within the T domain to generate hybrid NRPS.^28^ By assembling C-A didomains to the model NRPS GameXPeptide synthetase^29^ and expression in *E. coli*, the unknown A domain substrates are elongated with two leucines, which allows detection by high-performance liquid chromatography mass spectrometry (HPLC-MS). Applying ASCR we screened A domains from 12 different biosynthetically characterized and uncharacterized NRPS. For three novel NRPS we isolated their cyclic peptide products and elucidated the structures, which further validated the results of the assay.

## Results and Discussion

The five modular GameXPeptide synthetase (GxpS)^29^ from *Photorhabdus laumondii* TTO1 has been a frequently used model system to study various NRPS engineering strategies.^28,30–37^ Different parts of GxpS have been assembled to other NRPS at specific fusion sites to generate hybrid NRPS. Upon expression in *E. coli* DH10B::*mtaA* these engineered NRPS showed production of predictable peptides, which were detected *in vivo* by HPLC-MS. When we analyzed previously published (Fig. S1-S2) ^28,34^ and novel generated data (Fig. S3-S5) of engineered NRPS that used the last two modules of GxpS (M4 and M5) assembled to one or two modules of another NRPS, we observed the production of tripeptides as side products in all cases. The peptides consisted of two C-terminal leucines derived from GxpS, while the single N-terminal amino acids were derived from the upstream neighboring NRPS. Noteworthy, tripeptide side products were also reported in previous studies involving the modification and splitting of GxpS by synthetic zippers after the second C domain (Fig. S6).^28,33,35^

Since the tripeptide production was not only observed upon assembly of neighboring starter modules but also for elongation modules, we hypothesized that we could transfer this observation into a simple assay to determine the specificities of A domains. Thereby, any given A domain - independent of its positioning within the NRPS - could be fused to the last two modules of GxpS, which results upon expression in *E. coli* in production of tripeptides harboring N-terminally the amino acid selected by the targeted A domain (Fig. 1). The introduced two leucines from GxpS M4 and M5 serve as a tag and allow the detection of the peptide *in vivo* within the production culture by HPLC-MS. As an initial proof-of-concept, we selected the odilorhabdin synthetase (OdlS) from *X. nematophila*.^34,38,39^ Each individual A domain of OdlS was assembled to GxpS M4-5 at the XUT^IV^ fusion site.^28^ Beside the A domains, we decided to include in all following experiments the upstream C domains – if present – to maintain potential interactions with the A domain, which might influence amino acid selectivity.^40^ For eight out of nine tested C-A didomains of OdlS, we could observe the mass of the expected tripeptide (Fig. S7-S13). However, for OdlS A6 and A9 only the adjustment to the XUT^I^ fusion site showed production of the expected peptide while for A4 no production was observed for both the XUT^I^ and XUT^IV^ sites. These initial results encouraged the further development and testing of this concept for an A domain specificity assay application.

**Figure 1.**
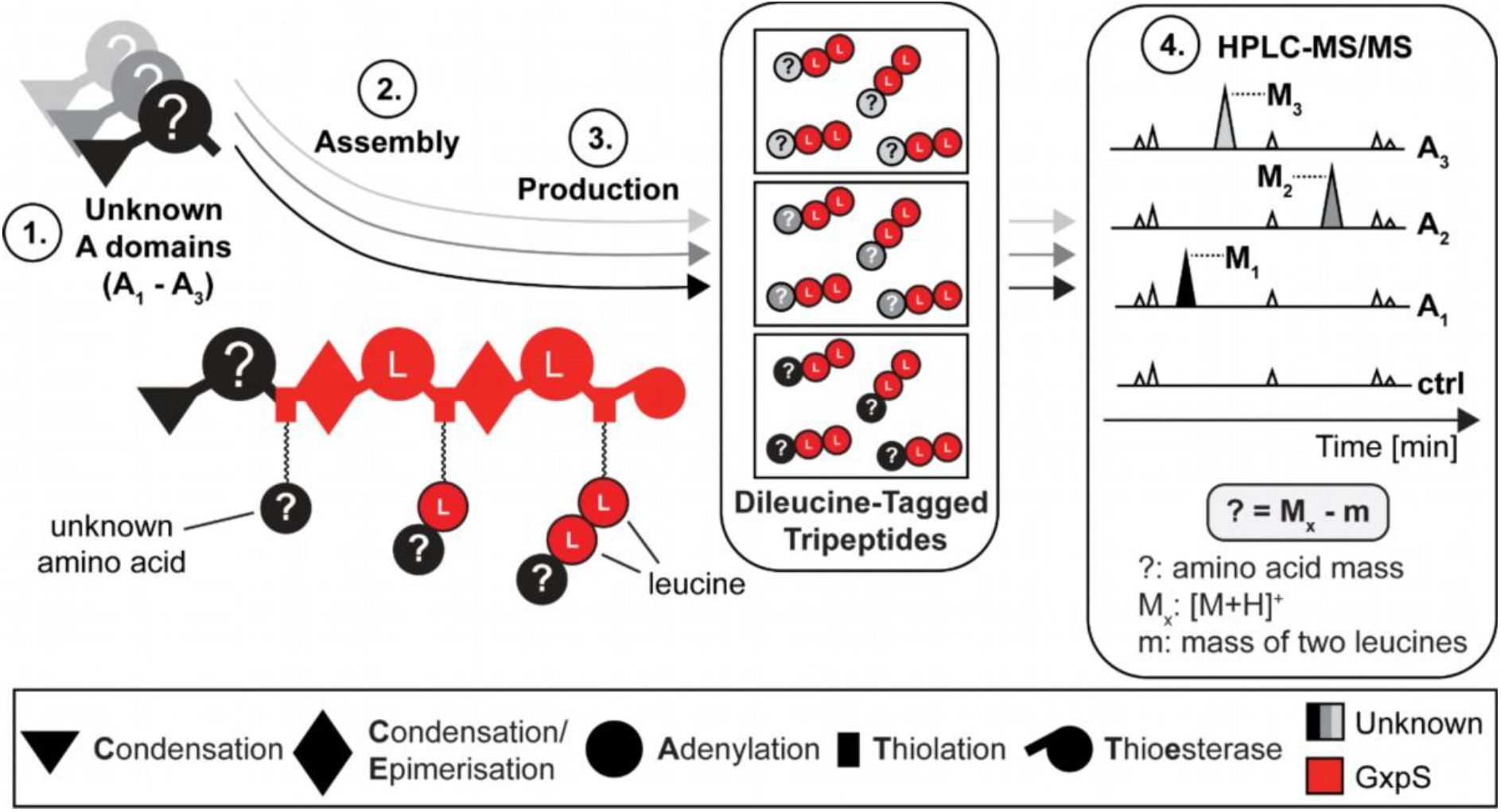
Concept of the A domain specificity screening. A domains with unknown specificity (1.) can be assembled to the last two modules of GxpS M4 and M5 at the XUT^I^ assembly site as C-A didomains (2.) to produce dileucine-tagged tripeptides (3.). The produced peptides are detected directly after cultivation and expression by HPLC-MS/MS by comparison to a control (ctrl) (4.), which allows calculation of the unknown amino acid mass. NRPS domain explanation and origins is depicted below. A domain specificity is indicated within the A domains as one-letter amino acid code.

To optimize and reduce the assay’s workflow for a semi high-throughput (HTP) application, three different cloning vectors were constructed (Fig. S14). The vectors pASCR2, pASCR3 and pASCR4 (pASCR = **A** domain **SCR**eening) contained *gxpS* M4 and M5 starting at the XUT^I^ fusion site and can be linearized by restriction digestion to introduce C-A didomain encoding gene fragments at this site via Gibson assembly^41^ (Fig. 2). Furthermore, pASCR3 and pASCR4 harbor C-terminally a monomeric superfolder GFP^42^ (*msfGFP*) fused via a GS-linker^43^, while pASCR4 contains a N-terminal *SUMO*-tag^44^ before the restriction site for linearization. In addition, four control plasmids were constructed - an empty plasmid control (pASCR_empty), an *msfGFP* expressing plasmid and two variations of a truncated *gxpS* M3-M5 with (pASCR5) and without (pASCR6) a C-terminal *msfGFP* (Fig. S15). The three T domains in the truncated GxpS were inactivated through mutagenesis of the post-translationally modified serines to alanine to obtain an inactivate NRPS, which represents according to the size of the gene a better control compared to expression of smaller genes. Next, the three cloning vectors were tested and quantitatively compared by specificity determination of six A domains from the uncharacterized NRPS PTEMK122_08040 from *P. temperata* K122 (Fig. 3a). Despite production detection for all six assembled C-A didomains, the growth was strongly reduced but could be mainly restored upon addition of 0.2% glucose, which significantly increased the production titers (Fig. S16-S18). The specificities were further confirmed by P_*BAD*_ promotor exchange based activation^45^ of PTEMK122_08040 in *P. temperata* K122 (Fig. 3). Four derivatives (**1**-**4**) were characterized by HPLC-MS with additional isotope and inverse isotope labelling experiments^29,46^, where **1** was isolated and its structure further elucidated by NMR and advanced Marfey’s analysis^47^, verifying a cyclic pentapeptide structure (Fig. 3 and Fig. S19-S29, Table S5). The last module of the NRPS does not seem to be involved in the biosynthesis of **1**-**4**. Nevertheless, the assay indicated an arginine specificity for A6, a determination in line with the prediction of PARAS.^18^ Overall, the direct comparison of the three cloning vectors indicated the best production yields for pASCR3 with supplementation of 0.2% glucose.

**Figure 2.**
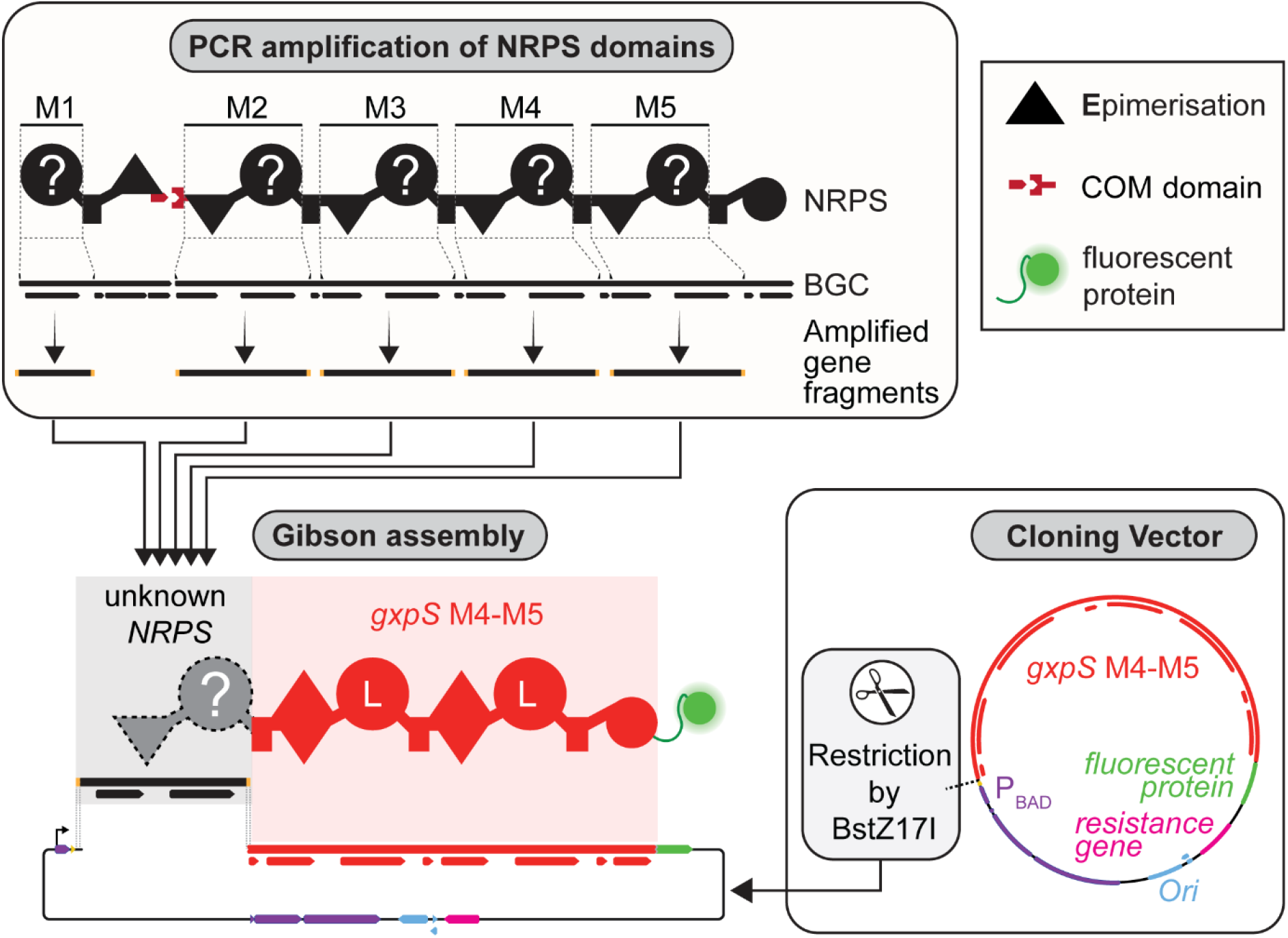
Cloning workflow of the A domain specificity screening assay. Illustration of a potential NRPS containing biosynthetic gene cluster (BGC), with antiSMASH^19^ derived domain annotations below, schematic representation of the encoded NRPS above with indications of A domains or C-A didomains, referred as modules (M) 1-5 (top-left). The potential primer pairs within the BGC and their respective PCR products are depicted. Representation of cloning vector plasmid map with indications of essential genes, promotor region, antiSMASH derived NRPS domain annotations for *gxpS* and the BstZ17I restriction site (bottom-right). Restriction by BstZ17H makes the vector suitable for Gibson assembly, which allows gene insertion of the individual amplified A domain or C-A didomain encoding genes into the vector (bottom-left).

**Figure 3.**
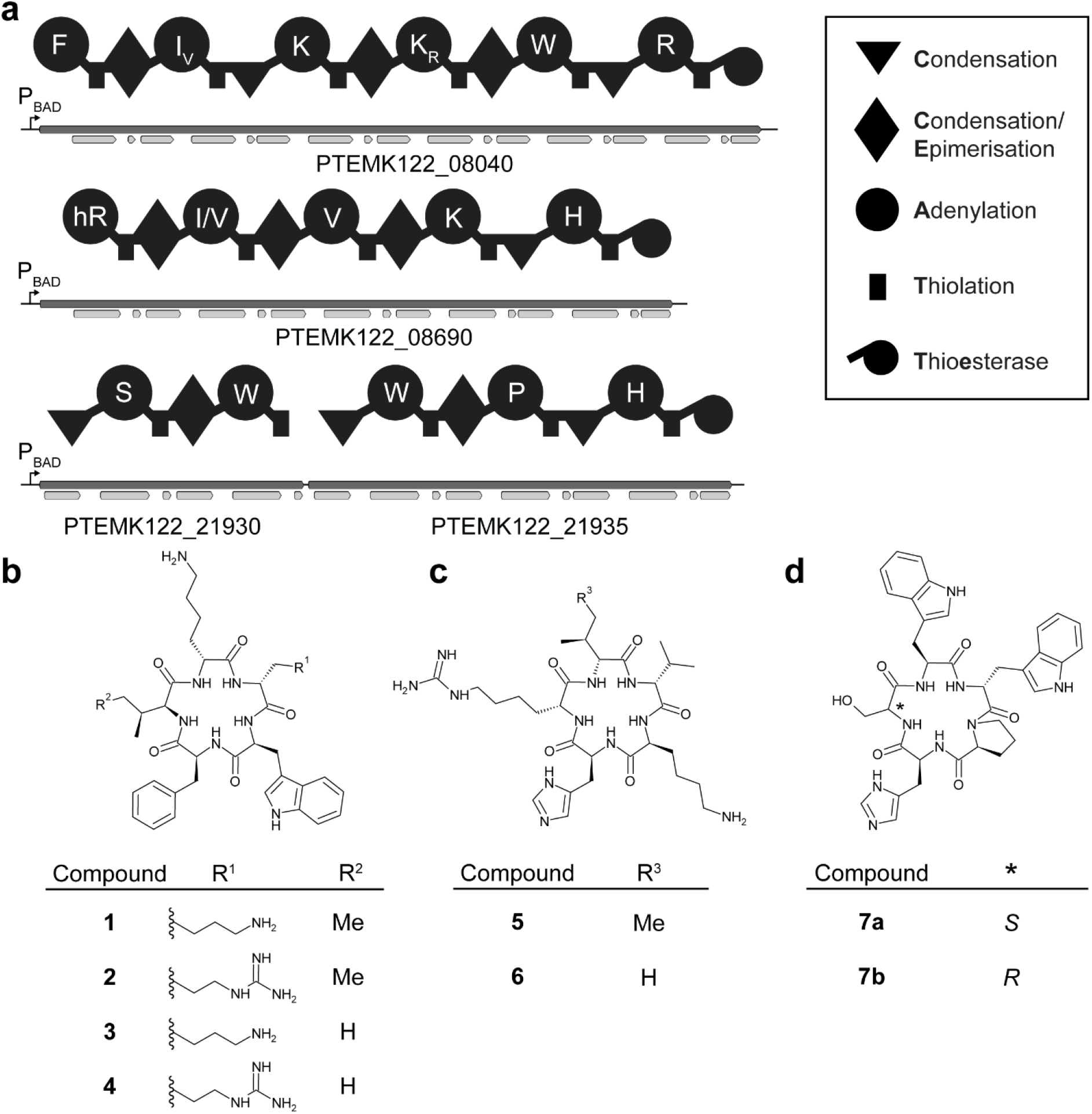
NRPS with previously unknown products identified in *Photorhabdus temperata* K122. (**a**) NRPS genes from *P. temperata* K122 activated by P_BAD_ promotor exchange activation with schematic illustration of encoded NRPS above. The antiSMASH^19^ derived domain annotations are shown below the genes. The A domain’s amino acid specificity is indicated by the one-letter amino acid code (hR = homoarginine). The identified peptide structures resulting from expression of the respective NRPS encoding genes PTEMK122_08040 (**b**), PTEMK122_08690 (**c**) and PTEMK122_21930 and PTEMK122_21935 (**d**) are depicted.

Next, the assay was further evaluated by testing a broader range of A domains in a semi HTP workflow. All following experiments were performed with cloning vector pASCR3. In addition, pASCR8, an analogous designed cloning vector was constructed, which varied in a different resistance gene, *araE* for broader expression host application and *mScarlet-I3*^48^ as a C-terminal fluorescent protein tag (Fig. S30). Furthermore, four analogous control plasmids were constructed for this new cloning vector. In total, 54 A domains derived from 12 different NRPS containing biosynthetic gene clusters (BGCs) were analyzed by the assay (Fig. 4). The dataset contained A domains from strains of the genera of *Aneurinibacillus, Xenorhabdus, Photorhabdus, Pseudomonas, Chromobacterium, Myxococcus, Xanthobacter* and *Goodfellowiella*, which covers low, medium and high GC content A domains. 17 proteinogenic and 10 non-proteinogenic amino acid A domain specificities are covered in this work, which demonstrates a broad substrate flexibility of the GxpS screening system. Within the dataset, two additional novel NRPS cluster from *P. temperata* K122 were elucidated and confirmed as described in the previous section and revealed the production of four head-to-tail cyclized pentapeptides **5, 6** (PTEMK122_08690) (Fig. S31-S48; Table S6-S7) and **7a** and **7b** (PTEMK122_21930_21935) (Fig. S49-66, Table S8-S9), where **5** and **6** showed high similarity to the recently reported piomide^49^ (Fig. 3 and 4b). Another novel NRPS cluster XSTOV2_09090 from *X. stockiae* was targeted by the assay and showed incorporation of an N-methylated threonine, threonine and a C-terminal histidine (Fig. S67-S68). However, no NRPS related product could be observed after heterologous expression of the entire NRPS encoding gene and co-expression of the downstream encoded transporter in *E. coli* DH10B::*mtaA* (data not shown).

**Figure 4.**
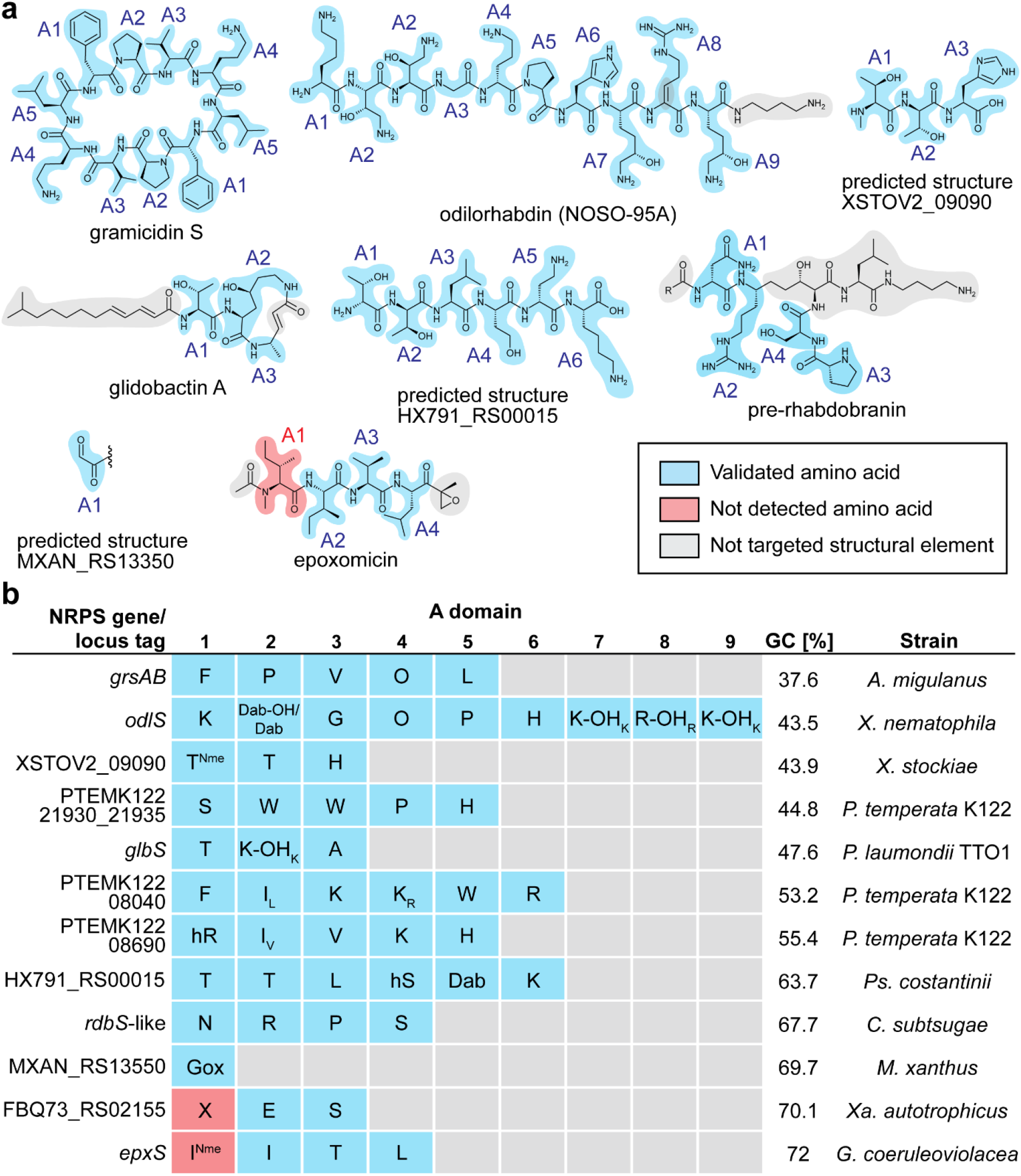
Overview of the A domain specificity screening. (**a**) Chemical structures of natural products or predicted structures produced by BGCs targeted by the A domain specificity assay. A domain origins with indication of verified (blue), not verified (red) or not targeted (grey) structural elements are depicted. (**b**) All A domains targeted by the A domain specificity assay with indications of the known NRPS genes or locus tags, the sequential A domain numbers, the BGC’s GC content and the strain the cluster is derived from. The determined amino acid specificity is indicated by the one-letter amino acid code (O = ornithine; Dab-OH = β-hydroxy diaminobutyric acid; Dab = diaminobuytric acid; K-OH = 4-hydroxylysine; R-OH = β-hydroxy-arginine; T^Nme^ = N-methylated threonine; hR = homoarginine; hS = homoserine; Gox = glyoxylic acid; I^Nme^ = N-methylated isoleucine). Amino acids written in lowercase indicate an additional secondary specificity due to A domain promiscuity detected in the ASCR assay.

For further validation of the assay, all A domains of the model system gramicidin synthetase (GrsAB)^50^ from *A. migulanus* were successfully screened with the ASCR assay (Fig. 4, Fig. S69-S70). A direct quantitative comparison of cloning vectors pASCR3 and pASCR8 showed an improved growth, expression and production profile for pASCR8 compared to pASCR3 (Fig. S71). Furthermore, all A domains from OdlS (Fig. S72-S73) and glidobactin synthetase (GlbS)^51^ from *P. laumondii* TTO1 (Fig. S74-S75) were targeted. Beside low intensities for OdlS A domains 4 and 6, all previously described specificities were verified. For both OdlS and GlbS it has been described that particular amino acids are modified by hydroxylases encoded by accessory genes located within the cluster.^34,51^ Therefore, the respective hydroxylases were co-expressed from a second plasmid, which resulted in the detection of the expected hydroxylated amino acids within the produced tripeptides (Fig. S76-S80). This demonstrate that the assay can be applied to clarify the role of accessory genes associated with A domain substrates in a defined but natural NRPS setup.

Next, A domains with higher GC contents were tested in the assay. First, all C-A didomains encoded by the undescribed HX791_RS00015 from *P. costantinii* were successfully analyzed (Fig. 4 and Fig. S81-S82). Noteworthy, A1, A2 and A4 showed the same mass and peptide fragmentation corresponding to a threonine but for A4 a differing retention time was determined during HPLC-MS (Fig. S81). The incorporation of two threonines by A1 and A2 was verified by isotopic labeling experiments, where for A4 no threonine incorporation was observed (Fig. S83). Therefore, we assigned homoserine as the amino acid specificity, which is further supported by prediction with PARAS.^18^ Next, amino acid specificities of four A domains from an NRPS/PKS hybrid containing BGC from *C. subtsugae* - assigned due to overall cluster similarity as a pre-rhabdobranin-like synthetase (RdbS)^52^ - were verified, which showed good signals in the HPLC-MS analysis (Fig. 4 and Fig. S84-S86) despite the high GC content. From *M. xanthus* an A domain with an integrated oxidase encoded by MXAN_RS13550 was analyzed by the assay, which resulted in the detection of an assigned tripeptide containing a glyoxal (Gox) moiety (Fig. 4 and Fig. S87-S88), as previously described for a similar domain in the melithiazol biosynthesis.^11^ From *X. autotrophicus*, A domains of a novel BGC were targeted by the assay, where only two out of three A domains showed production of tripeptides, which could be assigned (Fig. 4b and Fig. S89-S90). For A3 PARAS predicted with the highest score a cysteine specificity but the assay indicated serine incorporation instead. However, for A1 no specificity could be determined. Since PARAS prediction also resulted in a low score, we assume that the A domain might accept a substrate not present in the medium or metabolism of *E. coli*. Nevertheless, we also observed limitations when targeting the A domains of the epoxomicin synthetase (EpxS)^53^ from *G. coeruleoviolacea* (Fig. 4, Fig. S91-S92). Here, only the A domain specificities for three out of four domains could be validated, which shows certain limitations of the presented assay, probably still depending on proper expression and folding of the assembled C-A didomain and its compatibility towards GxpS. The expressions of all cloned NRPS hybrids from the vector pASCR8 were verified by determination of the fluorescence of the C-terminal located mScarlet-I3 and the cell growth, which in comparison to an empty vector control showed different expression profiles for the inserted C-A didomains or A domains but never no expression (Fig. S93-S94). Interestingly, for *epxS* A2 and *epxS* A4, specific to either isoleucine or leucine, we observed different retention times of the detected tripeptides during HPLC-MS (Fig. S91). To verify that the shift in the retention time can be utilized to determine either isoleucine or leucine, all A domains of the dataset specific towards these isomers were analyzed by feeding of [D_10_]-*L*-leucine and by inverse isotope feeding experiments with isoleucine.^29,46^ This further validated the observation and showed that the assay can differentiate between these two substrates by a decent retention time shift (Fig. S95-S96).

Next, the workflow of the A domain specificity screening assay was further adapted for HTP application. Thereby, we mainly focused on steps with high workload or scaling, which limits the assay’s throughput. Beside simple substitution of preparative gel electrophoresis by direct PCR clean-up purification, a major reduction of workload was achieved by omitting cell plating after plasmid assembly (Fig. 5). Instead, cells were directly cultivated in an overnight grown culture after transformation of Gibson assembled plasmids and were subsequently used for inoculation of the production culture. Even though a heterogeneous mixture of plasmids - including false assembly or mutations - might be present in culture, the major fraction of correct plasmids should result in production of tripeptides. The novel workflow was assessed with C-A didomains from three previously screened NRPS *grsAB*, PTEMK122_ 21930/PTEMK122_21935 and HX791_RS13350 by comparison to the verified plasmid (Fig. S97-S99). Despite reduced HPLC-MS intensities, clearly detectable signals were present for all A domains, which demonstrates the assay’s potential for future HTP application.

**Figure 5.**
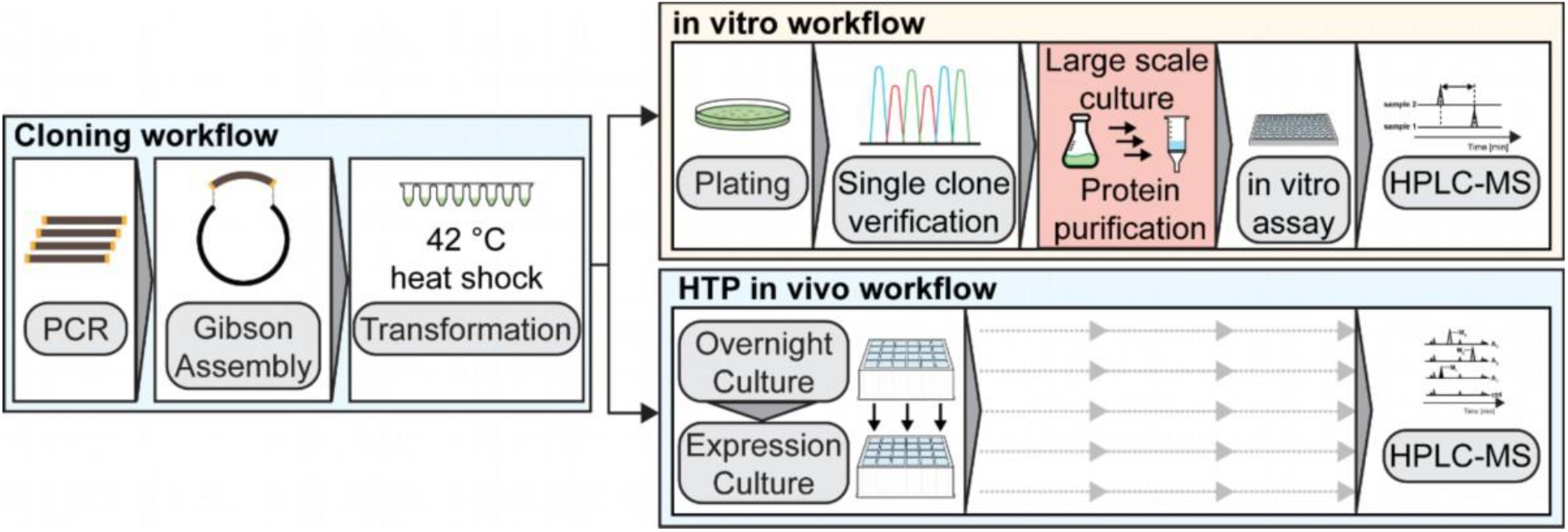
ASCR assay workflow overview. Schematic overview and comparison of an *in vitro* workflow and the novel described HTP *in vivo* ASCR workflow. The initial cloning procedure remains the same for both and can be performed in HTP. After transformation, both workflows differ significantly in workload, possible throughput and overall duration, especially when expression is performed in fast growing expression hosts like *Vibrio natriegens* (see Fig. S100).

## Conclusion

The accuracy of A domain prediction algorithms like PARAS depend on the availability of experimentally (structurally or biochemically) confirmed high quality data.^18^ Here, we describe a novel assay to determine the specificity of uncharacterized A domains *in vivo* by using NRPS engineering. We evaluated the applicability of the assay by screening A domains of 12 NRPS derived from various bacterial strains with a broad range of GC contents and taxonomic origins. Substrates with differing biophysical properties could be detected including non-proteinogenic amino acids, partially also derived from in *cis* acting enzymes like N-methyltransferases or oxidases. Even though, almost all A domains could be characterized, we observed some limitations for A domains with higher GC contents. Since the assay is based on heterologous expression in *E. coli*, absence of potential substrates of the A domain can be one explanation, which only can be circumvented by expression in the natural host or supplementation of the substrate. We are also aware that beside sufficient expression levels and correct folding, the generation of NRPS hybrids might be limiting since domain interactions must be functional to guarantee the assay to work.

Nevertheless, we also want to emphasize the advantages of the assay. In comparison to previously applied *in vitro* assays, the A domains are exposed to diverse substrates under cellular and natural competitive conditions with diverse amino acid substrates present in the culture. Furthermore, the A domain specificity is evaluated within the context of a full NRPS assembly line with upstream and downstream enzyme interactions, even though downstream interactions are artificially introduced through hybrid NRPS formation. This allows not only the determination of the A domain specificity but also its loading onto the T domain. In addition, accessory genes required for substrate modification are easily accessible through co-expression. The assay is suitable for high-throughput application with a minimal workload after cloning with low volume expression cultures that can be directly extracted and measured via HPLC-MS (Fig. 5). In comparison to high-throughput workflows for *in vitro* assays in a 96-well format, the active workload and required materials of to this assay are significantly reduced.^27,54^ Despite their overall workload (protein expression & purification), *in vitro* assays can in principle be performed faster than our *in vivo* ASCR assay which usually requires 72 hours of cultivation prior to HPLC-MS analysis. However, we could also significantly reduce this cultivation time to just 14 hours by applying *Vibrio natriegens* ATCC14048 Δ*dns*^55^ as expression strain (Fig. 100).

In conclusion, the ASCR assay successfully established a new *in vivo* high-throughput approach for determination of A domain specificities, which is fast and easy to perform for clarification of NRPS biosynthesis via HPLC-MS. Its broad application to several A domains with unknown substrate specificities even with the current state-of-the-art prediction tools like PARAS will also help to improve such bioinformatic tools in the future and identify NRPS incorporating rare and unusual non-canonical amino acids.

## Acknowledgements

Work in the Bode lab was supported by the Max-Planck Society. The authors are grateful to T.M. Mohiuddin for generation of plasmid pTH3 and Dr. Ismath Sadhir for providing template plasmids containing fluorescent proteins.

## Author contributions

J.L. generated all promoter exchange mutants. J.L. and Y.Z. performed experiments for structural elucidation of promoter exchange strain derived products. Y.Z. and A.B.W. analyzed all NMR data. O.N.L.S. performed experiments in *Vibrio natriegens*. L.P. generated the remaining data, wrote the manuscript, analyzed the data and generated all illustrations.

## Table of Content graphic

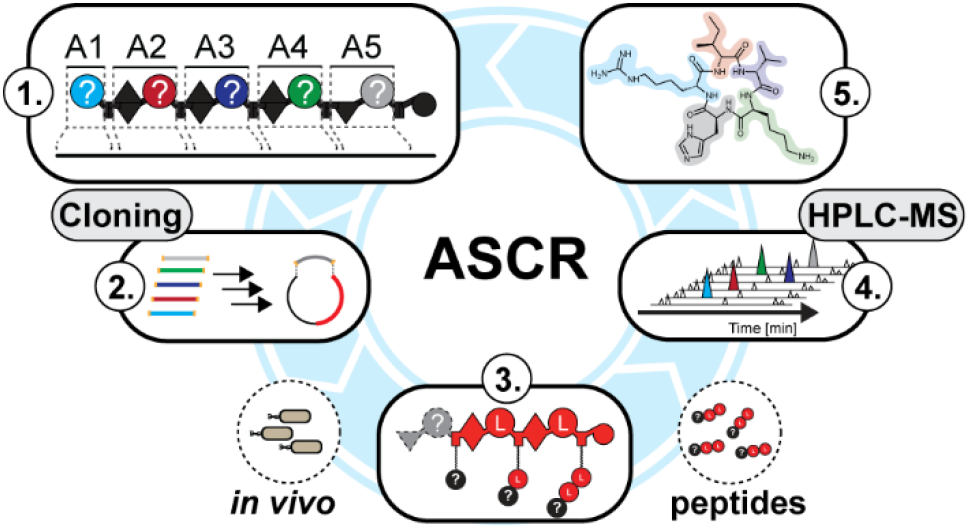

