## Supplementary Methods, Tables & Figures for "Development of a High-throughput *in vivo* Assay for the Determination of Adenylation Domain Specificities"

### 1 Materials and Methods

#### 1.1 Cultivation of *E. coli* DH10B:*mtaA* strains

All *E. coli* DH10B::*mtaA* were cultured either on liquid or solid low salt LB medium (pH 7.5, 10 g/l tryptone, 5 g/l yeast extract and 5 g/l NaCl). Solid medium was prepared with 1% agar (w/v). Depending on the experiment, gentamicin (15 µg/ml), chloramphenicol (34 µg/ml) and kanamycin (50 µg/ml) was added for selection. Cells were cultivated at 37 °C while shaking at 180 rpm.

#### 1.2 Cloning of vectors

Cloning was performed as described in Bozhüyük et al. 2024.[1] Genomic DNA (gDNA) was either isolated via Monarch® Genomic DNA Purification Kit (NEB) from the respective strains depicted in table S1 or was obtained from DSMZ. gDNA or plasmid DNA was used as template in PCR to generate plasmids listed in table S2. Q5® High-Fidelity DNA Polymerase (NEB) or Phusion® Hot Start Flex DNA Polymerase (NEB) were utilized to generate all DNA fragments in this work. All primer pair sequences and product sizes of the amplified fragments are documented in table S3. The PCR amplified fragments were purified by gel extraction from 1% (w/v) agarose gels taking the Monarch® DNA Gel Extraction Kit (NEB). If plasmid DNA was used as a template, the PCR mixture was digested with DpnI before gel extraction. Gibson cloning employing NEBuilder® HiFi DNA Assembly Cloning Kit (NEB) assembled the plasmid, which was subsequently transformed into chemical competent *E. coli* DH10B::*mtaA* (see section 1.4 and 1.5). Instead of plating only 15 µl as described in section 1.5, all cells were plated after transformation. Only pASCR_empty and pASCR8_empty were assembled using the KLD Enzyme Mix (NEB) as described by the manufactures instructions. Finally, a single colony was used to inoculate 3 ml low salt LB medium with respective antibiotics and was grown overnight at 37 °C and 180 rpm shaking. Plasmids were isolated by the PureYield™ Plasmid Miniprep System (Promega) and were further verified by oxford nanopore sequencing (ONT) by Microsynth Seqlab GmbH.

#### 1.3 Cloning of A domains or C-A didomains into pASCR vectors

Restriction digestion for pASCR2, pASCR3, pASCR8 and pASCR12 was performed using BstZ17I-HF (NEB), while BsaI-HF (NEB) was used for pASCR4. Restriction digestion was performed as described by the manufacturer. The mixture was incubated for 1 hour at 37 °C. Subsequently, DNA was purified using the Monarch Spin PCR & DNA Cleanup Kit (NEB) and the DNA concentration was adjusted to 40 ng/µl. Genes encoding for A domains or C-A didomains were amplified as described in section 1.2 from gDNA of strains listed in table S1 with the primer pairs as listed in table S4. For amplifications of genes with GC content above 60%, Phusion® Hot Start Flex DNA Polymerase (NEB) with Phusion® GC Reaction Buffer (NEB) in combination with Q5® GC Enhancer (NEB) was used. The primer were designed to amplify genes from the beginning of the antiSMASH[2] annotated C domain (or A domain if no C domain was present) and ended at the XUT[I] assembly site before the T domain.[1] Primer starting at the A or C domain of encoded elongation modules contained within the primer overhang the ATG start codon. All fragments were purified by gel extraction from 1% (w/v) agarose gel utilizing the Monarch® DNA Gel Extraction Kit (NEB) and the concentration was adjusted to 40 to 100 ng/µl. Plasmid and insert were mixed in a 1:1 (v/v) ratio. Finally, Gibson assembly was performed using NEBuilder® HiFi DNA Assembly Cloning Kit (NEB) with a final volume of 3 µl, which was incubated for 1 hour at 50 °C. 1 µl was subsequently transformed into chemical competent *E. coli* DH10B::*mtaA* (see section 1.4 and 1.5). For each plasmid, two colonies were tested via colony PCR to verify gene fragment insertion into the cloning vector (see section 1.6). A single colony was used to inoculate 3 ml low salt LB medium with respective antibiotics and was grown overnight at 37 °C. Plasmids were isolated by the PureYield™ Plasmid Miniprep System (Promega). All plasmids related to pASCR3 were further verified by oxford nanopore sequencing (ONT) by Microsynth Seqlab GmbH. For all plasmids related to pASCR8, the inserts were partially sequenced by Sanger sequencing by Microsynth Seqlab GmbH to verify the gene identity of the insert.

##### 1.4 Generation of chemical competent *E. coli* DH10B::*mtaA* cells

Chemical competent cells were prepared as described in Podolski et al. 2025.[3] Low salt LB medium was inoculated with a single colony of *E. coli* DH10B::*mtaA* and was grown overnight at 37 °C and shaking at 180 rpm. Fresh low salt LB medium supplemented with 10 mM $MgCl_2$ and 10 mM $MgSO_4$ was inoculated with 2% of the overnight grown culture and was incubated at 37 °C and 180 rpm shaking until an $OD_{600}$ between 0.6 and 0.8 was obtained. The culture was placed on ice for 30 minutes. The culture was centrifuged for 8 minutes and 4 °C at 3000 g and the supernatant was subsequently discarded. The cell pellet was suspended in one-third of the initial culture volume of ice-cold RF1 solution (100 mM RbCl, 50 mM $MnCl_2$, 30 mM Potassium acetate, 10 mM $CaCl_2$ and 15 % glycerol). The suspension was placed on ice for 30 minutes and was subsequently centrifuged at 3000 g for 8 minutes and 4 °C. After discarding the supernatant, the pellet was suspended in 5% of the initial culture volume of ice-cold RF2 solution (10 mM MOPS, 10 mM RbCl, 75 mM $CaCl_2$ and 15% glycerol). After incubation for 30 minutes on ice, aliquots of the cell suspension were frozen in liquid nitrogen and stored at -80 °C.

##### 1.5 Transformation of chemical competent cells via Heat Shock.

Add 50-100 ng plasmid DNA to 20 µl chemical competent *E. coli* DH10B::*mtaA* in PCR tube (250 µl volume). The mixture was incubated for 10 minutes on ice. Subsequently, cells were incubated for 45 seconds at 42 °C and were immediately placed on ice for 2 minutes. 150 µl low salt LB medium was added to the cells. The cells were incubated for 1 hour at 37 °C with no shaking. Subsequently 15 µl were plated by 12-well drop gravity flow plating with 15 µl as described by Köbel et al. 2025[4].

#### 1.6 Colony PCR

Cells of a single colony were transferred to 20 µl DMSO:$H_2O$ (1:1) (v/v). The mixture was incubated for 5 min at 98 °C. After briefly mixing the mixture, 0.5 µl was used as a template in a 20 µl PCR reaction using the Phire Hot Start II Polymerase (Thermo Scientific) with the Thermo Scientific Phire Green Reaction Buffer and the verification primer pair (Forward-Primer: CCATAAGATTAGCGGATCC, Reverse-Primer: CTGGAATAGCGTTTGCAC).

#### 1.7 Heterologous expression of NRPS-encoding genes in the A domains specificity assay

Heterologous expression of pFP8 and pJW75/76 was described in Bozhüyük et al. 2024[1] while expression of pPC1020 was described in Präve et al. 2024[5].

A single colony was used to inoculate 3 ml low salt LB medium with respective antibiotics and was grown overnight at 37 °C and 180 rpm shaking. Subsequently, 3 ml XPP medium[6] with 0.02% L-arabinose, 0.2 % glucose and respective antibiotics was prepared in a 24 deep-well plate and inoculated with 1% overnight grown culture. For expression of hybrid *NRPS*, which were not generated with pASCR3 or pASCR8 no glucose was added to the medium. The medium was further supplemented with 1 mM *L*-homoarginine, 1 mM *L*-diaminobutyric acid or 1 mM *L*-ornithine when the analyzed NRPS contained A domains specific for one of these amino acids. The culture was incubated for 72 hours at 200 rpm and 22 °C. Finally, 100 µl culture was added to 400 µl methanol and was shaken at 1500 rpm for 15 minutes at room temperature. The extract was centrifuged at 17000 g for 30 minutes at 22 °C and supernatant was analyzed by HPLC-MS (see section 1.9).

Inverse feeding experiments were performed as previously described.[7,8] The strains were cultivated overnight as described above. Subsequently, 1% overnight grown culture was used to inoculate 3 ml ISOGRO$^{TM}$-$^{13}$C medium (Sigma Aldrich, prepared according to manufactures instructions) containing respective antibiotics and 0.02% L-arabinose. 1 mM of non-isotope amino acids was added to verify its incorporation into the final peptide. No amino acid supplementation served as reference. Cultivation and extraction was performed as describe above. MS spectra at indicated retention times were compared to determine the present mass shifts by incorporation of non-isotope amino acids.

#### 1.8 High-throughput assembly and expression A domains specificity assay protocol

Cloning of vectors was performed as described in previous section 1.3. However, instead of gel purification of all amplified gene fragments encoding for A domains or C-A didomains the Monarch Spin PCR & DNA Cleanup Kit (NEB) was used. After Gibson assembly, 1 µl of the mixture was transformed into chemical competent *E. coli* DH10B::*mtaA* (see section 1.4 and 1.5). Instead of plating the cells on low salt LB-agar plates, 100 µl of the cell mixture was used to inoculate 3 ml low salt LB medium with respective antibiotics and was grown overnight at 37 °C and 180 rpm shaking. Finally, the cell culture was used to inoculate medium for heterologous expression of the NRPS-encoding genes as described in section 1.7.

##### 1.9 HPLC-MS analysis

Samples analyzed by high-performance liquid chromatography mass spectrometry (HPLC-MS) were prepared as described in section 1.7. All HPLC-MS analysis of samples not generated by pASCR cloning vectors were performed as described in Bozhüyük et al. 2024[1]. All samples related to pASCR cloning vectors and promotor exchanges mutants (see section 1.11) were measured by high-resolution mass spectrometry. The samples were separated by liquid chromatography utilizing Bruker Elute UPLC system with a C18 column (ACQUITY UPLCTM BEH, 130 Å, 2.1 mm x 100 mm, Waters) at 40 °C. For each sample 2 µl was injected and separated for 20 minutes and a flow rate of 0.4 ml/min using an acetonitrile/water gradient with 0.1 % formic acid (v/v) (0-2 minutes of an isocratic 5% acetonitrile flow; a gradient from 5% to 95% acetonitrile between 2-14 minutes; a gradient from 14-15 minutes from 95% to 100% acetonitrile, an isocratic flow of 100% acetonitrile between 15-17.80 minutes and between 17.80 – 20 minutes an equilibration to 5% acetonitrile). Internal calibration was performed by injection of a 10 mM sodium formate solution (water:isopropanol, 1:1) with a flow rate of 0.4 ml/min between 16-16.3 minutes. Mass spectrometric data was obtained between 0 and 14 minutes by a VIP-HESI (vacuum insulated probe heated-electrospray ionization) quadrupole time-of-flight (qTOF) mass spectrometer (timsTOF fleX MALDI-2, Bruker Daltonics). The MS spectra was recorded in positive ion mode over a mass range of 100-2000 m/z and was subsequently analyzed by DataAnalysis 6.1.

For quantitative comparison of domains from PTEMK122_08040 cloned into pASCR2, pASCR3 and pASCR4 as well as comparison of *grsAB* cloned into pASCR3 and pASCR8 mass spectrometry was performed on amaZon speed system. The samples were separated by liquid chromatography utilizing Agilent 1290 Infinity II UPLC system using a C18-column (ACQUITY UPLCTM BEH, 130 Å, 2.1 mm x 100 mm, Waters) at 40 °C. For each sample 5 µl was injected and separated for 20 minutes and a flow rate of 0.4 ml/min using an acetonitrile/water gradient with 0.1 % formic acid (v/v) (0-2 minutes of an isocratic 5% acetonitrile flow; a gradient from 5% to 95% acetonitrile between 2-14 minutes; a gradient from 14-15 minutes from 95% to 100% acetonitrile, an isocratic flow of 100% acetonitrile between 15-18 minutes and between 18-20 minutes an equilibration to 5% acetonitrile). Mass spectrometric data was obtained between 1 and 15 minutes by a ESI (electrospray ionization) ion trap spectrometer (amaZon speed, Bruker Daltonics). The MS spectra was recorded in positive ion mode over a mass range of 100-1200 m/z and was subsequently analyzed by DataAnalysis 6.1.

#### 1.10 Determination of expression and growth

A single colony was used to inoculate 3 ml low salt LB medium with respective antibiotics and was grown overnight at 37 °C and 180 rpm shaking. For each gene fragment from PTEMK122_08040 cloned into pASCR2, pASCR3 and pASCR4, 98 µl XPP medium[6] with 0.02% L-arabinose and respective antibiotics was prepared in a 96-well plate. The medium was inoculated with 2% overnight grown culture. If mentioned, 0.2% glucose was added to the medium. For all other analysis, 99 µl XPP medium with 0.02% L-arabinose, 0.2% glucose and respective antibiotics was prepared in a 96-well plate. The mediu was inoculated with 1% overnight grown culture. Fluorescence and growth was obtained by Tecan Spark® system. Growth was measured by determination of the $OD_{600}$. Fluorescence of msfGFP for pASCR3 derived constructs was determined with Excitation at 480 nm and Emission at 525 nm. Fluorescence of mScarlet-I3 for pASCR8 derived plasmids was determined at Excitation 560 nm and Emission of 605 nm. For both the gain was set to 80 and Z-position to 27000 µm, 30 flashes and an integration time of 40 µs was used. The plate was continuously shaken with an amplitude of 2.5 mm and a frequency of 630 rpm. Data points were obtained in 20 minute intervals for 72 hours at 22 °C. For domains from PTEMK122_08040 cloned into pASCR2, pASCR3 and pASCR4 as well as comparison of *grsAB* cloned into pASCR3 and pASCR8, all samples were determined in biological triplicates, while other samples were determined in a single measurement.

#### 1.11 Promoter exchange and expression in *P. temperata* K122

To obtain $P_{BAD}$ promoter exchange strains of *P. temperata* K122 for the genes PTEMK122_08040, PTEMK122_08690 and PTEMK122_21930, a protocol established by Bode et al. 2015 was used.[9] First, the three pCEP vectors pJL09, pJL05 and pJL14 were cloned as described in section 1.2 using the primer pairs listed in table S3. Next, the verified plasmids were transformed via electroporation into *E. coli* ST18 for subsequent conjugation into *P. temperata* K122 (see section 1.12). For plasmid conjugation, 5 ml low salt LB medium was inoculated with 2% overnight grown *P. temperata* K122 and was grown at 30 °C and 200 rpm to an $OD_{600}$ of 0.6 to 0.8. In parallel, 5 ml low salt LB medium supplemented with 5-aminolevulinic acid (ALA) (50µg/ml) was inoculated with 1% overnight grown *E. coli* ST18 containing the respective plasmid. The strain was grown at 37 °C and 200 rpm to an $OD_{600}$ of 0.6 to 0.8. 1 ml of each culture was centrifuged at 11000 rpm for 1 minute. *E. coli* ST18 cells were washed 3 times with 1 ml low salt LB medium supplemented with ALA to remove the antibiotics. Finally, each cell pellet was resuspended in 400 µl low salt LB medium. The two strains were mixed in 3:1 ratio (*E. coli* ST18 donor: *P. temperata* K122) and were plated on low salt LB-agar plates. After incubation for 3 hours at 37 °C, the plate was transferred to 30 °C and was incubated overnight. The cells were transferred into 1 ml low salt LB medium. 500 µl of the mixture was transferred to a low salt LB agar plate containing 50 µg/mL kanamycin and was grown at 30 °C until recombinant *P. temperata* K122 were visible. For obtained colonies, the correct insertion of the promoter exchange plasmid was verified by colony PCR utilizing the primer pairs (v_pCEP_fw = GCTATGCCATAGCATTTTTATCCATAAG and v_pCEP_rv = ACATGTGGAATTGTGAGCGG).

The verified *P. temperata* K122 $P_{BAD}$ promoter exchange strain was cultivated overnight in 5 ml low salt LB medium containing 50 µg/mL kanamycin at 30 °C and 180 rpm shaking. 1 % overnight culture was used to inoculate 5 ml low salt LB (*P. temperata* K122/JL09 and *P. temperata* K122/JL14) or XPP medium (*P. temperata* K122/JL05) supplemented with 50 µg/ml kanamycin and 0.2% L-arabinose. The culture was incubated for 72 hours at 30 °C while shaking at 200 rpm. For metabolic analysis, 50 µl culture was mixed with 500 µl methanol. The mixture was centrifuged for 30 minutes at 22 °C and the supernatant was analyzed as described in section 1.9.

For structure elucidation of detected compounds **1** to **11** via HPLC-MS, $^{13}C$ and $^{15}N$ isotope labelling experiments were performed with the *P. temperata* K122 $P_{BAD}$ promoter exchange strains as described previously.[7,8] The strains were cultivated overnight as described above. Subsequently, 1% overnight grown culture was used to inoculate 5 ml ISOGRO™-$^{13}C$ medium, ISOGRO™-$^{15}N$ medium (Sigma Aldrich, prepared according to manufactures instructions) and non-labelled medium (low salt LB medium or XPP medium) containing respective antibiotics and 0.2% L-arabinose, respectively. Cultivation and extraction was performed as describe above. MS spectra at indicated retention times were compared to determine the present carbon and nitrogen atoms by the isotope mass shifts. Furthermore, inverse feeding experiments were performed in ISOGRO™-$^{13}C$ medium by supplementation of 1 mM of non-isotope labelled amino acids to verify its incorporation into the final peptide by MS spectra mass shift.

##### 1.12 Generation of electro competent *E. coli* ST18 and transformation by electroporation

*E. coli* ST18 used for conjugation were transformed with plasmid DNA via electroporation.[10] Low salt LB medium with 5-aminolevulinic acid (ALA) (50µg/ml) was inoculated with *E. coli* ST18 and was grown overnight at 37 °C while shaking at 180 rpm. Fresh low salt LB medium supplemented with ALA was inoculated with 1% overnight grown culture and was incubated at 37 °C until an $OD_{600}$ between 0.8 - 1.0 was obtained. The cell culture was centrifuged at 4000 g and 4 °C for 15 minutes. The supernatant was discarded and the cell pellet was washed with 4/5 culture volume of ice-cold 10% glycerol. The washing step was repeated twice with 1/25 and 1/50 culture volume of ice-cold 10% glycerol. Finally, the cells were suspended in 1/500 culture volume of ice-cold 10% glycerol and 50 µl aliquots were stored at -80 °C. For transformation via electroporation, 50-100 ng of plasmid DNA was added to the cells and the mixture was transferred into an ice-cold 1 mm gap width cuvette. Electroporation was performed at 25 µF, 200 Ω, 1250 V and 1 pulse. The cell mixture was resuspended in 1 ml low salt LB medium supplemented with ALA and was incubated for 1 hour at 37 °C while shaking. Finally, the cell mixture was plated on low salt LB agar plates containing 50 µg/mL kanamycin and ALA and was incubated overnight at 37 °C.

##### 1.13 Purification of natural products and structure determination

For purification of natural products produced by *P. temperata* K122 $P_{BAD}$ promoter exchange strains *P. temperata* K122/JL09 (**1**), *P. temperata* K122/JL05 (**4** and **5**) and *P. temperata* K122/JL14 (**7a** and **7b**), 5 L Gibco™ Sf-900™ SFM II medium supplemented with 50 µg/ml kanamycin, 0.2% L-arabinose and 4% Amberlite® XAD-16 resin was inoculated with 1% overnight grown culture and was cultivated at 30 °C and 120 rpm for 72 hours. The separated XAD-16 resin was extracted 3 times with methanol and the solvent was subsequently evaporated to dryness. The extracts were dissolved in acetonitrile:water (1:1 ratio) and purified by preparative HPLC/MS (LC-MS-System 1260 Infinity II Preparative LC/MSD from Agilent) and semipreperative HPLC/MS (Agilent LC-MS-System 1260 Infinity II Analytical-Scale LC/MSD). Compound **1** was purified from the extract utilizing on the preparative HPLC system an Agilent 10 Prep-C18 (250 x 30.0 mm) column with a flow rate of 40 ml/min and an acetonitrile/water gradient from 5% to 50% (v/v) for 15 minutes. Subsequently, the extract was further purified by using a semipreparative HPLC system with a Cholester column (10 mm ID x 250 mm) a flow rate of 3 ml/min and an acetonitrile/water gradient from 19% to 20.8% (v/v) for 21 minutes. Compound **5** and **6** were purified from the same extract utilizing on the preparative HPLC system an Agilent 10 Prep-C18 (250 x 30.0 mm) column with a flow rate of 40 ml/min and an acetonitrile/water gradient from 5% to 25% (v/v) for 13 minutes. Fractions containing **5** were further purified by semipreparative HPLC system using an Eclipse XDB-C18 column with an isocratic flow of 20 ml/min with 6.8% acetonitrile in water for 25 minutes. Compound **7a** and **7b** were purified from the same extract utilizing on the preparative HPLC system an Agilent 10 Prep-C18 (250 x 30.0 mm) column with a flow rate of 40 ml/min and an acetonitrile/water gradient from 5% to 50% (v/v) for 16 minutes. Both compounds were further purified by semipreparative HPLC system using a Cholester column (10 mm ID x 250 mm) with a flow rate of 3 ml/min and an isocratic flow of 20% acetonitrile for 30 minutes. All samples were freeze-dried and solved in DMSO-$d_6$ for structure determination by NMR (Bruker AV-500 NMR spectrometer) recording a $^{1}$H-NMR spectra, $^{13}$C-NMR spectra, $^{1}$H-$^{1}$H COSY, HSQC and

HMBC (see table S5-S9). A NOESY spectra was measured for compound **1**, **5** and **6** in addition.

##### 1.14 Advanced Marfey's method

For determination of the stereochemistry of a purified peptide the advanced Marfey's method was applied.[11] 0.5-1 mg peptide (respectively for **1**, **4**, **5**, **7a** and **7b**) were solved in 200 µl methanol and were added to 800 µl HCl (6 M) solution in an Ace high-pressure tube and was incubated for 1 hour at 110 °C. Subsequently, the solvent was evaporated in a speed-vac system at 60 °C and material was solved in 100 µl $H_2O$. 45 µl hydrolyzed peptide or amino acid standard (corresponding to 0.1 mg) was mixed with 10 µl 1 M $NaCO_3$ and 80 µl 1% L-FDLA (Nα-(5-fluoro-2,4-dinitrophenyl)-L-leucinamide) or D-FDLA solution solved in acetone, respectively. The mixture was incubated at 40 °C for 1 hour under exclusion of light. The reaction was stopped upon addition of 10 µl 1M HCl. The solvent was evaporated in a speed-vac system at room temperature for 1 hour. The sample was solved in 400 µl methanol. For measurement on HPLC-MS systems the sample was centrifuged for 30 minutes at 17000 g and supernatant was transferred for measurement. For measurement of L-FDLA and D-FDLA derivatized samples, the individual reactions were mixed in a 1:1 ratio and were subsequently prepared for HPLC-MS on amaZon speed system (see section 1.9).

##### 1.15 Transformation via electroporation and heterologous expression of NRPS in *Vibrio natriegens*

Electrocompetent cells of *Vibrio natriegens* ATCC14048 Δ*dns*[12] were prepared following a modified version of established protocols.[13] A 5 ml culture of LB medium (pH 7.5, 10 g/l tryptone, 5 g/l yeast extract and 10 g/l NaCl) supplemented with V2 salts (10x stock solution: 2.04 M NaCl, 0.042 M KCl, and 0.2314 M $MgCl_2 \cdot 6H_2O$) was inoculated with *Vibrio natriegens* ATCC14048 Δ*dns* and was incubated in a 50 ml Erlenmeyer flask at 28 °C while shaking at 220 rpm overnight. The next day, 100 ml of the same medium was inoculated with 0.5% of overnight grown culture in a 500 ml Erlenmeyer flask and grown at 37 °C and 220 rpm until the $OD_{600}$ reached approximately 0.5. The culture was incubated on ice for 15 min and subsequently divided into four 25 ml aliquots in 50 mL Falcon tubes. Cells were pelleted by centrifugation at 6500 rpm for 20 min at 4 °C. The supernatant was discarded, and the cell pellet in each tube was re-suspended in 5 ml of ice-cold electroporation buffer (680 mM sucrose, 7 mM $K_2HPO_4$, pH 7.0), pooled into a single 50 ml Falcon tube, and diluted to a final volume of 30 ml with additional buffer. The suspension was mixed gently by inverting several times. Cells were pelleted again at 6500 rpm for 15 min at 4 °C, and the wash step was repeated three times in total. Finally, the cells were suspended in the electroporation buffer to a final $OD_{600}$ of ~16. 50 µL aliquots were frozen in liquid nitrogen and stored at -80 °C.

2 µL plasmid DNA (50–100 ng/µL) was added to the cells and was gently mixed. The mixture was transferred to an ice-cold electroporation cuvette with a 0.1 mm gap width. Electroporation was performed at 700 V, 25 µF, and 200 Ω. The cells were recovered in 1 ml of LB medium supplemented with V2 salts and were incubated for 2 hours at 37 °C with 800 rpm shaking. Following recovery, the culture was pelleted, re-suspended in 50 µL and then transferred to a 50 ml Erlenmeyer flask containing 5 ml of LB medium supplemented with V2 salts and 3.4 µg/mL chloramphenicol for selection. The culture was incubated at 28 °C overnight while shaking.

4 ml XPP medium supplemented with V2 salts, 3.4 µg/mL chloramphenicol, 0.2 % L-arabinose, 0.2 % glucose, and 1 mM stocks of required amino acids (e.g., ornithine) were inoculated with 1% of overnight grown culture obtained after electroporation. Cultures were grown in a 24-well deep-well plate at 28 °C and 250 rpm overnight. Finally, 100 µl culture was added to 400 µl methanol and was shaken at 1500 rpm for 15 minutes at room temperature. The extract was centrifuged at 17000 g for 30 minutes at 22 °C and supernatant was analyzed by HPLC-MS (see section 1.9).

#### 2 Supplementary Tables

**Table S1.** Strains used in this work.

| Strain | Genotype/NRPS | Reference | NCBI locus tags |
|---|---|---|---|
| *E. coli* DH10B | F_mcrA (*mrr-hsd*RMS-*mcr*BC), 80*lac*ZΔ, M15, Δ*lac*X74 *rec*A1 *end*A1 *ara*D 139Δ(*ara*, *leu*)7697 *gal*U *gal*K λ *rps*L (*Strr*) *nup*G / - | Invitrogen | - |
| *E. coli* DH10B::*mta*A | DH10B with *mtaA* from pCK_*mtaA*Δ*entD* / - | [14] | - |
| *E. coli* ST18 | *E. coli* S17-1 λpir *ΔhemA* | [15] | - |
| *Xenorhabdus stockiae* DSM 17904 | WT (*xabABC*[16]) | DMSZ | Xsto_RS05055 (*xabA*) XSTOV2_09090 (see supplementary data) |
| *Photorhabdus laumondii* TTO1 | WT (*gxpS*[8], *glbS*[17]) | DSMZ | PLU_RS16260 (*gxpS*), PLU_RS09375 (*glbC*), PLU_RS09365 (*glbE*) |
| *X. indica* DSM 17382 | WT (*xldS*[14]) | DSMZ | Xind_RS18785 (*xldS*) |
| *X. nematophila* ATCC 19061 | WT (*odlS*[18]) | ATCC | XNC1_RS10320 (*odlS1*), XNC1_RS10315 (*odlS2*), XNC1_RS10310 (*odlS3*), XNC1_RS10305 (*odlS4*), |
| *Photorhabdus temperata* K122 | WT | David Clarke Lab | PTEMK122_08040 (see supplementary data), PTEMK122_08690 (see supplementary data), PTEMK122_21930 & PTEMK122_21935 (see supplementary data) |
| *Aneurinibacillus migulanus* DSM 2895 | WT (*grsAB*[19]) | DSMZ | AF333_RS17650 (*grsA*), AF333_RS17645(*grsB*) |
| *Pseudomonas costantinii* DSM 16734 | WT | DSMZ | HX791_RS00015 |
| *Chromobacterium subtsugae* DSM 17043 | WT (*rdb-like*) | DSMZ | MY55_RS13030 (*rdbD*), MY55_RS13020 (*rdbF*), MY55_RS13005 (*rdbH*) |
| *Myxococcus xanthus* DK 1622 | WT | Lab collection | MXAN_RS13550 |
| *Xanthobacter autotrophicus* DSM 432 | WT | DSMZ | FBQ73_RS02155, FBQ73_RS02140 |
| *Goodfellowiella coeruleoviolacea* ATCC 53904 | WT (*epxS*[20]) | ATCC | *epxD, epxE* |
| *Photorhabdus temperata* K122/JL09 | WT conjugated with pJL09 | This work | PTEMK122_08040 |
| *Photorhabdus temperata* K122/JL05 | WT conjugated with pJL05 | This work | PTEMK122_08690 |
| *Photorhabdus temperata* K122/JL14 | WT conjugated with pJL14 | This work | PTEMK122_21930 |
| *Vibrio natriegens* ATCC14048 Δ*dns* | Δ*dns* | [12] | - |

**Table S2.** Plasmids and corresponding genotypes used in this work.

| Plasmids | Genotype | Reference |
|---|---|---|
| pACYC | ori p15A, $cm^R$, *araC*- $P_{BAD,}$ *tacI* and *araE* | [21] |
| pCOLA | ori ColA, $kan_R$, *araC*- $P_{BAD,}$ and *tacI* | [22] |
| pCEP | pDS132 based, R6K ori, oriT, $Km^R$, *araC*, $P_{BAD}$ | [9] |
| pACYC_Seva | $cm^R$, oriT, p15A ori, *araE*, *araC*, *araBAD* | iGEM Marburg 2025 (https://registry.igem.org/parts/bba-25oskput) |
| pCOLA_Seva | $Kan^R$, oriT, ColA ori, *araE*, *araC*, *araBAD* | iGEM Marburg 2025 (https://registry.igem.org/parts/bba-25u5gor0) |
| pJL09 | pCEP containing the first 625 bp of *PTEMK122_08040* | This work |
| pJL05 | pCEP containing the first 788 bp of *PTEMK122_08690* | This work |
| pJL14 | pCEP containing the first 718 bp of *PTEMK122_21930* | This work |
| pFP8 | ori ColA, $kan^R$, *araC*-$P_{BAD}$ *xabABC*_C1A1T1 (XUT$^{IV}$)-*gxpS*_T3CE4A4T4CE5A5T5TE and *tacI* | [1] |
| pLP179 | ori p15A, $cm^R$, *araC*-$P_{BAD}$ *xldS*_C1A1T1 (XUT$^{IV}$)-*gxpS*_T3CE4A4T4CE5A5T5TE *tacI* and *araE* | This work |
| pTH3 | ori ColA, $kan^R$, *araC*-$P_{BAD}$ *odlS*_C1A1T1 (XUT$^{III}$)-*gxpS*_T3CE4A4T4CE5A5T5TE and *tacI* | This work (Master Thesis T.M. Mohiuddin 'Engineering Enzymatic Assembly Lines') |
| pLP212 | ori p15A, $cm^R$, *araC*-$P_{BAD}$ *xldS*_C1A1T1CE2A2T2 (XUT$^{IV}$)-*gxpS*_T3CE4A4T4CE5A5T5TE *tacI* and *araE* | This work |
| pPC1020 | ori p15A, $cm^R$, *araC*-$P_{BAD}$ *odlS*_C1A1T1CE2A2T2 (XUT$^{IV}$)-*gxpS*_T3CE4A4T4CE5A5T5TE *tacI* and *araE* | [5] |
| pPC1016 | ori ColA, $kan^R$, *araC*-$P_{BAD}$ *odlE* and *tacI* | [5] |
| pPC1017 | ori ColA, $kan^R$, *araC*-$P_{BAD}$ *odlF* and *tacI* | [5] |
| pPC1018 | ori ColA, $kan^R$, *araC*-$P_{BAD}$ *odlB* and *tacI* | [5] |
| pJW75 | ori p15A, $cm^R$, *araC*-$P_{BAD}$ *gxpS*_A1T1CE2A2T2C3_SYNZIP17 *tacI* and *araE* | [5] |
| pJW76 | ori ColA, $kan^R$, *araC*-$P_{BAD}$ SYNZIP18 *gxpS*_A3T3CE4A4T4CE5A5T5TE and *tacI* | [5] |
| pLP228 | ori p15A, $cm^R$, *araC*-$P_{BAD}$ *odlS*_CE2A2T2 (XUT$^{IV}$)-*gxpS*_T3CE4A4T4CE5A5T5TE *tacI* and *araE* | This work |
| pLP236 | ori p15A, $cm^R$, *araC*-$P_{BAD}$ *odlS*_CE2A2T2(S1049A)C3T3 (XUT$^{IV}$)-*gxpS*_T3CE4A4T4CE5A5T5TE *tacI* and *araE* | This work |
| pLP229 | ori p15A, $cm^R$, *araC*-$P_{BAD}$ *odlS*_C4A3T4 (XUT$^{IV}$)-*gxpS*_T3CE4A4T4CE5A5T5TE *tacI* and *araE* | This work |
| pLP230 | ori p15A, $cm^R$, *araC*-$P_{BAD}$ *odlS*_C5A4T5 (XUT$^{IV}$)-*gxpS*_T3CE4A4T4CE5A5T5TE *tacI* and *araE* | This work |
| pLP237 | ori p15A, $cm^R$, *araC*-$P_{BAD}$ *odlS*_C5A4T5 (XUT$^{I}$)-*gxpS*_T3CE4A4T4CE5A5T5TE *tacI* and *araE* | This work |

| | | |
|---|---|---|
| pLP232 | ori p15A, $cm^R$, *araC-*$P_{BAD}$ *odlS*_CE6A5T6 (XUT$^{IV}$)-*gxpS*_T3CE4A4T4CE5A5T5TE *tacI* and *araE* | This work |
| pLP231 | ori p15A, $cm^R$, *araC-*$P_{BAD}$ *odlS*_C7A6T7 (XUT$^{IV}$)-*gxpS*_T3CE4A4T4CE5A5T5TE *tacI* and *araE* | This work |
| pPL238 | ori p15A, $cm^R$, *araC-*$P_{BAD}$ *odlS*_C7A6T7 (XUT$^{I}$)-*gxpS*_T3CE4A4T4CE5A5T5TE *tacI* and *araE* | This work |
| pLP233 | ori p15A, $cm^R$, *araC-*$P_{BAD}$ *odlS*_C8A7T8 (XUT$^{IV}$)-*gxpS*_T3CE4A4T4CE5A5T5TE *tacI* and *araE* | This work |
| pLP234 | ori p15A, $cm^R$, *araC-*$P_{BAD}$ *odlS*_C9A8T9 (XUT$^{IV}$)-*gxpS*_T3CE4A4T4CE5A5T5TE *ttacI* and *araE* | This work |
| pLP235 | ori p15A, $cm^R$, *araC-*$P_{BAD}$ *odlS*_C10A9T10 (XUT$^{IV}$)-*gxpS*_T3CE4A4T4CE5A5T5TE *tacI* and *araE* | This work |
| pLP239 | ori p15A, $cm^R$, *araC-*$P_{BAD}$ *odlS*_C10A9T10 (XUT$^{I}$)-*gxpS*_T3CE4A4T4CE5A5T5TE *tacI* and *araE* | This work |
| pSEVA661b-Neon | $gm^R$, oriT, ori p15A, *araC*, $P_{BAD}$ *riboJ, mNeonGreen* | [23] |
| pCIOX | $Kan^R$, pUC, T7 promotor, 8xHis-tag, *SUMO,* 6xHis | pCIOX was a gift from Andrea Mattevi (addgene plasmid #51300) |
| pISNP4 | p15A ori, $kan^R$, *msfGFP*, *mScarlet-I3* | Lab collection |
| pASCR2 | $gm^R$, oriT, ori p15A, *araC*, $P_{BAD}$ *riboJ* (BstZ17I) *gxpS*_T3CE4A4T4CE5A5T5TE G2315 (GGT→GGG) | This work |
| pASCR3 | $gm^R$, oriT, ori p15A, *araC*, $P_{BAD}$ *riboJ* (BstZ17I) *gxpS*_T3CE4A4T4CE5A5T5TE G2315 (GGT→GGG) GS-linker *msfGFP* | This work |
| pASCR4 | $gm^R$, oriT, ori p15A, *araC*, $P_{BAD}$ *riboJ SUMO*-tag (BsaI spacer) *gxpS*_T3CE4A4T4CE5A5T5TE G2315 (GGT→GGG) GS-linker *msfGFP* | This work |
| pASCR_empty | $gm^R$, oriT, ori p15A, *araC*, $P_{BAD}$ *riboJ* | This work |
| pASCR_msfGFP | $gm^R$, oriT, ori p15A, *araC*, $P_{BAD}$ *riboJ msfGFP* | This work |
| pASCR5 | $gm^R$, oriT, ori p15A, *araC*, $P_{BAD}$ *riboJ* *gxpS*_C3A3T3(S997A)CE4A4T4(S2073A)CE5A5T5(S3144A)TE G3274 (GGT→GGG) | This work |
| pASCR6 | $gm^R$, oriT, ori p15A, *araC*, $P_{BAD}$ *riboJ* *gxpS*_C3A3T3(S997A)CE4A4T4(S2073A)CE5A5T5(S3144A)TE G3274 (GGT→GGG) GS-linker *msfGFP* | This work |
| pASCR8 | $cm^R$, oriT, ori p15A, *araE, araC*, $P_{BAD}$ *riboJ* (BstZ17I) *gxpS*_T3CE4A4T4CE5A5T5TE G2315 (GGT→GGG) GS-linker *mScarlet-I3* | This work |
| pASCR10 | $cm^R$, oriT, ori p15A, *araE*, *araC*, $P_{BAD}$ *riboJ* *gxpS*_C3A3T3(S997A)CE4A4T4(S2073A)CE5A5T5(S3144A)TE G3274 (GGT→GGG) GS-linker *mScarlet-I3* | This work |
| pASCR11 | $cm^R$, oriT, ori p15A, *araE*, *araC*, $P_{BAD}$ *riboJ* *gxpS*_C3A3T3(S997A)CE4A4T4(S2073A)CE5A5T5(S3144A)TE G3274 (GGT→GGG) | This work |
| pASCR8_empty | $cm^R$, oriT, ori p15A, *araE*, *araC*, $P_{BAD}$ *riboJ* | This work |
| pASCR_mScarlet-I3 | $cm^R$, oriT, ori p15A, *araE*, *araC*, $P_{BAD}$ *riboJ* *mScarlet-I3* | This work |

| | | |
|---|---|---|
| pSEVA341 | *cm*$^R$, oriT, pRO1600/ColE1 | [24] |
| pASCR12 | *cm*$^R$, oriT, pRO1600/ColE1, *araE, araC*, $P_{BAD}$ *riboJ* (BstZ17I) *gxpS*_T3CE4A4T4CE5A5T5TE G2315 (GGT→GGG) GS-linker *mScarlet-I3* | This work |
| pASCR12_mScarlet-I3 | *cm*$^R$, oriT, pRO1600/ColE1, *araE, araC*, $P_{BAD}$ *riboJ mScarlet-I3* | This work |
| pASCR13 | *cm*$^R$, oriT, pRO1600/ColE1, *araE*, *araC*, $P_{BAD}$ *riboJ gxpS*_C3A3T3(S997A)CE4A4T4(S2073A)CE5A5T5(S3144A)TE G3274 (GGT→GGG) GS-linker *mScarlet-I3* | This work |
| pLP262 | *gm*$^R$, oriT, ori p15A, *araC*, $P_{BAD}$ *riboJ* PTEMK122_08040_A1 (XUT$^I$) *gxpS*_T3CE4A4T4CE5A5T5TE | This work |
| pLP263 | *gm*$^R$, oriT, ori p15A, *araC*, $P_{BAD}$ *riboJ* PTEMK122_08040_CE2A2 (XUT$^I$) *gxpS*_T3CE4A4T4CE5A5T5TE | This work |
| pLP264 | *gm*$^R$, oriT, ori p15A, *araC*, $P_{BAD}$ *riboJ* PTEMK122_08040_C3A3 (XUT$^I$) *gxpS*_T3CE4A4T4CE5A5T5TE | This work |
| pLP265 | *gm*$^R$, oriT, ori p15A, *araC*, $P_{BAD}$ *riboJ* PTEMK122_08040_CE4A4 (XUT$^I$) *gxpS*_T3CE4A4T4CE5A5T5TE | This work |
| pLP266 | *gm*$^R$, oriT, ori p15A, *araC*, $P_{BAD}$ *riboJ* PTEMK122_08040_CE5A5 (XUT$^I$) *gxpS*_T3CE4A4T4CE5A5T5TE | This work |
| pLP278 | *gm*$^R$, oriT, ori p15A, *araC*, $P_{BAD}$ *riboJ* PTEMK122_08040_C6A6 (XUT$^I$) *gxpS*_T3CE4A4T4CE5A5T5TE | This work |
| pLP256 | *gm*$^R$, oriT, ori p15A, *araC*, $P_{BAD}$ *riboJ* PTEMK122_08040_A1 (XUT$^I$) *gxpS*_T3CE4A4T4CE5A5T5TE GS-linker *msfGFP* | This work |
| pLP257 | *gm*$^R$, oriT, ori p15A, *araC*, $P_{BAD}$ *riboJ* PTEMK122_08040_CE2A2 (XUT$^I$) *gxpS*_T3CE4A4T4CE5A5T5TE GS-linker *msfGFP* | This work |
| pLP258 | *gm*$^R$, oriT, ori p15A, *araC*, $P_{BAD}$ *riboJ* PTEMK122_08040_C3A3 (XUT$^I$) *gxpS*_T3CE4A4T4CE5A5T5TE GS-linker *msfGFP* | This work |
| pLP259 | *gm*$^R$, oriT, ori p15A, *araC*, $P_{BAD}$ *riboJ* PTEMK122_08040_CE4A4 (XUT$^I$) *gxpS*_T3CE4A4T4CE5A5T5TE GS-linker *msfGFP* | This work |
| pLP260 | *gm*$^R$, oriT, ori p15A, *araC*, $P_{BAD}$ *riboJ* PTEMK122_08040_CE5A5 (XUT$^I$) *gxpS*_T3CE4A4T4CE5A5T5TE GS-linker *msfGFP* | This work |
| pLP261 | *gm*$^R$, oriT, ori p15A, *araC*, $P_{BAD}$ *riboJ* PTEMK122_08040_C6A6 (XUT$^I$) *gxpS*_T3CE4A4T4CE5A5T5TE GS-linker *msfGFP* | This work |
| pLP250 | *gm*$^R$, oriT, ori p15A, *araC*, $P_{BAD}$ *riboJ SUMO*-tag PTEMK122_08040_A1 (XUT$^I$) *gxpS*_T3CE4A4T4CE5A5T5TE GS-linker *msfGFP* | This work |
| pLP251 | *gm*$^R$, oriT, ori p15A, *araC*, $P_{BAD}$ *riboJ SUMO*-tag PTEMK122_08040_CE2A2 (XUT$^I$) *gxpS*_T3CE4A4T4CE5A5T5TE GS-linker *msfGFP* | This work |

| | | |
|---|---|---|
| pLP252 | *gm*$^{R}$, oriT, ori p15A, *araC*, $P_{BAD}$ *riboJ SUMO*-tag<br>PTEMK122_08040_C3A3 (XUT$^{I}$)<br>*gxpS*_T3CE4A4T4CE5A5T5TE GS-linker *msfGFP* | This work |
| pLP253 | *gm*$^{R}$, oriT, ori p15A, *araC*, $P_{BAD}$ *riboJ SUMO*-tag<br>PTEMK122_08040_CE4A4 (XUT$^{I}$)<br>*gxpS*_T3CE4A4T4CE5A5T5TE GS-linker *msfGFP* | This work |
| pLP254 | *gm*$^{R}$, oriT, ori p15A, *araC*, $P_{BAD}$ *riboJ SUMO*-tag<br>PTEMK122_08040_CE5A5 (XUT$^{I}$)<br>*gxpS*_T3CE4A4T4CE5A5T5TE GS-linker *msfGFP* | This work |
| pLP255 | *gm*$^{R}$, oriT, ori p15A, *araC*, $P_{BAD}$ *riboJ SUMO*-tag<br>PTEMK122_08040_C6A6 (XUT$^{I}$)<br>*gxpS*_T3CE4A4T4CE5A5T5TE GS-linker *msfGFP* | This work |
| pASCR3_D9 | *gm*$^{R}$, oriT, ori p15A, *araC*, $P_{BAD}$ *riboJ*<br>PTEMK122_08690_A1 (XUT$^{I}$)<br>*gxpS*_T3CE4A4T4CE5A5T5TE GS-linker *msfGFP* | This work |
| pASCR3_D10 | *gm*$^{R}$, oriT, ori p15A, *araC*, $P_{BAD}$ *riboJ*<br>PTEMK122_08690_CE2A2 (XUT$^{I}$)<br>*gxpS*_T3CE4A4T4CE5A5T5TE GS-linker *msfGFP* | This work |
| pASCR3_D11 | *gm*$^{R}$, oriT, ori p15A, *araC*, $P_{BAD}$ *riboJ*<br>PTEMK122_08690_CE3A3 (XUT$^{I}$)<br>*gxpS*_T3CE4A4T4CE5A5T5TE GS-linker *msfGFP* | This work |
| pASCR3_D12 | *gm*$^{R}$, oriT, ori p15A, *araC*, $P_{BAD}$ *riboJ*<br>PTEMK122_08690_CE4A4 (XUT$^{I}$)<br>*gxpS*_T3CE4A4T4CE5A5T5TE GS-linker *msfGFP* | This work |
| pASCR3_E1 | *gm*$^{R}$, oriT, ori p15A, *araC*, $P_{BAD}$ *riboJ*<br>PTEMK122_08690_C5A5 (XUT$^{I}$)<br>*gxpS*_T3CE4A4T4CE5A5T5TE GS-linker *msfGFP* | This work |
| pASCR3_E2 | *gm*$^{R}$, oriT, ori p15A, *araC*, $P_{BAD}$ *riboJ*<br>PTEMK122_21930_C1A1 (XUT$^{I}$)<br>*gxpS*_T3CE4A4T4CE5A5T5TE GS-linker *msfGFP* | This work |
| pASCR3_E3 | *gm*$^{R}$, oriT, ori p15A, *araC*, $P_{BAD}$ *riboJ*<br>PTEMK122_21930_CE2A2 (XUT$^{I}$)<br>*gxpS*_T3CE4A4T4CE5A5T5TE GS-linker *msfGFP* | This work |
| pASCR3_E4 | *gm*$^{R}$, oriT, ori p15A, *araC*, $P_{BAD}$ *riboJ*<br>PTEMK122_21935_C3A3 (XUT$^{I}$)<br>*gxpS*_T3CE4A4T4CE5A5T5TE GS-linker *msfGFP* | This work |
| pASCR3_E5 | *gm*$^{R}$, oriT, ori p15A, *araC*, $P_{BAD}$ *riboJ*<br>PTEMK122_21935_CE4A4 (XUT$^{I}$)<br>*gxpS*_T3CE4A4T4CE5A5T5TE GS-linker *msfGFP* | This work |
| pASCR3_E6 | *gm*$^{R}$, oriT, ori p15A, *araC*, $P_{BAD}$ *riboJ*<br>PTEMK122_21935_C5A5 (XUT$^{I}$)<br>*gxpS*_T3CE4A4T4CE5A5T5TE GS-linker *msfGFP* | This work |
| pASCR3_Xsto_M1 | *gm*$^{R}$, oriT, ori p15A, *araC*, $P_{BAD}$ *riboJ*<br>XSTOV2_09090_A1nMT (XUT$^{I}$)<br>*gxpS*_T3CE4A4T4CE5A5T5TE GS-linker *msfGFP* | This work |
| pASCR3_Xsto_M2 | *gm*$^{R}$, oriT, ori p15A, *araC*, $P_{BAD}$ *riboJ*<br>XSTOV2_09090_C2A2 (XUT$^{I}$)<br>*gxpS*_T3CE4A4T4CE5A5T5TE GS-linker *msfGFP* | This work |
| pASCR3_Xsto_M3 | *gm*$^{R}$, oriT, ori p15A, *araC*, $P_{BAD}$ *riboJ*<br>XSTOV2_09090_CE3A3 (XUT$^{I}$)<br>*gxpS*_T3CE4A4T4CE5A5T5TE GS-linker *msfGFP* | This work |
| pLP285 | *cm*$^{R}$, oriT, ori p15A, *araE*, *araC*, araBAD<br>XSTOV2_09090_09085 | This work |

| | | |
|---|---|---|
| pLP286 | *kan*$^R$, oriT, ori ColA, *araE*, *araC*, araBAD 09085 | This work |
| pASCR3_A1 | *gm*$^R$, oriT, ori p15A, *araC*, $P_{BAD}$ *riboJ grsA*_A1 (XUT$^I$) *gxpS*_T3CE4A4T4CE5A5T5TE GS-linker *msfGFP* | This work |
| pASCR3_A2 | *gm*$^R$, oriT, ori p15A, *araC*, $P_{BAD}$ *riboJ grsB*_C2A2 (XUT$^I$) *gxpS*_T3CE4A4T4CE5A5T5TE GS-linker *msfGFP* | This work |
| pASCR3_A3 | *gm*$^R$, oriT, ori p15A, *araC*, $P_{BAD}$ *riboJ grsB*_C3A3 (XUT$^I$) *gxpS*_T3CE4A4T4CE5A5T5TE GS-linker *msfGFP* | This work |
| pASCR3_A5 | *gm*$^R$, oriT, ori p15A, *araC*, $P_{BAD}$ *riboJ grsB*_C4A4 (XUT$^I$) *gxpS*_T3CE4A4T4CE5A5T5TE GS-linker *msfGFP* | This work |
| pASCR3_A5 | *gm*$^R$, oriT, ori p15A, *araC*, $P_{BAD}$ *riboJ grsB*_C5A5 (XUT$^I$) *gxpS*_ GS-linker *msfGFP* | This work |
| pASCR3_*odlS*_M1 | *gm*$^R$, oriT, ori p15A, *araC*, $P_{BAD}$ *riboJ odlS1*_C1A1 (XUT$^I$) *gxpS*_T3CE4A4T4CE5A5T5TE GS-linker *msfGFP* | This work |
| pASCR3_*odlS*_M2 | *gm*$^R$, oriT, ori p15A, *araC*, $P_{BAD}$ *riboJ odlS1*_CE2A2 (XUT$^I$) *gxpS*_T3CE4A4T4CE5A5T5TE GS-linker *msfGFP* | This work |
| pASCR3_*odlS*_M4 | *gm*$^R$, oriT, ori p15A, *araC*, $P_{BAD}$ *riboJ odlS2*_C4A3 (XUT$^I$) *gxpS*_T3CE4A4T4CE5A5T5TE GS-linker *msfGFP* | This work |
| pASCR3_*odlS*_M5 | *gm*$^R$, oriT, ori p15A, *araC*, $P_{BAD}$ *riboJ odlS2*_C5A4 (XUT$^I$) *gxpS*_T3CE4A4T4CE5A5T5TE GS-linker *msfGFP* | This work |
| pASCR3_*odlS*_M6 | *gm*$^R$, oriT, ori p15A, *araC*, $P_{BAD}$ *riboJ odlS2*_CE6A5 (XUT$^I$) *gxpS*_T3CE4A4T4CE5A5T5TE GS-linker *msfGFP* | This work |
| pASCR3_*odlS*_M7 | *gm*$^R$, oriT, ori p15A, *araC*, $P_{BAD}$ *riboJ odlS3*_C7A6 (XUT$^I$) *gxpS*_T3CE4A4T4CE5A5T5TE GS-linker *msfGFP* | This work |
| pASCR3_*odlS*_M8 | *gm*$^R$, oriT, ori p15A, *araC*, $P_{BAD}$ *riboJ odlS4*_C8A7 (XUT$^I$) *gxpS*_T3CE4A4T4CE5A5T5TE GS-linker *msfGFP* | This work |
| pASCR3_*odlS*_M9 | *gm*$^R$, oriT, ori p15A, *araC*, $P_{BAD}$ *riboJ odlS4*_C9A8 (XUT$^I$) *gxpS*_T3CE4A4T4CE5A5T5TE GS-linker *msfGFP* | This work |
| pASCR3_*odlS*_M10 | *gm*$^R$, oriT, ori p15A, *araC*, $P_{BAD}$ *riboJ odlS4*_C10A9 (XUT$^I$) *gxpS*_T3CE4A4T4CE5A5T5TE GS-linker *msfGFP* | This work |
| pASCR3_*glbS*_M1 | *gm*$^R$, oriT, ori p15A, *araC*, $P_{BAD}$ *riboJ glbE*_C1A1 (XUT$^I$) *gxpS*_T3CE4A4T4CE5A5T5TE GS-linker *msfGFP* | This work |
| pASCR3_*glbS*_M2 | *gm*$^R$, oriT, ori p15A, *araC*, $P_{BAD}$ *riboJ glbC*_C2A2 (XUT$^I$) *gxpS*_T3CE4A4T4CE5A5T5TE GS-linker *msfGFP* | This work |
| pASCR3_*glbS*_M3 | *gm*$^R$, oriT, ori p15A, *araC*, $P_{BAD}$ *riboJ glbC*_C3A3 (XUT$^I$) *gxpS*_T3CE4A4T4CE5A5T5TE GS-linker *msfGFP* | This work |

| | | |
|---|---|---|
| pASCR3_A6 | *gm*$^R$, oriT, ori p15A, *araC*, $P_{BAD}$ *riboJ* HX791_RS00015_CE1A1 (XUT$^I$) *gxpS*_T3CE4A4T4CE5A5T5TE GS-linker *msfGFP* | This work |
| pASCR3_A7 | *gm*$^R$, oriT, ori p15A, *araC*, $P_{BAD}$ *riboJ* HX791_RS00015_CE2A2 (XUT$^I$) *gxpS*_T3CE4A4T4CE5A5T5TE GS-linker *msfGFP* | This work |
| pASCR3_A8 | *gm*$^R$, oriT, ori p15A, *araC*, $P_{BAD}$ *riboJ* HX791_RS00015_CE3A3 (XUT$^I$) *gxpS*_T3CE4A4T4CE5A5T5TE GS-linker *msfGFP* | This work |
| pASCR3_A9 | *gm*$^R$, oriT, ori p15A, *araC*, $P_{BAD}$ *riboJ* HX791_RS00015_C4A4 (XUT$^I$) *gxpS*_T3CE4A4T4CE5A5T5TE GS-linker *msfGFP* | This work |
| pASCR3_A10 | *gm*$^R$, oriT, ori p15A, *araC*, $P_{BAD}$ *riboJ* HX791_RS00015_C5A5 (XUT$^I$) *gxpS*_T3CE4A4T4CE5A5T5TE GS-linker *msfGFP* | This work |
| pASCR3_A11 | *gm*$^R$, oriT, ori p15A, *araC*, $P_{BAD}$ *riboJ* HX791_RS00015_C6A6 (XUT$^I$) *gxpS*_T3CE4A4T4CE5A5T5TE GS-linker *msfGFP* | This work |
| pASCR3_B4 | *gm*$^R$, oriT, ori p15A, *araC*, $P_{BAD}$ *riboJ* MY55_13005_C1A1 (XUT$^I$) *gxpS*_T3CE4A4T4CE5A5T5TE GS-linker *msfGFP* | This work |
| pASCR3_B5 | *gm*$^R$, oriT, ori p15A, *araC*, $P_{BAD}$ *riboJ* MY55_12955_C2A2 (XUT$^I$) *gxpS*_T3CE4A4T4CE5A5T5TE GS-linker *msfGFP* | This work |
| pASCR3_B6 | *gm*$^R$, oriT, ori p15A, *araC*, $P_{BAD}$ *riboJ* MY55_12980_A3 (XUT$^I$) *gxpS*_T3CE4A4T4CE5A5T5TE GS-linker *msfGFP* | This work |
| pASCR3_B7 | *gm*$^R$, oriT, ori p15A, *araC*, $P_{BAD}$ *riboJ* MY55_12980_CE3A4 (XUT$^I$) *gxpS*_T3CE4A4T4CE5A5T5TE GS-linker *msfGFP* | This work |
| pASCR3_C12 | *gm*$^R$, oriT, ori p15A, *araC*, $P_{BAD}$ *riboJ* MXAN_RS13550_A-Ox1 (XUT$^I$) *gxpS*_T3CE4A4T4CE5A5T5TE GS-linker *msfGFP* | This work |
| pASCR3_F8 | *gm*$^R$, oriT, ori p15A, *araC*, $P_{BAD}$ *riboJ* FBQ73_RS02155_A1 (XUT$^I$) *gxpS*_T3CE4A4T4CE5A5T5TE GS-linker *msfGFP* | This work |
| pASCR3_F9 | *gm*$^R$, oriT, ori p15A, *araC*, $P_{BAD}$ *riboJ* FBQ73_RS02140_C1A2 (XUT$^I$) *gxpS*_T3CE4A4T4CE5A5T5TE GS-linker *msfGFP* | This work |
| pASCR3_F10 | *gm*$^R$, oriT, ori p15A, *araC*, $P_{BAD}$ *riboJ* FBQ73_RS02140_Cy2A3 (XUT$^I$) *gxpS*_T3CE4A4T4CE5A5T5TE GS-linker *msfGFP* | This work |
| pASCR3_epoxy_ M1 | *gm*$^R$, oriT, ori p15A, *araC*, $P_{BAD}$ *riboJ* *epxD*_C1A1nMT (XUT$^I$) *gxpS*_T3CE4A4T4CE5A5T5TE GS-linker *msfGFP* | This work |
| pASCR3_ epoxy _M2 | *gm*$^R$, oriT, ori p15A, *araC*, $P_{BAD}$ *riboJ* *epxD*_C2A2 (XUT$^I$) *gxpS*_T3CE4A4T4CE5A5T5TE GS-linker *msfGFP* | This work |
| pASCR3_ epoxy _M3 | *gm*$^R$, oriT, ori p15A, *araC*, $P_{BAD}$ *riboJ* *epxD*_C3A3 (XUT$^I$) *gxpS*_T3CE4A4T4CE5A5T5TE GS-linker *msfGFP* | This work |
| pASCR3_ epoxy _M4 | *gm*$^R$, oriT, ori p15A, *araC*, $P_{BAD}$ *riboJ* *epxD*_C4A4 (XUT$^I$) *gxpS*_T3CE4A4T4CE5A5T5TE GS-linker *msfGFP* | This work |

| | | |
|---|---|---|
| pASCR8_D3 | *$cm^{R}$*, oriT, ori p15A, *araE, araC*, $P_{BAD}$ *riboJ*<br>PTEMK122_08040_A1 ($XUT^{I}$)<br>*gxpS*_T3CE4A4T4CE5A5T5TE GS-linker<br>*mScarlet-I3* | This work |
| pASCR8_D4 | *$cm^{R}$*, oriT, ori p15A, *araE, araC*, $P_{BAD}$ *riboJ*<br>PTEMK122_08040_CE2A2 ($XUT^{I}$)<br>*gxpS*_T3CE4A4T4CE5A5T5TE GS-linker<br>*mScarlet-I3* | This work |
| pASCR8_D5 | *$cm^{R}$*, oriT, ori p15A, *araE, araC*, $P_{BAD}$ *riboJ*<br>PTEMK122_08040_C3A3 ($XUT^{I}$)<br>*gxpS*_T3CE4A4T4CE5A5T5TE GS-linker<br>*mScarlet-I3* | This work |
| pASCR8_D6 | *$cm^{R}$*, oriT, ori p15A, *araE, araC*, $P_{BAD}$ *riboJ*<br>PTEMK122_08040_CE4A4 ($XUT^{I}$)<br>*gxpS*_T3CE4A4T4CE5A5T5TE GS-linker<br>*mScarlet-I3* | This work |
| pASCR8_D7 | *$cm^{R}$*, oriT, ori p15A, *araE, araC*, $P_{BAD}$ *riboJ*<br>PTEMK122_08040_CE5A5 ($XUT^{I}$)<br>*gxpS*_T3CE4A4T4CE5A5T5TE GS-linker<br>*mScarlet-I3* | This work |
| pASCR8_D8 | *$cm^{R}$*, oriT, ori p15A, *araE, araC*, $P_{BAD}$ *riboJ*<br>PTEMK122_08040_C6A6 ($XUT^{I}$)<br>*gxpS*_T3CE4A4T4CE5A5T5TE GS-linker<br>*mScarlet-I3* | This work |
| pASCR8_D9 | *$cm^{R}$*, oriT, ori p15A, *araE, araC*, $P_{BAD}$ *riboJ*<br>PTEMK122_08690_A1 ($XUT^{I}$)<br>*gxpS*_T3CE4A4T4CE5A5T5TE GS-linker<br>*mScarlet-I3* | This work |
| pASCR8_D10 | *$cm^{R}$*, oriT, ori p15A, *araE, araC*, $P_{BAD}$ *riboJ*<br>PTEMK122_08690_CE2A2 ($XUT^{I}$)<br>*gxpS*_T3CE4A4T4CE5A5T5TE GS-linker<br>*mScarlet-I3* | This work |
| pASCR8_D11 | *$cm^{R}$*, oriT, ori p15A, *araE, araC*, $P_{BAD}$ *riboJ*<br>PTEMK122_08690_CE3A3 ($XUT^{I}$)<br>*gxpS*_T3CE4A4T4CE5A5T5TE GS-linker<br>*mScarlet-I3* | This work |
| pASCR8_D12 | *$cm^{R}$*, oriT, ori p15A, *araE, araC*, $P_{BAD}$ *riboJ*<br>PTEMK122_08690_CE4A4 ($XUT^{I}$)<br>*gxpS*_T3CE4A4T4CE5A5T5TE GS-linker<br>*mScarlet-I3* | This work |
| pASCR8_E1 | *$cm^{R}$*, oriT, ori p15A, *araE, araC*, $P_{BAD}$ *riboJ*<br>PTEMK122_08690_C5A5 ($XUT^{I}$)<br>*gxpS*_T3CE4A4T4CE5A5T5TE GS-linker<br>*mScarlet-I3* | This work |
| pASCR8_E2 | *$cm^{R}$*, oriT, ori p15A, *araE, araC*, $P_{BAD}$ *riboJ*<br>PTEMK122_21930_C1A1 ($XUT^{I}$)<br>*gxpS*_T3CE4A4T4CE5A5T5TE GS-linker<br>*mScarlet-I3* | This work |
| pASCR8_E3 | *$cm^{R}$*, oriT, ori p15A, *araE, araC*, $P_{BAD}$ *riboJ*<br>PTEMK122_21930_CE2A2 ($XUT^{I}$)<br>*gxpS*_T3CE4A4T4CE5A5T5TE GS-linker<br>*mScarlet-I3* | This work |

| | | |
|---|---|---|
| pASCR8_E4 | *cm*$^R$, oriT, ori p15A, *araE, araC*, $P_{BAD}$ *riboJ* PTEMK122_21935_C3A3 (XUT$^I$) *gxpS*_T3CE4A4T4CE5A5T5TE GS-linker *mScarlet-I3* | This work |
| pASCR8_E5 | *cm*$^R$, oriT, ori p15A, *araE, araC*, $P_{BAD}$ *riboJ* PTEMK122_21935_CE4A4 (XUT$^I$) *gxpS*_T3CE4A4T4CE5A5T5TE GS-linker *mScarlet-I3* | This work |
| pASCR8_E6 | *cm*$^R$, oriT, ori p15A, *araE, araC*, $P_{BAD}$ *riboJ* PTEMK122_21935_C5A5 (XUT$^I$) *gxpS*_T3CE4A4T4CE5A5T5TE GS-linker *mScarlet-I3* | This work |
| pASCR8_Xsto_M1 | *cm*$^R$, oriT, ori p15A, *araE, araC*, $P_{BAD}$ *riboJ* XSTOV2_09090_A1nMT (XUT$^I$) *gxpS*_T3CE4A4T4CE5A5T5TE GS-linker *mScarlet-I3* | This work |
| pASCR8_Xsto_M2 | *cm*$^R$, oriT, ori p15A, *araE, araC*, $P_{BAD}$ *riboJ* XSTOV2_09090_C2A2 (XUT$^I$) *gxpS*_T3CE4A4T4CE5A5T5TE GS-linker *mScarlet-I3* | This work |
| pASCR8_Xsto_M3 | *cm*$^R$, oriT, ori p15A, *araE, araC*, $P_{BAD}$ *riboJ* XSTOV2_09090_CE3A3 (XUT$^I$) *gxpS*_T3CE4A4T4CE5A5T5TE GS-linker *mScarlet-I3* | This work |
| pASCR8_A1 | *cm*$^R$, oriT, ori p15A, *araE, araC*, $P_{BAD}$ *riboJ* *grsA*_A1 (XUT$^I$) *gxpS*_T3CE4A4T4CE5A5T5TE GS-linker *mScarlet-I3* | This work |
| pASCR8_A2 | *cm*$^R$, oriT, ori p15A, *araE, araC*, $P_{BAD}$ *riboJ* *grsB*_C2A2 (XUT$^I$) *gxpS*_T3CE4A4T4CE5A5T5TE GS-linker *mScarlet-I3* | This work |
| pASCR8_A3 | *cm*$^R$, oriT, ori p15A, *araE, araC*, $P_{BAD}$ *riboJ* *grsB*_C3A3 (XUT$^I$) *gxpS*_T3CE4A4T4CE5A5T5TE GS-linker *mScarlet-I3* | This work |
| pASCR8_A4 | *cm*$^R$, oriT, ori p15A, *araE, araC*, $P_{BAD}$ *riboJ* *grsB*_C4A4 (XUT$^I$) *gxpS*_T3CE4A4T4CE5A5T5TE GS-linker *mScarlet-I3* | This work |
| pASCR8_A5 | *cm*$^R$, oriT, ori p15A, *araE, araC*, $P_{BAD}$ *riboJ* *grsB*_C5A5 (XUT$^I$) *gxpS*_T3CE4A4T4CE5A5T5TE GS-linker *mScarlet-I3* | This work |
| pASCR8_*odlS*_M1 | *cm*$^R$, oriT, ori p15A, *araE, araC*, $P_{BAD}$ *riboJ* *odlS1*_C1A1 (XUT$^I$) *gxpS*_T3CE4A4T4CE5A5T5TE GS-linker *mScarlet-I3* | This work |
| pASCR8_*odlS*_M2 | *cm*$^R$, oriT, ori p15A, *araE, araC*, $P_{BAD}$ *riboJ* *odlS1*_CE2A2 (XUT$^I$) *gxpS*_T3CE4A4T4CE5A5T5TE GS-linker *mScarlet-I3* | This work |
| pASCR8_*odlS*_M4 | *cm*$^R$, oriT, ori p15A, *araE, araC*, $P_{BAD}$ *riboJ* *odlS2*_C4A3 (XUT$^I$) *gxpS*_T3CE4A4T4CE5A5T5TE GS-linker *mScarlet-I3* | This work |
| pASCR8_*odlS*_M5 | *cm*$^R$, oriT, ori p15A, *araE, araC*, $P_{BAD}$ *riboJ* *odlS2*_C5A4 (XUT$^I$) | This work |

| | | |
|---|---|---|
| | *gxpS*_T3CE4A4T4CE5A5T5TE GS-linker *mScarlet-I3* | |
| pASCR8_*odlS*_M6 | *cm*$^R$, oriT, ori p15A, *araE, araC*, $P_{BAD}$ *riboJ odlS2*_CE6A5 (XUT$^I$) *gxpS*_T3CE4A4T4CE5A5T5TE GS-linker *mScarlet-I3* | This work |
| pASCR8_*odlS*_M7 | *cm*$^R$, oriT, ori p15A, *araE, araC*, $P_{BAD}$ *riboJ odlS3*_C7A6 (XUT$^I$) *gxpS*_T3CE4A4T4CE5A5T5TE GS-linker *mScarlet-I3* | This work |
| pASCR8_*odlS*_M8 | *cm*$^R$, oriT, ori p15A, *araE, araC*, $P_{BAD}$ *riboJ odlS4*_C8A7 (XUT$^I$) *gxpS*_T3CE4A4T4CE5A5T5TE GS-linker *mScarlet-I3* | This work |
| pASCR8_*odlS*_M9 | *cm*$^R$, oriT, ori p15A, *araE, araC*, $P_{BAD}$ *riboJ odlS4*_C9A8 (XUT$^I$) *gxpS*_T3CE4A4T4CE5A5T5TE GS-linker *mScarlet-I3* | This work |
| pASCR8_*odlS*_M10 | *cm*$^R$, oriT, ori p15A, *araE, araC*, $P_{BAD}$ *riboJ odlS4*_C10A9 (XUT$^I$) *gxpS*_T3CE4A4T4CE5A5T5TE GS-linker *mScarlet-I3* | This work |
| pASCR8_*glbS*_M1 | *cm*$^R$, oriT, ori p15A, *araE, araC*, $P_{BAD}$ *riboJ glbE*_C1A1 (XUT$^I$) *gxpS*_T3CE4A4T4CE5A5T5TE GS-linker *mScarlet-I3* | This work |
| pASCR8_*glbS*_M2 | *cm*$^R$, oriT, ori p15A, *araE, araC*, $P_{BAD}$ *riboJ glbC*_C2A2 (XUT$^I$) *gxpS*_T3CE4A4T4CE5A5T5TE GS-linker *mScarlet-I3* | This work |
| pASCR8_*glbS*_M3 | *cm*$^R$, oriT, ori p15A, *araE, araC*, $P_{BAD}$ *riboJ glbC*_C3A3 (XUT$^I$) *gxpS*_T3CE4A4T4CE5A5T5TE GS-linker *mScarlet-I3* | This work |
| pASCR8_A6 | *cm*$^R$, oriT, ori p15A, *araE, araC*, $P_{BAD}$ *riboJ* HX791_RS00015_CE1A1 (XUT$^I$) *gxpS*_T3CE4A4T4CE5A5T5TE GS-linker *mScarlet-I3* | This work |
| pASCR8_A7 | *cm*$^R$, oriT, ori p15A, *araE, araC*, $P_{BAD}$ *riboJ* HX791_RS00015_CE2A2 (XUT$^I$) *gxpS*_T3CE4A4T4CE5A5T5TE GS-linker *mScarlet-I3* | This work |
| pASCR8_A8 | *cm*$^R$, oriT, ori p15A, *araE, araC*, $P_{BAD}$ *riboJ* HX791_RS00015_CE3A3 (XUT$^I$) *gxpS*_T3CE4A4T4CE5A5T5TE GS-linker *mScarlet-I3* | This work |
| pASCR8_A9 | *cm*$^R$, oriT, ori p15A, *araE, araC*, $P_{BAD}$ *riboJ* HX791_RS00015_C4A4 (XUT$^I$) *gxpS*_T3CE4A4T4CE5A5T5TE GS-linker *mScarlet-I3* | This work |
| pASCR8_A10 | *cm*$^R$, oriT, ori p15A, *araE, araC*, $P_{BAD}$ *riboJ* HX791_RS00015_C5A5 (XUT$^I$) *gxpS*_T3CE4A4T4CE5A5T5TE GS-linker *mScarlet-I3* | This work |
| pASCR8_A11 | *cm*$^R$, oriT, ori p15A, *araE, araC*, $P_{BAD}$ *riboJ* HX791_RS00015_C6A6 (XUT$^I$) | This work |

| | | |
|---|---|---|
| | *gxpS*_T3CE4A4T4CE5A5T5TE GS-linker *mScarlet-I3* | |
| pASCR8_B4 | *cm*$^{R}$, oriT, ori p15A, *araE, araC*, P$_{BAD}$ *riboJ* MY55_13005_C1A1 (XUT$^{I}$) *gxpS*_T3CE4A4T4CE5A5T5TE GS-linker *mScarlet-I3* | This work |
| pASCR8_B5 | *cm*$^{R}$, oriT, ori p15A, *araE, araC*, P$_{BAD}$ *riboJ* MY55_12955_C2A2 (XUT$^{I}$) *gxpS*_T3CE4A4T4CE5A5T5TE GS-linker *mScarlet-I3* | This work |
| pASCR8_B6 | *cm*$^{R}$, oriT, ori p15A, *araE, araC*, P$_{BAD}$ *riboJ* MY55_12980_A3 (XUT$^{I}$) *gxpS*_T3CE4A4T4CE5A5T5TE GS-linker *mScarlet-I3* | This work |
| pASCR8_B7 | *cm*$^{R}$, oriT, ori p15A, *araE, araC*, P$_{BAD}$ *riboJ* MY55_12980_CE3A4 (XUT$^{I}$) *gxpS*_T3CE4A4T4CE5A5T5TE GS-linker *mScarlet-I3* | This work |
| pASCR8_C12 | *cm*$^{R}$, oriT, ori p15A, *araE, araC*, P$_{BAD}$ *riboJ* MXAN_RS13550_A-Ox1 (XUT$^{I}$) *gxpS*_T3CE4A4T4CE5A5T5TE GS-linker *mScarlet-I3* | This work |
| pASCR8_F8 | *cm*$^{R}$, oriT, ori p15A, *araE, araC*, P$_{BAD}$ *riboJ* FBQ73_RS02155_A1 (XUT$^{I}$) *gxpS*_T3CE4A4T4CE5A5T5TE GS-linker *mScarlet-I3* | This work |
| pASCR8_F9 | *cm*$^{R}$, oriT, ori p15A, *araE, araC*, P$_{BAD}$ *riboJ* FBQ73_RS02140_C1A2 (XUT$^{I}$) *gxpS*_T3CE4A4T4CE5A5T5TE GS-linker *mScarlet-I3* | This work |
| pASCR8_F10 | *cm*$^{R}$, oriT, ori p15A, *araE, araC*, P$_{BAD}$ *riboJ* FBQ73_RS02140_Cy2A3 (XUT$^{I}$) *gxpS*_T3CE4A4T4CE5A5T5TE GS-linker *mScarlet-I3* | This work |
| pASCR8_epxS_ M1 | *cm*$^{R}$, oriT, ori p15A, *araE, araC*, P$_{BAD}$ *riboJ* *epxD*_C1A1nMT (XUT$^{I}$) *gxpS*_T3CE4A4T4CE5A5T5TE GS-linker *mScarlet-I3* | This work |
| pASCR8_ epxS _M2 | *cm*$^{R}$, oriT, ori p15A, *araE, araC*, P$_{BAD}$ *riboJ* *epxD*_C2A2 (XUT$^{I}$) *gxpS*_T3CE4A4T4CE5A5T5TE GS-linker *mScarlet-I3* | This work |
| pASCR8_ epxS _M3 | *cm*$^{R}$, oriT, ori p15A, *araE, araC*, P$_{BAD}$ *riboJ* *epxD*_C3A3 (XUT$^{I}$) *gxpS*_T3CE4A4T4CE5A5T5TE GS-linker *mScarlet-I3* | This work |
| pASCR8_ epxS _M4 | *cm*$^{R}$, oriT, ori p15A, *araE, araC*, P$_{BAD}$ *riboJ* *epxD*_C4A4 (XUT$^{I}$) *gxpS*_T3CE4A4T4CE5A5T5TE GS-linker *mScarlet-I3* | This work |
| pASCR12_A1 | *cm*$^{R}$, oriT, pRO1600/ColE1, *araE, araC*, P$_{BAD}$ *riboJ* *grsA*_A1 (XUT$^{I}$) *gxpS*_T3CE4A4T4CE5A5T5TE GS-linker *mScarlet-I3* | This work |
| pASCR12_A2 | *cm*$^{R}$, oriT, pRO1600/ColE1, *araE, araC*, P$_{BAD}$ *riboJ* *grsB*_C2A2 (XUT$^{I}$) *gxpS*_T3CE4A4T4CE5A5T5TE GS-linker *mScarlet-I3* | This work |

| | | |
|---|---|---|
| pASCR12_A3 | *cm*$^R$, oriT, pRO1600/ColE1, *araE, araC*, $P_{BAD}$ *riboJ grsB*_C3A3 (XUT$^I$) *gxpS*_T3CE4A4T4CE5A5T5TE GS-linker *mScarlet-I3* | This work |
| pASCR12_A4 | *cm*$^R$, oriT, pRO1600/ColE1, *araE, araC*, $P_{BAD}$ *riboJ grsB*_C4A4 (XUT$^I$) *gxpS*_T3CE4A4T4CE5A5T5TE GS-linker *mScarlet-I3* | This work |
| pASCR12_A5 | *cm*$^R$, oriT, pRO1600/ColE1, *araE, araC*, $P_{BAD}$ *riboJ grsB*_C5A5 (XUT$^I$) *gxpS*_T3CE4A4T4CE5A5T5TE GS-linker *mScarlet-I3* | This work |

**Table S3.** Primer and template used in this work to generate indicated plasmids. Sizes of the PCR products are depicted below the template.

| Plasmids | Oligo-nucleotides | Sequence 5'→3 | Template/ Product size in bp |
|---|---|---|---|
| pLP179 | pc1080 | TTTTTGGGCTAACAGGAGGAATTCCATGAATATGACACGTAACCATACATCC | *X. indica*<br>gDNA<br>3.089 kb |
| | 30 | GCGAGTGACCGCCCAAGGCAAAGAAATGATCGTGGCGACCGACAC | |
| | pc1101 | TTCTTTGCCTTGGGCGGTCAC | *P. laumondii*<br>TTO1 gDNA<br>4.475 kb |
| | pc1102 | GCCAGTGCCAACGAGGCATATTCG | |
| | pc1103 | CGAATATGCCTCGTTGGCACTGGC | *P. laumondii*<br>TTO1 gDNA<br>3.019 kb |
| | pc1104 | ATACGAGCCGATGATTAATTGTCACAGCGCCTCCGCTTCACAATTC | |
| | pc1091 | TGACAATTAATCATCGGCTCGTATAATGTG | pACYC<br>5.220 kb |
| | pc1090 | GGAATTCCTCCTGTTAGCCCAAAAAAACG | |
| pTH3 | TH3-1-O | AGGAGGAATTCCATGTTTCTAGATAAAGTCGGGCAGC | *X. nematophila*<br>gDNA<br>3.087 kb |
| | TH3-2-O | CAAGGCAAAGAAGTTGTCATGGCGGCTGACTTTATC | |
| | TH3-4-O | CCGCCATGACAACTTCTTTGCCTTGGGCGGTCAC | *P. laumondii*<br>TTO1 gDNA<br>5.156 kb |
| | TH3-RV | CCGCATCCGTGAGCATATAAGCTAACC | |
| | TH3-FW | GGTTAGCTTATATGCTCACGGATGCGG | *P. laumondii*<br>TTO1 gDNA<br>2.351 kb |
| | jw0127_rev | CGAGCCGATGATTAATTGTCACAGCGCCTCCGCTTC | |
| | jw0061 | TGACAATTAATCATCGGCTCG | pCOLA<br>3.322 kb |
| | TH3-3-O | TATCTAGAAACATGGAATTCCTCCTGTTAGCCC | |
| pLP212 | 26 | TTTTTGGGCTAACAGGAGGAATTCCATGAATATGACACGTAACCATACATCC | *X. indica*<br>gDNA<br>6.337 kb |
| | LP30_rev | AAGTTTAAAGAAATGGTCATAACGTCCGACG | |
| | LP29_fw | GTTATGACCATTTCTTTAAACTTGGCGGTCACTCGCTGTTGG | *P. laumondii*<br>TTO1 gDNA<br>4.486 kb |
| | pc1102 | GCCAGTGCCAACGAGGCATATTCG | |
| | pc1103 | CGAATATGCCTCGTTGGCACTGGC | *P. laumondii*<br>TTO1 gDNA<br>3.019 kb |
| | pc1104 | ATACGAGCCGATGATTAATTGTCACAGCGCCTCCGCTTCACAATTC | |
| | pc1091 | TGACAATTAATCATCGGCTCGTATAATGTG | pACYC<br>5.220 kb |
| | pc1090 | GGAATTCCTCCTGTTAGCCCAAAAAAACG | |
| pLP228 | LP582 | TTTTTGGGCTAACAGGAGGAATTCCATGGCTGTCAACAAGCTG | *X. nematophila*<br>gDNA<br>3.185 kb |
| | pc1105 | TGACTGCCAACAGCGAGTGACCGCCGAGGTGGAAAAAGTTGTCGTTTC | |
| | pc1106 | GGCGGTCACTCGCTGTTGGCAG | *P. laumondii*<br>TTO1 gDNA<br>4.463 kb |
| | pc1102 | GCCAGTGCCAACGAGGCATATTCG | |
| | pc1103 | CGAATATGCCTCGTTGGCACTGGC | *P. laumondii*<br>TTO1 gDNA<br>3.019 kb |
| | pc1104 | ATACGAGCCGATGATTAATTGTCACAGCGCCTCCGCTTCACAATTC | |

| | | | |
|---|---|---|---|
| | pc1091 | TGACAATTAATCATCGGCTCGTATAATGTG | pACYC 5.220 kb |
| | pc1090 | GGAATTCCTCCTGTTAGCCCAAAAAAACG | |
| pLP236 | LP582 | TTTTTGGGCTAACAGGAGGAATTCCATGG CTGTCAACAAGCTG | *X. nematophila* gDNA 3.185 kb |
| | LP598 | GGATAACCAACAAAGCATGGCCTCCGAG G | |
| | LP597 | GCTTTGTTGGTTATCCGGGTCATTTCCAA GATACG | *X. nematophila* gDNA 1.780 kb |
| | LP599 | TGACTGCCAACAGCGAGTGACCGCCTATT GCGAAGAAGTTATCGTTAC | |
| | pc1106 | GGCGGTCACTCGCTGTTGGCAG | *P. laumondii* TTO1 gDNA 4.463 kb |
| | pc1102 | GCCAGTGCCAACGAGGCATATTCG | |
| | pc1103 | CGAATATGCCTCGTTGGCACTGGC | *P. laumondii* TTO1 gDNA 3.019 kb |
| | pc1104 | ATACGAGCCGATGATTAATTGTCACAGCG CCTCCGCTTCACAATTC | |
| | pc1091 | TGACAATTAATCATCGGCTCGTATAATGTG | pACYC 5.220 kb |
| | pc1090 | GGAATTCCTCCTGTTAGCCCAAAAAAACG | |
| pLP229 | LP583 | TTTTTGGGCTAACAGGAGGAATTCCATGA TGGATATGCTGAAATTAGTC | *X. nematophila* gDNA 3.263 kb |
| | LP584 | TGACTGCCAACAGCGAGTGACCGCCCGC CATAAAAAAATTGTCATCC | |
| | pc1106 | GGCGGTCACTCGCTGTTGGCAG | *P. laumondii* TTO1 gDNA 4.463 kb |
| | pc1102 | GCCAGTGCCAACGAGGCATATTCG | |
| | pc1103 | CGAATATGCCTCGTTGGCACTGGC | *P. laumondii* TTO1 gDNA 3.019 kb |
| | pc1104 | ATACGAGCCGATGATTAATTGTCACAGCG CCTCCGCTTCACAATTC | |
| | pc1091 | TGACAATTAATCATCGGCTCGTATAATGTG | pACYC 5.220 kb |
| | pc1090 | GGAATTCCTCCTGTTAGCCCAAAAAAACG | |
| pLP230 | LP585 | TTTTTGGGCTAACAGGAGGAATTCCATGA TTATTGAACGCAAAACAC | *X. nematophila* gDNA 3.083 kb |
| | LP586 | TGACTGCCAACAGCGAGTGACCGCCCAA CTCAAAGAAGTTGG | |
| | pc1106 | GGCGGTCACTCGCTGTTGGCAG | *P. laumondii* TTO1 gDNA 4.463 kb |
| | pc1102 | GCCAGTGCCAACGAGGCATATTCG | |
| | pc1103 | CGAATATGCCTCGTTGGCACTGGC | *P. laumondii* TTO1 gDNA 3.019 kb |
| | pc1104 | ATACGAGCCGATGATTAATTGTCACAGCG CCTCCGCTTCACAATTC | |
| | pc1091 | TGACAATTAATCATCGGCTCGTATAATGTG | pACYC 5.220 kb |
| | pc1090 | GGAATTCCTCCTGTTAGCCCAAAAAAACG | |
| pLP237 | LP585 | TTTTTGGGCTAACAGGAGGAATTCCATGA TTATTGAACGCAAAACAC | *X. nematophila* gDNA 2.981 kb |
| | LP601 | CAATTTCCCCTTGTGGCGCTTGGTATTCA ATTTTCCGTTGAGC | |
| | LP600 | TACCAAGCGCCACAAGGGGAAATTGAG | *P. laumondii* TTO1 gDNA 4.565 kb |
| | pc1102 | GCCAGTGCCAACGAGGCATATTCG | |
| | pc1103 | CGAATATGCCTCGTTGGCACTGGC | *P. laumondii* TTO1 gDNA 3.019 kb |
| | pc1104 | ATACGAGCCGATGATTAATTGTCACAGCG CCTCCGCTTCACAATTC | |
| | pc1091 | TGACAATTAATCATCGGCTCGTATAATGTG | pACYC |

| | | | |
|---|---|---|---|
| | pc1090 | GGAATTCCTCCTGTTAGCCCAAAAAAACG | 5.220 kb |
| pLP232 | LP589 | TTTTTGGGCTAACAGGAGGAATTCCATGC AGTTAACGGCCCATG | *X. nematophila* gDNA 3.149 kb |
| | LP590 | TGACTGCCAACAGCGAGTGACCGCCTAAA TCAAAAAAGCTGTCGGCC | |
| | pc1106 | GGCGGTCACTCGCTGTTGGCAG | *P. laumondii* TTO1 gDNA 4.463 kb |
| | pc1102 | GCCAGTGCCAACGAGGCATATTCG | |
| | pc1103 | CGAATATGCCTCGTTGGCACTGGC | *P. laumondii* TTO1 gDNA 3.019 kb |
| | pc1104 | ATACGAGCCGATGATTAATTGTCACAGCG CCTCCGCTTCACAATTC | |
| | pc1091 | TGACAATTAATCATCGGCTCGTATAATGTG | pACYC 5.220 kb |
| | pc1090 | GGAATTCCTCCTGTTAGCCCAAAAAAACG | |
| pLP231 | LP587 | TTTTTGGGCTAACAGGAGGAATTCCATGG GGATTATGAACCTAAAAGAC | *X. nematophila* gDNA 3.278 kb |
| | LP588 | TGACTGCCAACAGCGAGTGACCGCCGAG CTGTACGAACCGG | |
| | pc1106 | GGCGGTCACTCGCTGTTGGCAG | *P. laumondii* TTO1 gDNA 4.463 kb |
| | pc1102 | GCCAGTGCCAACGAGGCATATTCG | |
| | pc1103 | CGAATATGCCTCGTTGGCACTGGC | *P. laumondii* TTO1 gDNA 3.019 kb |
| | pc1104 | ATACGAGCCGATGATTAATTGTCACAGCG CCTCCGCTTCACAATTC | |
| | pc1091 | TGACAATTAATCATCGGCTCGTATAATGTG | pACYC 5.220 kb |
| | pc1090 | GGAATTCCTCCTGTTAGCCCAAAAAAACG | |
| pPL238 | LP587 | TTTTTGGGCTAACAGGAGGAATTCCATGG GGATTATGAACCTAAAAGAC | *X. nematophila* gDNA 3.176 kb |
| | LP602 | CAATTTCCCCTTGTGGCGCTTGGTAAGGC TCTTCGACGATATTTTCC | |
| | LP600 | TACCAAGCGCCACAAGGGGAAATTGAG | *P. laumondii* TTO1 gDNA 4.565 kb |
| | pc1102 | GCCAGTGCCAACGAGGCATATTCG | |
| | pc1103 | CGAATATGCCTCGTTGGCACTGGC | *P. laumondii* TTO1 gDNA 3.019 kb |
| | pc1104 | ATACGAGCCGATGATTAATTGTCACAGCG CCTCCGCTTCACAATTC | |
| | pc1091 | TGACAATTAATCATCGGCTCGTATAATGTG | pACYC 5.220 kb |
| | pc1090 | GGAATTCCTCCTGTTAGCCCAAAAAAACG | |
| pLP233 | LP591 | TTTTTGGGCTAACAGGAGGAATTCCATGA ATATTGCGGCATTTTTAGCTG | *X. nematophila* gDNA 3.257 kb |
| | LP592 | TGACTGCCAACAGCGAGTGACCGCCGAG CCTGAAAAAGTTATCAGTAC | |
| | pc1106 | GGCGGTCACTCGCTGTTGGCAG | *P. laumondii* TTO1 gDNA 4.463 kb |
| | pc1102 | GCCAGTGCCAACGAGGCATATTCG | |
| | pc1103 | CGAATATGCCTCGTTGGCACTGGC | *P. laumondii* TTO1 gDNA 3.019 kb |
| | pc1104 | ATACGAGCCGATGATTAATTGTCACAGCG CCTCCGCTTCACAATTC | |
| | pc1091 | TGACAATTAATCATCGGCTCGTATAATGTG | pACYC 5.220 kb |
| | pc1090 | GGAATTCCTCCTGTTAGCCCAAAAAAACG | |
| pLP234 | LP593 | TTTTTGGGCTAACAGGAGGAATTCCATGA TAATCCCTGTGGCGAAAACAG | *X. nematophila* gDNA 3.104 kb |
| | LP594 | TGACTGCCAACAGCGAGTGACCGCCTAAA GCGAAGAAATTATCGTAACGACC | |

| | | | |
|---|---|---|---|
| | pc1106 | GGCGGTCACTCGCTGTTGGCAG | *P. laumondii* TTO1 gDNA 4.463 kb |
| | pc1102 | GCCAGTGCCAACGAGGCATATTCG | |
| | pc1103 | CGAATATGCCTCGTTGGCACTGGC | *P. laumondii* TTO1 gDNA 3.019 kb |
| | pc1104 | ATACGAGCCGATGATTAATTGTCACAGCG CCTCCGCTTCACAATTC | |
| | pc1091 | TGACAATTAATCATCGGCTCGTATAATGTG | pACYC 5.220 kb |
| | pc1090 | GGAATTCCTCCTGTTAGCCCAAAAAAACG | |
| pLP235 | LP595 | TTTTTGGGCTAACAGGAGGAATTCCATGT ATCACGCTTTTTCATTACTC | *X. nematophila* gDNA 3.110 kb |
| | LP596 | TGACTGCCAACAGCGAGTGACCGCCCATT TCCATAAAAGATGCG | |
| | pc1106 | GGCGGTCACTCGCTGTTGGCAG | *P. laumondii* TTO1 gDNA 4.463 kb |
| | pc1102 | GCCAGTGCCAACGAGGCATATTCG | |
| | pc1103 | CGAATATGCCTCGTTGGCACTGGC | *P. laumondii* TTO1 gDNA 3.019 kb |
| | pc1104 | ATACGAGCCGATGATTAATTGTCACAGCG CCTCCGCTTCACAATTC | |
| | pc1091 | TGACAATTAATCATCGGCTCGTATAATGTG | pACYC 5.220 kb |
| | pc1090 | GGAATTCCTCCTGTTAGCCCAAAAAAACG | |
| pLP239 | LP595 | TTTTTGGGCTAACAGGAGGAATTCCATGT ATCACGCTTTTTCATTACTC | *X. nematophila* gDNA 3.002 kb |
| | LP603 | CAATTTCCCCTTGTGGCGCTTGGTATGAT CCGCGCACAAGACAGGG | |
| | LP600 | TACCAAGCGCCACAAGGGGAAATTGAG | *P. laumondii* TTO1 gDNA 4.565 kb |
| | pc1102 | GCCAGTGCCAACGAGGCATATTCG | |
| | pc1103 | CGAATATGCCTCGTTGGCACTGGC | *P. laumondii* TTO1 gDNA 3.019 kb |
| | pc1104 | ATACGAGCCGATGATTAATTGTCACAGCG CCTCCGCTTCACAATTC | |
| | pc1091 | TGACAATTAATCATCGGCTCGTATAATGTG | pACYC 5.220 kb |
| | pc1090 | GGAATTCCTCCTGTTAGCCCAAAAAAACG | |
| pASCR2 | LP612 | GTCGTGACTGGGAAAACCCTGGCG | pSEVA661b-Neon 3.586 kb |
| | LP635 | TACGTATTTCCCCTCTTTCTCTAGTATTAA AC | |
| | LP636 | GTTTAATACTAGAGAAAGAGGGGAAATAC GTA TACCAAGCGCCACAAGGGGAAATTG | *P. laumondii* TTO1 gDNA 4.597 kb |
| | pc1102 | GCCAGTGCCAACGAGGCATATTCG | |
| | pc1103 | CGAATATGCCTCGTTGGCACTGGC | *P. laumondii* TTO1 gDNA 2.415 kb |
| | LP637 | TAAGCCAGTATCCCGCCAGAAGAGTAGC | |
| | LP638 | ACTCTTCTGGCGGGATACTGGCTTATGC | *P. laumondi* TTO1 gDNA 633 bp |
| | LP611 | TCGCCAGGGTTTTCCCAGTCACGACT TACAGCGCCTCCGCTTCACAATTC | |
| pASCR3 | LP612 | GTCGTGACTGGGAAAACCCTGGCG | pSEVA661b-Neon[23] 3.586 kb |
| | LP635 | TACGTATTTCCCCTCTTTCTCTAGTATTAA AC | |

| | | | |
|---|---|---|---|
| | LP636 | GTTTAATACTAGAGAAAGAGGGGAAATAC<br>GTA TACCAAGCGCCACAAGGGGAAATTG | *P. laumondii*<br>TTO1<br>gDNA<br>4.597 kb |
| | pc1102 | GCCAGTGCCAACGAGGCATATTCG | |
| | pc1103 | CGAATATGCCTCGTTGGCACTGGC | *P. laumondii*<br>TTO1<br>gDNA<br>2.415 kb |
| | LP637 | TAAGCCAGTATCCCGCCAGAAGAGTAGC | |
| | LP638 | ACTCTTCTGGCGGGATACTGGCTTATGC | *P. laumondi*<br>TTO1<br>gDNA<br>630 bp |
| | LP688 | AACCAGCAGCGGAGCCAGCGGATCC<br>CAGCGCCTCCGCTTCACAATTCATTG | |
| | LP689 | GGATCCGCTGGCTCCGCTGCTGGTTCTG<br>GCGAATTCATGTCCAAGGGTGAAGAG | pISNP4<br>784 bp |
| | LP690 | TCGCCAGGGTTTTCCCAGTCACGAC<br>TTACGACCCCTTATAAAGC | |
| pASCR4 | LP612 | GTCGTGACTGGGAAAACCCTGGCG | pSEVA661b-<br>Neon<br>3.586 kb |
| | LP635 | TACGTATTTCCCCTCTTTCTCTAGTATTAA<br>AC | |
| | LP691 | ACTAGAGAAAGAGGGGAAATACGTA<br>ATGTCGGACTCAGAAGTCAATC | pCIOX<br>344 bp |
| | LP692 | TGAGACCTTGCATTCGTTGGTCTCG<br>ACCACCAATCTGTTCTCTGTG | |
| | LP693 | CGAGACCAACGAATGCAAGGTCTCA<br>TACCAAGCGCCACAAGGGGAAATTG | *P. laumondii*<br>TTO1<br>gDNA<br>4.590 kb |
| | pc1102 | GCCAGTGCCAACGAGGCATATTCG | |
| | pc1103 | CGAATATGCCTCGTTGGCACTGGC | *P. laumondii*<br>TTO1<br>gDNA<br>2.415 kb |
| | LP637 | TAAGCCAGTATCCCGCCAGAAGAGTAGC | |
| | LP638 | ACTCTTCTGGCGGGATACTGGCTTATGC | *P. laumondi*<br>TTO1<br>gDNA<br>630 bp |
| | LP688 | AACCAGCAGCGGAGCCAGCGGATCC<br>CAGCGCCTCCGCTTCACAATTCATTG | |
| | LP689 | GGATCCGCTGGCTCCGCTGCTGGTTCTG<br>GCGAATTCATGTCCAAGGGTGAAGAG | pISNP4<br>784 bp |
| | LP690 | TCGCCAGGGTTTTCCCAGTCACGAC<br>TTACGACCCCTTATAAAGC | |
| pASCR_mf<br>GPF | LP612 | GTCGTGACTGGGAAAACCCTGGCG | pSEVA661b-<br>Neon<br>3.586 kb |
| | LP635 | TACGTATTTCCCCTCTTTCTCTAGTATTAA<br>AC | |
| | LP697 | ACTAGAGAAAGAGGGGAAATACGTA<br>ATGTCCAAGGGTGAAGAG | pISNP4<br>773 bp |
| | LP690 | TCGCCAGGGTTTTCCCAGTCACGAC<br>TTACGACCCCTTATAAAGC | |
| pASCR_e<br>mpty | LP612 | GTCGTGACTGGGAAAACCCTGGCG | pSEVA661b-<br>Neon<br>3.586 kb |
| | LP635 | TACGTATTTCCCCTCTTTCTCTAGTATTAA<br>AC | |
| pASCR5 | LP612 | GTCGTGACTGGGAAAACCCTGGCG | pSEVA661b-<br>Neon<br>3.586 kb |
| | LP635 | TACGTATTTCCCCTCTTTCTCTAGTATTAA<br>AC | |
| | LP740 | ACTAGAGAAAGAGGGGAAATACGTA<br>ATGGTCTTGCCGTTATCATTTGG | *P. laumondii*<br>TTO1 |

| | | | |
|---|---|---|---|
| | LP741 | CGTGACCGCCCAAGGCAAAGAAACTGTC | gDNA 3.014 kb |
| | LP742 | AGTTTCTTTGCCTTGGGCGGTCACGCGCT GTTGGCAGTCAGGATGATCG | *P. laumondii* TTO1 gDNA 3.253 kb |
| | LP743 | CATGGCCCCCCAAGGCAAAGAAGCTG | |
| | LP744 | AGCTTCTTTGCCTTGGGGGGCCATG CGTTGCTTGCGGTACGGATGGTTG | pASCR3 3.238 kb |
| | LP745 | CATGTCCACCCAAGGTAAAGAAGTTGTCA TGC | |
| | LP746 | AACTTCTTTACCTTGGGTGGACATGCGCT GCTCGCTATGCGGATGATC | pASCR3 1.033 kb |
| | LP747 | AACCAGCAGCGGAGCCAGCGGATCCCAG CGCCTCCGCTTCACAATTCATTG | |
| | LP689 | GGATCCGCTGGCTCCGCTGCTGGTTCTG GCGAATTCATGTCCAAGGGTGAAGAG | pASCR3 784 bp |
| | LP690 | TCGCCAGGGTTTTCCCAGTCACGAC TTACGACCCCTTATAAAGC | |
| pASCR6 | LP612 | GTCGTGACTGGGAAAACCCTGGCG | pSEVA661b-Neon 3.586 kb |
| | LP635 | TACGTATTTCCCCTCTTTCTCTAGTATTAA AC | |
| | LP740 | ACTAGAGAAAGAGGGGAAATACGTA ATGGTCTTGCCGTTATCATTTGG | *P. laumondii* TTO1 gDNA 3.014 kb |
| | LP741 | CGTGACCGCCCAAGGCAAAGAAACTGTC | |
| | LP742 | AGTTTCTTTGCCTTGGGCGGTCACGCGCT GTTGGCAGTCAGGATGATCG | *P. laumondii* TTO1 gDNA 3.253 kb |
| | LP743 | CATGGCCCCCCAAGGCAAAGAAGCTG | |
| | LP744 | AGCTTCTTTGCCTTGGGGGGCCATG CGTTGCTTGCGGTACGGATGGTTG | pASCR3 3.238 kb |
| | LP745 | CATGTCCACCCAAGGTAAAGAAGTTGTCA TGC | |
| | LP746 | AACTTCTTTACCTTGGGTGGACATGCGCT GCTCGCTATGCGGATGATC | pASCR3 1.036 kb |
| | LP611 | TCGCCAGGGTTTTCCCAGTCACGACT TACAGCGCCTCCGCTTCACAATTC | |
| pASCR8 | LP759 | GTCGTGACTGGGAAAACCC | pACYC_Seva 3.620 kb |
| | LP800 | TGATGTAGCCGTCAAGTTGTCATG | |
| | LP797 | TCATGACAACTTGACGGCTACATCATTCA C | pASCR3 6.000 kb |
| | pc1102 | GCCAGTGCCAACGAGGCATATTCG | |
| | pc1103 | CGAATATGCCTCGTTGGCACTGGC | pASCR3 3.020 kb |
| | LP688 | AACCAGCAGCGGAGCCAGCGGATCCCAG CGCCTCCGCTTCACAATTCATTG | |
| | LP798 | GATCCGCTGGCTCCGCTGCTGGTTCTGG CGAATTCGATAGCACTGAAGCAGTCATTA AAG | pISNP4 747 bp |
| | LP799 | TCGCCAGGGTTTTCCCAGTCACGACTTAC GATCCTCCAGACCCG | |
| pASCR10 | LP797 | TCATGACAACTTGACGGCTACATCATTCA C | pASCR5 4.424 kb |
| | LP741 | CGTGACCGCCCAAGGCAAAGAAACTGTC | |

| | | | |
|---|---|---|---|
| | LP742 | AGTTTCTTTGCCTTGGGCGGTCACGCGCTGTTGGCAGTCAGGATGATCG | pASCR5 7.474 kb |
| | LP688 | AACCAGCAGCGGAGCCAGCGGATCCCAGCGCCTCCGCTTCACAATTCATTG | |
| | LP798 | GATCCGCTGGCTCCGCTGCTGGTTCTGGCGAATTCGATAGCACTGAAGCAGTCATTAAAG | pASCR8 4.342 kb |
| | LP800 | TGATGTAGCCGTCAAGTTGTCATG | |
| pASCR11 | LP742 | AGTTTCTTTGCCTTGGGCGGTCACGCGCTGTTGGCAGTCAGGATGATCG | pASCR6 7.435 kb |
| | LP611 | TCGCCAGGGTTTTCCCAGTCACGACTTACAGCGCCTCCGCTTCACAATTC | |
| | LP759 | GTCGTGACTGGGAAAACCC | pACYC_Seva 3.620 kb |
| | LP800 | TGATGTAGCCGTCAAGTTGTCATG | |
| | LP797 | TCATGACAACTTGACGGCTACATCATTCAC | pASCR6 4.424 kb |
| | LP741 | CGTGACCGCCCAAGGCAAAGAAACTGTC | |
| pASCR_mScarlet-I3 | LP635 | TACGTATTTCCCCTCTTTCTCTAGTATTAAAC | pASCR8 5.030 kb |
| | LP759 | GTCGTGACTGGGAAAACCC | |
| | LP812 | ACTAGAGAAAGAGGGGAAATACGTAATGGATAGCACTGAAGCAGTC | pISNP4 740 bp |
| | LP799 | TCGCCAGGGTTTTCCCAGTCACGACTTACGATCCTCCAGACCCG | |
| pASCR8_empty | LP635 | TACGTATTTCCCCTCTTTCTCTAGTATTAAAC | pASCR8 5.030 kb |
| | LP759 | GTCGTGACTGGGAAAACCC | |
| pASCR12 | AR900 | GTCGTGACTGGGAAAACCCT | pSEVA341[24] 3.327 kb |
| | AR1081 | TCCTGTGTGAAATTGTTATCCGCT | |
| | AR703n | CCTTTAATTAAAGCGGATAACAATTTCACACAGGA | pASCR8 7.495 kb |
| | pc1102 | GCCAGTGCCAACGAGGCATATTCG | |
| | pc1103 | CGAATATGCCTCGTTGGCACTGGC | pASCR8 3.718 kb |
| | LP799 | TCGCCAGGGTTTTCCCAGTCACGACTTACGATCCTCCAGACCCG | |
| pASCR13 | LP798 | GGATCCGCTGGCTCCGCTGCTGGTTCTGGCGAATTCGATAGCACTGAAGCAGTCATTAAAG | pASCR12 5.534 kb |
| | LP800 | TGATGTAGCCGTCAAGTTGTCATG | |
| | LP797 | TCATGACAACTTGACGGCTACATCATTCAC | pASCR5 4.424 kb |
| | LP741 | CGTGACCGCCCAAGGCAAAGAAACTGTC | |
| | LP742 | AGTTTCTTTGCCTTGGGCGGTCACGCGCTGTTGGCAGTCAGGATGATCG | pASCR5 7.474 kb |
| | LP688 | AACCAGCAGCGGAGCCAGCGGATCCCAGCGCCTCCGCTTCACAATTCATTG | |
| pASCR12_mScarlet-I3 | LP759 | GTCGTGACTGGGAAAACCC | pASCR12 6.222 kb |
| | LP635 | TACGTATTTCCCCTCTTTCTCTAGTATTAAAC | |
| | LP812 | ACTAGAGAAAGAGGGGAAATACGTAATGGATAGCACTGAAGCAGTC | pASCR8 740 bp |
| | LP799 | TCGCCAGGGTTTTCCCAGTCACGACTTACGATCCTCCAGACCCG | |

| | | | |
|---|---|---|---|
| pJL09 | pCEP_kan_bb_fw | ATGTGCATGCTCGAGCTC | pCEP 4.841 kb |
| | pCEP_kan_bb_rv | ATGCTAGCCTCCTGTTAGC | |
| | JL25 | TTTGGGCTAACAGGAGGCTAGCATATGAAAGATAACATTGCTAACGTGG | *P. temperata* K122 gDNA 673 bp |
| | JL26 | TCTGCAGAGCTCGAGCATGCACATCATCCTGCTCTCCTGATTTGTC | |
| pJL05 | pCEP_kan_bb_fw | ATGTGCATGCTCGAGCTC | pCEP 4.841 kb |
| | pCEP_kan_bb_rv | ATGCTAGCCTCCTGTTAGC | |
| | JL13 | TTTGGGCTAACAGGAGGCTAGCATATGAAAGATAGCATTGCTCAAGCAG | *P. temperata* K122 gDNA 836 bp |
| | JL14 | TCTGCAGAGCTCGAGCATGCACATGACAACACATTAAGCTGTCCGAC | |
| pJL14 | pCEP_kan_bb_fw | ATGTGCATGCTCGAGCTC | pCEP 4.841 kb |
| | pCEP_kan_bb_rv | ATGCTAGCCTCCTGTTAGC | |
| | JL40 | TTTGGGCTAACAGGAGGCTAGCATATGTTAGATGAATATTTAAGTGCTCTTGATG | *P. temperata* K122 gDNA 766 bp |
| | JL41 | TCTGCAGAGCTCGAGCATGCACATGTGGTAGATCAACAAGCTGTGC | |
| pLP285 | LP759 | GTCGTGACTGGGAAAACCC | pACYC_Seva 4.817 kb |
| | LP758 | GGAATTCCTCCTGTTAGCCC | |
| | LP786 | TTTTTGGGCTAACAGGAGGAATTCCATGTATAAATATCAAGATTATCTGTGTATAC | *X. stockiae* gDNA 7.167 kb |
| | LP792 | ACATGACACAAGCTTGCCAGACTCAC | |
| | LP793 | GTGAGTCTGGCAAGCTTGTGTCATG | *X. stockiae* gDNA 4.596 kb |
| | LP795 | TCGCCAGGGTTTTCCCAGTCACGACTTAGTTATCATGTTCATTCAGCGATG | |
| pLP286 | LP759 | GTCGTGACTGGGAAAACCC | pCOLA_Seva 4.875 kb |
| | LP758 | GGAATTCCTCCTGTTAGCCC | |
| | LP796 | TTTTTGGGCTAACAGGAGGAATTCCATGGCTAATTCCTATCGTGACC | *X. stockiae* gDNA 1.265 kb |
| | LP795 | TCGCCAGGGTTTTCCCAGTCACGACTTAGTTATCATGTTCATTCAGCGATG | |

**Table S4.** Primer name and sequence used for amplification of genes encoding for A domains and C-A didomains used for cloning into A domain screening assay cloning vectors with indication of the template DNA and resulting fragment size. The first column indicates the fragment name (sometimes plasmid name) and the cloning vectors the fragment was inserted to by Gibson assembly. (pASCR2 = 11.125 kb; pASCR3 = 11.881 kb; pASCR4 = 12.200 kb; pASCR8 = 13.289 kb, pASCR12 = 14.481 kb)

| Fragment and Cloning vector | Oligo-nucleotides | Sequence 5'→3 | Template/ Product size in bp |
|---|---|---|---|
| pLP262 (pASCR2)/ pLP256 (pASCR3) | AS_077 | ACTAGAGAAAGAGGGGAAATACGTAATG AAAGATAACATTGCTAACGTGG | *P. temperata* K122 gDNA 2.414 kb |
| | AS_083 | CAATTTCCCCTTGTGGCGCTTGGTA AACCTGACGGGCGAAGGCTTCC | |
| pLP263 (pASCR2)/ pLP257 (pASCR3) | AS_078 | ACTAGAGAAAGAGGGGAAATACGTAATG CAGGATATCTATGCATTGTCAC | *P. temperata* K122 gDNA 2.912 kb |
| | AS_084 | CAATTTCCCCTTGTGGCGCTTGGTA AACCTGACGGGCAAAGGCTTCC | |
| pLP264 (pASCR2)/ pLP258 (pASCR3) | AS_079 | ACTAGAGAAAGAGGGGAAATACGTAATG GGTGAATTGCCGCTGTCATTCG | *P. temperata* K122 gDNA 2.936 kb |
| | AS_085 | CAATTTCCCCTTGTGGCGCTTGGTA AACCTGACGGGCAAAGGCCTCC | |
| pLP265 (pASCR2)/ pLP259 (pASCR3) | AS_080 | ACTAGAGAAAGAGGGGAAATACGTAATG CAGGATATCTATGCCCTGTCGC | *P. temperata* K122 gDNA 2.918 kb |
| | AS_086 | CAATTTCCCCTTGTGGCGCTTGGTA AACCTGACGGGCAAAAGCGTTC | |
| pLP266 (pASCR2)/ pLP260 (pASCR3) | AS_081 | ACTAGAGAAAGAGGGGAAATACGTAATG CAGGATATCTATGGATTGTCGC | *P. temperata* K122 gDNA 2.912 kb |
| | AS_087 | CAATTTCCCCTTGTGGCGCTTGGTA GACCTGACGGGCAAAAGACGCC | |
| pLP278 (pASCR2)/ pLP261 (pASCR3) | AS_082 | ACTAGAGAAAGAGGGGAAATACGTAATG GGCAAATTGCCGTTGTCATTTG | *P. temperata* K122 gDNA 2.948 kb |
| | AS_088 | CAATTTCCCCTTGTGGCGCTTGGTA AACCAAACGGGCAAAAGCC | |
| pLP250 (pASCR4) | LP698 | GGCTCACAGAGAACAGATTGGTGGTAAA GATAACATTGCTAACGTGG | *P. temperata* K122 gDNA 2.411 kb |
| | AS_083 | CAATTTCCCCTTGTGGCGCTTGGTAAACC TGACGGGCGAAGGCTTCC | |
| pLP251 (pASCR4) | LP699 | GGCTCACAGAGAACAGATTGGTGGTCAG GATATCTATGCATTGTCAC | *P. temperata* K122 gDNA 2.909 kb |
| | AS_084 | CAATTTCCCCTTGTGGCGCTTGGTAAACC TGACGGGCAAAGGCTTCC | |
| pLP252 (pASCR4) | LP700 | GGCTCACAGAGAACAGATTGGTGGTGGT GAATTGCCGCTGTCATTCG | *P. temperata* K122 gDNA 2.933 kb |
| | AS_085 | CAATTTCCCCTTGTGGCGCTTGGTAAACC TGACGGGCAAAGGCCTCC | |
| pLP253 (pASCR4) | LP701 | GGCTCACAGAGAACAGATTGGTGGTCAG GATATCTATGCCCTGTCGC | *P. temperata* K122 gDNA 2.915 kb |
| | AS_086 | CAATTTCCCCTTGTGGCGCTTGGTAAACC TGACGGGCAAAAGCGTTC | |
| pLP254 | LP702 | GGCTCACAGAGAACAGATTGGTGGTCAG GATATCTATGGATTGTCGC | *P. temperata* K122 |

| | | | |
|---|---|---|---|
| | AS_087 | CAATTTCCCCTTGTGGCGCTTGGTAGACC TGACGGGCAAAGACGCC | gDNA 2.909 kb |
| pLP255 (pASCR4) | LP703 | GGCTCACAGAGAACAGATTGGTGGTGGC AAATTGCCGTTGTCATTTG | *P. temperata* K122 gDNA 2.945 kb |
| | AS_088 | CAATTTCCCCTTGTGGCGCTTGGTAAACC AAACGGGCAAAGCCTCTC | |
| D9 (pASCR3 or pASCR8) | AS_089 | ACTAGAGAAAGAGGGGAAATACGTAATG AAAGATAGCATTGCTCAAGCAG | *P. temperata* K122 gDNA 2.420 kb |
| | AS_094 | CAATTTCCCCTTGTGGCGCTTGGTA AATCTGCCGGACAAAATCGTCTTC | |
| D10 (pASCR3 or pASCR8) | AS_090 | ACTAGAGAAAGAGGGGAAATACGTAATG CAGGATATCTATGGGTTGTCGC | *P. temperata* K122 gDNA 2.912 kb |
| | AS_095 | CAATTTCCCCTTGTGGCGCTTGGTA AACTTGATAAGCAAAGGCTTCATTG | |
| D11 (pASCR3 or pASCR8) | AS_091 | ACTAGAGAAAGAGGGGAAATACGTAATG CAGGATATCTATGCCCTGTCGC | *P. temperata* K122 gDNA 2.912 kb |
| | AS_096 | CAATTTCCCCTTGTGGCGCTTGGTA AACCTGACGGGCAACAGCTTCAC | |
| D12 (pASCR3 or pASCR8) | AS_092 | ACTAGAGAAAGAGGGGAAATACGTAATG CAGGATATCTATGCCCTGTCAC | *P. temperata* K122 gDNA 2.918 kb |
| | AS_097 | CAATTTCCCCTTGTGGCGCTTGGTA GATCTGACGGGCAAAGGCTTCC | |
| E1 (pASCR3 or pASCR8) | AS_093 | ACTAGAGAAAGAGGGGAAATACGTAATG ACGTTGCCGTTGTCATTCGCCC | *P. temperata* K122 gDNA 2.933 kb |
| | AS_098 | CAATTTCCCCTTGTGGCGCTTGGTA AATCTGACGGGCAACCGCTTTTTC | |
| E2 (pASCR3 or pASCR8) | AS_099 | ACTAGAGAAAGAGGGGAAATACGTAATG TTTCCATTATCATTTTCGCAAAATAG | *P. temperata* K122 gDNA 2.969 kb |
| | AS_109 | CAATTTCCCCTTGTGGCGCTTGGTA ACGCTGATGTTGGTAAGCCAAAG | |
| E3 (pASCR3 or pASCR8) | AS_100 | ACTAGAGAAAGAGGGGAAATACGTAATG CAGGATATTTATGGGCTTTCGC | *P. temperata* K122 gDNA 2.954 kb |
| | AS_110 | CAATTTCCCCTTGTGGCGCTTGGTA AACTTGTTGTTCATAATCAGCATC | |
| E4 (pASCR3 or pASCR8) | AS_101 | ACTAGAGAAAGAGGGGAAATACGTAATG AATATTCCCTTGAGTTTTGTTC | *P. temperata* K122 gDNA 2.957 kb |
| | AS_111 | CAATTTCCCCTTGTGGCGCTTGGTA CGTTTGATGTTCATAATCCGTATC | |
| E5 (pASCR3 or pASCR8) | AS_102 | ACTAGAGAAAGAGGGGAAATACGTAATG CAGGATATCTATTCACTCTCTC | *P. temperata* K122 gDNA 2.927 kb |
| | AS_112 | CAATTTCCCCTTGTGGCGCTTGGTA AACCTTATGAGCCAAGGCAGTC | |
| E6 (pASCR3 or pASCR8) | AS_103 | ACTAGAGAAAGAGGGGAAATACGTAATG TTACCACTCTCATTTGCTCAAC | *P. temperata* K122 gDNA 2.924 kb |
| | AS_113 | CAATTTCCCCTTGTGGCGCTTGGTA AACTTTGCGGGCCAATGCCATC | |
| A1 (pASCR3, pASCR8 or pASCR12) | AS_001 | ACTAGAGAAAGAGGGGAAATACGTAATG TTAAACAGTTCTAAAAGTATATTG | *A. migulanus* gDNA 1.655 kb |
| | AS_006 | CAATTTCCCCTTGTGGCGCTTGGTA GTCTACCCTCATCCCG | |
| A2 (pASCR3, pASCR8 or pASCR12) | AS_002 | ACTAGAGAAAGAGGGGAAATACGTAATG CAGGATATGTATCGTTTATCTC | *A. migulanus* gDNA 2.930 kb |
| | AS_007 | CAATTTCCCCTTGTGGCGCTTGGTA TTTTGCGTTTGTATTCACAATC | |

| | | | |
|---|---|---|---|
| A3 (pASCR3, pASCR8 or pASCR12) | AS_003 | ACTAGAGAAAGAGGGGAAATACGTAATG GAGTACTATCCTGTATCATCAG | *A. migulanus* gDNA 2.885 kb |
| | AS_008 | CAATTTCCCCTTGTGGCGCTTGGTA TTCGGTTGCTGTACCAAATTC | |
| A4 (pASCR3, pASCR8 or pASCR12) | AS_004 | ACTAGAGAAAGAGGGGAAATACGTAATG GATTACTATCCAGTATCATCTG | *A. migulanus* gDNA 2.915 kb |
| | AS_009 | CAATTTCCCCTTGTGGCGCTTGGTA TTCTGTTCCTATCGATATGGAAC | |
| A5 (pASCR3, pASCR8 or pASCR12) | AS_005 | ACTAGAGAAAGAGGGGAAATACGTAATG GACTATTATCCAGTATCATCAG | *A. migulanus* gDNA 2.894 kb |
| | AS_010 | CAATTTCCCCTTGTGGCGCTTGGTA CTCCCTTGCCATTAAGC | |
| A6 (pASCR3 or pASCR8) | AS_011 | ACTAGAGAAAGAGGGGAAATACGTAATG AAGGACATCTACCCGCTGGCGC | *P. costantinii* gDNA 2.945 kb |
| | AS_017 | CAATTTCCCCTTGTGGCGCTTGGTA CTGCCGACGCGCCAACGCGTCG | |
| A7 (pASCR3 or pASCR8) | AS_012 | ACTAGAGAAAGAGGGGAAATACGTAATG CAGGAAATCTACAGCCTGGCGC | *P. costantinii* gDNA 2.948 kb |
| | AS_018 | CAATTTCCCCTTGTGGCGCTTGGTA TTCGCGACGGGCCAGGGCATCG | |
| A8 (pASCR3 or pASCR8) | AS_013 | ACTAGAGAAAGAGGGGAAATACGTAATG CAGGATATCTACAGCCTGGCGC | *P. costantinii* gDNA 2.882 kb |
| | AS_019 | CAATTTCCCCTTGTGGCGCTTGGTA ACCGCGACTGAGCAAGGCGTCC | |
| A9 (pASCR3 or pASCR8) | AS_014 | ACTAGAGAAAGAGGGGAAATACGTAATG GAGTCCTTGCCGCTGTCGTTTG | *P. costantinii* gDNA 2.930 kb |
| | AS_020 | CAATTTCCCCTTGTGGCGCTTGGTA CACCCCGCTGACCACCGCATCG | |
| A10 (pASCR3 or pASCR8) | AS_015 | ACTAGAGAAAGAGGGGAAATACGTAATG CAGGATTTGCCATTGTCATTTG | *P. costantinii* gDNA 2.945 kb |
| | AS_021 | CAATTTCCCCTTGTGGCGCTTGGTA ATCGCGGCTGGCCACCGCCGAC | |
| A11 (pASCR3 or pASCR8) | AS_016 | ACTAGAGAAAGAGGGGAAATACGTAATG CAAGACATCTATGGCATGACGC | *P. costantinii* gDNA 2.927 kb |
| | AS_022 | CAATTTCCCCTTGTGGCGCTTGGTA GTCGCGGCTGAGCATCACCTGC | |
| Xsto_M1 (pASCR3 or pASCR8) | LP786 | ACTAGAGAAAGAGGGGAAATACGTA ATGTATAAATATCAAGATTATCTGTGTATA C | *X. stockiae* gDNA 2.834 kb |
| | LP787 | CAATTTCCCCTTGTGGCGCTTGGTA AACCGTTTGTACAAAAGCAC | |
| Xsto_M2 (pASCR3 or pASCR8) | LP788 | ACTAGAGAAAGAGGGGAAATACGTAATG GAACCTTTACCGCTATCTTTTTCC | *X. stockiae* gDNA 3.026 kb |
| | LP789 | CAATTTCCCCTTGTGGCGCTTGGTA GGTCGCATGGACAAAGGCTTCGTTC | |
| Xsto_M3 (pASCR3 or pASCR8) | LP790 | ACTAGAGAAAGAGGGGAAATACGTAATG CAGGATATCTATGCACTTTCTCC | *X. stockiae* gDNA 2.966 kb |
| | LP791 | CAATTTCCCCTTGTGGCGCTTGGTA GACTTCATGGGCGAGGACCTCG | |
| B4 (pASCR3 or pASCR8) | AS_031 | ACTAGAGAAAGAGGGGAAATACGTAATG ACTTTTGAATTGTCCCGTTCGC | *C. subtsuage* gDNA 2.906 kb |
| | AS_033 | CAATTTCCCCTTGTGGCGCTTGGTA CTCGCCAGCGGCGCGCGCGGCTTC | |

| | | | |
|---|---|---|---|
| B5 (pASCR3 or pASCR8) | AS_032 | ACTAGAGAAAGAGGGGAAATACGTAATG GAAGAAGTGCTGCAGGCGACGC | *C. subtsuage* gDNA 2.933 kb |
| | AS_034 | CAATTTCCCCTTGTGGCGCTTGGTA GGCGGGCGCCGCCTGCTCCTGC | |
| B6 (pASCR3 or pASCR8) | AS_035 | ACTAGAGAAAGAGGGGAAATACGTAATG TTTGCTGAGTCCTTGCTGGAAAAG | *C. subtsuage* gDNA 1.643 kb |
| | AS_037 | CAATTTCCCCTTGTGGCGCTTGGTA GGCCTCCGCCGCCGGCGCCTGG | |
| B7 (pASCR3 or pASCR8) | AS_036 | ACTAGAGAAAGAGGGGAAATACGTAATG TGCCCGCTGGGGCCGCTGCAGC | gDNA *C. subtsuage* 2.957 kb |
| | AS_038 | CAATTTCCCCTTGTGGCGCTTGGTA CGGCTGTCCGGCCGCGACGCCG | |
| C12 (pASCR3 or pASCR8) | AS_071 | ACTAGAGAAAGAGGGGAAATACGTAATG AGCGCGGCTCCGGAGTCGCCC | *M. xanthus* gDNA 4.394 kb |
| | AS_072 | CAATTTCCCCTTGTGGCGCTTGGTA CTCACCGCGCGCCTGCGCCTGC | |
| F8 (pASCR3 or pASCR8) | AS_135 | ACTAGAGAAAGAGGGGAAATACGTAATG AACGTCGCCAGCATTTTCGCCC | *X. autotrophicus* gDNA 1.529 kb |
| | AS_136 | CAATTTCCCCTTGTGGCGCTTGGTA TTCCGAGCCGCGCTCGCGGCG | |
| F9 (pASCR3 or pASCR8) | AS_137 | ACTAGAGAAAGAGGGGAAATACGTAATG TTTCCCCTGTCGTTCGCGCAGG | *X. autotrophicus* gDNA 3.734 kb |
| | AS_138 | CAATTTCCCCTTGTGGCGCTTGGTA TTGCGGCGCGGCGGGGGC | |
| F10 (pASCR3 or pASCR8) | AS_139 | ACTAGAGAAAGAGGGGAAATACGTAATG CACGAGCCGTTCCCACTCACCC | *X. autotrophicus* gDNA 3.707 kb |
| | AS_140 | CAATTTCCCCTTGTGGCGCTTGGTA CGCATCCGCCACAGGCAGGACG | |
| epoxy_M1 (pASCR3 or pASCR8) | LP730 | ACTAGAGAAAGAGGGGAAATACGTAATG GCCACAACACTGCCGCACGAGCAG | *G. coeruleoviolacea* gDNA 4.181 kb |
| | LP731 | CAATTTCCCCTTGTGGCGCTTGGTA GCCCTCGGCGGTGTCCAGTTTG | |
| pASCR3_epoxy_M2 (nested PCR) | LP732 | ACTAGAGAAAGAGGGGAAATACGTAATG GACCGGGTGCCGCTGTCGGACG | *G. coeruleoviolacea* gDNA 5.030 kb |
| | LP737 | TCATCCGGGTGTGGTCGAACCGGGCGTC | |
| epoxy_M2 (pASCR3 or pASCR8) | LP732 | ACTAGAGAAAGAGGGGAAATACGTAATG GACCGGGTGCCGCTGTCGGACG | pASCR3_epoxy_M2 (nested PCR) 3.530 kb |
| | LP733 | CAATTTCCCCTTGTGGCGCTTGGTA GTGCCGCTGCCGCAACGGCACAC | |
| pASCR3_epoxy_M3 (nested PCR) | LP734 | ACTAGAGAAAGAGGGGAAATACGTAATG GAGTCCGCACCCGCGTCCAGCTTC | *G. coeruleoviolacea* gDNA 3.657 kb |
| | LP738 | GCTTCGGTCCCGGTGTAGCACCGGCCCAGTTCG | |
| epoxy_M3 (pASCR3 or pASCR8) | LP734 | ACTAGAGAAAGAGGGGAAATACGTAATG GAGTCCGCACCCGCGTCCAGCTTC | pASCR3_epoxy_M3 (nested PCR) 2.948 kb |
| | LP733 | CAATTTCCCCTTGTGGCGCTTGGTA GTGCCGCTGCCGCAACGGCACAC | |

| | | | |
|---|---|---|---|
| epoxy_M4 (pASCR3 or pASCR8) | LP735 | ACTAGAGAAAGAGGGGAAATACGTAATG GACACCTTCCCGGCATCCGGGTTC | *G. coeruleoviolacea* gDNA 3.689 kb |
| | LP736 | CAATTTCCCCTTGTGGCGCTTGGTA CGGCGCCGCCCCCACCAGGTCG | |
| glbS_M1 (pASCR3 or pASCR8) | LP650 | ACTAGAGAAAGAGGGGAAATACGTAATG TATTACCCCCTGACCGCAGCAC | *T. laumondii* TTO1 gDNA 2.966 kb |
| | LP647 | CAATTTCCCCTTGTGGCGCTTGGTA ATTGATTACTACACTGTTCCACATG | |
| glbS_M2 (pASCR3 or pASCR8) | LP651 | ACTAGAGAAAGAGGGGAAATACGTAATG CCGGCTCAACGCAGGTTATGGC | *T. laumondii* TTO1 gDNA 2.975 kb |
| | LP648 | CAATTTCCCCTTGTGGCGCTTGGTA AGACTCCTGATTTTCTGGTAGTGAG | |
| glbS_M3 (pASCR3 or pASCR8) | LP652 | ACTAGAGAAAGAGGGGAAATACGTAATG CTCTCATTCGGTCAACGCAGTTTG | *T. laumondii* TTO1 gDNA 2.891 kb |
| | LP649 | CAATTTCCCCTTGTGGCGCTTGGTA AGGAGAAGATTGATGCTGACCGGC | |
| odlS_M1 (pASCR3 or pASCR8) | LP760 | ACTAGAGAAAGAGGGGAAATACGTAATG TTTCTAGATAAAGTCGGGCAGC | *X. nematophila* gDNA 3.023 kb |
| | LP761 | CAATTTCCCCTTGTGGCGCTTGGTA TTCCTGCTGACTGTAATCTTGTTTTTG | |
| odlS_M2 (pASCR3 or pASCR8) | LP762 | ACTAGAGAAAGAGGGGAAATACGTAATG CAAGATATTTACCCATTATCAC | *X. nematophila* gDNA 2.960 kb |
| | LP763 | CAATTTCCCCTTGTGGCGCTTGGTA TTTGGCATGATGATTAATGAAACC | |
| odlS_M4 (pASCR3 or pASCR8) | LP765 | ACTAGAGAAAGAGGGGAAATACGTAATG ATGGATATGCTGAAATTAGTCC | *X. nematophila* gDNA 3.161 kb |
| | LP766 | CAATTTCCCCTTGTGGCGCTTGGTA GGCTTTAACAGCAATATCACTGTC | |
| odlS_M5 (pASCR3 or pASCR8) | LP739 | ACTAGAGAAAGAGGGGAAATACGTAATG ATTATTGAACGCAAAACAC | *X. nematophila* gDNA 2.981 kb |
| | LP601 | CAATTTCCCCTTGTGGCGCTTGGTA TTCAATTTTCCGTTGAGC | |
| odlS_M6 (pASCR3 or pASCR8) | LP767 | ACTAGAGAAAGAGGGGAAATACGTAGGC ATTTATCCTTTGGCACC | *X. nematophila* gDNA 2.941 kb |
| | LP768 | AATTTCCCCTTGTGGCGCTTGGTATTCGG CTTTGACAAAGTTATCTG | |
| odlS_M7 (pASCR3 or pASCR8) | LP769 | ACTAGAGAAAGAGGGGAAATACGTAATG GGGATTATGAACCTAAAAGAC | *X. nematophila* gDNA 3.176 kb |
| | LP602 | CAATTTCCCCTTGTGGCGCTTGGTA AGGCTCTTCGACGATATTTTCC | |
| odlS_M8 (pASCR3 or pASCR8) | LP770 | ACTAGAGAAAGAGGGGAAATACGTAATG AATATTGCGGCATTTTTAGCTG | *X. nematophila* gDNA 3.154 kb |
| | LP771 | AATTTCCCCTTGTGGCGCTTGGTA ACCCTCTGGTAATAATTCTTGCG | |
| odlS_M9 (pASCR3 or pASCR8) | LP772 | ACTAGAGAAAGAGGGGAAATACGTAATG GAACGATTTCCCTTGTCGTTTG | *X. nematophila* gDNA 2.981 kb |
| | LP773 | CAATTTCCCCTTGTGGCGCTTGGTA TTCAGTACGGGCAAAAGCGC | |
| odlS_M10 (pASCR3 or pASCR8) | LP774 | ACTAGAGAAAGAGGGGAAATACGTAATG TATCACGCTTTTTCATTACTCAC | *X. nematophila* gDNA 3.002 kb |
| | LP603 | CAATTTCCCCTTGTGGCGCTTGGTA TGATCCGCGCACAAGACAGGG | |

**Table S5.** $^{1}H$ NMR and $^{13}C$ NMR data of compound **1** ($\delta$ in ppm).

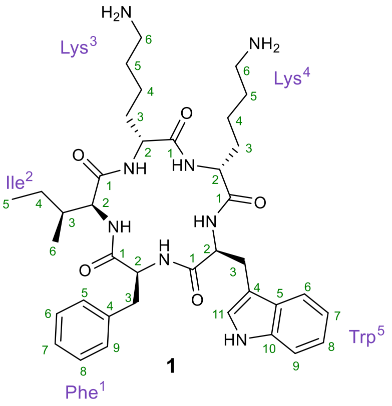

1

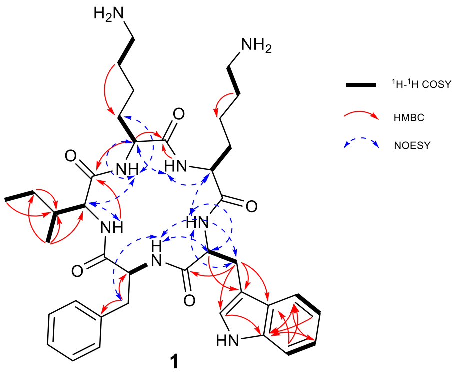

1

| subunit | position | $\delta_C{}^a$, type | $\delta_H{}^b$, (*J* in Hz) | subunit | position | $\delta_C{}^a$, type | $\delta_H{}^b$, (*J* in Hz) |
|---|---|---|---|---|---|---|---|
| Phe[1] | 1 | 170.7-171,0 C | | Lys[4] | 1 | 170.7-171.0, C | |
| | 2 | 58.0, CH | 4.08, overlap | | 2 | 52.2, CH | 4.07, overlap |
| | 2-NH | | 8.54, d (8.66) | | 2-NH | | 7.40, d (7.27) |
| | 3a | 36.6, $CH_2$ | 3.00, dd (13.63, 9.89) | | 3a | 30.6, $CH_2$ | 1.46, overlap |
| | 3b | | 3.00, m | | 3b | | 1.53, overlap |
| | 4 | 137.5, C | | | 4a | 22.1, $CH_2$ | 1.06, m |
| | 5 | 129.0, CH | 7.08, overlap | | 4b | | 0.99, m |
| | 6 | 128.0, CH | 7.15, overlap | | 5a | 27.1, $CH_2$ | 1.39, overlap |
| | 7 | 126.1, CH | 7.10, m | | 5b | | 1.39, overlap |
| | 8 | 128.0, CH | 7.15, overlap | | 6a | 38.4, $CH_2$ | 2.57, overlap |
| | 9 | 129.0, CH | 7.08, overlap | | 6b | | 2.57, overlap |
| Ile[2] | 1 | 170.9, C | | | 6-NH | not found | |
| | 2 | 56.8, CH | 4.04, overlap | Trp[5] | 1 | 171.1, C | |
| | 2-NH | | 8.09, d (7.77) | | 2 | 55.6, CH | 4.18, dt (7.18) |
| | 3 | 37.2, CH | 1.64, m | | 2-NH | | 8.26, brs |
| | 4a | 24.6, $CH_2$ | 1.297, overlap | | 3a | 26.6, $CH_2$ | 2.82, overlap |
| | 4b | | 0.92, m | | 3b | | 2.82, overlap |
| | 5 | 11.3, $CH_3$ | 0.77, overlap | | 4 | 109.6, C | |
| | 6 | 14.9, $CH_3$ | 0.76, overlap | | 5 | 126.9, C | |
| Lys[3] | 1 | 171.8, C | | | 6 | 118.0, CH | 7.45, d (7.87) |
| | 2 | 55.4, CH | 3.83, dt (7.01) | | 7 | 118.2, CH | 6.91, m |
| | 2-NH | | 8.63, d (6.29) | | 8 | 120.8, CH | 6.98, m |
| | 3a | 29.7, $CH_2$ | 1.54, overlap | | 9 | 111.2, CH | 7.26, d (8.1) |
| | 3b | | 1.54, overlap | | 10 | 136.0, C | |
| | 4a | 22.4, $CH_2$ | 1.24, overlap | | 10-NH | | 10.84 |
| | 4b | | 1.30, m | | 11 | 123.2, CH | 6.89,brs |
| | 5a | 27.0, $CH_2$ | 1.45, overlap | | | | |
| | 5b | | 1.45, overlap | | | | |
| | 6a | 38.4, $CH_2$ | 2.64, overlap | | | | |
| | 6b | | 2.65, overlap | | | | |
| | 6-NH | not found | | | | | |

[a]Recorded at 125 MHz, in DMSO-*d*6. [b]Recorded at 500 MHz, in DMSO-*d*6.

**Table S6.** ¹H NMR and ¹³C NMR data of compound **5** ($\delta$ in ppm).

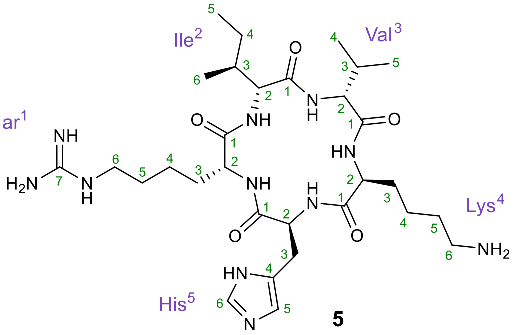

**5**

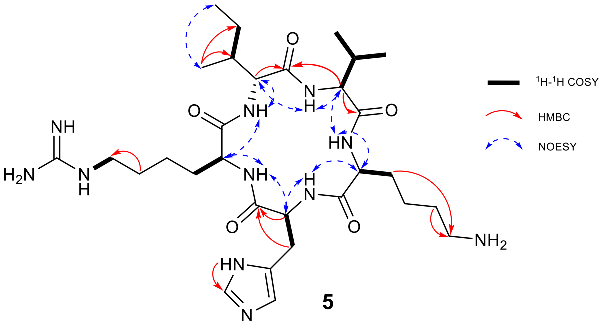

**5**

| subunit | position | $\delta_C$[a], type | $\delta_H$[b], (*J* in Hz) | subunit | position | $\delta_C$[a], type | $\delta_H$[b], (*J* in Hz) |
|---|---|---|---|---|---|---|---|
| Homo-Arg[1] | 1 | 171.7-172.2, C | | Lys[4] | 1 | 171.7-172.2, C | |
| | 2 | 55.4, CH | 3.98 overlap | | 2 | 55.4, CH | 3.95, overlap |
| | 2-NH | | 8.26, brs | | 2-NH | | 8.57, d (6.75) |
| | 3a | 30.9, $CH_2$ | 1.63, overlap | | 3a | 30.5, $CH_2$ | 1.60, overlap |
| | 3b | | 1.99, overlap | | 3b | | 1.60, overlap |
| | 4a | 23.0, $CH_2$ | 1.26, overlap | | 4a | 23.2, $CH_2$ | 1.33, overlap |
| | 4b | | 1.26, overlap | | 4b | | 1.33, overlap |
| | 5a | 28.5, $CH_2$ | 1.48, overlap | | 5a | 27.7, $CH_2$ | 1.55, overlap |
| | 5b | | 1.48, overlap | | 5b | | 1.55, overlap |
| | 6a | 40.9, $CH_2$ | 3.08, overlap | | 6a | 39.1, $CH_2$ | 2.76, overlap |
| | 6b | | 3.08, overlap | | 6b | | 2.76, overlap |
| | 6-NH | | 8.78, s | His[5] | 1 | 171.1, C | |
| | 7 | 157.9, C | | | 2 | 53.7, CH | 4.44, dt (7.2,?) |
| Ile[2] | 1 | 171.7, C | | | 2-NH | | 7.49, d (7.3) |
| | 2 | 61.7, CH | 3.83, m | | 3a | 29.6, $CH_2$ | 2.92, overlap |
| | 2-NH | | 8.39, d (9.1) | | 3b | | 2.92, overlap |
| | 3 | 36.3, CH | 2.06, m | | 4 | Not found | |
| | 4a | 26.1 $CH_2$ | 1.11, overlap | | 5 | 129.5, CH | 7.75, m |
| | 4b | | 1.48, m | | 6 | 135.0, CH | 7.54 |
| | 5 | 11.6, $CH_3$ | 0.92, overlap | | 6-NH | 6.71 | |
| | 6 | 15.6, $CH_3$ | 0.82, overlap | | | | |
| Val[3] | 1 | 171.2, C | | | | | |
| | 2 | 58.2, CH | 4.15, dd (7.5,?) | | | | |
| | 2-NH | | 8.04, d (7.9) | | | | |
| | 3 | 31.4, CH | 1.97, m | | | | |
| | 4 | 18.9, $CH_3$ | 0.89, overlap | | | | |
| | 5 | 19.34, $CH_3$ | 0.90, overlap | | | | |

[a]Recorded at 125 MHz, in DMSO-*d*6. [b]Recorded at 500 MHz, in DMSO-*d*6.

**Table S7.** $^{1}$H NMR and $^{13}$C NMR data of compound **6** ($\delta$ in ppm).

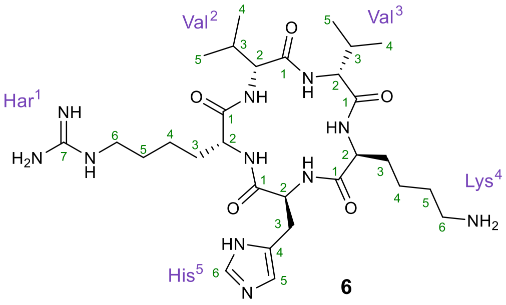

**6**

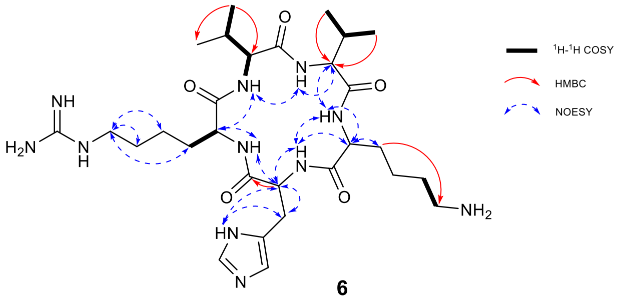

**6**

| subunit | position | $\delta_C$[a], type | $\delta_H$[b], (J in Hz) | subunit | position | $\delta_C$[a], type | $\delta_H$[b], (J in Hz) |
|---|---|---|---|---|---|---|---|
| Homo-Arg[1] | 1 | 170.5, C | | Lys[4] | 1 | 171.4, C | |
| | 2 | 55.0, CH | 3.86, overlap | | 2 | 55.1, CH | 3.81, overlap |
| | 2-NH | | 7.98, brs | | 2-NH | | 8.44, d (6.65) |
| | 3a | 30.2, $CH_2$ | 1.50, overlap | | 3a | 29.9, $CH_2$ | 1.47, overlap |
| | 3b | | 1.50, overlap | | 3b | | 1.47, overlap |
| | 4a | 22.3, $CH_2$ | 1.20, overlap | | 4a | 22.6, $CH_2$ | 1.13, overlap |
| | 4b | | 1.21, overlap | | 4b | | 1.13, overlap |
| | 5a | 27.8, $CH_2$ | 1.36, overlap | | 5a | 27.3, $CH_2$ | 1.41, overlap |
| | 5b | | 1.36, overlap | | 5b | | 1.41, overlap |
| | 6a | 40.2, $CH_2$ | 2.96, overlap | | 6a | 38.6, $CH_2$ | 2.62, overlap |
| | 6b | | 2.97, overlap | | 6b | | 2.62, overlap |
| | 7 | 157.0, C | | His[5] | 1 | 170.5, C | |
| | 7-NH | | 8.41, m | | 2 | 52.5 CH | 4.13, dt (7.2) |
| Val[2] | 1 | 170.9, C | | | 2-NH | | 7.36, brs |
| | 2 | 62.9, CH | 3.56, m | | 3a | Not found | 2.79, overlap |
| | 2-NH | | 8.19, d (9.13) | | 3b | | 2.79, overlap |
| | 3 | 29.4, CH | 2.14, m | | 4 | 128.1, C | |
| | 4 | 19.21, $CH_3$ | 0.72, d (6.5) | | 5 | 129.0, CH | 7.2, m |
| | 5 | 19.38, $CH_3$ | 0.8, d (6.7) | | 6 | 134.4, CH | Not found |
| Val[3] | 1 | 170.7, C | | | | 6-NH | 6.59 |
| | 2 | 57.5, CH | 4.02, dd (7.55, 7.78) | | | | |
| | 2-NH | | 7.90, d (7.78) | | | | |
| | 3 | 30.8, CH | 1.83, m | | | | |
| | 4 | 18.35, $CH_3$ | 0.78, d/overlap | | | | |
| | 5 | 18.35, $CH_3$ | 0.78, d/overlap | | | | |

[a]Recorded at 125 MHz, in DMSO-$d_6$. [b]Recorded at 500 MHz, in DMSO-$d_6$.

**Table S8.** $^{1}$H NMR and $^{13}$C NMR data of compound **7a** (*δ* in ppm).

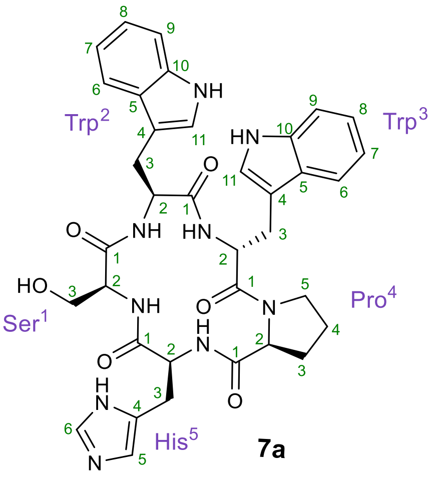

**7a**

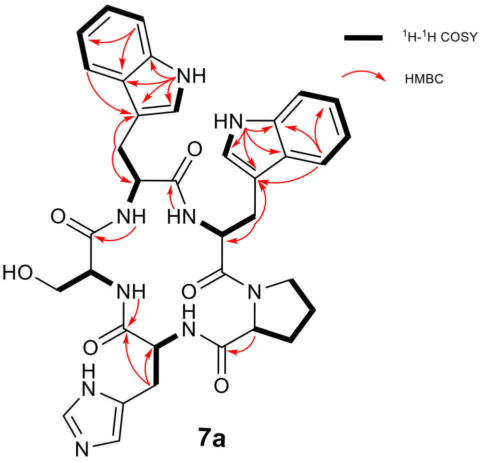

**7a**

| subunit | position | $\delta_C{}^a$, type | $\delta_H{}^b$, (*J* in Hz) | subunit | position | $\delta_C{}^a$, type | $\delta_H{}^b$, (*J* in Hz) |
|---|---|---|---|---|---|---|---|
| Ser$^1$ | 1 | 170.3, C | | Pro$^4$ | 1 | 171.4, C | |
| | 2 | 57.3, CH | 4.04, m | | 2 | 60.7, CH | 3.93, dd (8.1, 3.7) |
| | 2-NH | | 8.28, t (6.2) | | 3a | 29.0, $CH_2$ | 1.51, overlap |
| | 3a | 60.6, $CH_2$ | 3.71, dd (11.0, 5.3) | | 3b | | 1.41, m |
| | 3b | | 3.56, dd (11.0, 5.3) | | 4a | 23.7, $CH_2$ | 1.51, overlap |
| Trp$^2$ | 1 | 170.9, C | | | 4b | | 1.22, m |
| | 2 | 53.4, CH | 4.51, m | | 5a | 46.4, $CH_2$ | 3.51, overlap |
| | 2-NH | | 7.88, dd (8.1, 3.8) | | 5b | | 2.49, overlap |
| | 3a | 26.5, $CH_2$ | 3.15, dd (14.5, 7.2) | His$^5$ | 1 | 170.1, C | |
| | 3b | | 2.96, m | | 2 | 53.6, CH | 4.51, m |
| | 4 | 110.4, C | | | 2-NH | | 7.12, overlap |
| | 5 | 127.5, C | | | 3a | 29.8, $CH_2$ | 2.88, overlap |
| | 6 | 118.5, CH | 7.54, d (7.9) | | 3b | | 2.80, overlap |
| | 7 | 118.2, CH | 6.94, m | | 4 | 129.4, C | |
| | 8 | 120.9, CH | 7.06, overlap | | 5 | 118.3, CH | 7.06, overlap |
| | 9 | 111.3, CH | 7.33, overlap | | 6 | 134.8, CH | 7.45, overlap |
| | 10 | 136.0, C | | | | | |
| | 10-NH | | 10.75, d (2.4) | | | | |
| | 11 | 123.2, CH | 7.06, overlap | | | | |
| Trp$^3$ | 1 | 171.0, C | | | | | |
| | 2 | 52.4, CH | 4.61, ddd (9.2, 7.2, 5.8) | | | | |
| | 2-NH | | 8.46, d (7.2) | | | | |
| | 3a | 27.3, $CH_2$ | 3.02, m | | | | |
| | 3b | | 2.88, overlap | | | | |
| | 4 | 109.3, C | | | | | |
| | 5 | 127.0, C | | | | | |
| | 6 | 118.3, CH | 7.45, d (7.3) | | | | |
| | 7 | 118.4, CH | 6.97, m | | | | |
| | 8 | 121.1, CH | 7.06, overlap | | | | |
| | 9 | 111.5, CH | 7.33, overlap | | | | |
| | 10 | 136.1, C | | | | | |
| | 10-NH | | 10.81, d (2.4) | | | | |
| | 11 | 123.8, CH | 7.12, overlap | | | | |

$^a$Recorded at 125 MHz, in DMSO-*d*$_6$. $^b$Recorded at 500 MHz, in DMSO-*d*$_6$.

**Table S9.** $^{1}H$ NMR and $^{13}C$ NMR data of compound **7b** ($\delta$ in ppm).

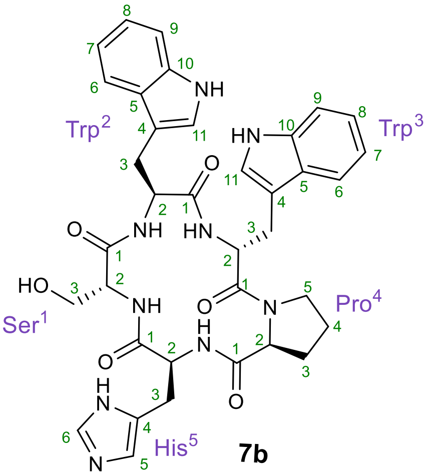

**7b**

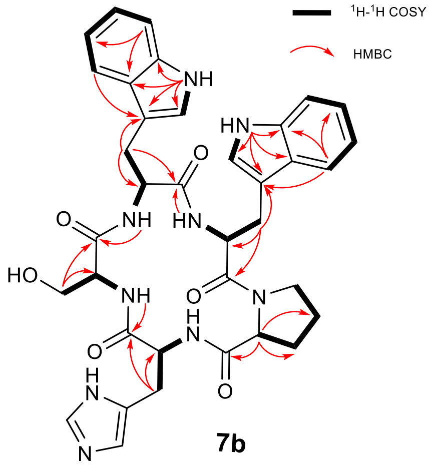

**7b**

| subunit | position | $\delta_C$[a], type | $\delta_H$[b], (*J* in Hz) | subunit | position | $\delta_C$[a], type | $\delta_H$[b], (*J* in Hz) |
|---|---|---|---|---|---|---|---|
| Ser[1] | 1 | 170.5, C | | Pro[4] | 1 | 171.0, C | |
| | 2 | 54.4, CH | 4.28, td (7.7, 5.7) | | 2 | 60.9, CH | 3.87, dd (7.9, 4.5) |
| | 2-NH | | 7.57, d (7.8) | | 3a | 28.7, $CH_2$ | 1.52, overlap |
| | 3a | 59.3, $CH_2$ | 3.63, m | | 3b | | 1.42, d (5.6) |
| | 3b | | 3.42, m | | 4a | 23.7, $CH_2$ | 1.52, overlap |
| Trp[2] | 1 | 171.3, C | | | 4b | | 1.17, m |
| | 2 | 52.3, CH | 4.74, td (8.5, 6.1) | | 5a | 46.2, $CH_2$ | 3.47, overlap |
| | 2-NH | | 8.01, d (9.0) | | 5b | | 2.53, overlap |
| | 3a | 26.1, $CH_2$ | 3.13, dd (14.9, 6.0) | His[5] | 1 | 170.4, C | |
| | 3b | | 2.86, dd (14.9, 8.2) | | 2 | 52.2, CH | 4.41, q (7.5) |
| | 4 | 110.1, C | | | 2-NH | | 7.53, overlap |
| | 5 | 127.3, C | | | 3a | 28.1, $CH_2$ | 3.02, overlap |
| | 6 | 118.3, CH | 7.53, overlap | | 3b | | |
| | 7 | 118.1, CH | 6.94, overlap | | 4 | 128.1, C | |
| | 8 | 120.8, CH | 7.06, overlap | | 5 | 118.2, CH | 7.53, overlap |
| | 9 | 111.3, CH | 7.34, overlap | | 6 | 134.5, CH | 7.44, overlap |
| | 10 | 136.0, C | | | | | |
| | 10-NH | | 10.80, d (2.4) | | | | |
| | 11 | 123.3, CH | 7.13, d (2.3) | | | | |
| Trp[3] | 1 | 171.5, C | | | | | |
| | 2 | 52.8, CH | 4.56, dt (8.8, 6.4) | | | | |
| | 2-NH | | 8.34, d (7.0) | | | | |
| | 3a | 27.4, $CH_2$ | 3.02, overlap | | | | |
| | 3b | | 2.96, dd (14.1, 5.9) | | | | |
| | 4 | 109.1, C | | | | | |
| | 5 | 127.1, C | | | | | |
| | 6 | 118.0, CH | 7.41, d (7.9) | | | | |
| | 7 | 118.2, CH | 6.96, overlap | | | | |
| | 8 | 121.0, CH | 7.06, overlap | | | | |
| | 9 | 111.4, CH | 7.33, overlap | | | | |
| | 10 | 136.0, C | | | | | |
| | 10-NH | | 10.87, d (2.4) | | | | |
| | 11 | 124.0, CH | 7.20, d (2.4) | | | | |

[a]Recorded at 125 MHz, in DMSO-*d*6. [b]Recorded at 500 MHz, in DMSO-*d*6.

#### 3 Supplementary Figures

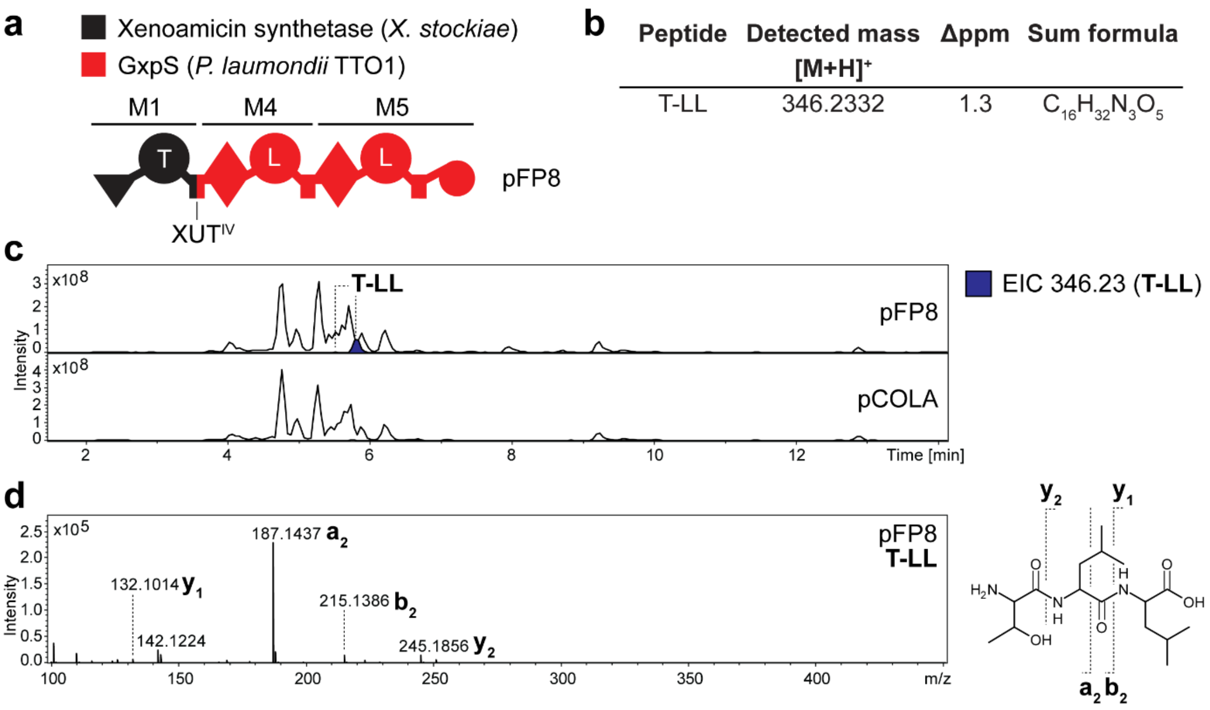

**Figure S1.** HPLC-MS/MS data referred to production of pFP8. (**a**) Schematic representation of of the NRPS encoded by plasmid pFP8 described by Bozyhüyük *et al.* 2024.[1] Domain origin is depicted in the legend. Indication of the utilized fusion site. A domain specitifity is indicated by the one-letter amino acid code. (**b**) Table of HPLC-MS/MS data indicating peptide side product sequence, the detected mass with the corresponding error and the sum formula. (**c**) HPLC-MS/MS data of *E. coli* DH10B::*mtaA* expressing pFP8 or an empty vector control. Base Peak Chromatogram (BPC, top) with Extracted Ion Chromatogram (EIC, below with colors according to the depicted legend) of **T-LL** (*m/z* $[M+H]^+$ = 346.23). (**d**) The $MS^2$ data for the respective compound is displayed, along with indications of the assigned fragments.

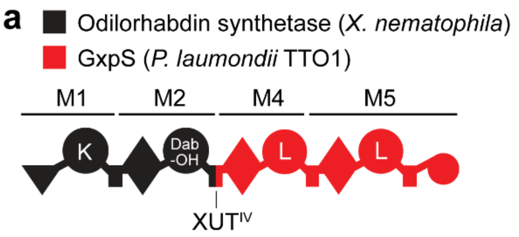

**b**

| Peptide | Detected mass [M+H]+ | Δppm | Sum formula |
|---|---|---|---|
| Dab-LL | 345.2493 | 1.0 | $C_{16}H_{33}N_4O_4$ |
| Dab-OH-LL | 361.2437 | 2.4 | $C_{16}H_{33}N_4O_5$ |

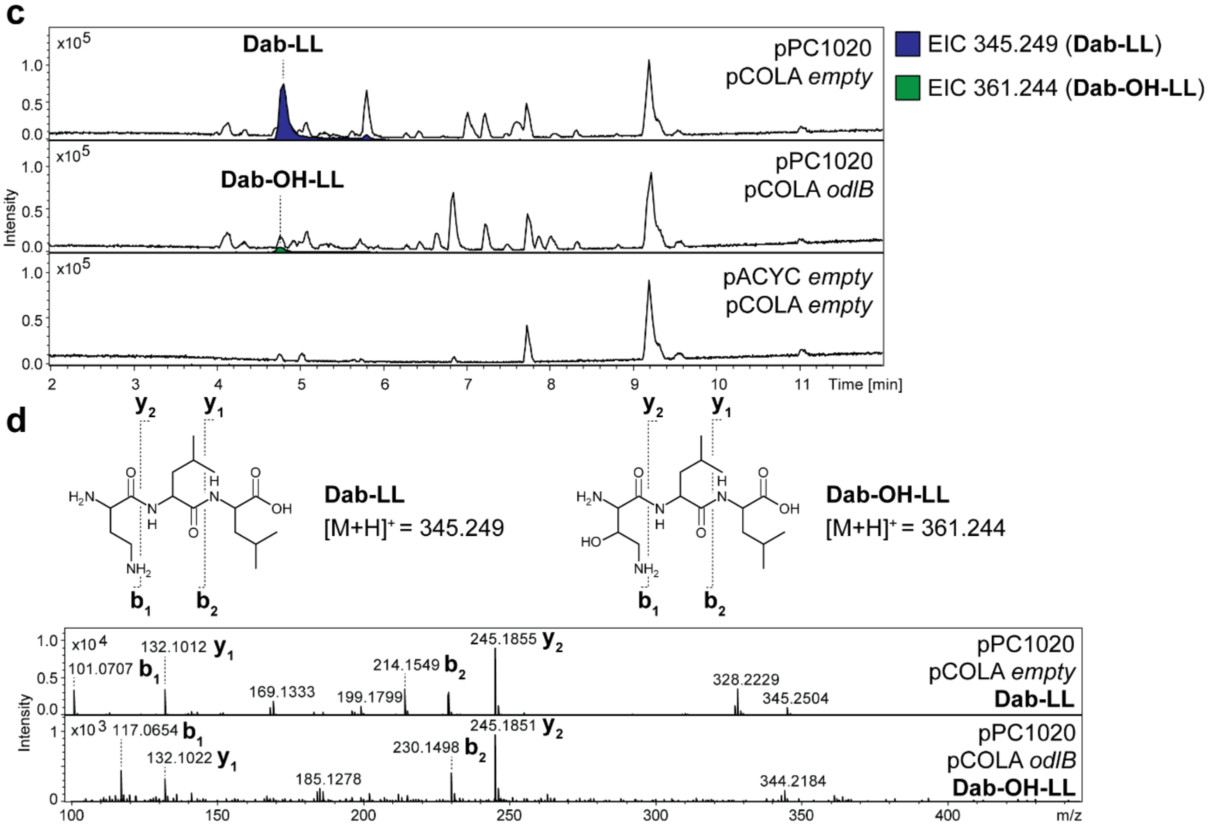

**Figure S2.** HPLC-MS/MS data referred to production of pPC1020. (**a**) Schematic representation of the NRPS encoded by plasmid pPC1020 described by Präve et al. 2024.[5] Domain origin is depicted in the domain legend. Indication of the utilized fusion site. A domain specitifity is indicated by the one-letter amino acid code (Dab = diamino butyric acid; Dab-OH = β-hydroxy diamino butyric acid). (**b**) Table of HPLC-MS/MS data indicating peptide side product sequences, the detected masses with the corresponding errors and the sum formulas. (**c**) HPLC-MS/MS data of *E. coli* DH10B::*mtaA* co-expressing pPC1020 with either an empty vector or *odlB*. Two empty vectors served as negative control. Base Peak Chromatogram (BPC, top) with Extracted Ion Chromatogram (EIC, below with colors according to the depicted legend) of **Dab-LL** (*m/z* $[M+H]^+$ = 345.24) and **Dab-OH-LL** (*m/z* $[M+H]^+$ = 361.24). (**d**) The $MS^2$ data for the respective compounds are displayed, along with indications of the assigned fragments.

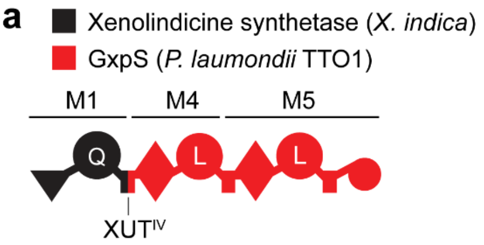

**b**

| Peptide | Detected mass $[M+H]^+$ | Δppm | Sum formula |
|---|---|---|---|
| $C_{14}$-Q-LL | 583.4427 | 0.3 | $C_{31}H_{59}N_4O_6$ |
| Q-LL | 373.2443 | 0.8 | $C_{17}H_{33}N_4O_5$ |

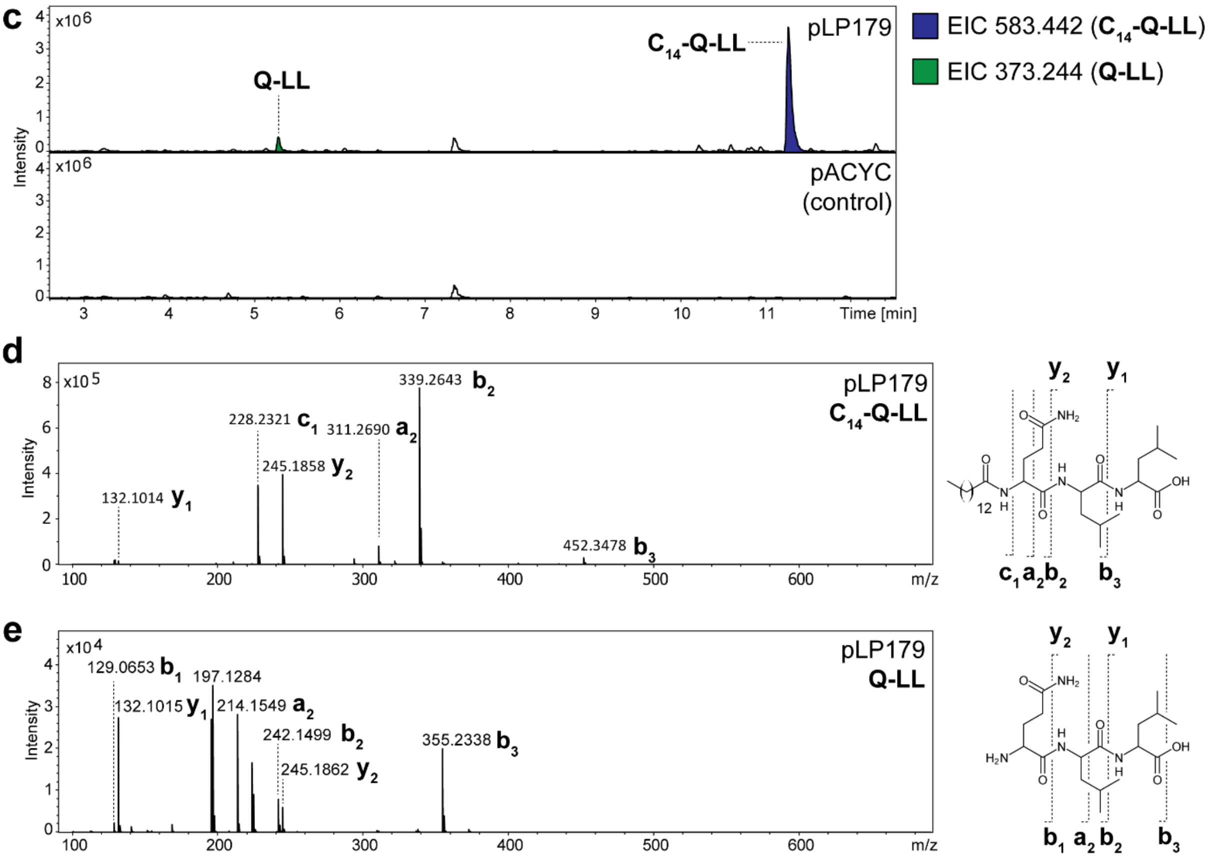

**Figure S3.** HPLC-MS/MS data referred to production of pLP179. (**a**) Schematic representation of the NRPS encoded by plasmid pLP179. Domain origin is depicted in the legend. Indication of the utilized fusion site. A domain specitifity is indicated by the one-letter amino acid code. (**b**) Table of HPLC-MS/MS data indicating peptide main and side product sequence, the detected masses with the corresponding errors and the sum formulas. (**c**) HPLC-MS/MS data of *E. coli* DH10B::*mtaA* expressing pLP179 or an empty vector control. Base Peak Chromatogram (BPC, top) with Extracted Ion Chromatogram (EIC, below with colors according to the depicted legend) of **$C_{14}$-Q-LL** (*m/z* $[M+H]^+$ = 583.442) and **Q-LL** (*m/z* $[M+H]^+$ = 373.244). (**d-e**) The $MS^2$ data for the respective compound is displayed, along with indications of the assigned fragments.

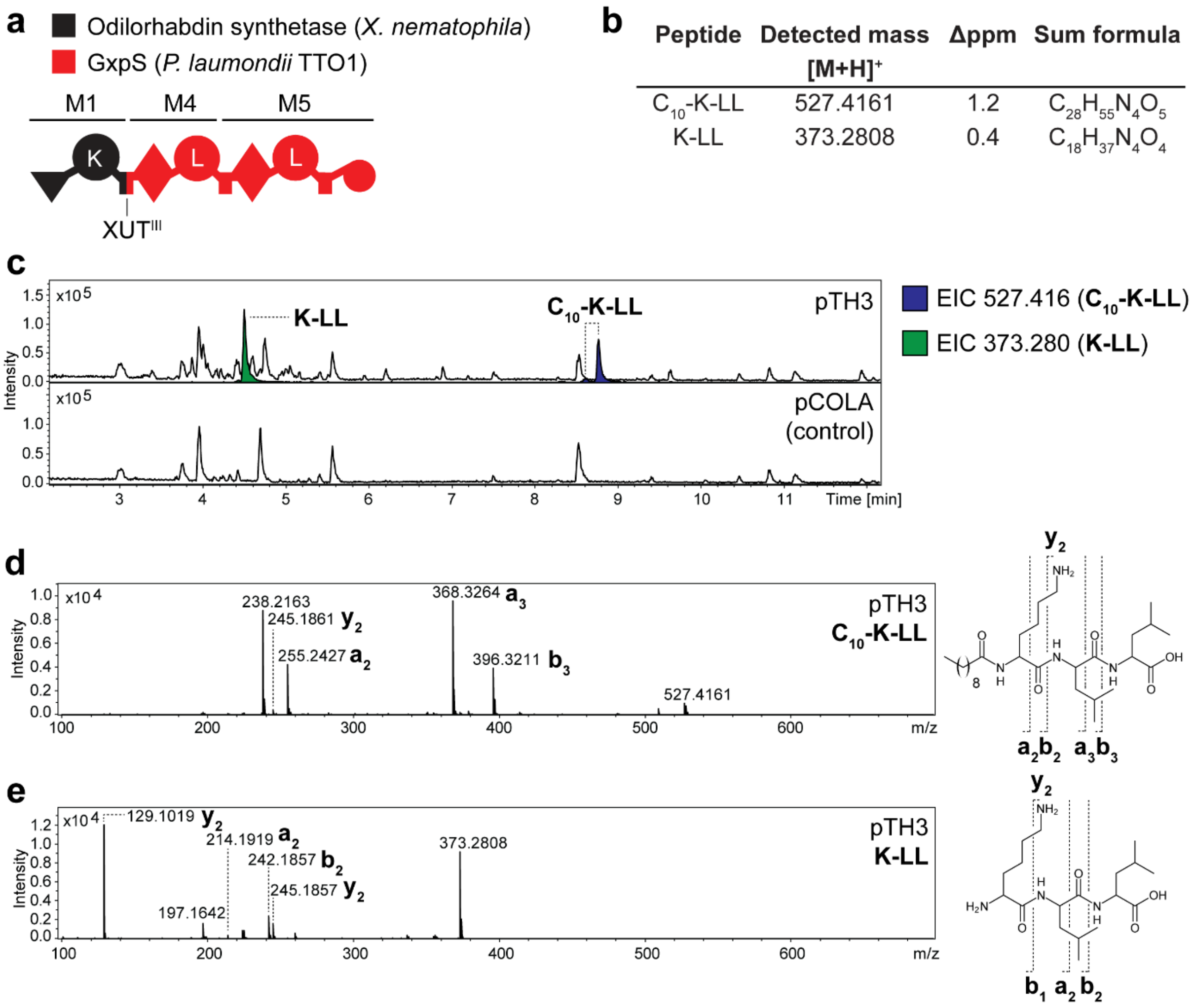

| Peptide | Detected mass [M+H]$^+$ | Δppm | Sum formula |
|---|---|---|---|
| $C_{10}$-K-LL | 527.4161 | 1.2 | $C_{28}H_{55}N_4O_5$ |
| K-LL | 373.2808 | 0.4 | $C_{18}H_{37}N_4O_4$ |

**Figure S4.** HPLC-MS/MS data referred to production of pTH3. (**a**) Schematic representation of the NRPS encoded by plasmid pTH3. Domain origin is depicted in the legend. Indication of the utilized fusion site. A domain specitifity is indicated by the one-letter amino acid code. (**b**) Table of HPLC-MS/MS data indicating peptide main and side product sequence, the detected masses with the corresponding errors and the sum formulas. (**c**) HPLC-MS/MS data of *E. coli* DH10B::*mtaA* expressing pTH3 or an empty vector control. Base Peak Chromatogram (BPC, top) with Extracted Ion Chromatogram (EIC, below with colors according to the depicted legend) of **$C_{10}$-K-LL** (*m/z* [M+H]$^+$ = 527.416) and **K-LL** (*m/z* [M+H]$^+$ = 373.280). (**d-e**) The $MS^2$ data for the respective compound is displayed, along with indications of the assigned fragments.

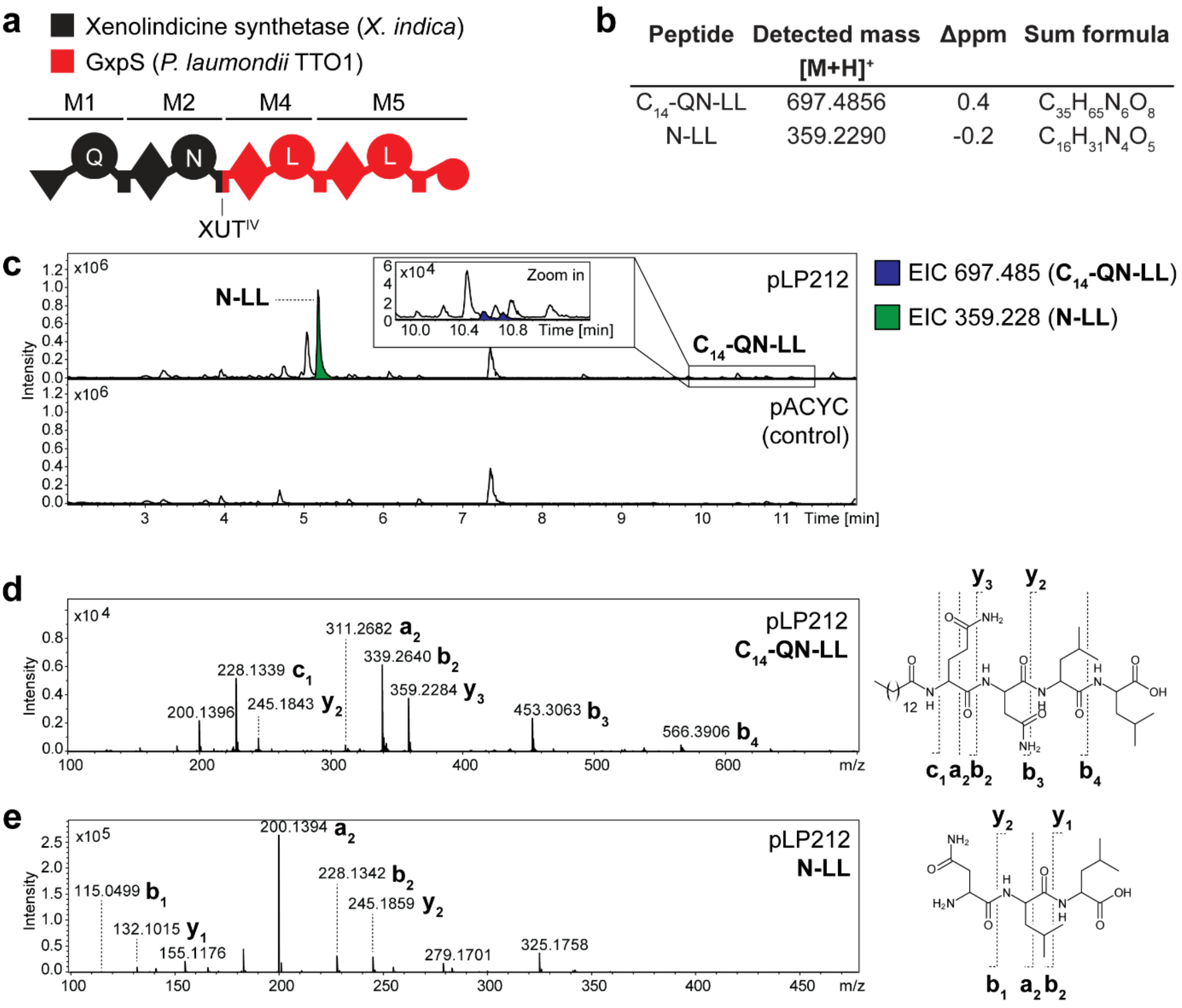

b

| Peptide | Detected mass [M+H]+ | Δppm | Sum formula |
|---|---|---|---|
| $C_{14}$-QN-LL | 697.4856 | 0.4 | $C_{35}H_{65}N_6O_8$ |
| N-LL | 359.2290 | -0.2 | $C_{16}H_{31}N_4O_5$ |

**Figure S5.** HPLC-MS/MS data referred to production of pLP212. (**a**) Schematic representation of the NRPS encoded by plasmid pLP212. Domain origin is depicted in the legend. Indication of the utilized fusion site. A domain specitifity is indicated by the one-letter amino acid code. (**b**) Table of HPLC-MS/MS data indicating peptide main and side product sequence, the detected masses with the corresponding errors and the sum formulas. (**c**) HPLC-MS/MS data of *E. coli* DH10B::*mtaA* expressing pLP212 or an empty vector control. Base Peak Chromatogram (BPC, top) with Extracted Ion Chromatogram (EIC, below with colors according to the depicted legend) of **$C_{14}$-QN-LL** (*m/z* $[M+H]^+$ = 697.485) and **N-LL** (*m/z* $[M+H]^+$ = 359.228). (**d-e**) The $MS^2$ data for the respective compound is displayed, along with indications of the assigned fragments.

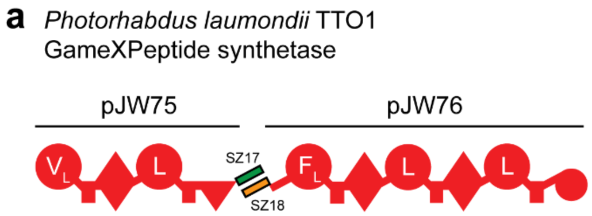

**b**

| Peptide | Detected mass [M+H]+ | Δppm | Sum formula |
|---|---|---|---|
| F-LL | 392.2535 | 2.2 | $C_{21}H_{34}N_3O_4$ |
| L-LL | 358.2694 | 1.7 | $C_{18}H_{36}N_3O_4$ |

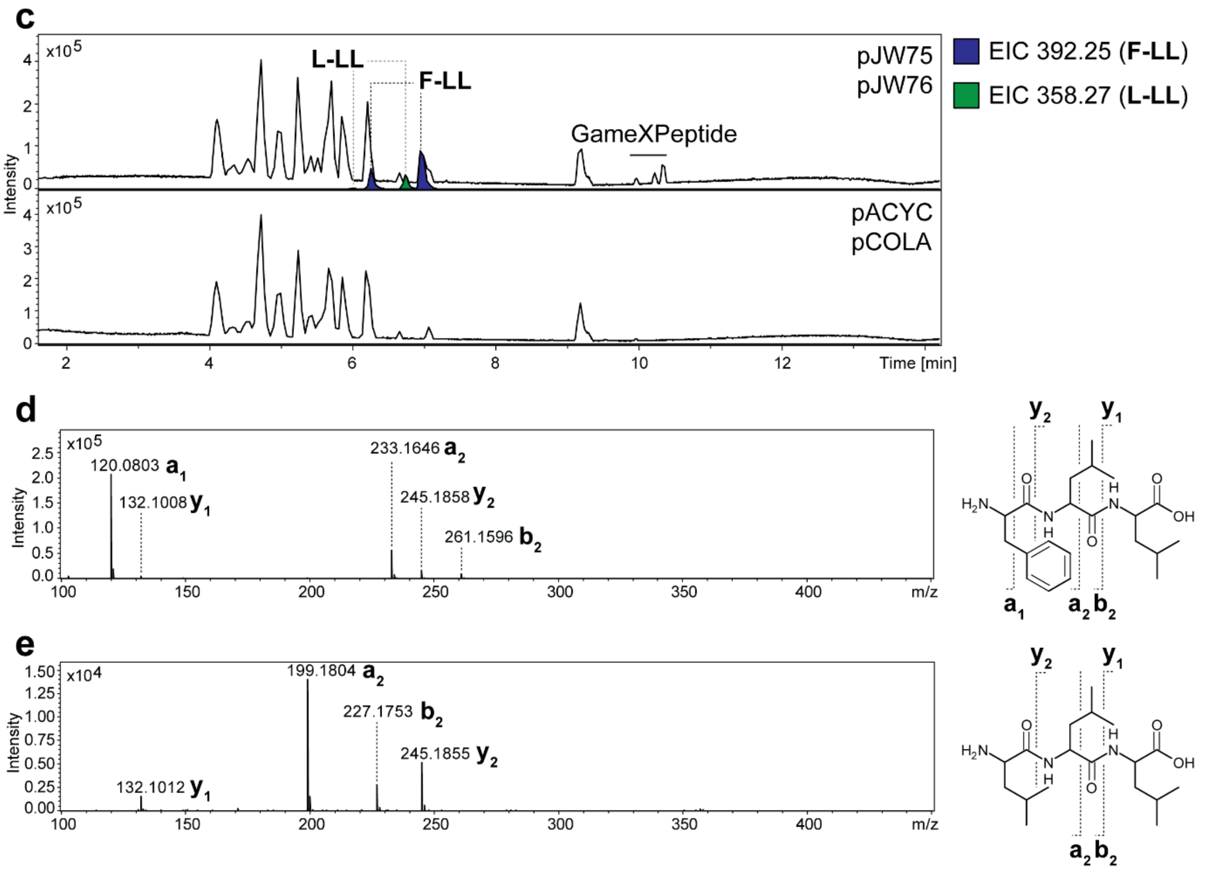

**Figure S6.** HPLC-MS/MS data referred to production of GxpS. (**a**) Schematic representation of GxpS from *P. laumondii* TTO1 described by Bozhyüyk *et al.* 2021.[1,21] The NRPS is split by a pair of synthetic zippers, SZ17 and SZ18, and encoded on two plasmids, pJW75 and pJW76. A domain specitifity is indicated by the one-letter amino acid code. (**b**) Table of HPLC-MS/MS data indicating peptide side product sequences, the detected masses with the corresponding errors and the sum formulas. (**c**) HPLC-MS/MS data of *E. coli* DH10B::*mtaA* expressing *gxpS* (pJW75 and pJW76) or empty vector controls. Base Peak Chromatogram (BPC, top) with Extracted Ion Chromatogram (EIC, below with colors according to the depicted legend) of **F-LL** (*m/z* $[M+H]^+$ = 392.25) and **L-LL** (*m/z* $[M+H]^+$ = 358.27). (**d-e**) The $MS^2$ data for the respective compounds are displayed, along with indications of the assigned fragments.

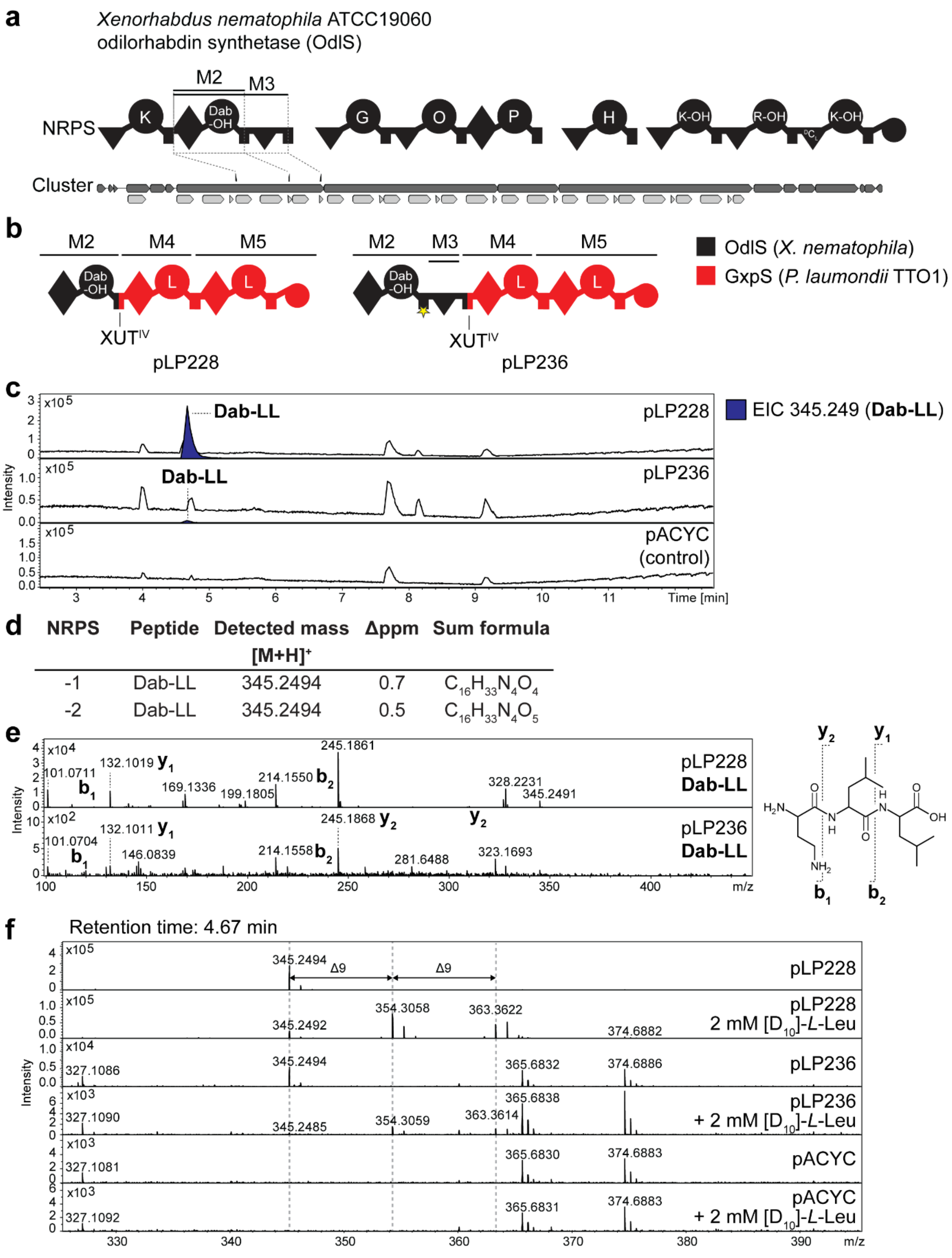

**Figure S7.** HPLC-MS/MS data referred to NRPS hybrids containing OdlS M2 and M3. (**a**) Schematic representation of the OdlS with indication of module 2 and 3 (M2 & M3) targeted by engineering (Dab-OH = β-hydroxy diamino butyric acid, O = ornithine, K-OH = δ-hydroxylysine, R-OH = β-hydroxy-arginine). (**b**) Schematic representation of the NRPS encoded by pLP228 and PLP236. Domain origin is depicted in the legend. Indication of the utilized fusion sites. A domain specitifity is indicated by the one-letter amino acid code (Dab =

diamino butyric acid). Asterisk indicates a point mutation (S1049A) to mutate the serine to an alanine for domain inactivation. (**c**) HPLC-MS/MS data of *E. coli* DH10B::*mtaA* expressing pLP228 and PLP236. An empty vector served as negative control. Base Peak Chromatogram (BPC, top) with Extracted Ion Chromatogram (EIC, below with colors according to the depicted legend) of **Dab-LL** (*m/z* $[M+H]^+$ = 345.24). (**d**) Table of HPLC-MS/MS data indicating peptide side product sequences, the detected masses with the corresponding errors and the sum formulas. (**e**) The $MS^2$ data for the respective compounds are displayed, along with indications of the assigned fragments. (**f**) MS spectra of empty vector control, pLP228 and PLP236 with and without supplementation of 2 mM $[D_{10}]$-*L*-leucine at a retention time of 4.67 min.

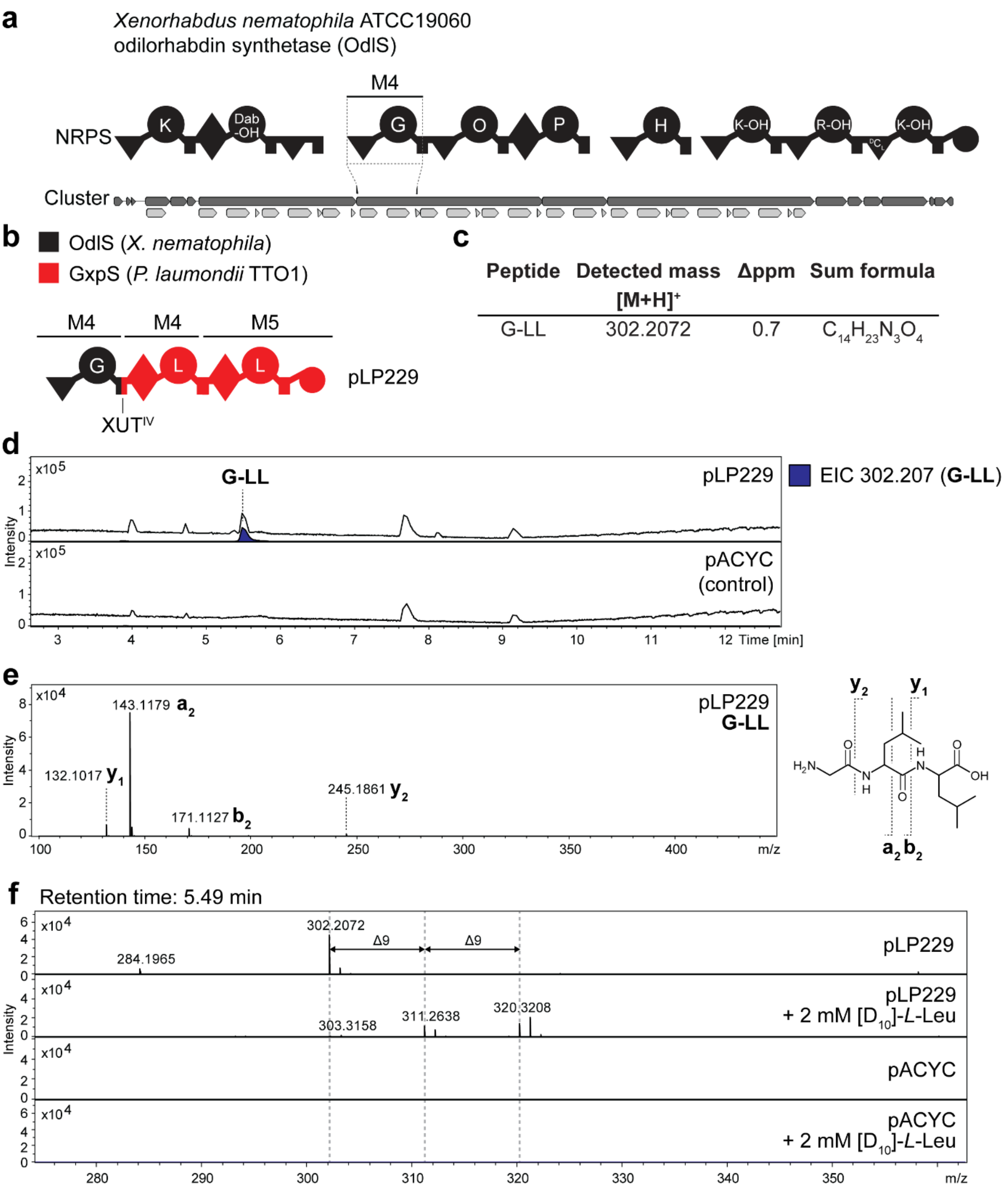

**Figure S8.** HPLC-MS/MS data referred to NRPS hybrids containing OdlS M4. (**a**) Schematic representation of the OdlS with indication of module 4 (M4) targeted by engineering (Dab-OH = β-hydroxy diamino butyric acid, O = ornithine, K-OH = δ-hydroxylysine, R-OH = β-hydroxy-arginine). (**b**) Schematic representation of the NRPS encoded by pLP229. Domain origin is depicted in the legend. Indication of the utilized fusion sites. A domain specitifity is indicated by the one-letter amino acid code. (**c**) Table of HPLC-MS/MS data indicating peptide side product sequence, the detected mass with the corresponding error and the sum formula. (**d**) HPLC-MS/MS data of *E. coli* DH10B::*mtaA* expressing pLP229. An empty vector served as negative control. Base Peak Chromatogram (BPC, top) with Extracted Ion Chromatogram (EIC, below with colors according to the depicted legend) of **G-LL** (*m/z* $[M+H]^+$ = 302.20). (**e**) The $MS^2$ data for the respective compound is displayed, along with indications of the assigned fragments. (**f**) MS spectra of empty vector control and pLP229 with and without supplementation of 2 mM $[D_{10}]$-*L*-leucine at a retention time of 5.49 min.

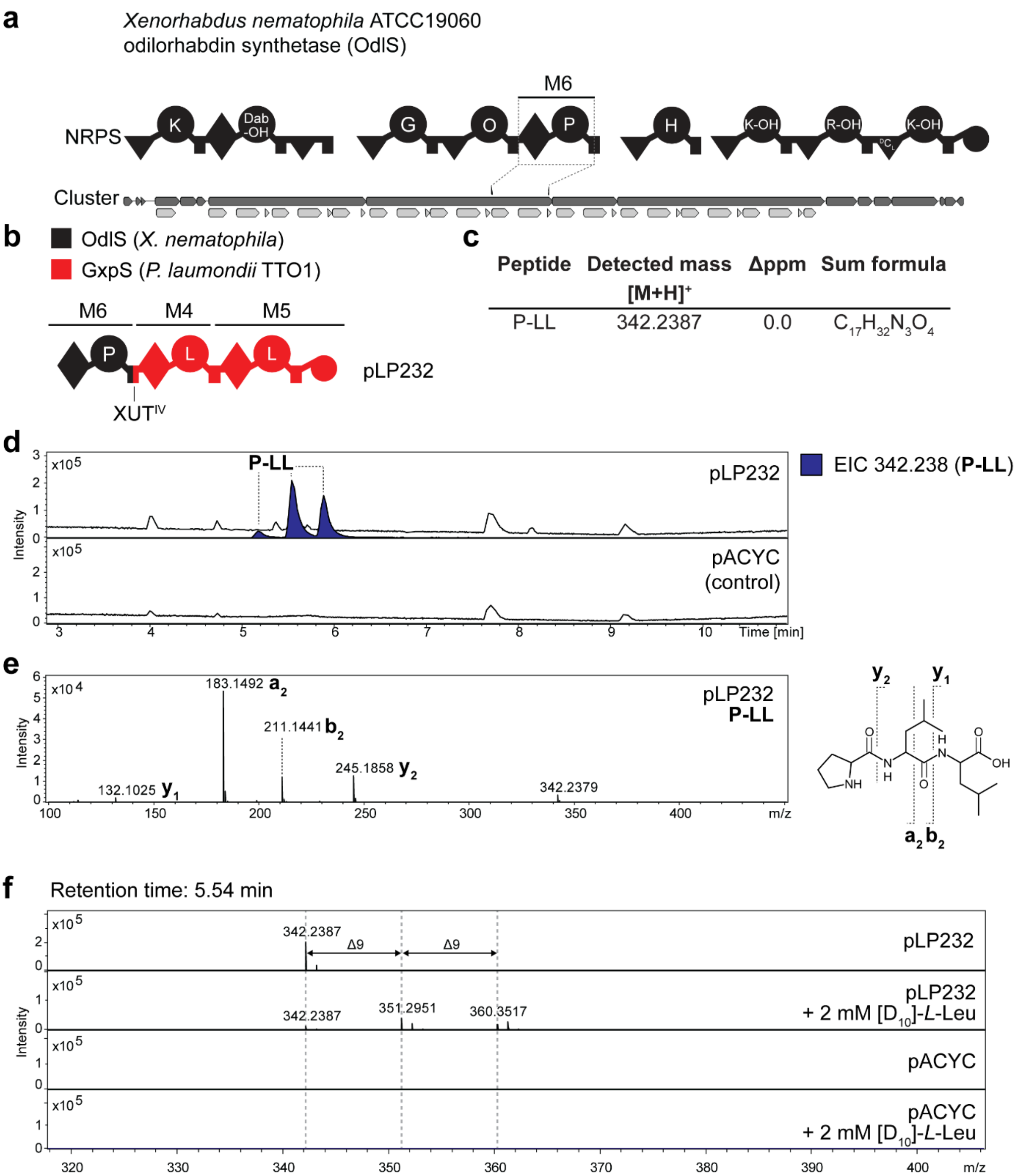

**Figure S9.** HPLC-MS/MS data referred to NRPS hybrids containing OdlS M6. (**a**) Schematic representation of the OdlS with indication of module 6 (M6) targeted by engineering (Dab-OH = β-hydroxy diamino butyric acid, O = ornithine, K-OH = δ-hydroxylysine, R-OH = β-hydroxy-arginine). (**b**) Schematic representation of the NRPS encoded by pLP232. Domain origin is depicted in the legend. Indication of the utilized fusion sites. A domain specitifity is indicated by the one-letter amino acid code. (**c**) Table of HPLC-MS/MS data indicating peptide side product sequence, the detected mass with the corresponding error and the sum formula. (**d**) HPLC-MS/MS data of *E. coli* DH10B::*mtaA* expressing pLP232. An empty vector served as negative control. Base Peak Chromatogram (BPC, top) with Extracted Ion Chromatogram (EIC, below with colors according to the depicted legend) of **P-LL** (*m/z* $[M+H]^+$ = 342.238). (**e**) The $MS^2$ data for the respective compound is displayed, along with indications of the assigned fragments. (**f**) MS spectra of empty vector control and pLP232 with and without supplementation of 2 mM $[D_{10}]$-*L*-leucine at a retention time of 5.54 min.

**a**

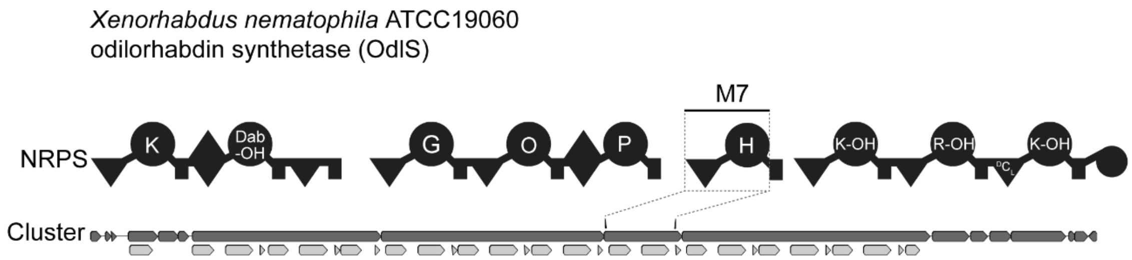

**b**

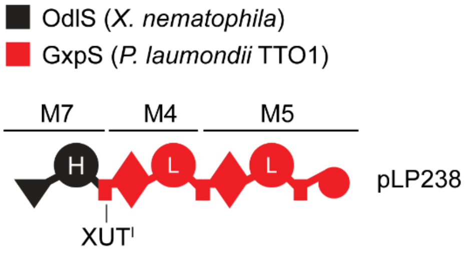

**c**

| Peptide | Detected mass $[M+H]^+$ | Δppm | Sum formula |
|---|---|---|---|
| H-LL | 382.2445 | 1.0 | $C_{18}H_{32}N_5O_4$ |

**d**

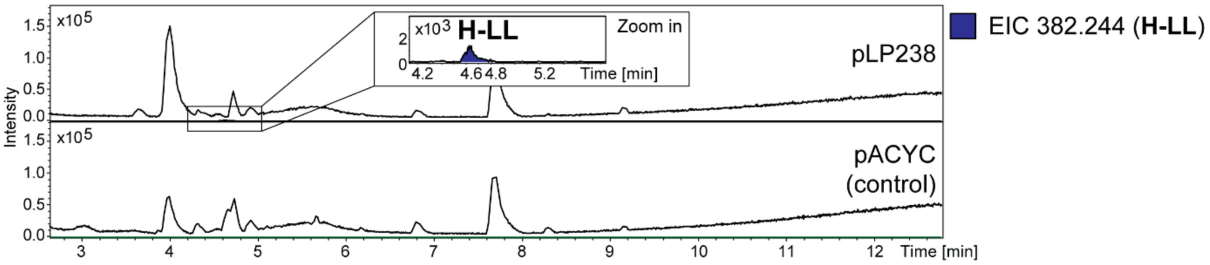

**Figure S10.** HPLC-MS/MS data referred to NRPS hybrids containing OdlS M7. (**a**) Schematic representation of the OdlS with indication of module 7 (M7) targeted by engineering (Dab-OH = β-hydroxy diamino butyric acid, O = ornithine, K-OH = δ-hydroxylysine, R-OH = β-hydroxy-arginine). (**b**) Schematic representation of the NRPS encoded by pLP238. Domain origin is depicted in the legend. Indication of the utilized fusion sites. A domain specitifity is indicated by the one-letter amino acid code. (**c**) Table of HPLC-MS/MS data indicating peptide side product sequence, the detected mass with the corresponding error and the sum formula. (**d**) HPLC-MS/MS data of *E. coli* DH10B::*mtaA* expressing pLP238. An empty vector served as negative control. Base Peak Chromatogram (BPC, top) with Extracted Ion Chromatogram (EIC, below with colors according to the depicted legend) of **H-LL** (*m/z* $[M+H]^+$ = 382.244).

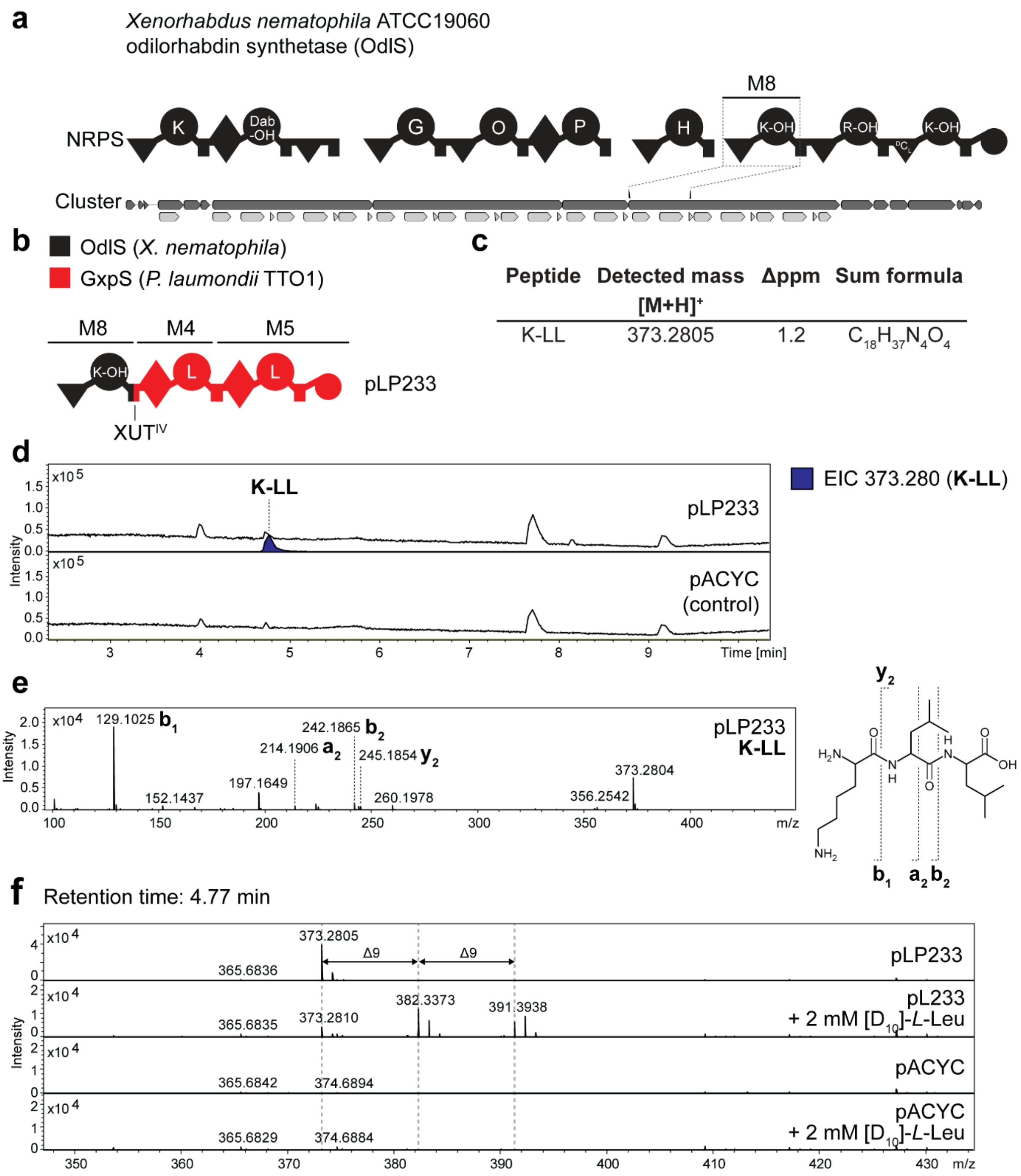

**Figure S11.** HPLC-MS/MS data referred to NRPS hybrids containing OdlS M8. (**a**) Schematic representation of the OdlS with indication of module 8 (M8) targeted by engineering (Dab-OH = β-hydroxy diamino butyric acid, O = ornithine, K-OH = δ-hydroxylysine, R-OH = β-hydroxy-arginine). (**b**) Schematic representation of the NRPS encoded by pLP233. Domain origin is depicted in the legend. Indication of the utilized fusion sites. A domain specitifity is indicated by the one-letter amino acid code (K-OH = δ-hydroxy-lysine). (**c**) Table of HPLC-MS/MS data indicating peptide side product sequence, the detected mass with the corresponding error and the sum formula. (**d**) HPLC-MS/MS data of *E. coli* DH10B::*mtaA* expressing pLP233. An empty vector served as negative control. Base Peak Chromatogram (BPC, top) with Extracted Ion Chromatogram (EIC, below with colors according to the depicted legend) of **K-LL** (*m/z* $[M+H]^+$ = 373.280). (**e**) The $MS^2$ data for the respective compound is displayed, along with indications of the assigned fragments. (**f**) MS spectra of empty vector control and pLP233 with and without supplementation of 2 mM $[D_{10}]$-*L*-leucine at a retention time of 4.77 min.

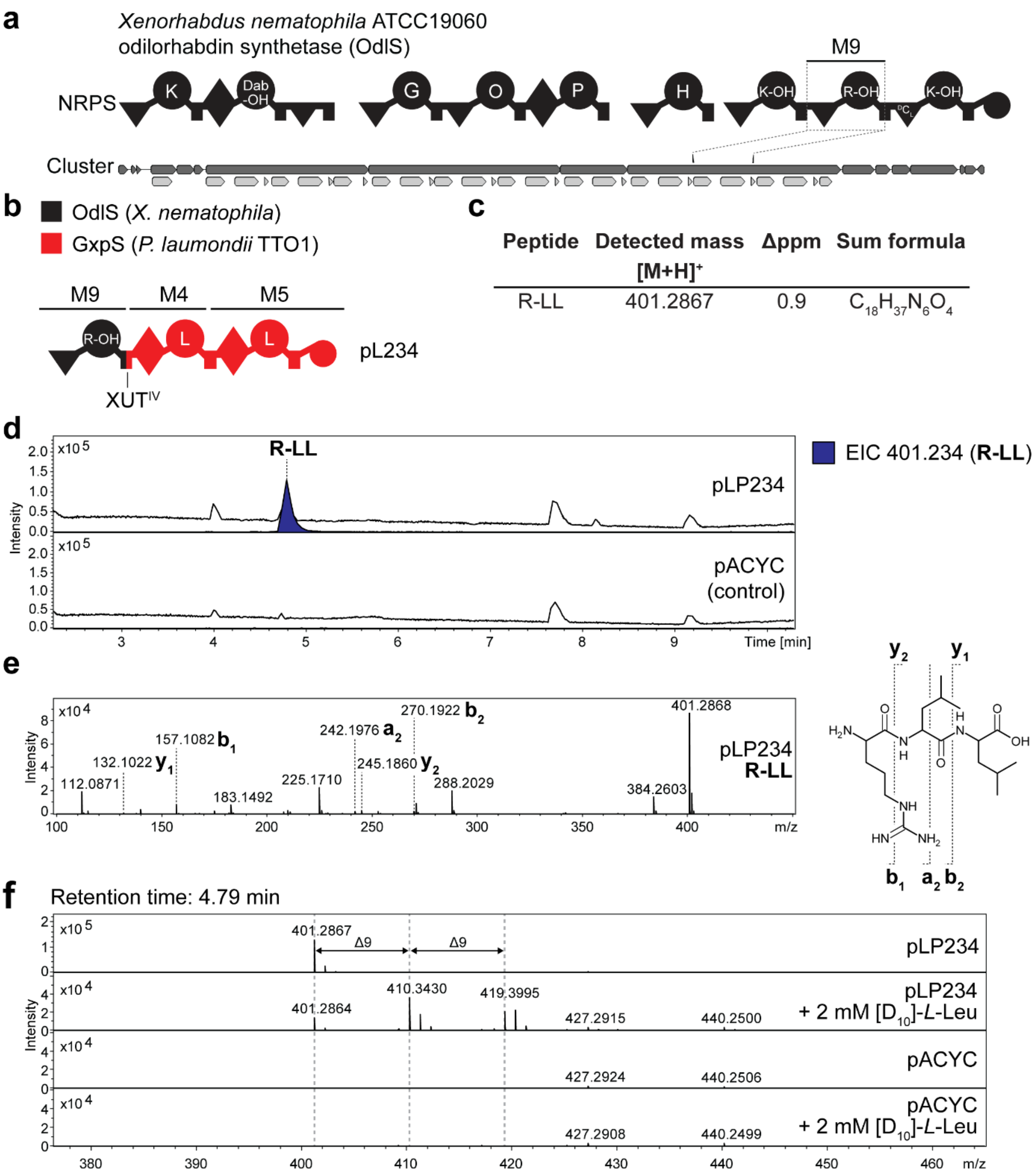

**Figure S12.** HPLC-MS/MS data referred to NRPS hybrids containing OdlS M9. (**a**) Schematic representation of the OdlS with indication of module 9 (M9) targeted by engineering (Dab-OH = β-hydroxy diamino butyric acid, O = ornithine, K-OH = δ-hydroxylysine, R-OH = β-hydroxy-arginine). (**b**) Schematic representation of the NRPS encoded by pLP234. Domain origin is depicted in the legend. Indication of the utilized fusion sites. A domain specitifity is indicated by the one-letter amino acid code (R-OH = β-hydroxy-arginine). (**c**) Table of HPLC-MS/MS data indicating peptide side product sequence, the detected mass with the corresponding error and the sum formula. (**d**) HPLC-MS/MS data of *E. coli* DH10B::*mtaA* expressing pLP234. An empty vector served as negative control. Base Peak Chromatogram (BPC, top) with Extracted Ion Chromatogram (EIC, below with colors according to the depicted legend) of **R-LL** (*m/z* $[M+H]^+$ = 401.234). (**e**) The $MS^2$ data for the respective compound is displayed, along with indications of the assigned fragments. (**f**) MS spectra of empty vector control and pLP234 with and without supplementation of 2 mM $[D_{10}]$-*L*-leucine at a retention time of 4.79 min.

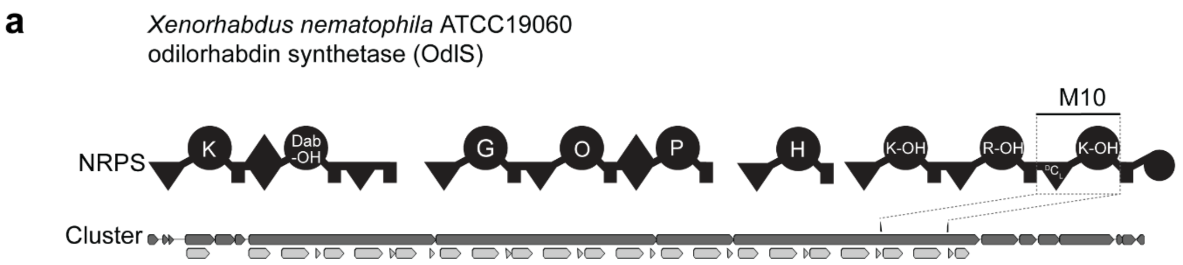

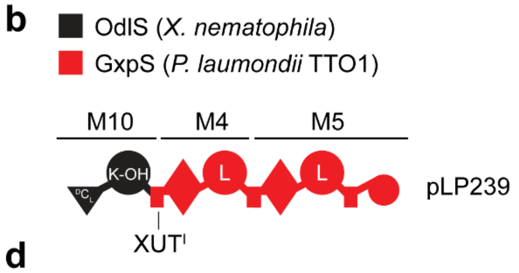

c

| Peptide | Detected mass $[M+H]^+$ | Δppm | Sum formula |
|---|---|---|---|
| K-LL | 373.2809 | 0.0 | $C_{18}H_{37}N_4O_4$ |

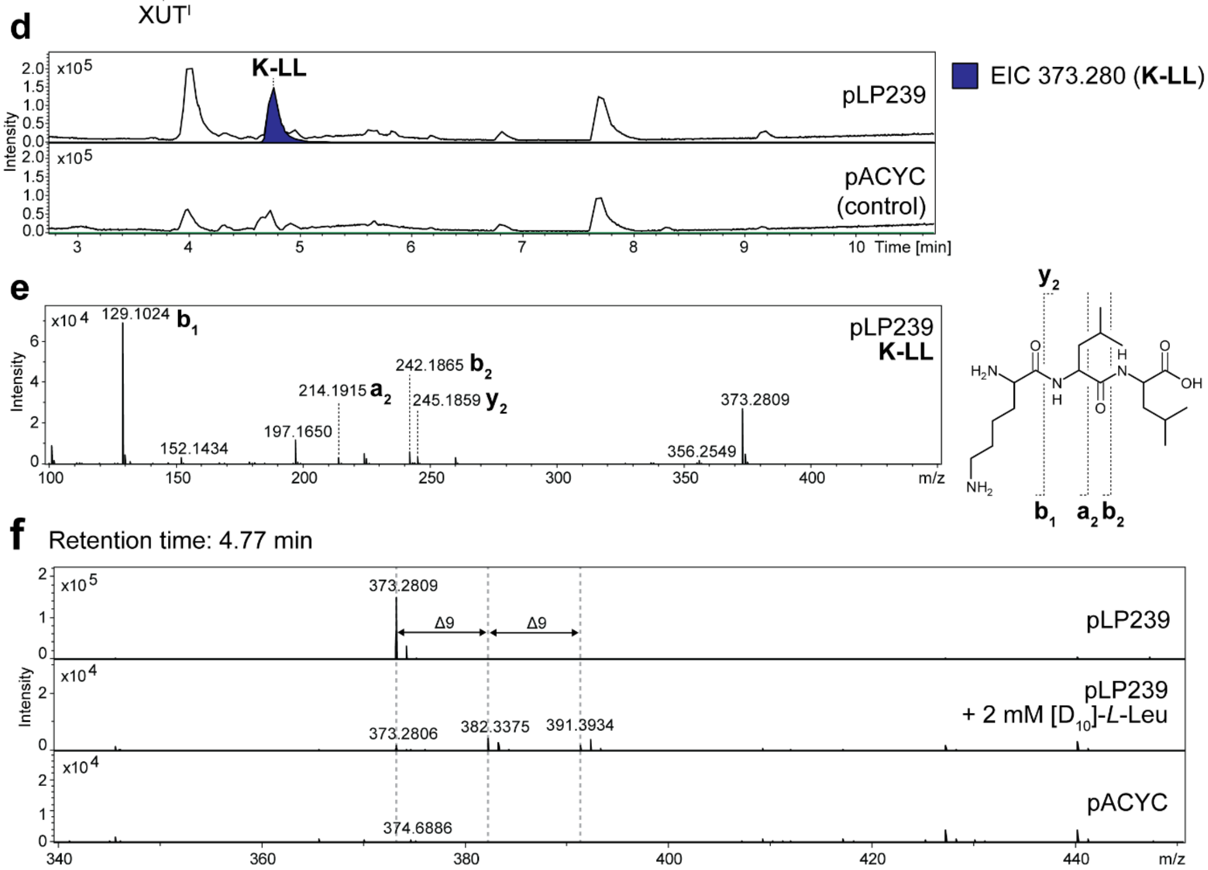

**Figure S13.** HPLC-MS/MS data referred to NRPS hybrids containing OdlS M10. (**a**) Schematic representation of the OdlS with indication of module 10 (M10) targeted by engineering (Dab-OH = β-hydroxy diamino butyric acid, O = ornithine, K-OH = δ-hydroxylysine, R-OH = β-hydroxy-arginine). (**b**) Schematic representation of the NRPS encoded by pLP239. Domain origin is depicted in the legend. Indication of the utilized fusion sites. A domain specitifity is indicated by the one-letter amino acid code (K-OH = δ-hydroxy-lysine). (**c**) Table of HPLC-MS/MS data indicating peptide side product sequence, the detected mass with the corresponding error and the sum formula. (**d**) HPLC-MS/MS data of *E. coli* DH10B::*mtaA* expressing pLP239. An empty vector served as negative control. Base Peak Chromatogram (BPC, top) with Extracted Ion Chromatogram (EIC, below with colors according to the depicted legend) of **K-LL** (*m/z* $[M+H]^+$ = 373.280). (**e**) The $MS^2$ data for the respective compound is displayed, along with indications of the assigned fragments. (**f**) MS spectra of empty vector control and pLP239 with and without supplementation of 2 mM $[D_{10}]$-*L*-leucine at a retention time of 4.77 min.

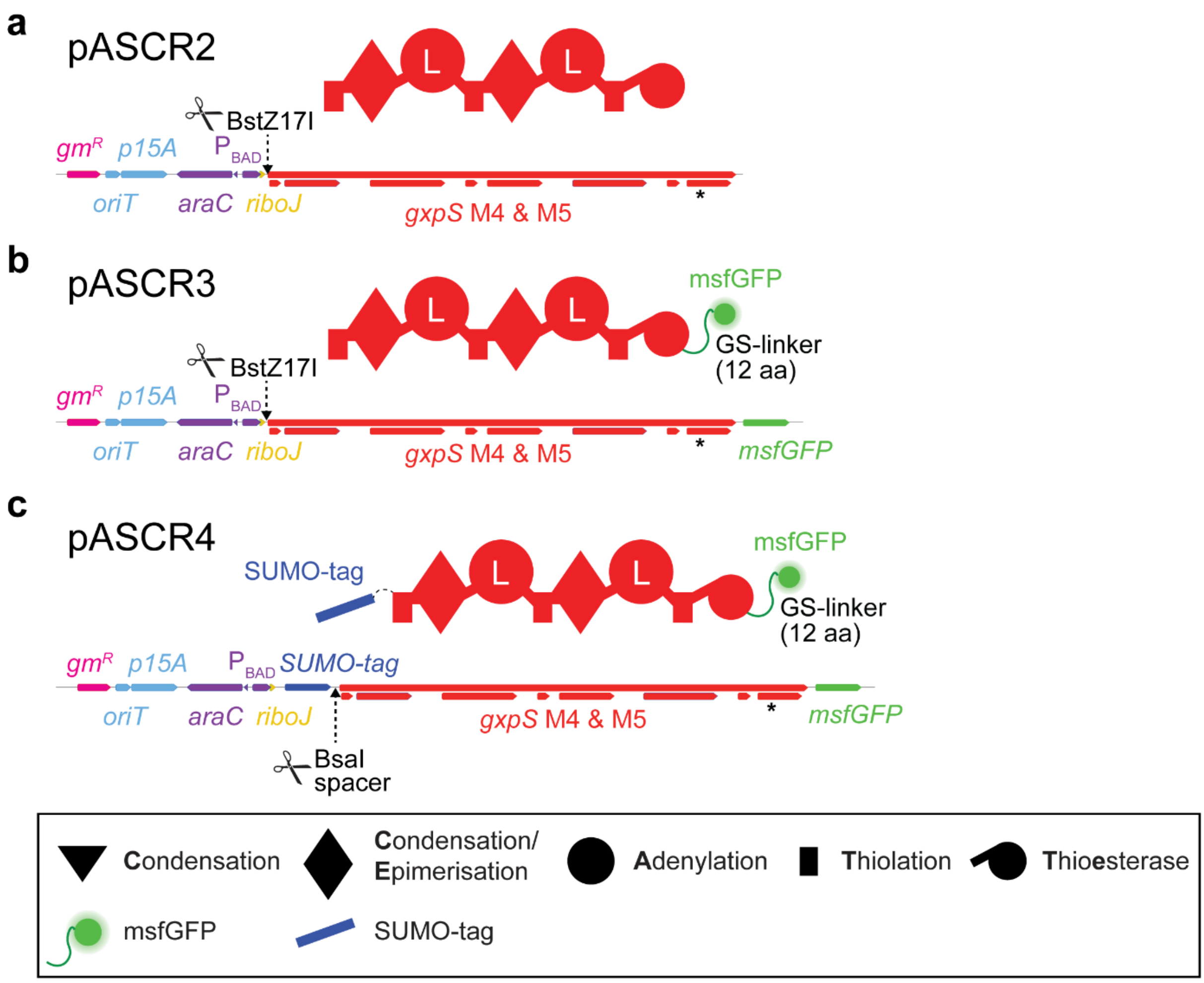

**Figure S14.** Schematic illustration of the cloning vectors. Linearized illustration of the plasmid maps of pASCR2 (**a**), pASCR3 (**b**) and pASCR4 (**c**). Indication of the resistance genes, origin of replications, promoter regions, introduced restriction sites, NRPS-encoding regions and added tags. An asterisk (*) indicates the introduced silent mutation at G2315 (GGT→GGG) to ensure single cleavage by the restriction enzyme. Schematic illustration of the encoded protein above. The corresponding NRPS domain and protein-tag legend is depicted below.

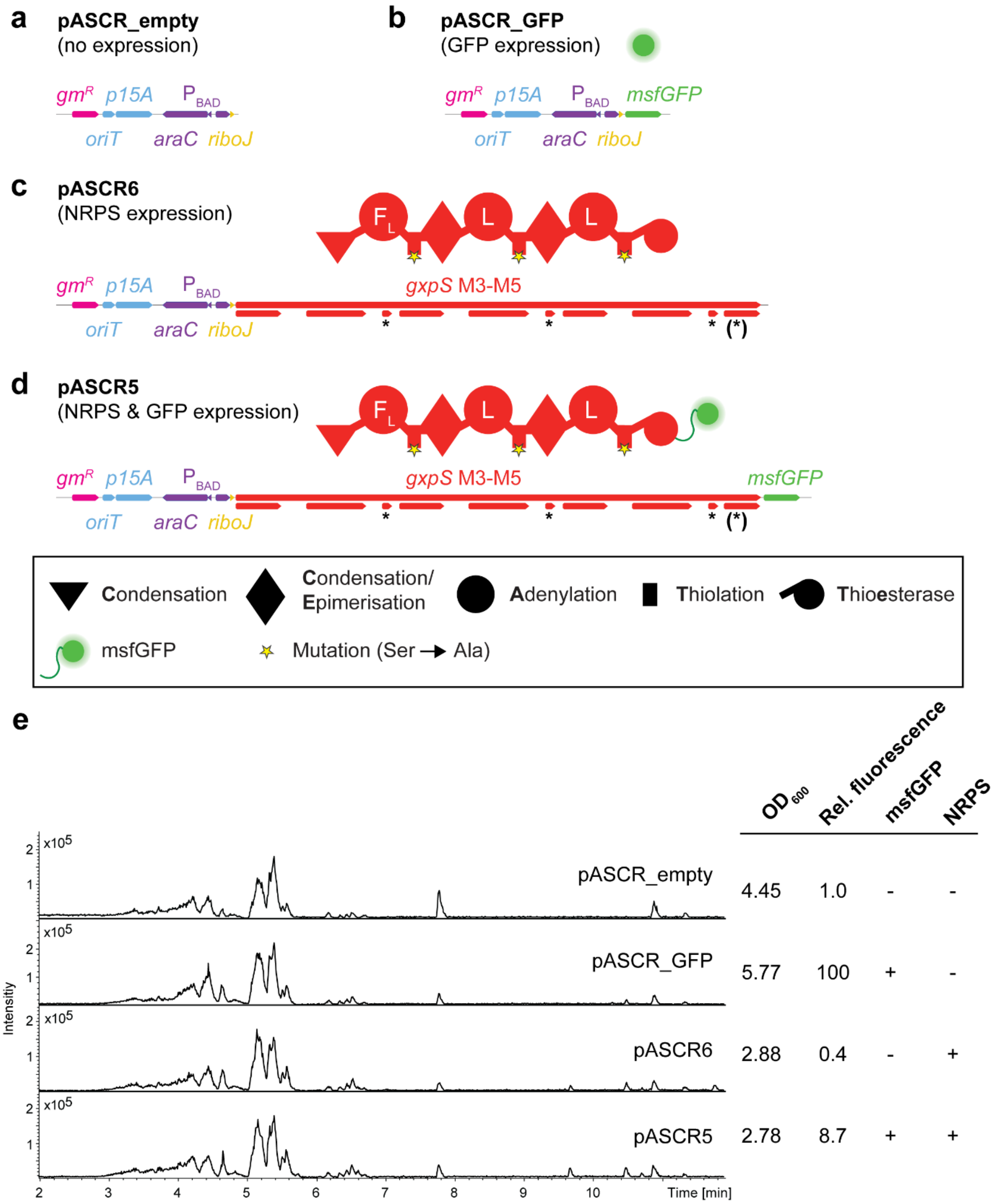

**Figure S15.** Control vector construction. Linearized illustration of the plasmid maps of pASCR_empty (**a**), pASCR_GFP (**b**), pASCR6 (**c**) and pASCR5 (**d**). Indication of the resistance genes, origin of replications, promoter regions, NRPS-encoding regions and added tags. An asterisk (*) indicates the introduced mutations within the T domains to inactivate the NRPS. An asterisk within brackets indicates a silent mutation at G3274 (GGT→GGG). Schematic illustration of the encoded protein above. The corresponding NRPS domain and protein-tag legend is depicted below. (**e**) HPLC-MS/MS data of *E. coli* DH10B::*mtaA* expressing the individual control vectors. The Base Peak Chromatogram (BPC) is shown. On the right the msfGFP fluorescence (Ex.: 480 nm; Em.: 525 nm) and the $OD_{600}$ is depicted for the same production. (-) and (+) indicates presence or absence of the protein.

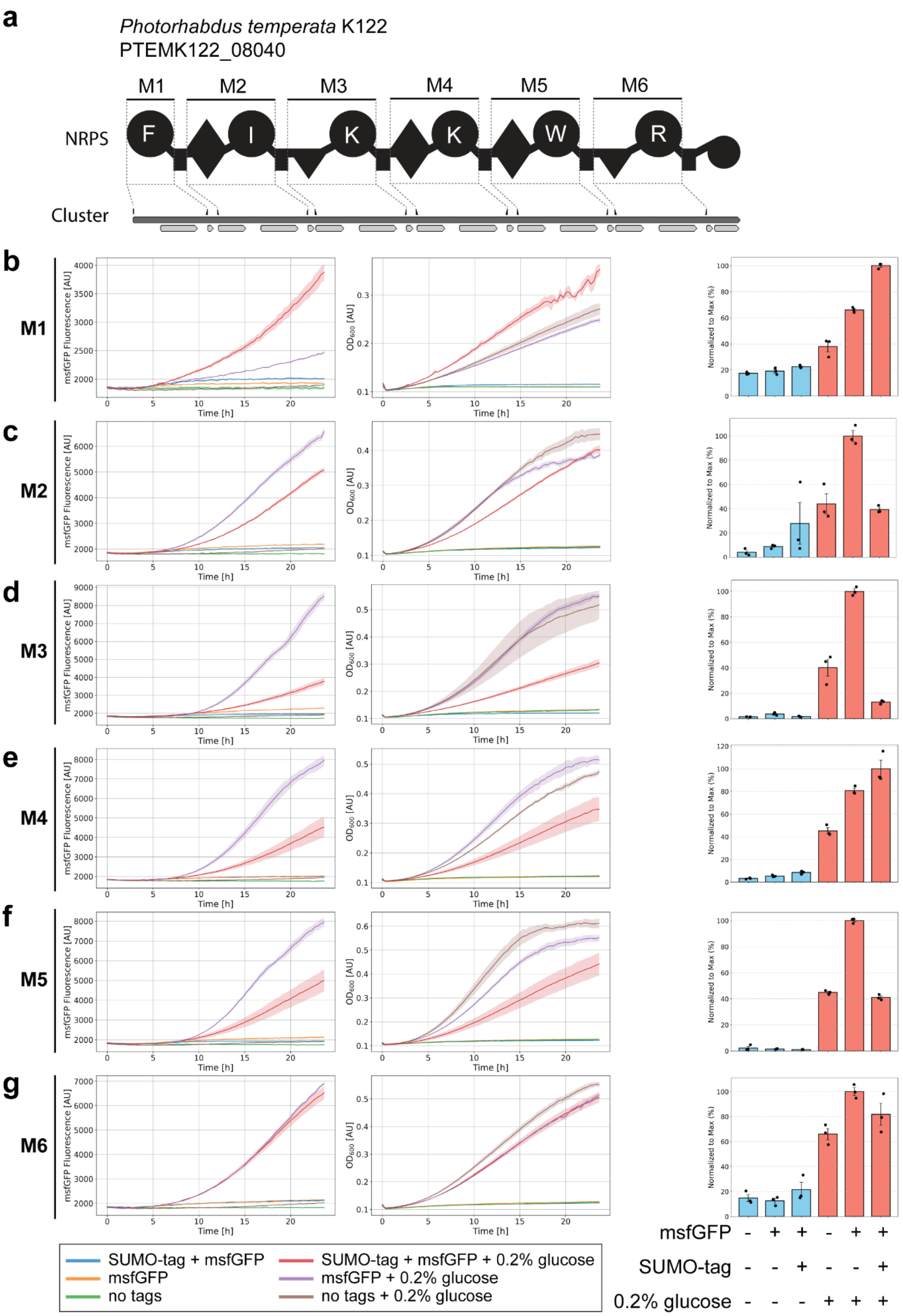

**Figure S16.** Comparison of expression, growth and peptide production for cloning vectors pASCR2, pASCR3 and pASCR4. (**a**) Illustration of the gene cluster PTEMK122_08040 from

*P. temperata* K122 with schematic illustration of the encoded NRPS. Indication of the region of cloned modules on gene and protein level. Utilized primers are indicated above the cluster A domain specitifity is indicated by the one-letter amino acid code. (**b-g**) The genes of the individual six modules were cloned onto pASCR2 (no tags), pASCR3 (msfGFP) and pASCR4 (SUMO-tag and msfGFP) and were expressed in *E. coli* DH10B::*mtaA*. Subsequently, msfGFP fluorescence, the $OD_{600}$ and peptide production were determined in XPP medium with and without supplementation of 0.2 % glucose. The figure displays plots of the individual targeted modules of PTEMK122_08040 (indicated on the left site) for msfGFP fluorescence (Ex.: 480 nm; Em.: 525 nm, left) and the $OD_{600}$ (middle) determined for 24 hours. Experimental conditions are indicated by the color code in the legend below. The bar plot (right) shows the relative production of the expected tripeptide for the respective module for production after 72 hours. The experimental conditions are indicated by the legend below, where (+) indicates presence and (-) absence of this condition. In all plots, the standard errors of the means (SEM) are displayed.

**a**

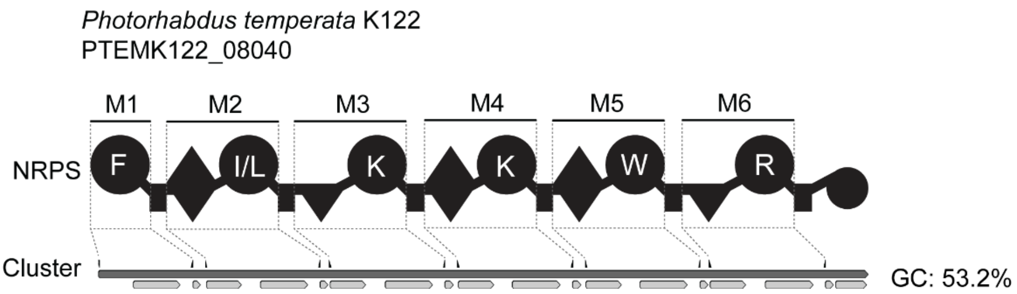

**b**

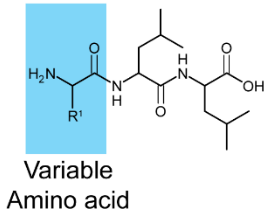

| Module | Detected mass [M+H]+ | Δppm | Sum formula | Estimated amino acid |
|---|---|---|---|---|
| M1 | 392.2546 | -0.6 | $C_{21}H_{34}N_3O_4$ | Phe |
| M2 | 358.2701 | -0.1 | $C_{18}H_{36}N_3O_4$ | Ile/Leu |
| M2 | 373.2811 | -0.3 | $C_{18}H_{37}N_4O_4$ | Lys |
| M3 | 373.2808 | 0.4 | $C_{18}H_{37}N_4O_4$ | Lys |
| M4 | 401.2855 | 3.9 | $C_{18}H_{37}N_6O_4$ | Arg |
| M5 | 431.2651 | 0.5 | $C_{23}H_{35}N_4O_4$ | Trp |
| M6 | 401.2879 | 0.3 | $C_{18}H_{37}N_6O_4$ | Arg |

**c**

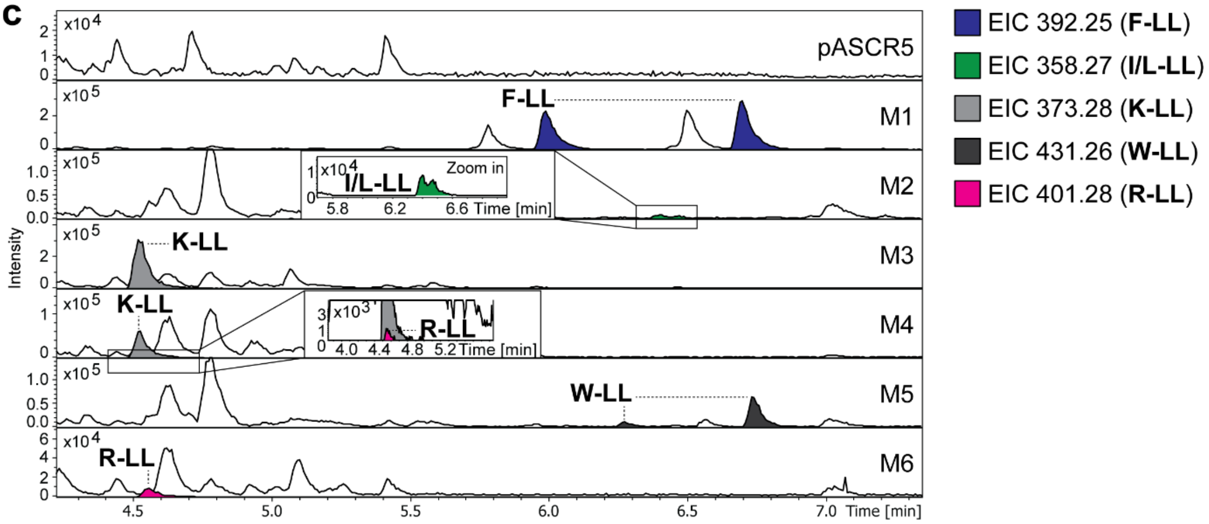

**Figure S17.** HPLC-MS/MS data referred to module screening of PTEMK122_08040 from *P. temperata* K122. (**a**) Illustration of the gene cluster PTEMK122_08040 from *P. temperata* K122 with schematic illustration of the encoded NRPS. Indication of the region of cloned modules M1 to M6 on gene and protein level. Utilized primers are shown above the cluster. A domain specitifity is indicated by the one-letter amino acid code. GC content is depicted next to the cluster. (**b**) Table of HPLC-MS/MS data indicating the targeted module cloned into pASCR3, the detected mass of the compound, the corresponding error, the sum formula and the incoporated amino acid introduced to the peptide at the highlighted position in the chemical structure (left). The amino acid is shown by the three-letter amino acid code. (**c**) HPLC-MS/MS data of *E. coli* DH10B::*mtaA* expressing M1 to M6 cloned into pASCR3. Expression of pASCR5 served as negative control. Base Peak Chromatogram (BPC, top) with Extracted Ion Chromatogram (EIC, below with colors according to the depicted legend) of **F-LL** (*m/z* $[M+H]^+$ = 392.254), **I/L-LL** (*m/z* $[M+H]^+$ = 358.270), **K-LL** (*m/z* $[M+H]^+$ = 373.280), **W-LL** (*m/z* $[M+H]^+$ = 431.265) and **R-LL** (*m/z* $[M+H]^+$ = 401.287). Indication of the targeted modules on the right of the chromatograms.

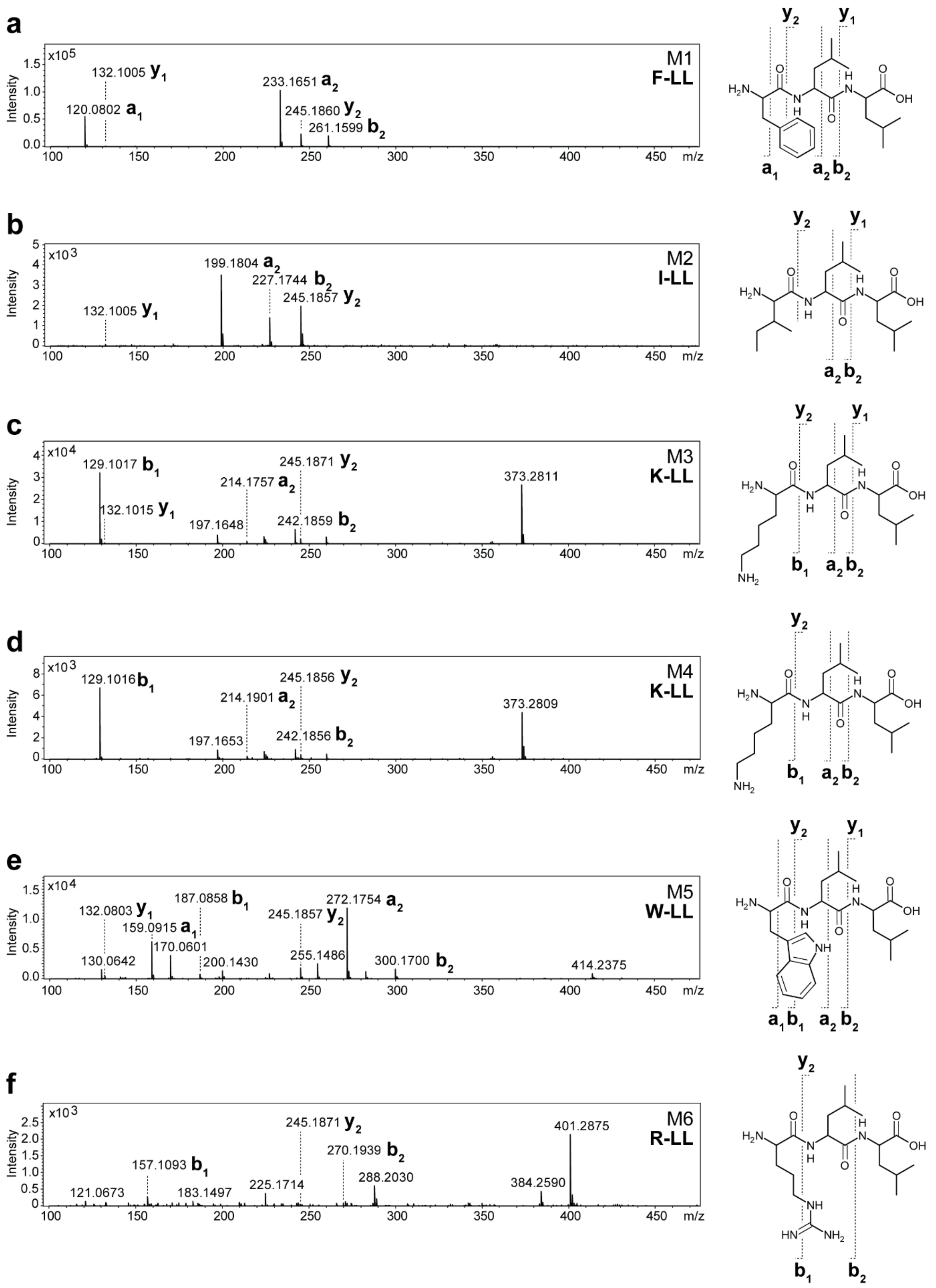

**Figure S18.** MS$^2$ spectra of HPLC-MS/MS data referred to figure S17. (**a-f**) The respective compounds are displayed, along with indications of the assigned fragments. The targeted module and peptide name is depicted within the spectra.

**a**

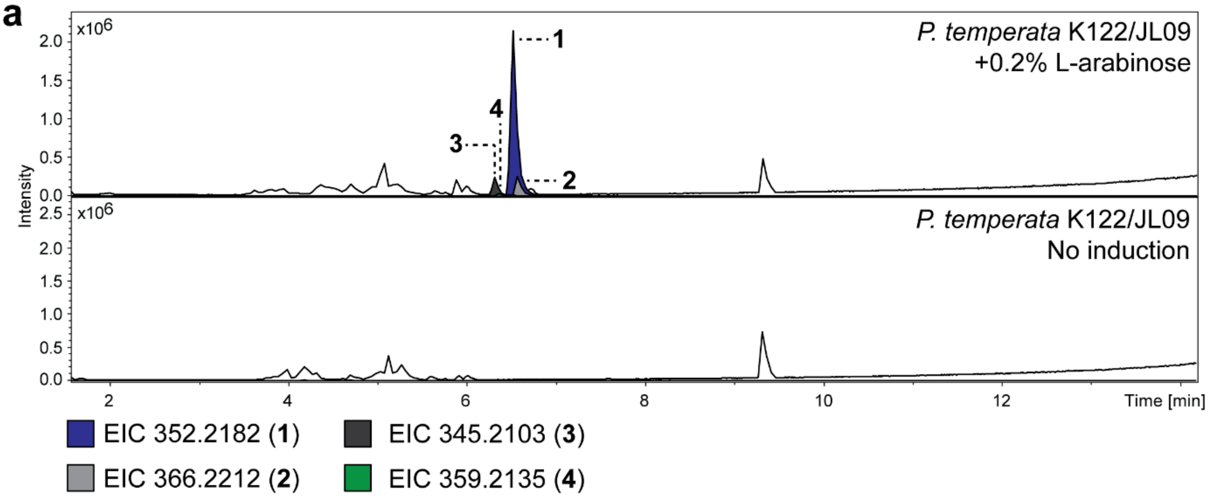

**b**

| Compound | Detected mass $[M+2H]^{2+}$ | Δppm | Sum formula | Assigned peptide |
|---|---|---|---|---|
| **1** | 352.2182 | -0.1 | $C_{38}H_{56}N_8O_5$ | cyclic(FIkkW) |
| **2** | 366.2211 | 0.4 | $C_{38}H_{56}N_{10}O_5$ | cylic(FIkrW) |
| **3** | 345.2100 | 0.8 | $C_{37}H_{54}N_8O_5$ | cylic(FVkkW) |
| **4** | 359.2133 | 0.2 | $C_{37}H_{54}N_{10}O_5$ | cylic(FVkrW) |

**Figure S19.** HPLC-MS/MS data of *P. temperata* K122 $P_{BAD}$ promoter exchange based activation of PTEMK122_08040. (**a**) HPLC-MS analysis of *P. temperata* K122/JL09 cultivated in low salt LB medium with and without induction with 0.2% L-arabinose. Comparison of Base Peak Chromatogram (BPC) is displayed with corresponding Extracted Ion Chromatogram (EIC) of compound **1** (*m/z* $[M+2H]^{2+}$ = 352.218), **2** (*m/z* $[M+2H]^{2+}$ = 366.221), **3** (*m/z* $[M+2H]^{2+}$ = 345.210) and **4** (*m/z* $[M+2H]^{2+}$ = 359.2135) depicted below. (**b**) High-resolution MS data of compound **1** to **4** with indications of the detected mass, the error, the sum formula and an assigned peptide structure represented by the one-letter amino acid code. Capital letter indicate L-configuration and lowercase letter D-configuration.

**a**

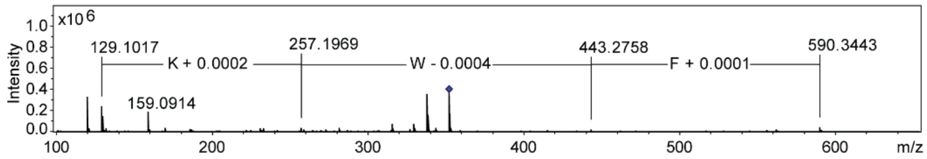

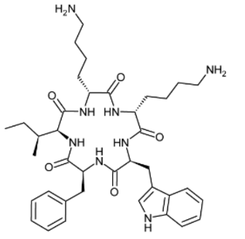

**1**

**b**

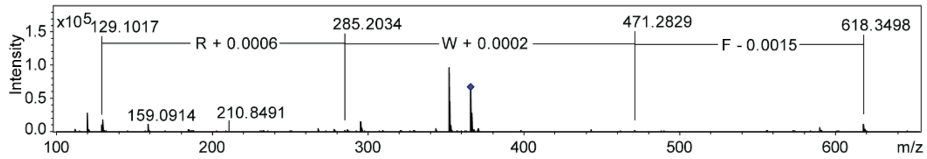

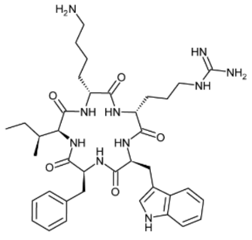

**2**

**c**

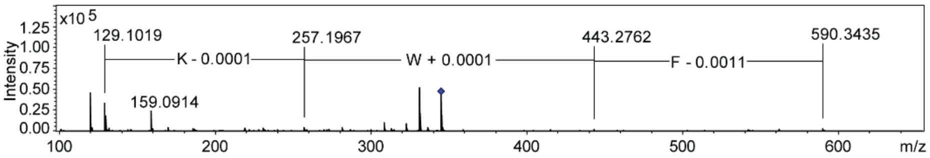

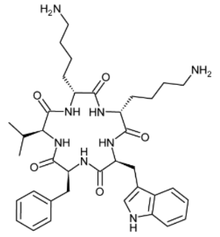

**3**

**d**

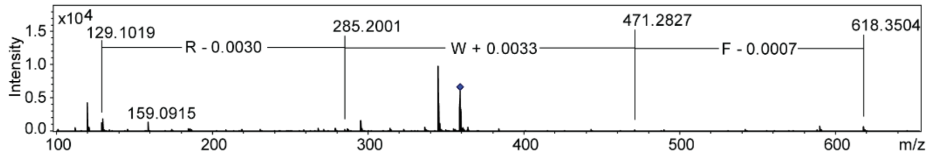

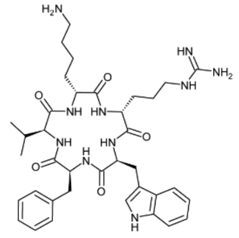

**4**

**Figure S20.** $MS^2$ spectra of HPLC-MS/MS data referred to figure S19. (**a-d**) Compounds **1**-**4** are displayed, along with indications of the $MS^2$ fragmentation spectra. Mass differences are indicated within the spectra.

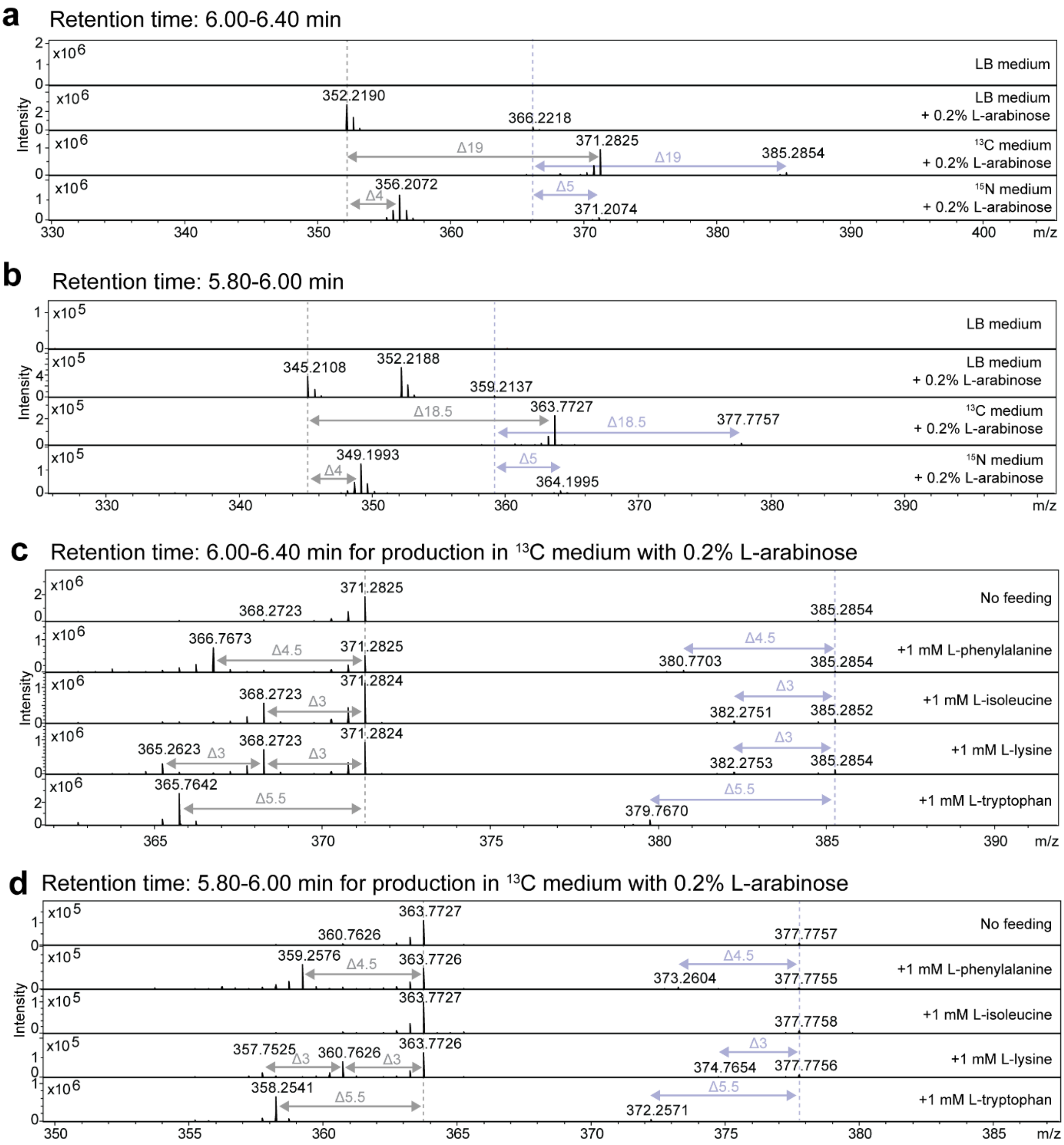

**Figure S21.** Isotope labeling experiments for structure determination of compounds **1**-**4**. HPLC-MS analysis of *P. temperata* K122/JL09. Determination of carbon and nitrogen atoms in compounds **1** (*m/z* $[M+2H]^{2+}$ = 352.218) (**a**), **2** (*m/z* $[M+2H]^{2+}$ = 366.221) (**a**), **3** (*m/z* $[M+2H]^{2+}$ = 345.210) (**b**) and **4** (*m/z* $[M+2H]^{2+}$ = 359.213) (**b**) by isotope labelling experiments. MS spectra are displayed for the strain cultivated in low salt LB medium ($^{12}C$, $^{14}N$), fully labelled $^{13}C$ medium ($^{13}C$, $^{14}N$) and $^{15}N$ medium ($^{12}C$, $^{15}N$) with indication of the retention time and mass shifts. In addition, MS spectra of inverse feeding experiments are shown, performed in fully labelled $^{13}C$ medium ($^{13}C$, $^{14}N$) supplemented with unlabelled L-phenylalanine, L-isoleucine, L-lysine and L-tryptophan. MS spectra retention times are shown with indications of mass shifts for the respective compounds **1**/**2** (**c**) and **3**/**4** (**d**).

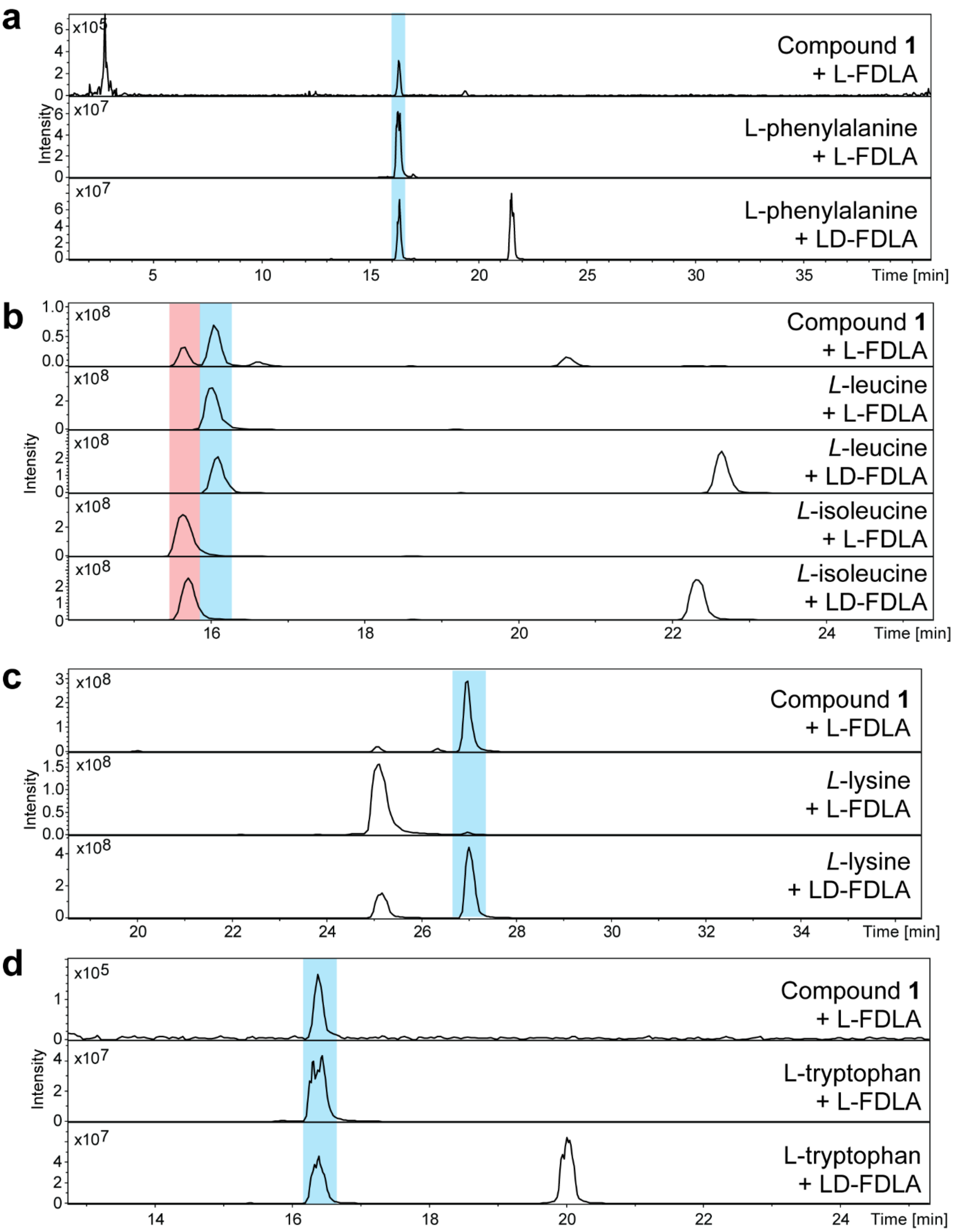

**Figure S22.** Configuration determination of amino acids in **1** using the advanced Marfey's method. HPLC-MS analysis of hydrolyzed and L-FDLA derivatized **1** along with derivatized amino acid standards with L-FDLA and LD-FDLA. Depicted are EIC traces for Marfey's derivatives of L-phenylalanine (Phe, *m/z* 460.21 $[M+H]^+$) (**a**), L-leucine and L-isoleucine (Ile, *m/z* 426.20 $[M+H]^+$)(**b**), L-di-lysine Marfey derivative (Lys, *m/z* 735.31 $[M+H]^+$)(**c**) and L-tryptophan (Trp, *m/z* 499.17 $[M+H]^+$)(**d**).

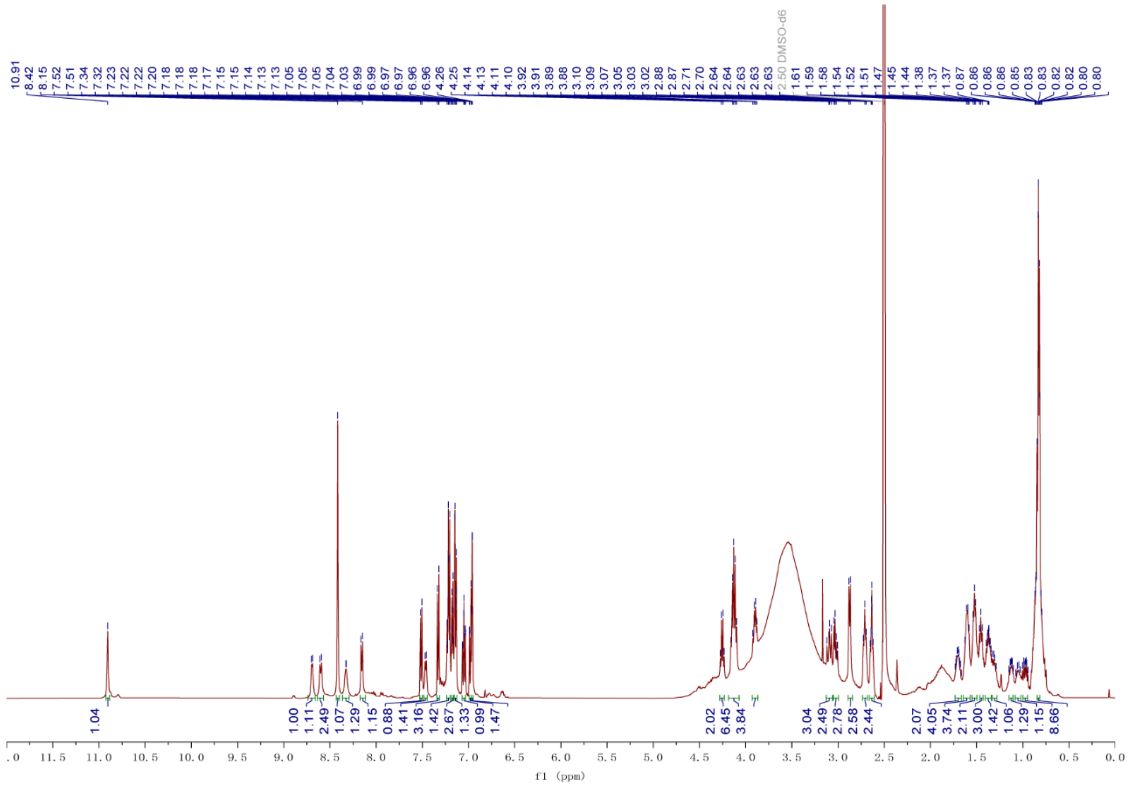

**Figure S23.** $^1H$ NMR spectrum (DMSO-$d_6$, 500 MHz) of **1**.

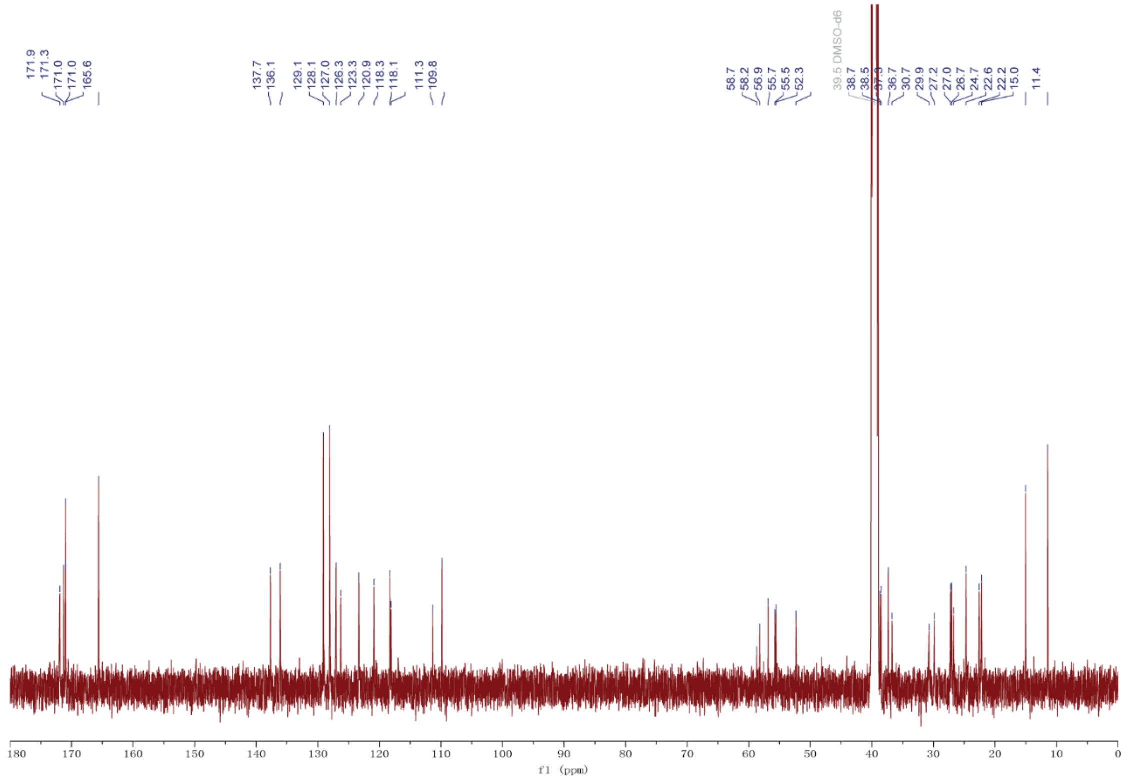

**Figure S24.** $^{13}C$ NMR spectrum (DMSO-$d_6$, 125 MHz) of **1**.

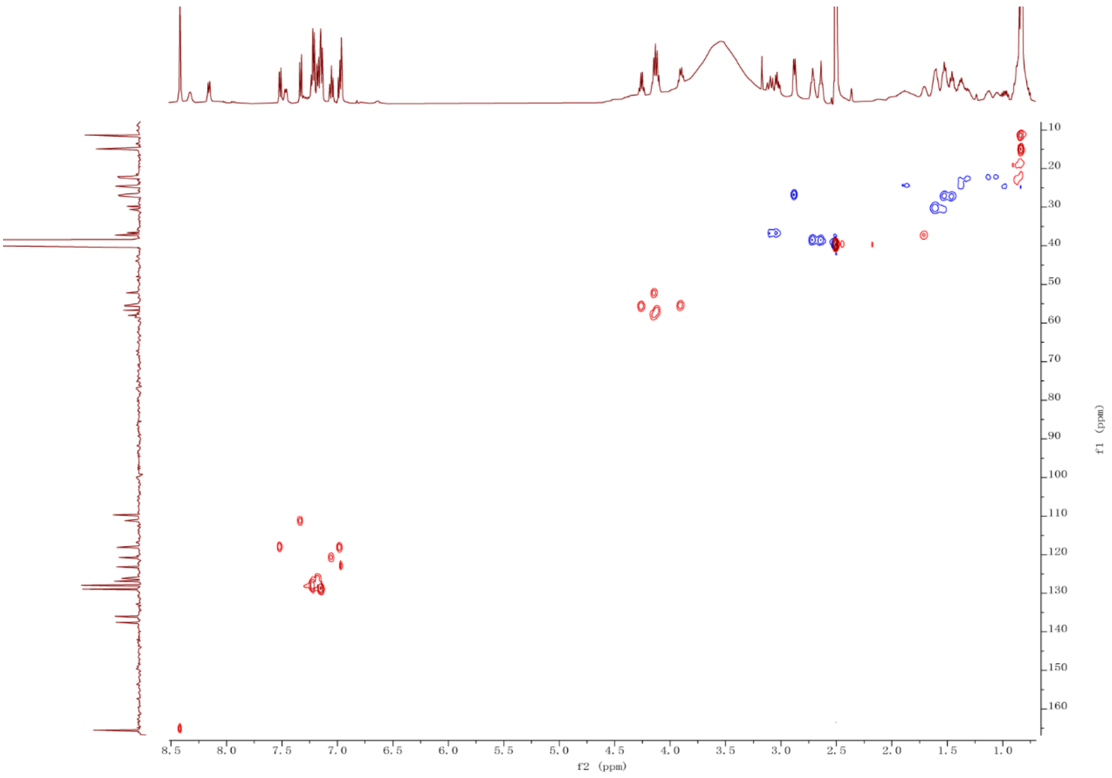

**Figure S25.** HSQC NMR spectrum (DMSO-$d_6$, 500 MHz) of **1**.

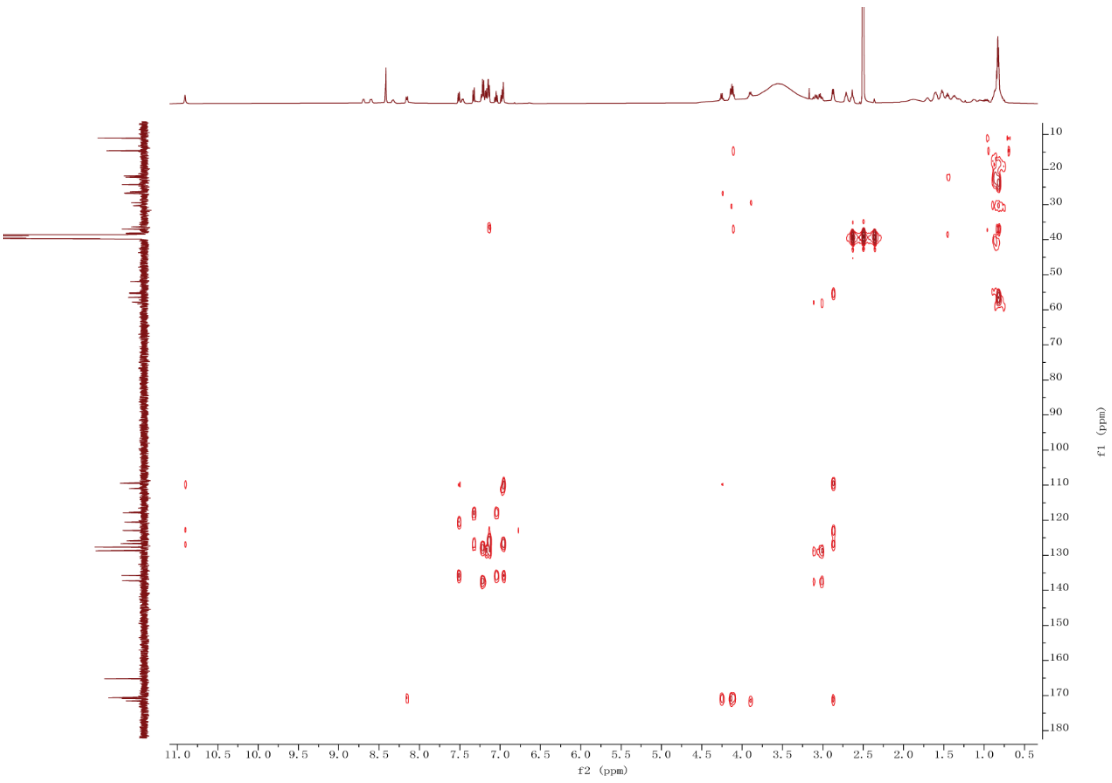

**Figure S26.** HMBC spectrum (DMSO-$d_6$, 500 MHz) of **1**.

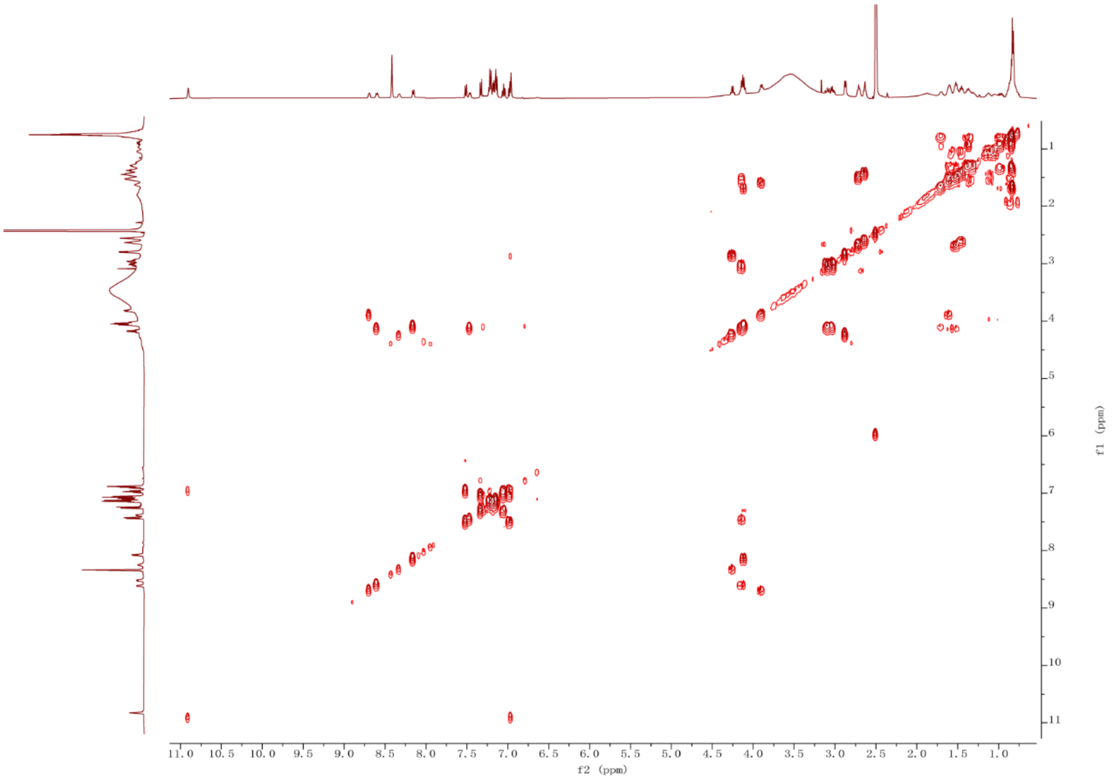

**Figure S27.** $^{1}H$-$^{1}H$ COSY spectrum (DMSO-$d_6$, 500 MHz) of **1**.

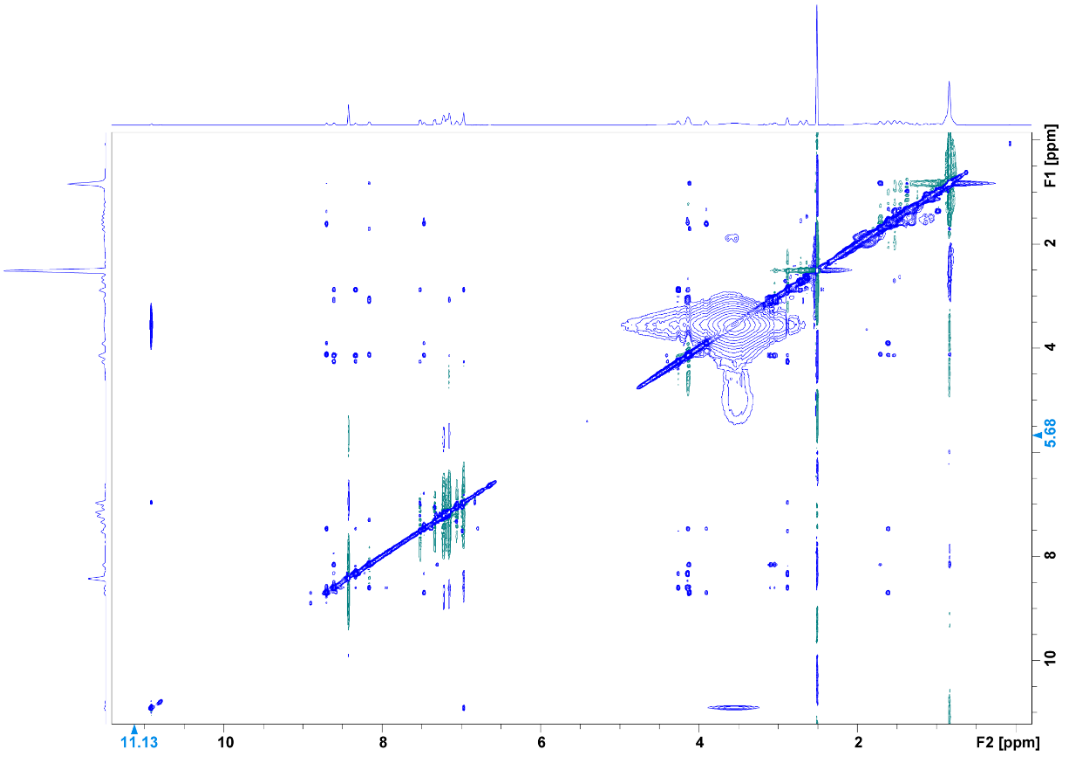

**Figure S28.** NOESY spectrum (DMSO-$d_6$, 500 MHz) of **1**.

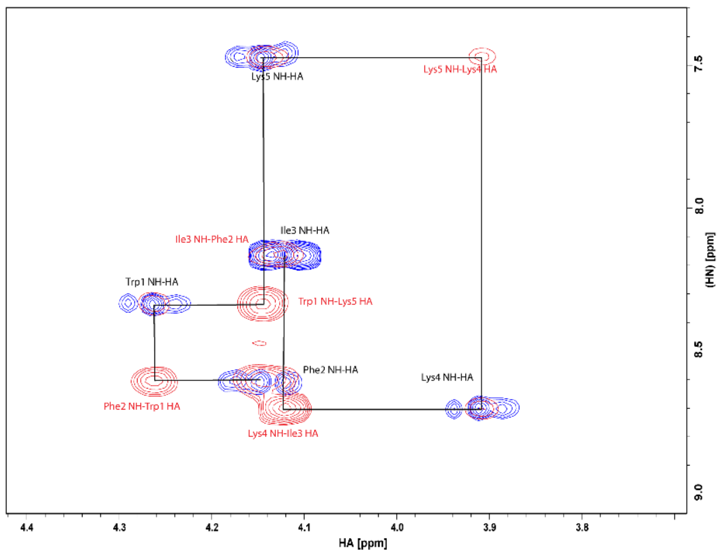

**Figure S29.** Overlay of $^1$H-$^1$H NOESY and $^1$H-$^1$H COSY spectrum (DMSO-d$_6$, 500 MHz) of **1**.

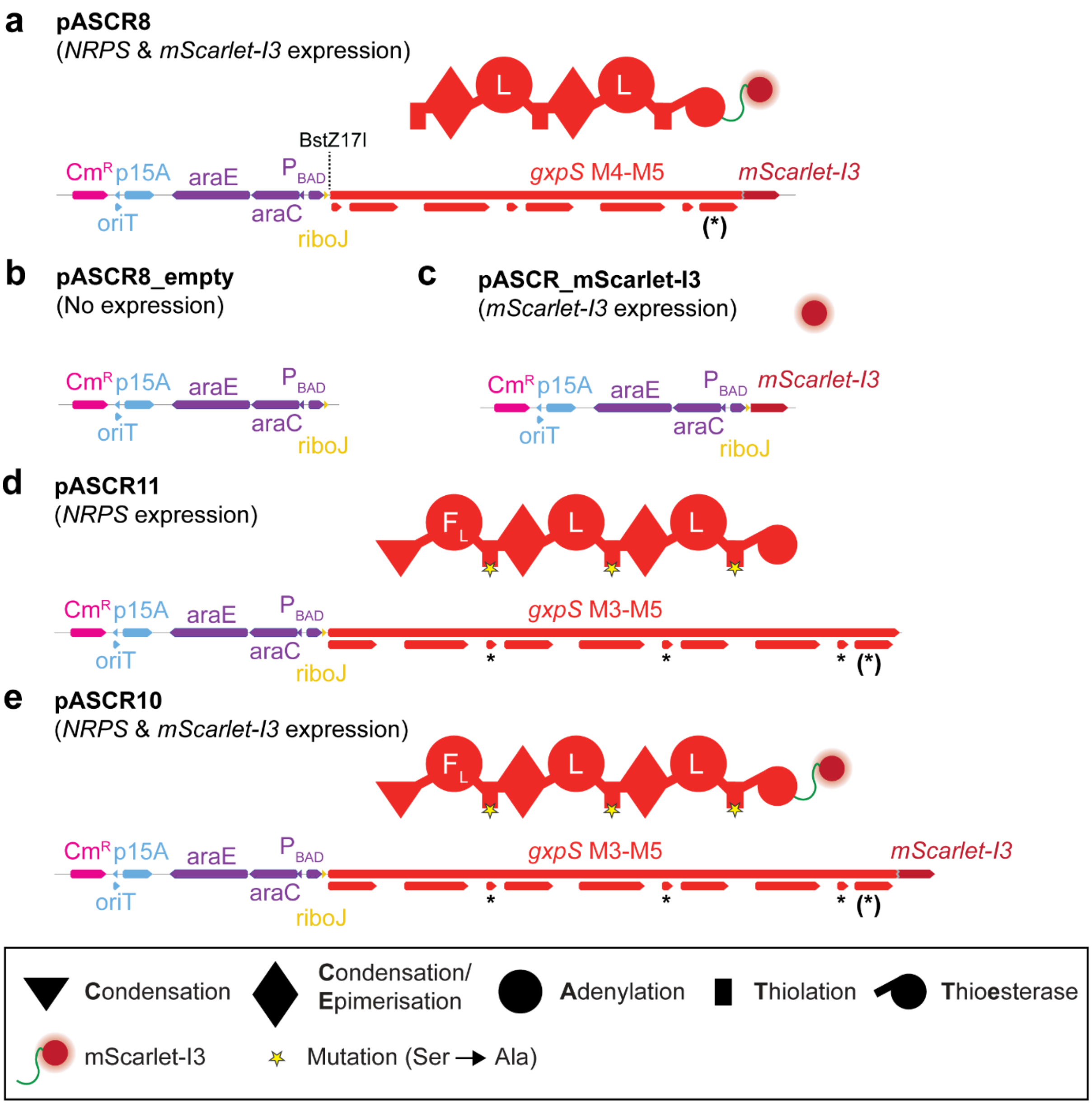

**Figure S30.** pASCR8 and control vector construction. Linearized illustration of the plasmid maps of pASCR8 (**a**), pASCR8_empty (**b**), pASCR_mScarlet-I3 (**c**), pASCR11 (**d**) and pASCR10 (**e**). Indication of the resistance genes, origin of replications, promoter regions, NRPS-encoding regions and added tags. An asterisk (*) indicates the introduced mutations within the T domains to inactivate the NRPS. An asterisk within brackets indicates a silent mutation at G2315 (GGT→GGG) to ensure single cleavage by the restriction enzyme. For pASCR11 and pASCR12 the point mutation is present at G3274 (GGT→GGG). Schematic illustration of the encoded protein above. The corresponding NRPS domain and protein-tag legend is depicted below.

**a**

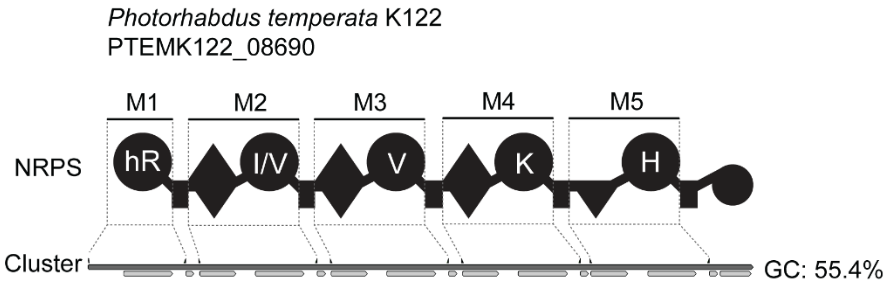

**b**

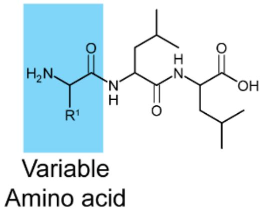

| Module | Detected mass $[M+H]^+$ | Δppm | Sum formula | Estimated amino acid |
|---|---|---|---|---|
| M1 | 415.3026 | 0.4 | $C_{19}H_{39}N_6O_4$ | hR |
| M2 | 344.2545 | -0.2 | $C_{17}H_{34}N_3O_4$ | Val |
| M2 | 358.2700 | 0.0 | $C_{18}H_{36}N_3O_4$ | Ile |
| M3 | 344.2542 | 0.5 | $C_{17}H_{34}N_3O_4$ | Val |
| M4 | 373.2808 | 0.4 | $C_{18}H_{37}N_4O_4$ | Lys |
| M5 | 382.2449 | -0.1 | $C_{18}H_{32}N_5O_4$ | His |

**c**

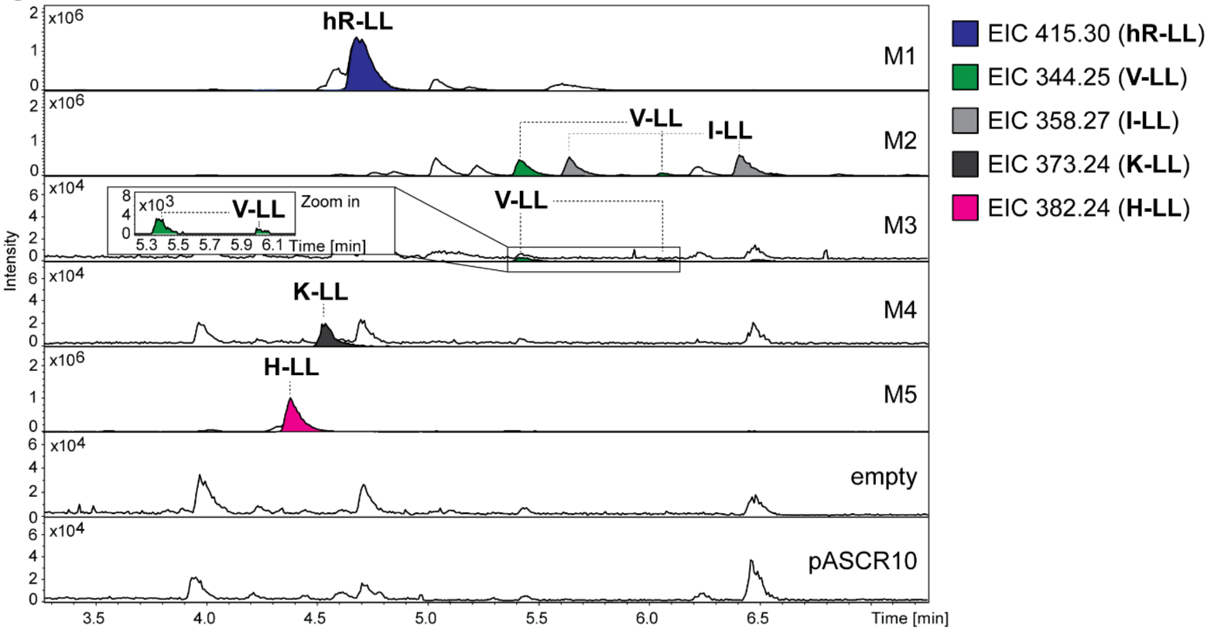

**Figure S31.** HPLC-MS/MS data referred to module screening of PTEMK122_08690 from *P. temperata* K122. (**a**) Illustration of the gene cluster PTEMK122_08690 from *P. temperata* K122 with schematic illustration of the encoded NRPS. Indication of the region of cloned modules M1 to M5 on gene and protein level. Utilized primers are shown above the cluster. A domain specitifity is indicated by the one-letter amino acid code (hR = homoarginine). GC content is depicted next to the cluster. (**b**) Table of HPLC-MS/MS data indicating the targeted module cloned into pASCR8, the detected mass of the compound, the corresponding error, the sum formula and the incoporated amino acid introduced to the peptide at the highlighted position in the chemical structure (left). The amino acid is shown by the three-letter amino acid code (hR = homoarginine). (**c**) HPLC-MS/MS data of *E. coli* DH10B::*mtaA* expressing M1 to M5 cloned into pASCR8. Expression of an empty vector and pASCR10 served as negative control. Base Peak Chromatogram (BPC, top) with Extracted Ion Chromatogram (EIC, below with colors according to the depicted legend) of **hR-LL** (*m/z* $[M+H]^+$ = 415.302), **V-LL** (*m/z* $[M+H]^+$ = 344.254), **I-LL** (*m/z* $[M+H]^+$ = 358.270), **K-LL** (*m/z* $[M+H]^+$ = 373.280) and **H-LL** (*m/z* $[M+H]^+$ = 382.244). Indication of the targeted modules on the right of the chromatograms.

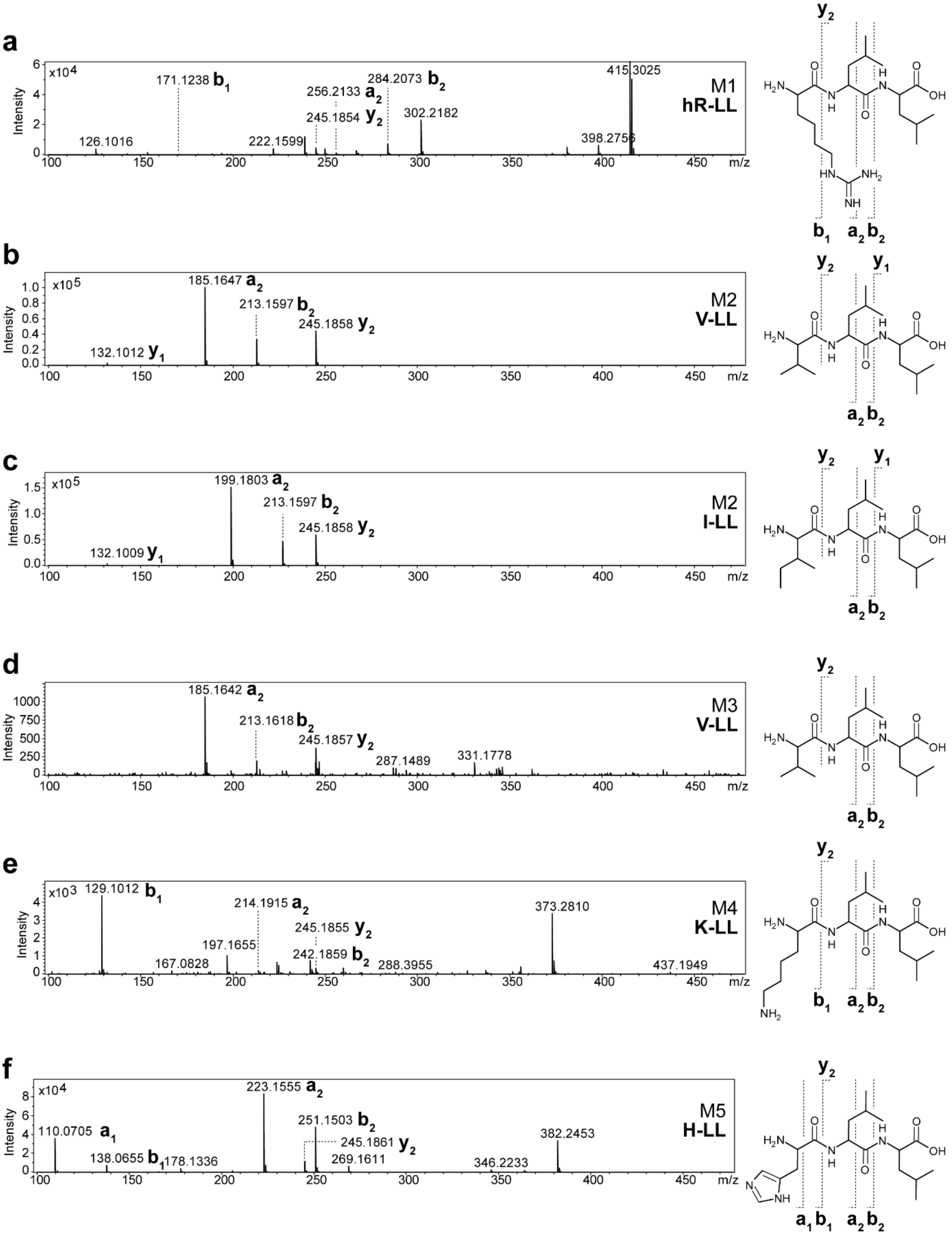

**Figure S32.** $MS^2$ spectra of HPLC-MS/MS data referred to figure S31. (**a-f**) The respective compounds are displayed, along with indications of the assigned fragments. The targeted module and peptide name is depicted within the $MS^2$ spectra.

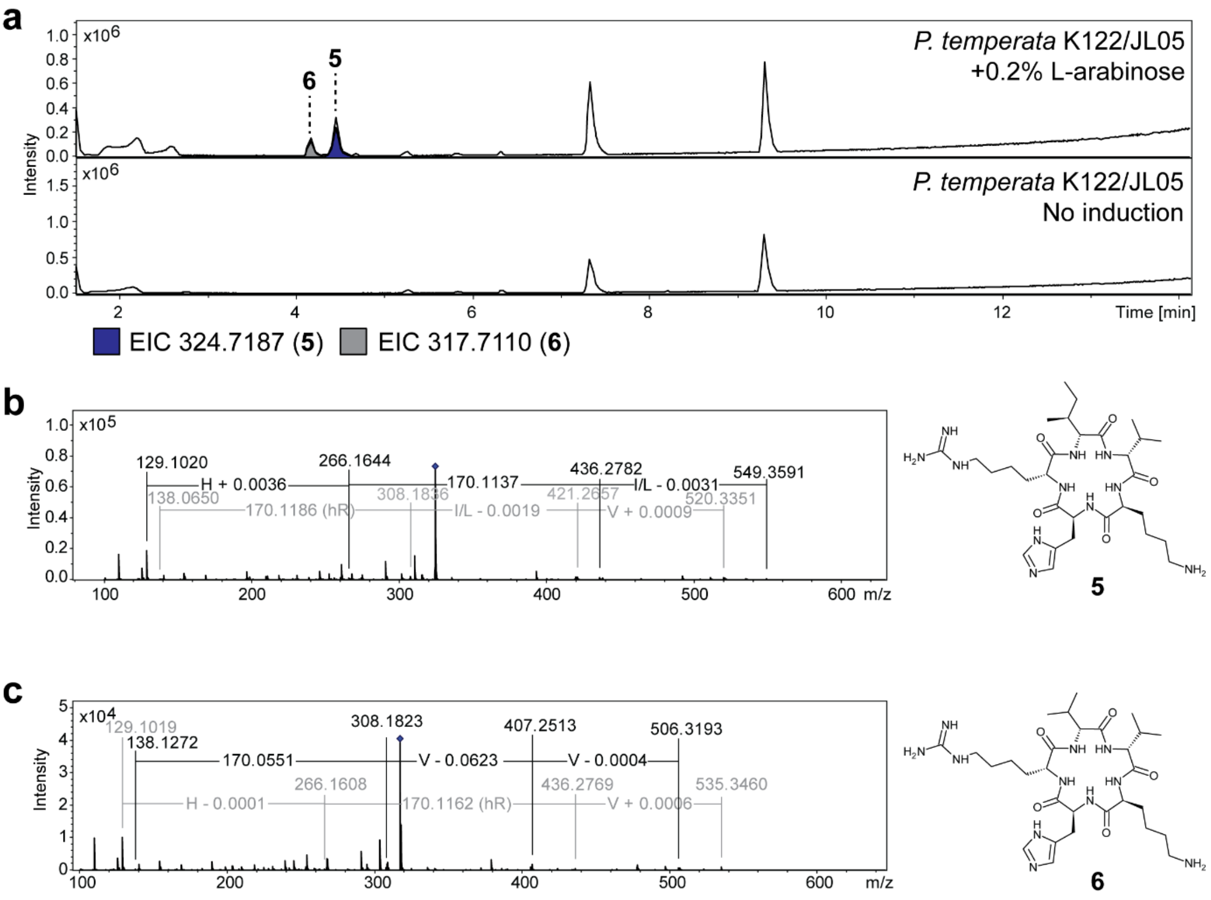

**d**

| Compound | Detected mass $[M+2H]^{2+}$ | Δppm | Sum formula |
|---|---|---|---|
| **5** | 324.7187 | 0.3 | $C_{30}H_{55}N_{11}O_5$ |
| **6** | 317.7109 | 0.4 | $C_{29}H_{53}N_{11}O_5$ |

**Figure S33.** HPLC-MS/MS data of *P. temperata* K122 $P_{BAD}$ promoter exchange based activation of PTEMK122_08690. (**a**) HPLC-MS/MS data of *P. temperata* K122/JL05 cultivated in XPP medium with and without induction with 0.2% L-arabinose. Comparison of Base Peak Chromatogram (BPC) is displayed with corresponding Extracted Ion Chromatogram (EIC) of compound **5** (*m/z* $[M+2H]^{2+}$ = 324.7179) and **6** (*m/z* $[M+2H]^{2+}$ = 317.7102) depicted below. $MS^2$ spectra of HPLC-MS/MS data of **5** (**b**) and **6** (**c**) are displayed, along with the chemical structures. Mass differences are indicated within the spectra. (**d**) High-resolution MS data of compound **5** and **6** with indications of the detected mass, the error and the sum formula.

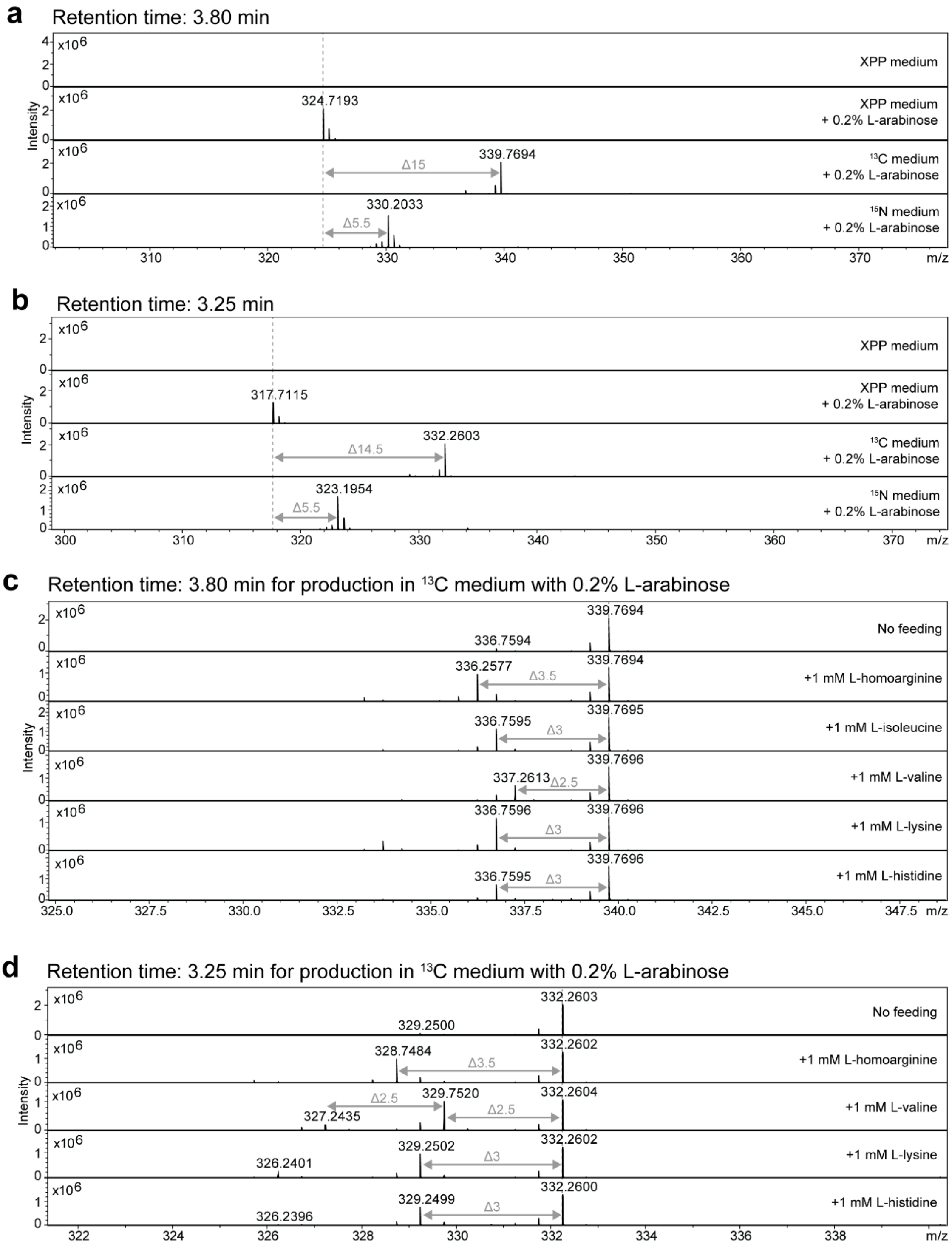

**Figure S34.** Isotope and inverse istope feeding experiments for structure determination of compounds **5** and **6**. HPLC-MS analysis of *P. temperata* K122/JL05. Determination of carbon and nitrogen atoms in compounds **5** (*m/z* $[M+2H]^{2+}$ = 324.7187) (**a**) and **6** (*m/z* $[M+2H]^{2+}$ = 317.7110) (**b**) by isotope labelling experiments. MS spectra are displayed for the strain cultivated in XPP medium ($^{12}C$, $^{14}N$), fully labelled $^{13}C$ medium ($^{13}C$, $^{14}N$) and $^{15}N$ medium ($^{12}C$, $^{15}N$) with indication of the retention time and mass shifts. In addition, MS spectra of inverse feeding experiments are shown, performed in fully labelled $^{13}C$ medium ($^{13}C$, $^{14}N$) supplemented with unlabelled L-homoarginine, L-isoleucine, L-valine, L-lysine and L-hisitidine. MS spectra retention times are shown with indications of mass shifts for the respective compounds **5** (**c**) and **6** (**d**).

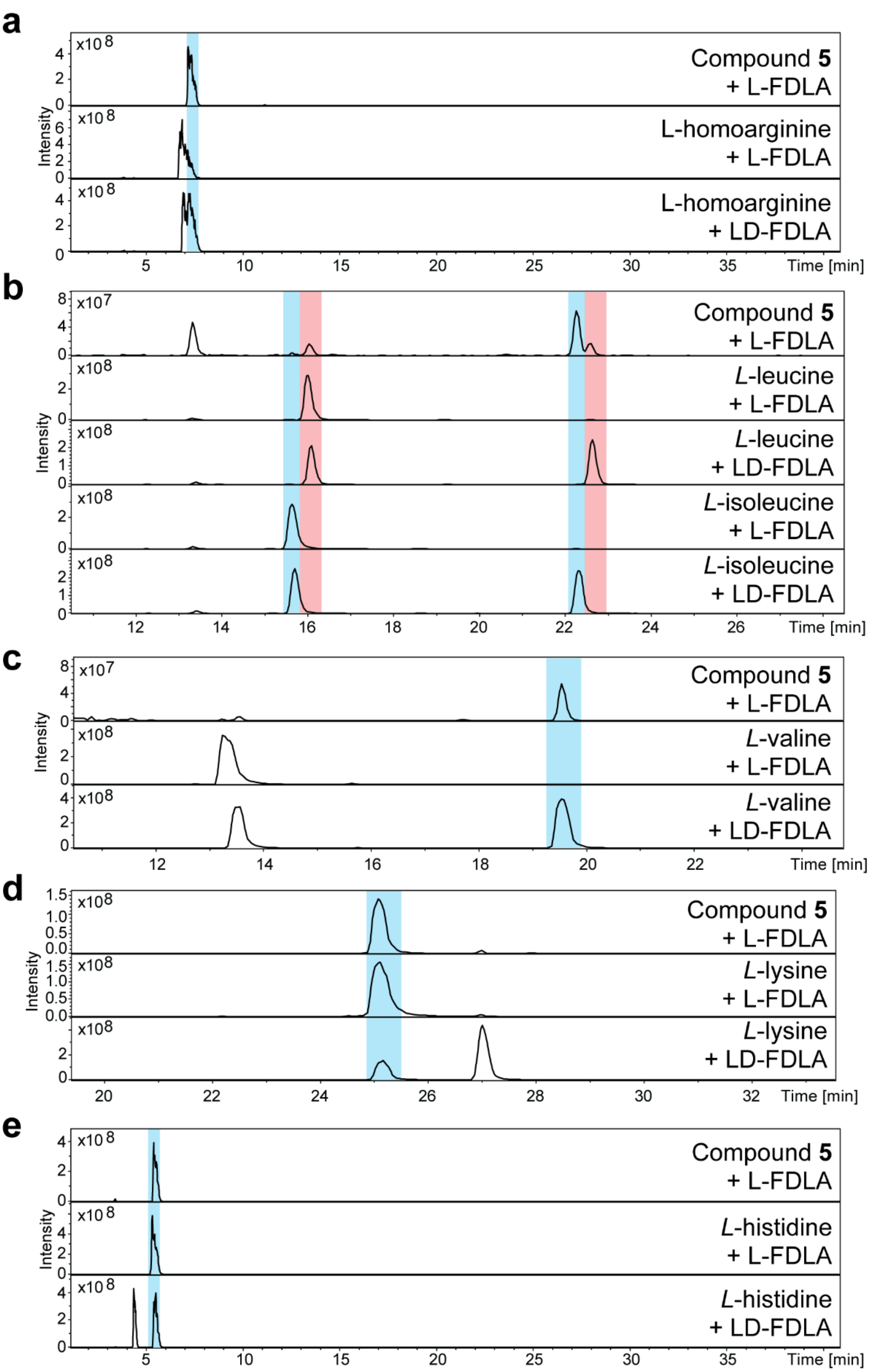

**Figure 35**. Configuration determination of amino acids in **5** using the advanced Marfey's method. HPLC-MS analysis of hydrolyzed and L-FDLA derivatized **5** along with derivatized amino acid standards with L-FDLA and LD-FDLA. Depicted are EIC traces for Marfey's derivatives of L-homoarginine (hR, *m/z* 483.23 $[M+H]^+$)(**a**), L-leucine and L-leucine (ile,leu *m/z* 426.20 $[M+H]^+$)(**b**), L-valine (Val, *m/z* 412.20 $[M+H]_+$)(**c**), L-di-lysine Marfey derivative (Lys, *m/z* 735.31 $[M+H]^+$)(**d**) and L-histidine (His, *m/z* 450.17 $[M+H]^+$)(**e**).

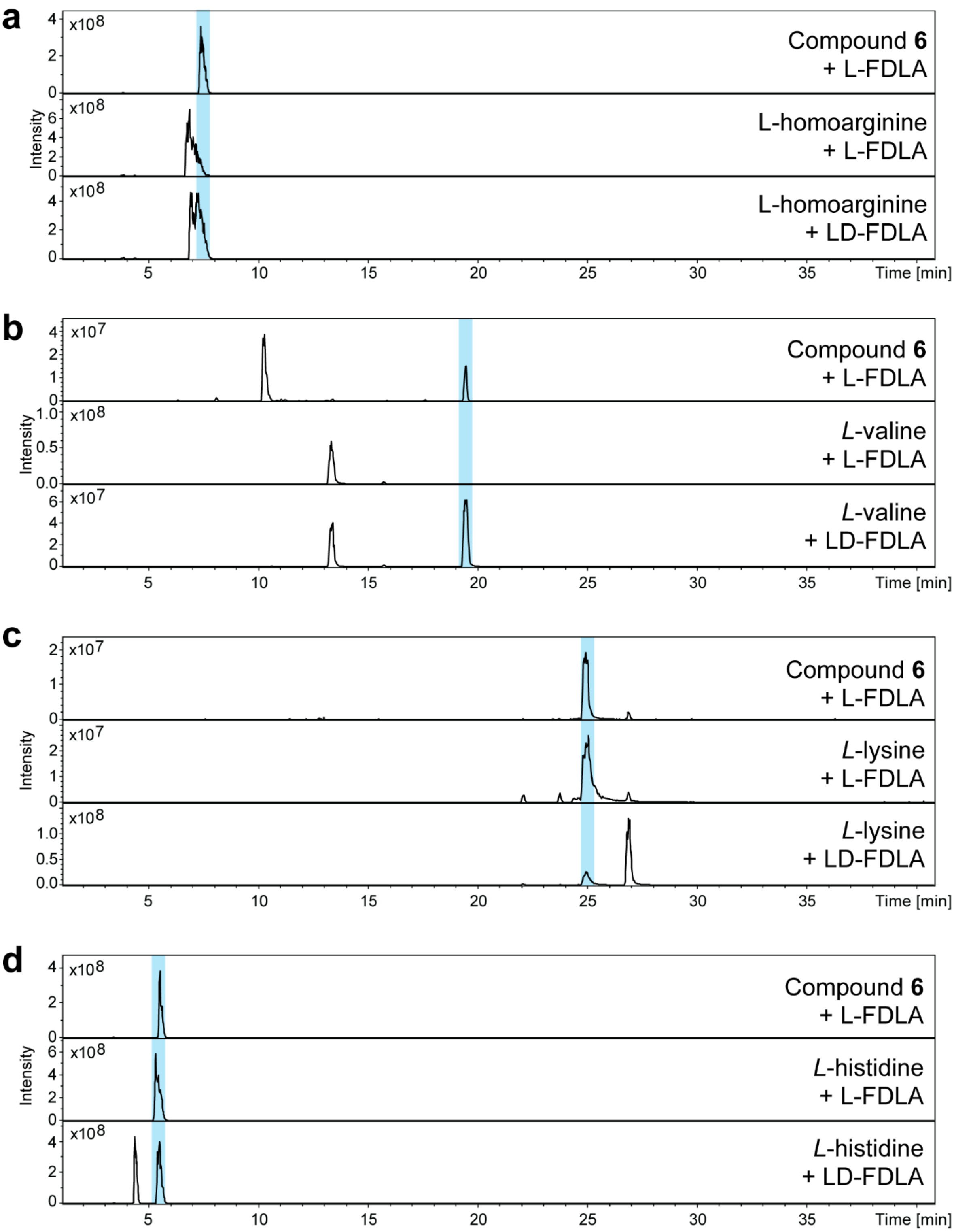

**Figure 36**. Configuration determination of amino acids in **6** using the advanced Marfey's method. HPLC-MS analysis of hydrolyzed and L-FDLA derivatized **6** along with derivatized amino acid standards with L-FDLA and LD-FDLA. Depicted are EIC traces for Marfey's derivatives of L-homoarginine (hR, *m/z* 483.23 $[M+H]^+$)(**a**), L-valine (Val, *m/z* 412.20 $[M+H]^+$)(**b**), L-di-lysine Marfey derivative (Lys, *m/z* 735.31 $[M+H]^+$)(**c**) and L-histidine (His, *m/z* 450.17 $[M+H]^+$)(**d**).

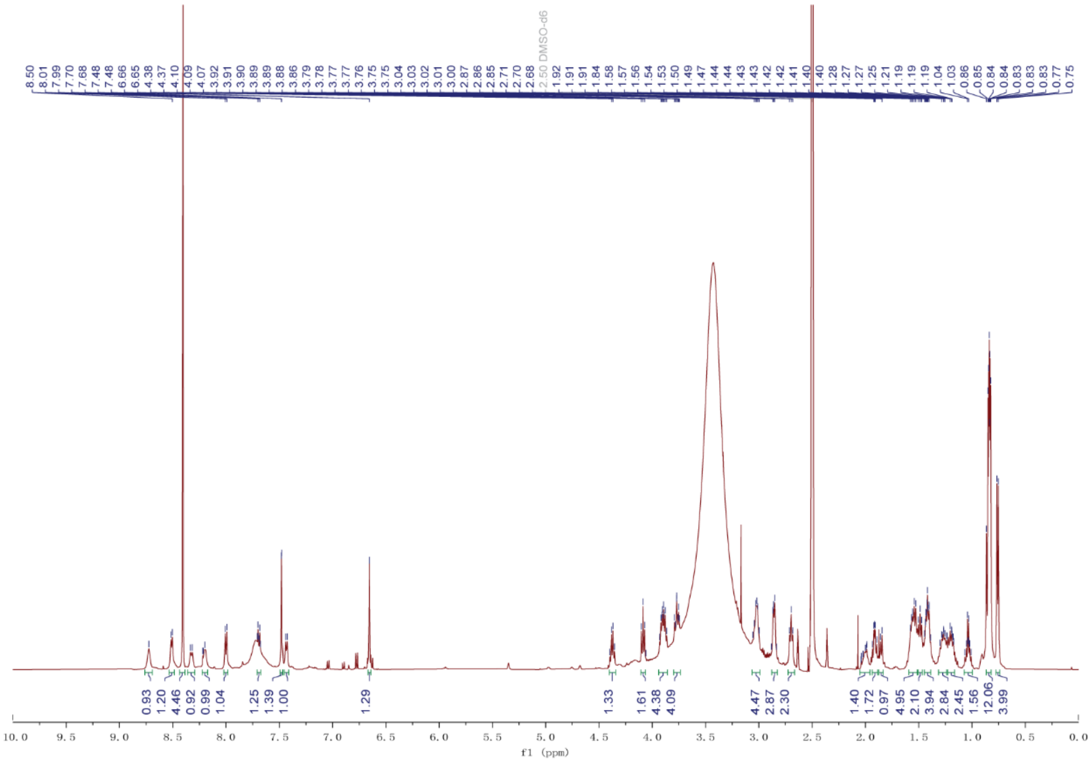

**Figure S37.** $^{1}H$ NMR spectrum (DMSO-$d_6$, 500 MHz) of **5**.

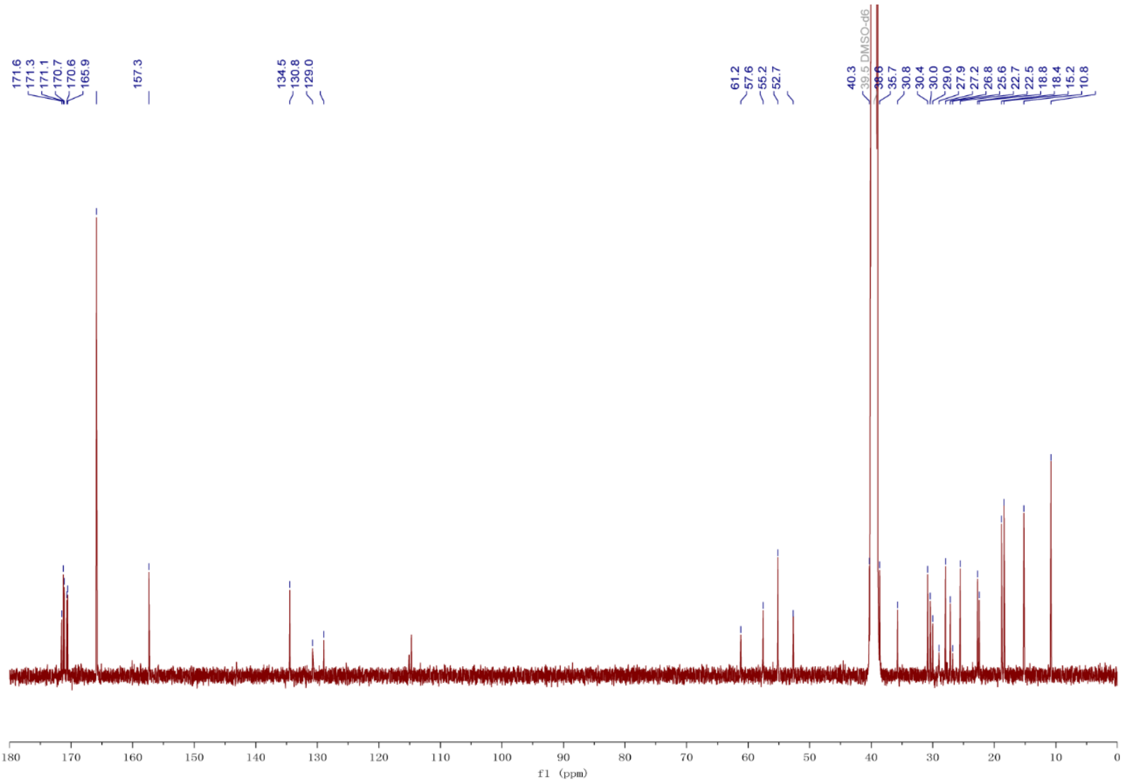

**Figure S38.** $^{13}C$ NMR spectrum (DMSO-$d_6$, 125 MHz) of **5**.

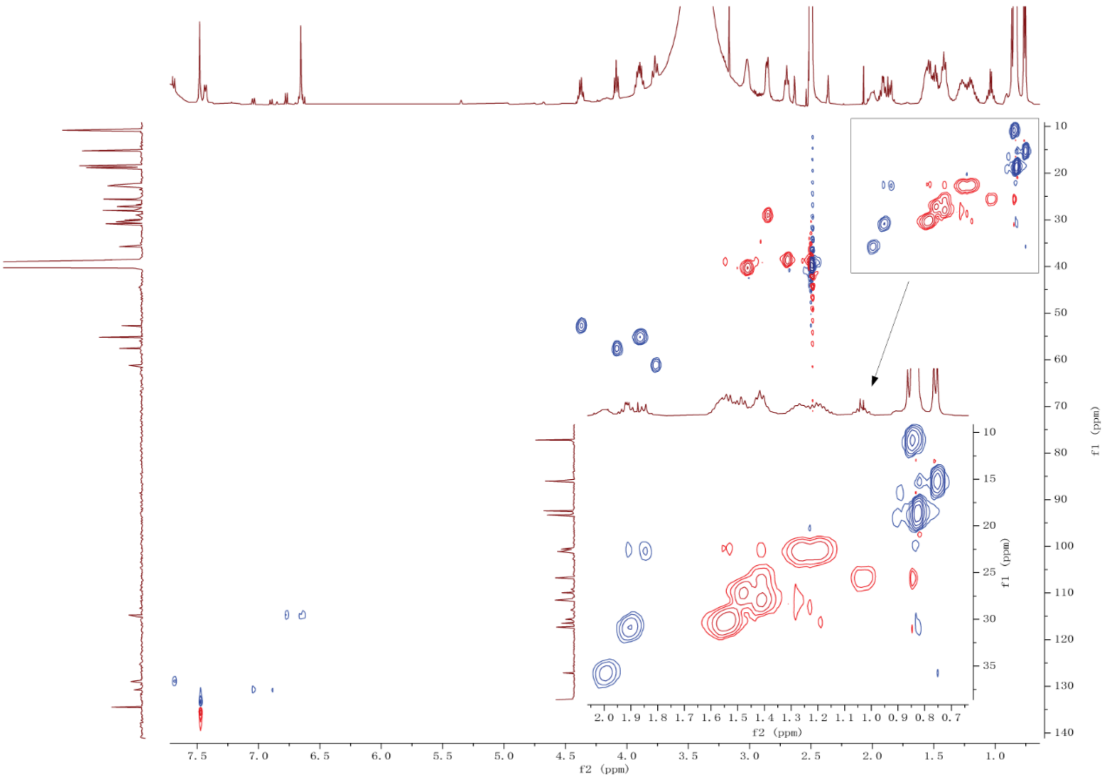

**Figure S39.** HSQC NMR spectrum (DMSO-d$_6$, 500 MHz) of **5**.

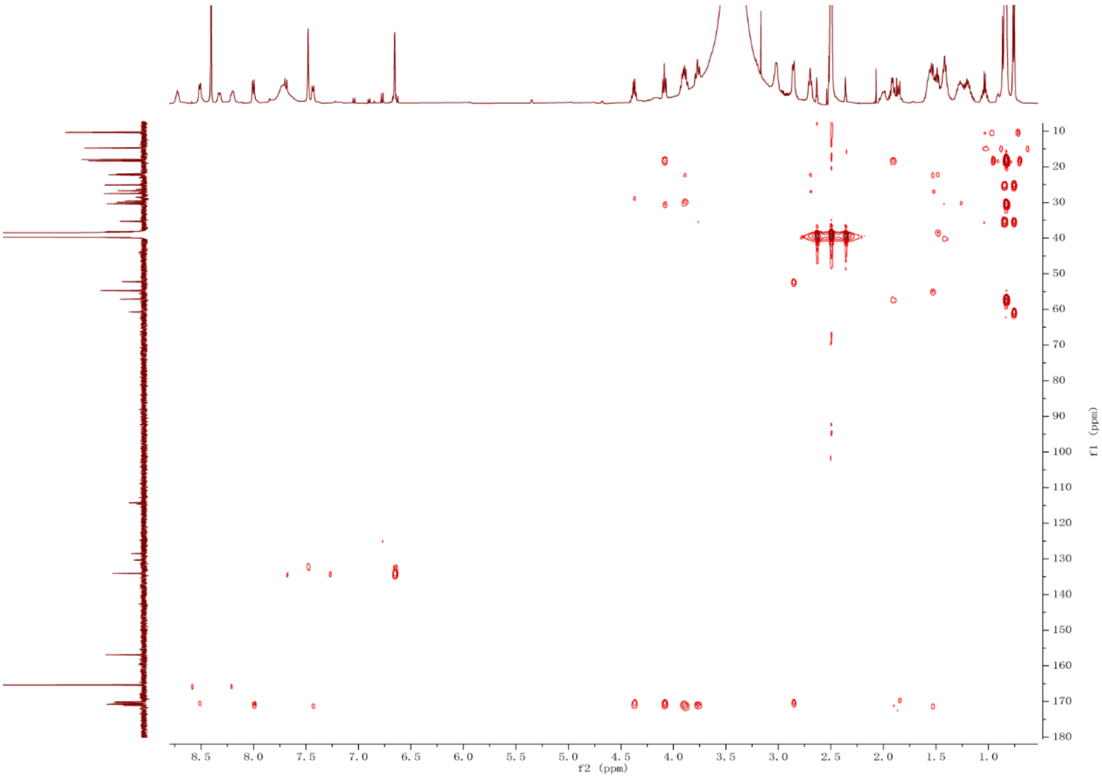

**Figure S40.** HMBC spectrum (DMSO-d$_6$, 500 MHz) of **5**.

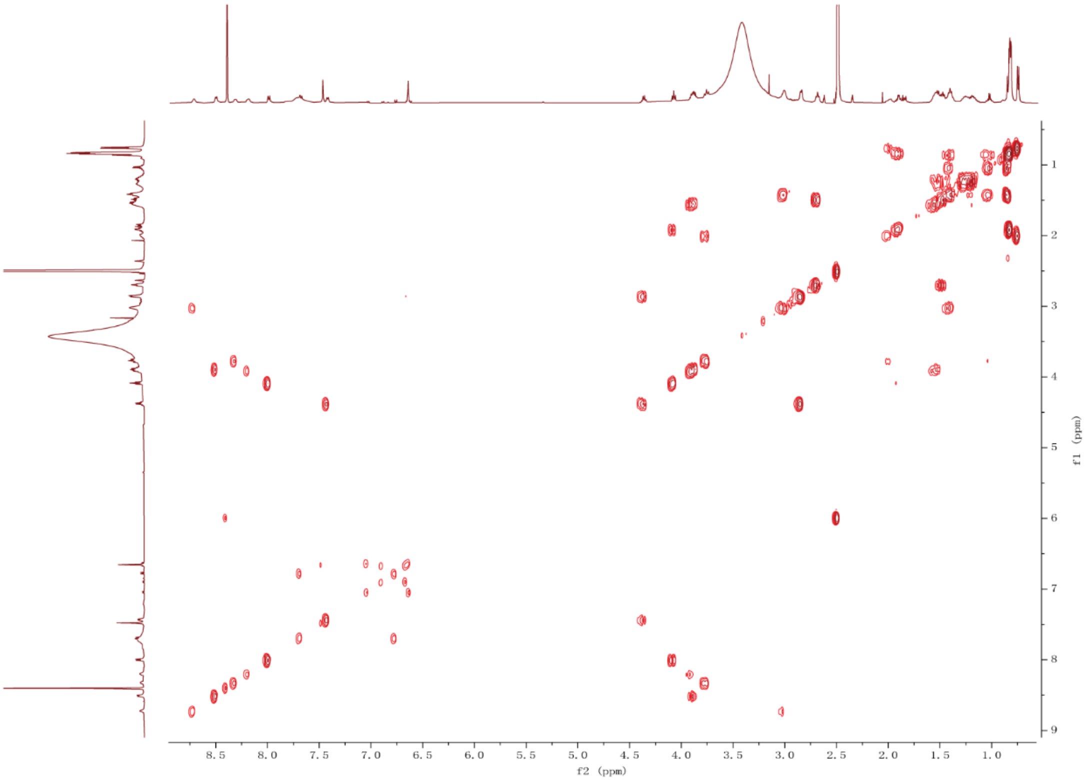

**Figure S41.** $^{1}H$-$^{1}H$ COSY spectrum (DMSO-d$_6$, 500 MHz) of **5**.

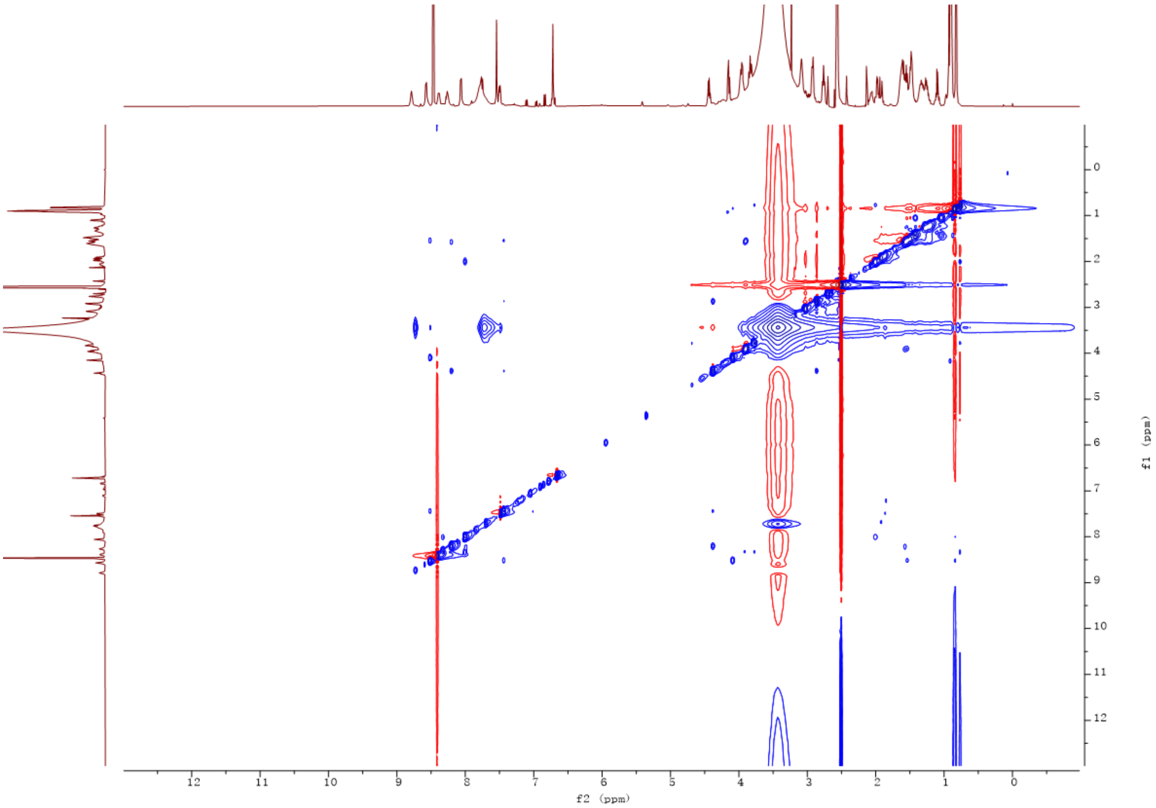

**Figure S42.** NOESY spectrum (DMSO-d$_6$, 500 MHz) of **5**.

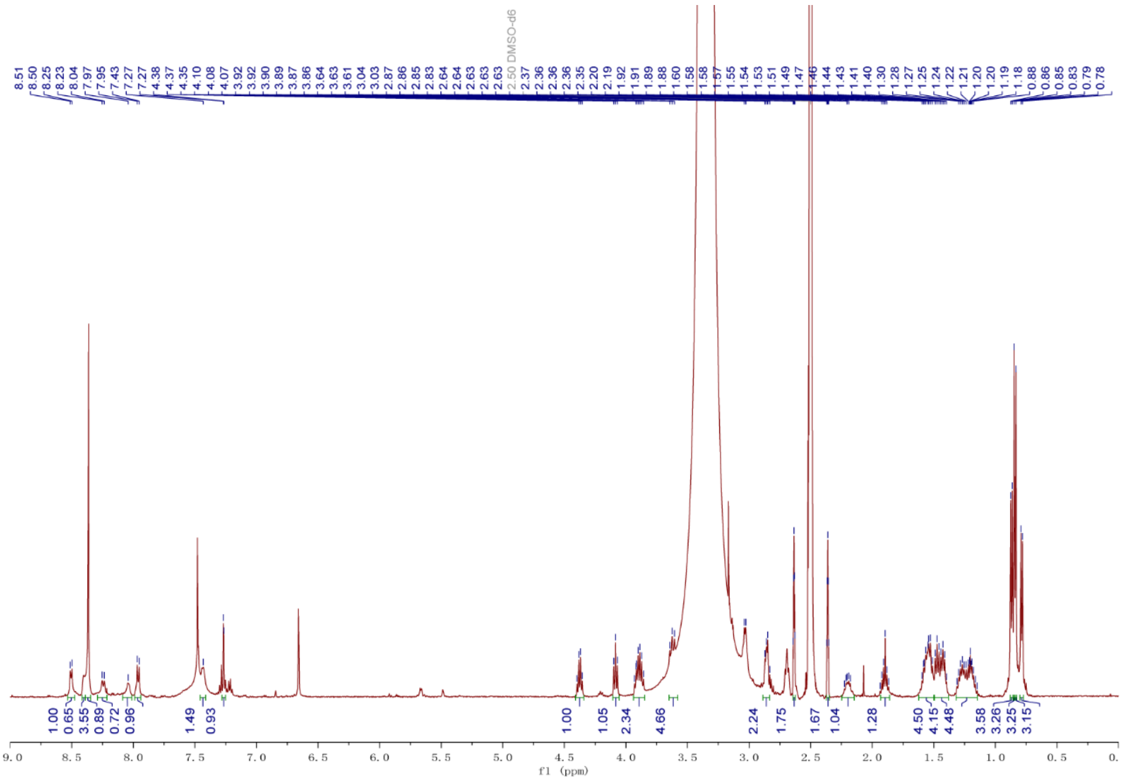

**Figure S43.** $^{1}H$ NMR spectrum (DMSO-$d_6$, 500 MHz) of **6**.

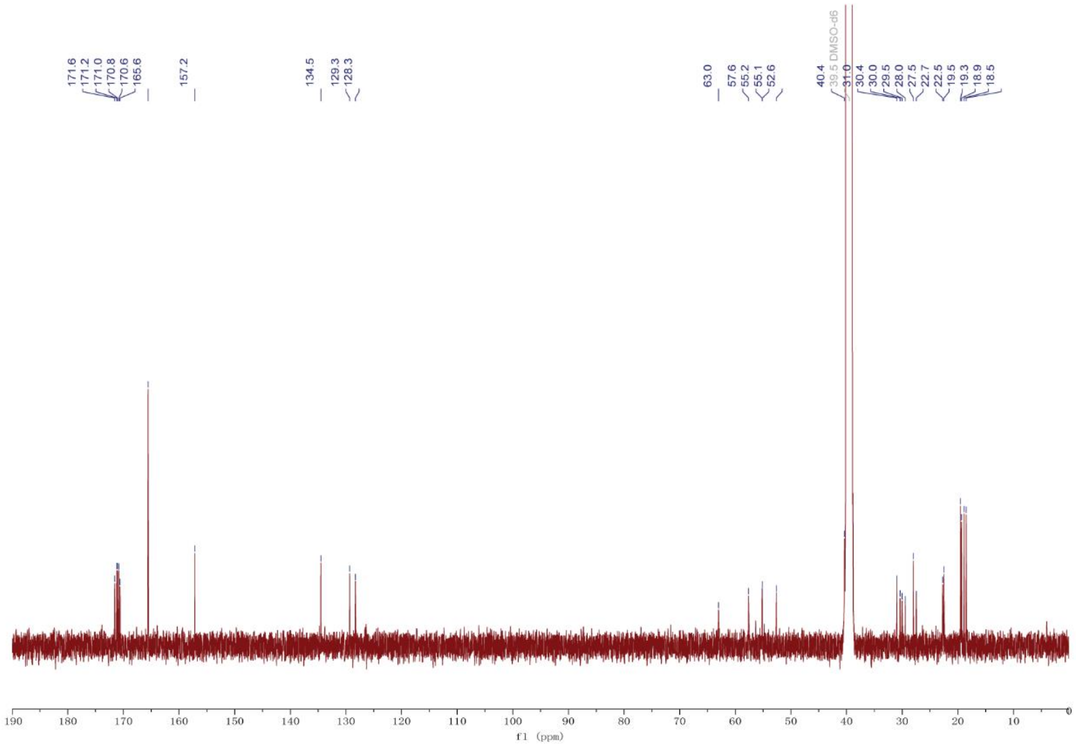

**Figure S44.** $^{13}C$ NMR spectrum (DMSO-$d_6$, 125 MHz) of **6**.

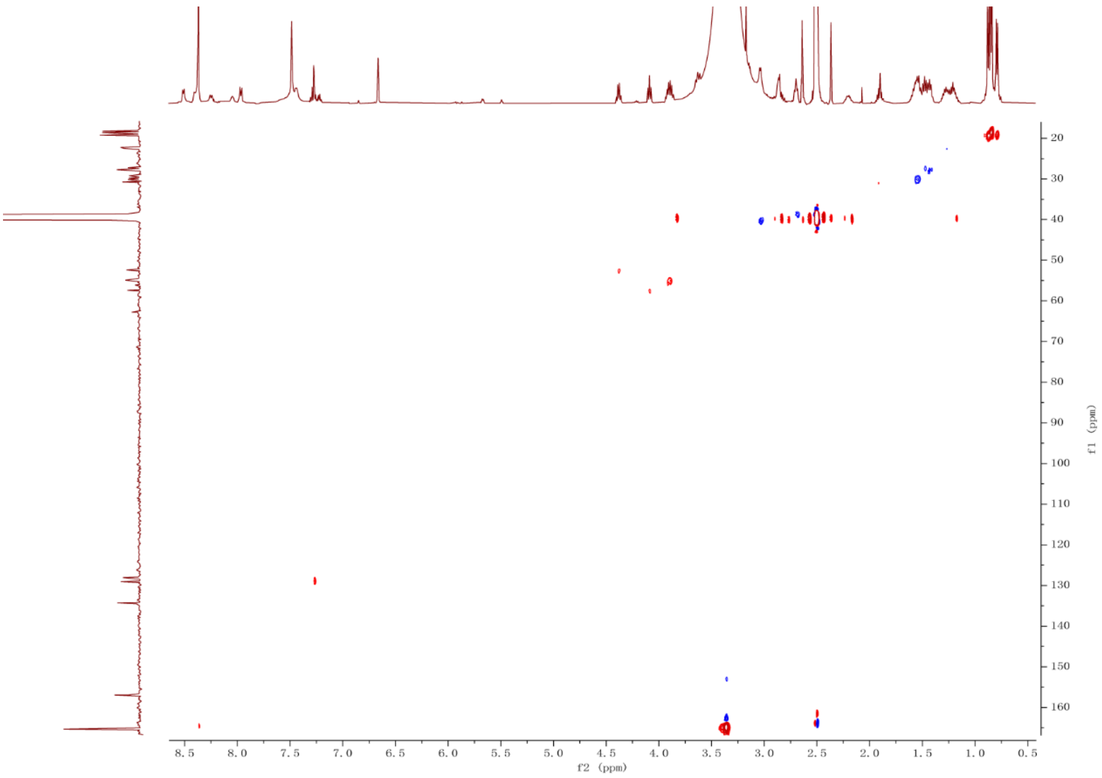

**Figure S45.** HSQC NMR spectrum (DMSO-d$_6$, 500 MHz) of **6**.

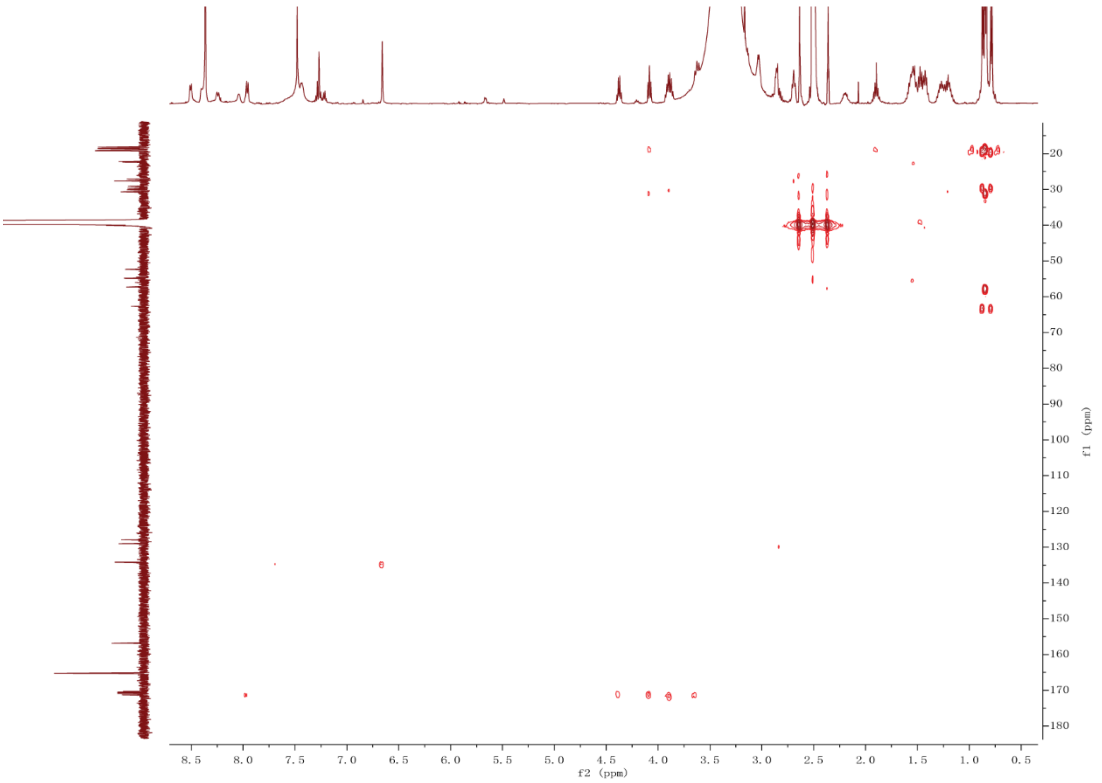

**Figure S46.** HMBC spectrum (DMSO-d$_6$, 500 MHz) of **6**.

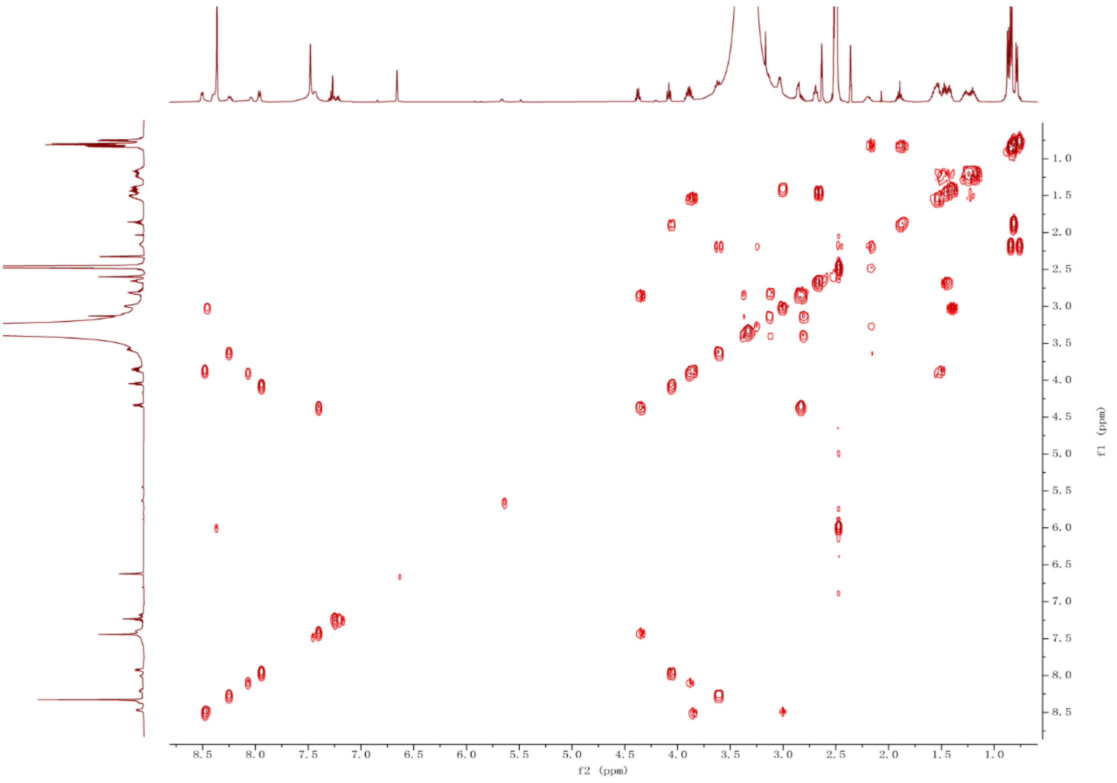

**Figure S47.** $^{1}H$-$^{1}H$ COSY spectrum (DMSO-$d_6$, 500 MHz) of **6**.

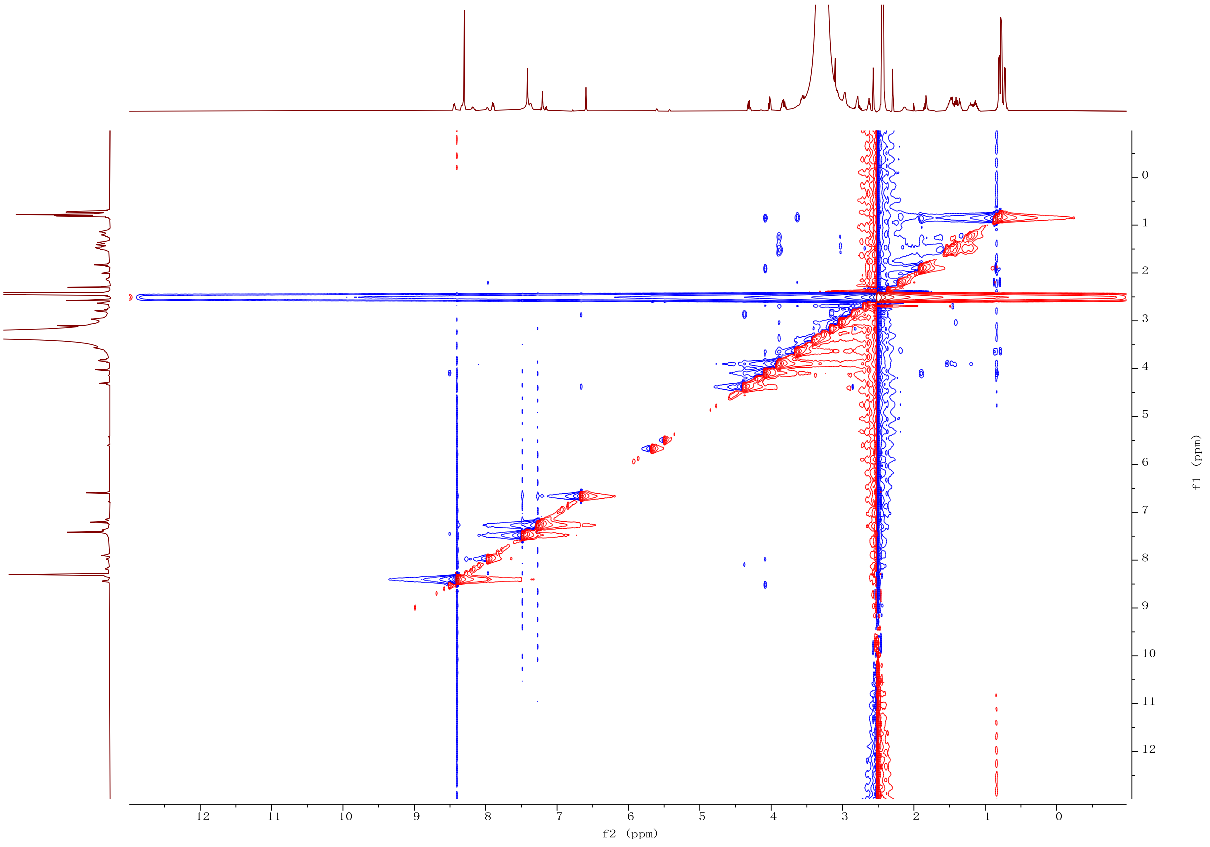

**Figure S48.** NOESY spectrum (DMSO-$d_6$, 500 MHz) of **6**.

**a**

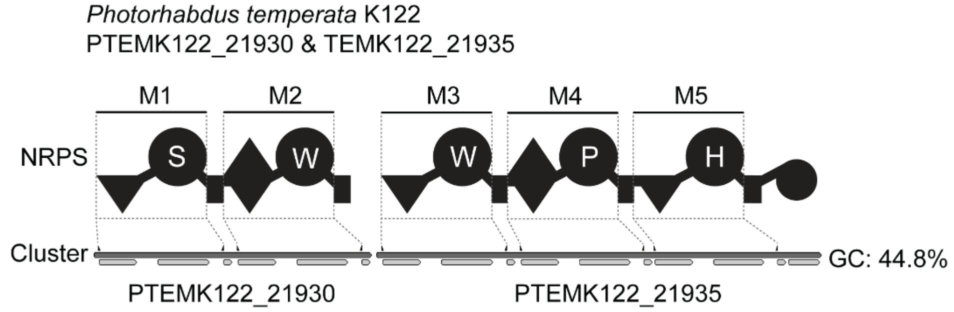

**b**

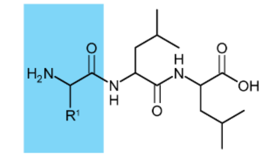

| Module | Detected mass $[M+H]^+$ | Δppm | Sum formula | Estimated amino acid |
|---|---|---|---|---|
| M1 | 332.2182 | -0.7 | $C_{15}H_{30}N_3O_5$ | Ser |
| M2 | 431.2653 | 0.0 | $C_{23}H_{35}N_4O_4$ | Trp |
| M3 | 431.2651 | 0.5 | $C_{23}H_{35}N_4O_4$ | Trp |
| M4 | 342.2390 | -0.7 | $C_{17}H_{32}N_3O_4$ | Pro |
| M5 | 382.2450 | -0.3 | $C_{18}H_{32}N_5O_4$ | His |

**c**

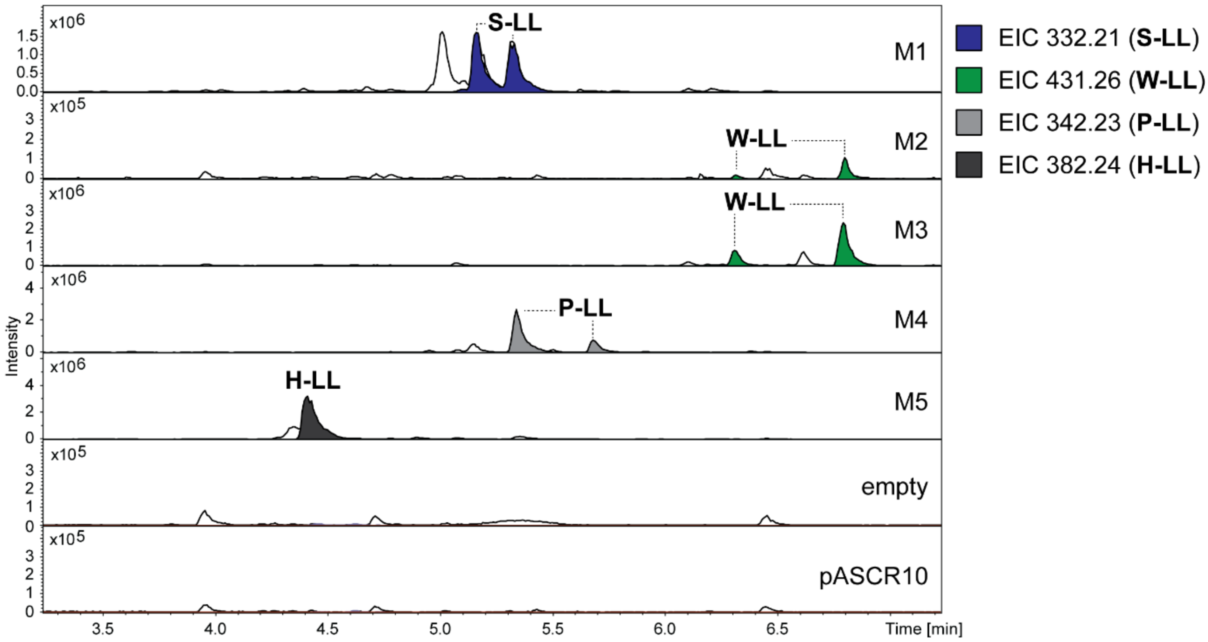

**Figure S49.** HPLC-MS/MS data referred to module screening of PTEMK122_21930 and PTEMK122_21935 from *P. temperata* K122. (**a**) Illustration of the gene cluster PTEMK122_21930 and PTEMK122_21935 from *P. temperata* K122 with schematic illustration of the encoded NRPS. Indication of the region of cloned modules M1 to M5 on gene and protein level. Utilized primers are shown above the cluster. A domain specitifity is indicated by the one-letter amino acid code. GC content is depicted next to the cluster. (**b**) Table of HPLC-MS/MS data indicating the targeted module cloned into pASCR8, the detected mass of the compound, the corresponding error, the sum formula and the incoporated amino acid introduced to the peptide at the highlighted position in the chemical structure (left). The amino acid is shown by the three-letter amino acid code. (**c**) HPLC-MS/MS data of *E. coli* DH10B::*mtaA* expressing M1 to M5 cloned into pASCR8. Expression of an empty vector and pASCR10 served as negative control. Base Peak Chromatogram (BPC, top) with Extracted Ion Chromatogram (EIC, below with colors according to the depicted legend) of **S-LL** (*m/z* $[M+H]^+$ = 332.217), **W-LL** (*m/z* $[M+H]^+$ = 431.265), **P-LL** (*m/z* $[M+H]^+$ = 342.254) and **H-LL** (*m/z* $[M+H]^+$ = 382.244). Indication of the targeted modules on the right of the chromatograms.

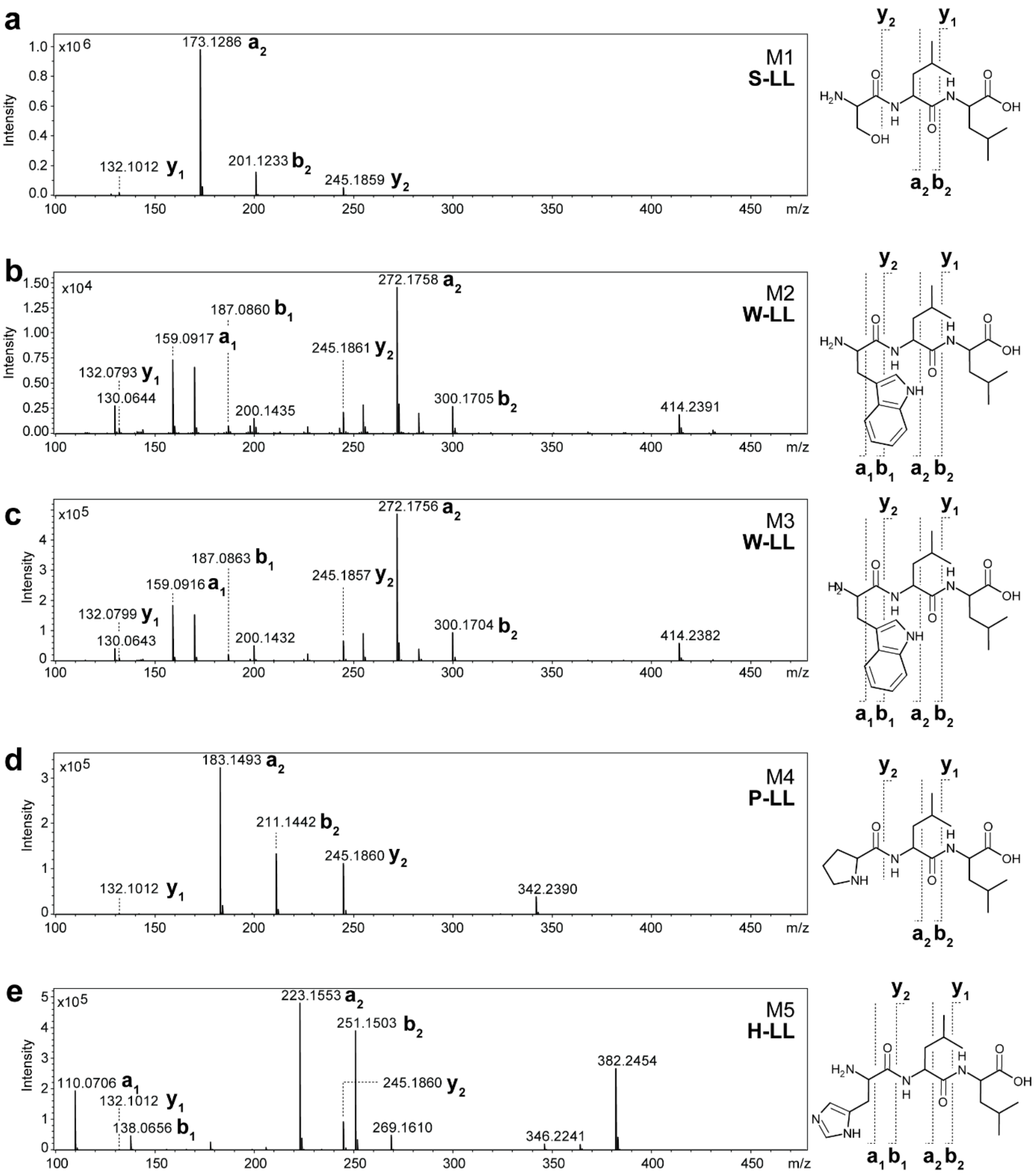

**Figure S50.** $MS^2$ spectra of HPLC-MS/MS data referred to figure S49. (**a-e**) The respective compounds are displayed, along with indications of the assigned fragments. The targeted module and peptide name is depicted within the $MS^2$ spectra.

a

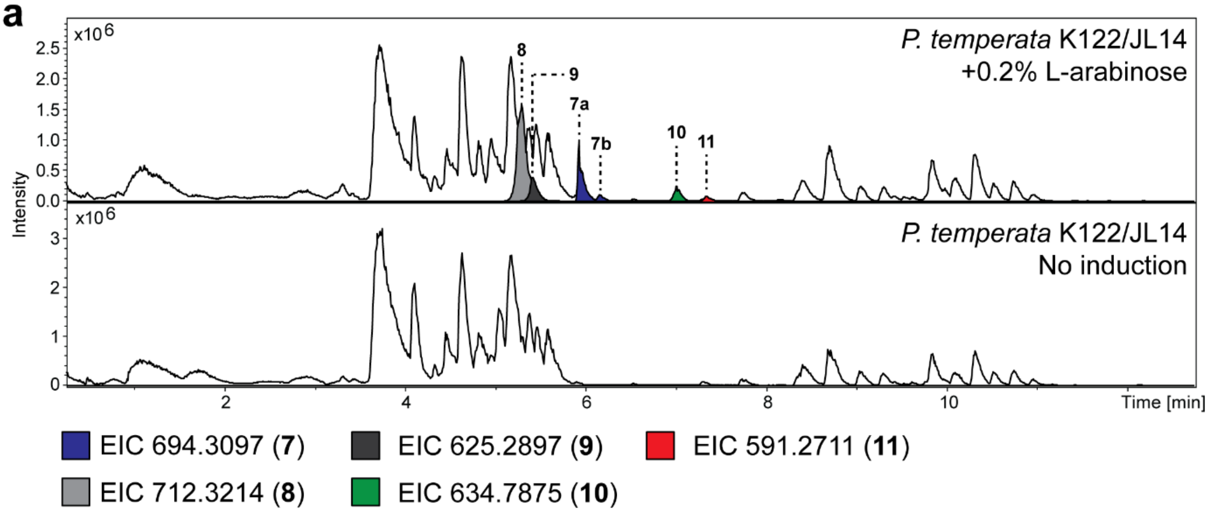

b

| Compound | Detected mass $[M+H]^+$ | Δppm | Sum formula | Assigned peptide |
|---|---|---|---|---|
| **7a** | 694.3097 | -0.1 | $C_{36}H_{40}N_9O_6$ | cyclic(SwWPH) |
| **7b** | 694.3096 | 0.0 | $C_{36}H_{40}N_9O_6$ | cylic(swWPH) |
| **8** | 712.3206 | -0.6 | $C_{36}H_{42}N_9O_7$ | SwWPH |
| **9** | 625.2880 | 0.3 | $C_{33}H_{37}N_8O_5$ | wWPH |
| **10** | 634.7854* | 0.0 | $C_{66}H_{75}N_{15}O_{12}$ | SwWPSwWPH |
| **11** | 591.2695* | -0.2 | $C_{63}H_{70}N_{14}O_{10}$ | wWPSwWPH |

* indicates detection of $[M+2H]^{2+}$

**Figure S51.** HPLC-MS/MS data of *P. temperata* K122 $P_{BAD}$ promoter exchange based activation of PTEMK122_21930. (**a**) HPLC-MS/MS data of *P. temperata* K122/JL14 cultivated in low salt LB medium with and without induction with 0.2% L-arabinose. Comparison of Base Peak Chromatogram (BPC) is displayed with corresponding Extracted Ion Chromatogram (EIC) of compound **7a/b** (*m/z* $[M+H]^+$ = 694.3105), **8** (*m/z* $[M+H]^+$ = 712.3214), **9** (*m/z* $[M+H]^+$ = 625.2897), **10** (*m/z* $[M+2H]^{2+}$ = 634.7875) and **11** (*m/z* $[M+2H]^{2+}$ = 591.2711) depicted below. (**b**) High-resolution MS data of compound **7** to **11** with indications of the detected mass, the error, the sum formula and an assigned peptide structure represented by the one-letter amino acid code. Capital letter indicate L-configuration and lowercase letter D-configuration.

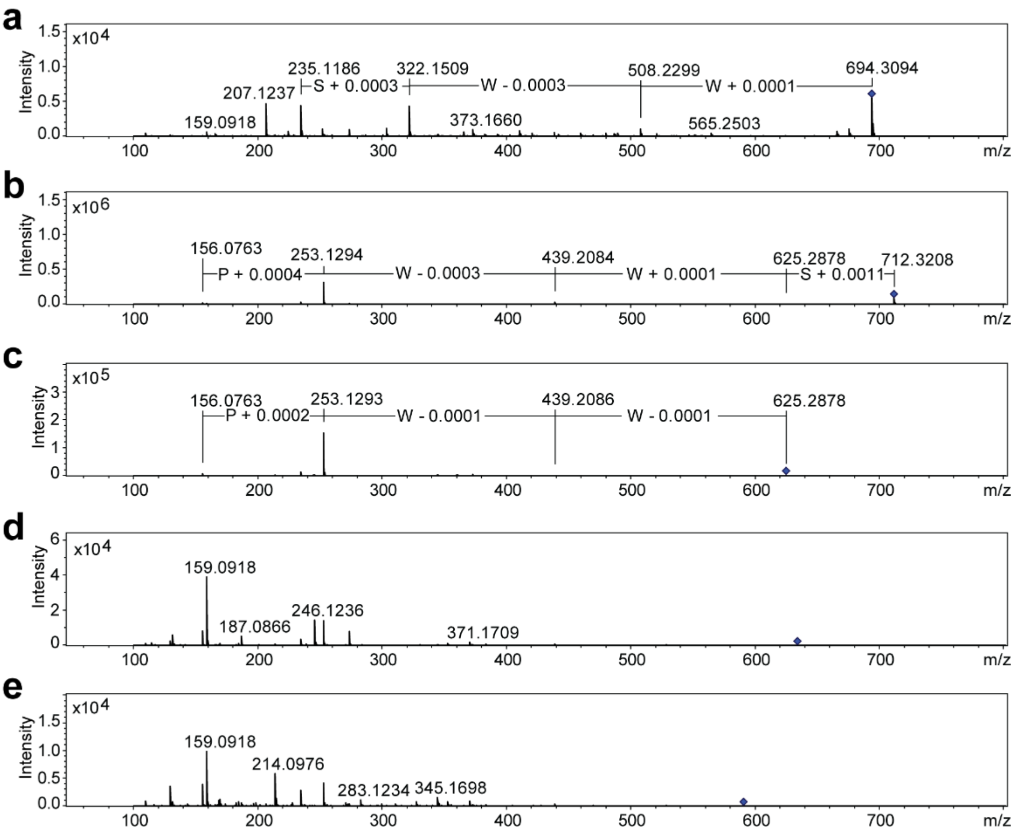

**7a** cyclic(SwWPH)

**8** SwWPH

**9** wWPH

**10** SwWPSwWPH

**11** wWPSwWPH

**Figure S52.** $MS^2$ spectra of HPLC-MS/MS data referred to figure S51. (**a-e**) Assigned peptide structures of **7a** to **11** are displayed as one-letter amino acid code peptide sequence along with the $MS^2$ fragmentation spectra. Capital letter indicate L-configuration and lowercase letter D-configuration. Mass differences are indicated within the spectra.

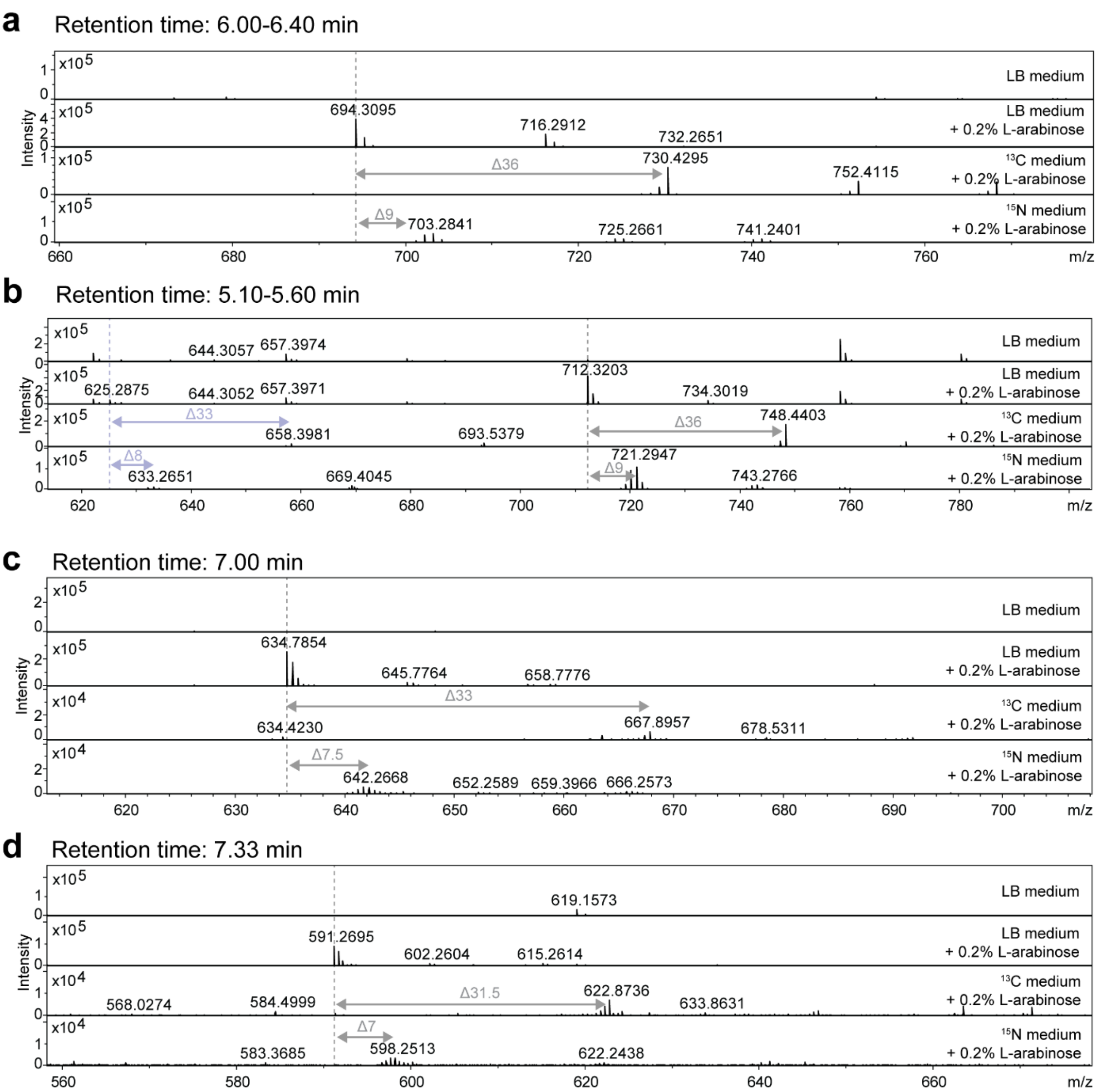

**Figure S53.** Isotope medium experiments for structure determination of compounds **7** to **11**. HPLC-MS analysis of *P. temperata* K122/JL14. Determination of carbon and nitrogen atoms in compounds **7** (*m/z* $[M+H]^+$ = 694.3096) (**a**), **8** (*m/z* $[M+H]^+$ = 712.3214) (**b**), **9** (*m/z* $[M+H]^+$ = 625.2897) (**b**), **10** (*m/z* $[M+2H]^{2+}$ = 634.7875) (**c**) and **11** (*m/z* $[M+2H]^{2+}$ = 591.2711) (**d**) by isotope labelling experiments. MS spectra are displayed for the strain cultivated in low salt LB medium ($^{12}C$, $^{14}N$), fully labelled $^{13}C$ medium ($^{13}C$, $^{14}N$) and $^{15}N$ medium ($^{12}C$, $^{15}N$) with indication of the retention time and mass shifts.

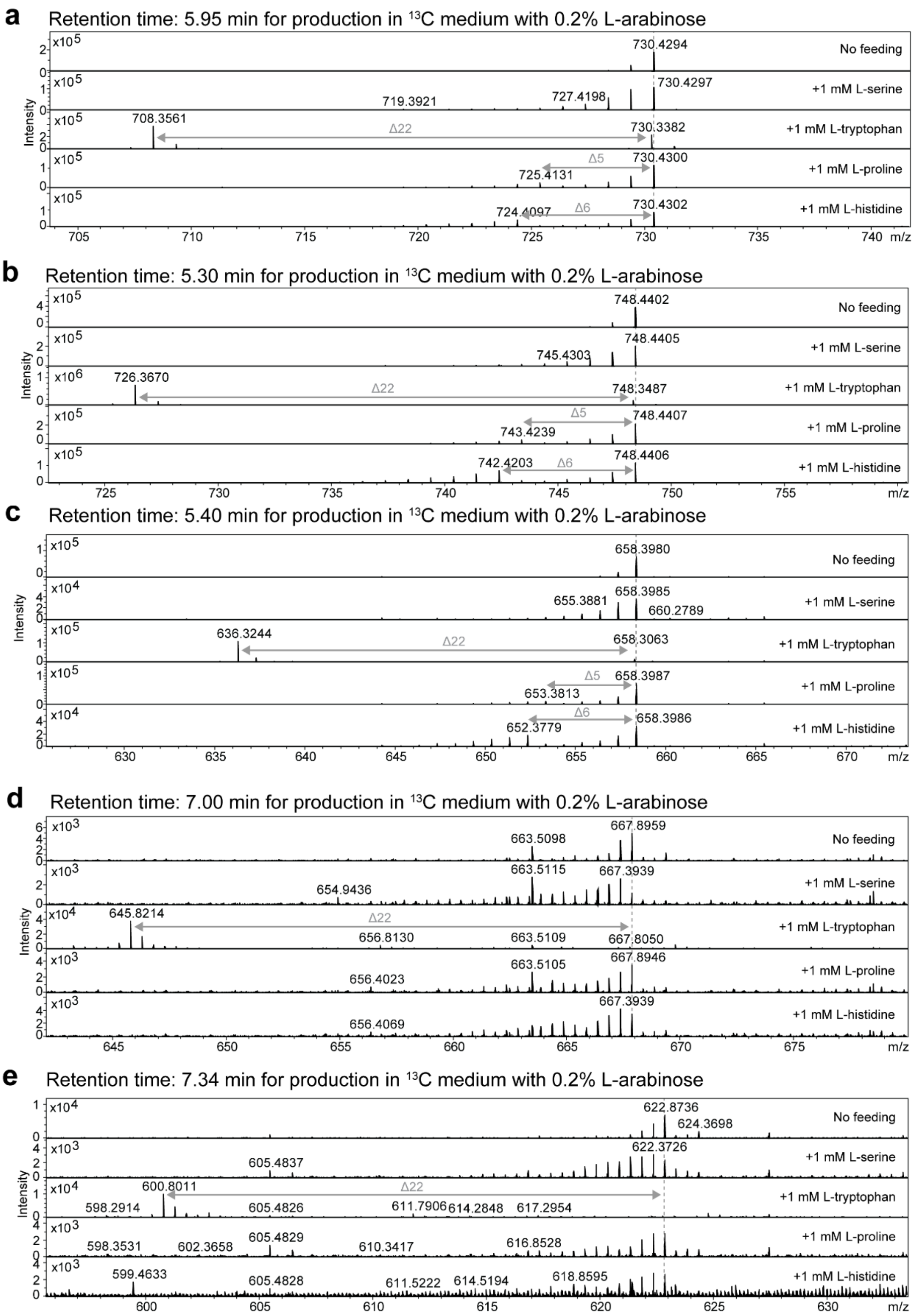

**Figure S54.** Inverse isotope feeding experiments for structure determination of compounds **7** to **11**. HPLC-MS analysis of *P. temperata* K122/JL14. Inverse feeding experiment performed with fully labelled $^{13}C$ medium ($^{13}C$, $^{14}N$) supplemented with unlabelled L-serine, L-tryptophan, L-proline and L-histidine. MS spectra for indicated retention times are shown with indications of mass shifts for the respective compounds **7** (**a**), **8** (**b**), **9** (**c**), **10** (**d**) and **11** (**e**). Inverse feeding with L-serine showed for all compounds no conclusive incorporation. For compounds **10** and **11** only incorporation of tryptophan could be verified.

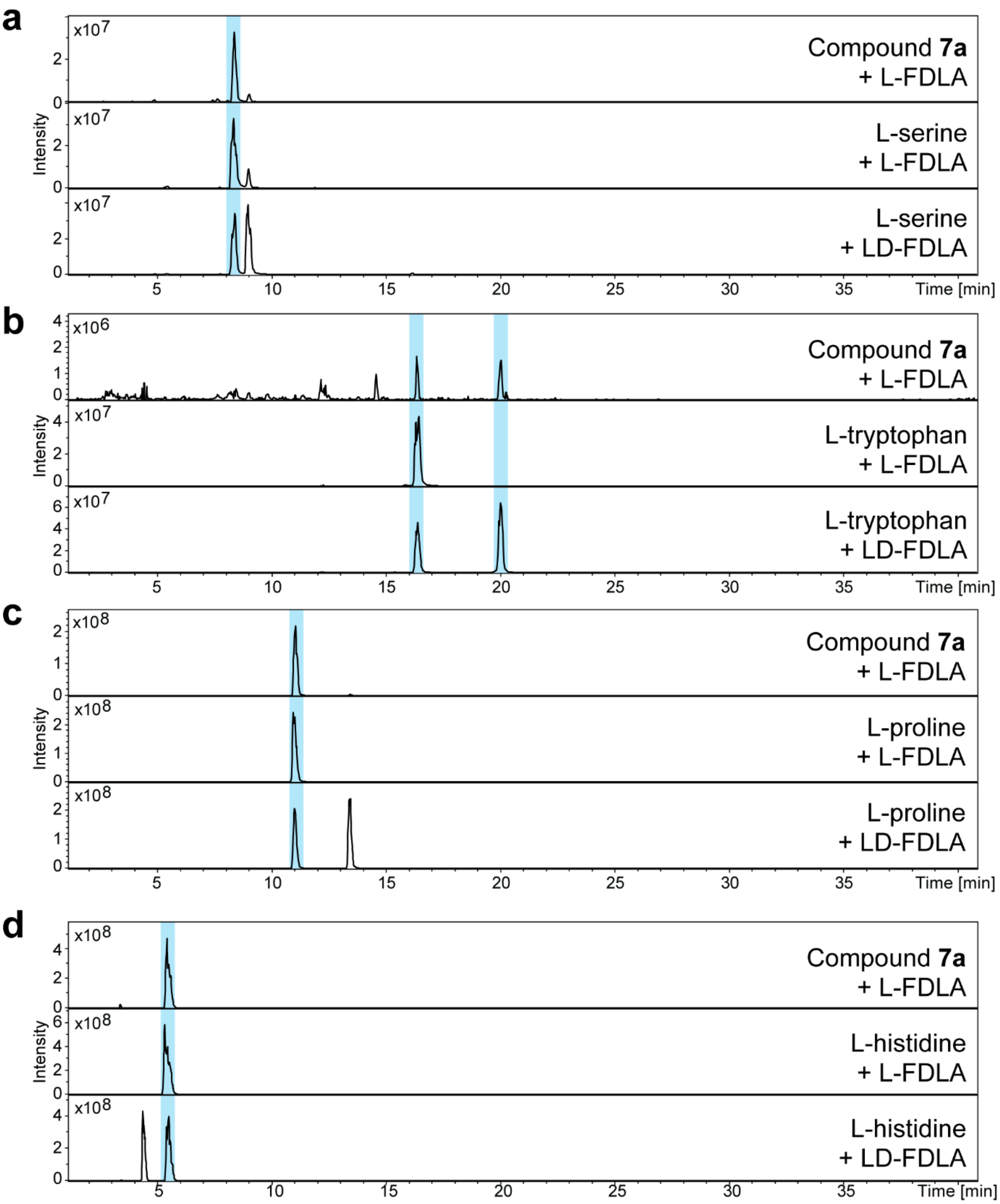

**Figure S55**. Configuration determination of amino acids in **7a** using the advanced Marfey's method. HPLC-MS analysis of hydrolyzed and L-FDLA derivatized **7a** along with derivatized amino acid standards with L-FDLA and LD-FDLA. Depicted are EIC traces for Marfey's derivatives of L-serine (Ser, *m/z* 400.14 $[M+H]^+$)(**a**), L-tryptophan (Trp *m/z* 499.19 $[M+H]^+$)(**b**), L-proline (Pro, *m/z* 410.16 [M+H]+)(**c**) and L-histidine (His, *m/z* 450.17 $[M+H]^+$)(**d**).

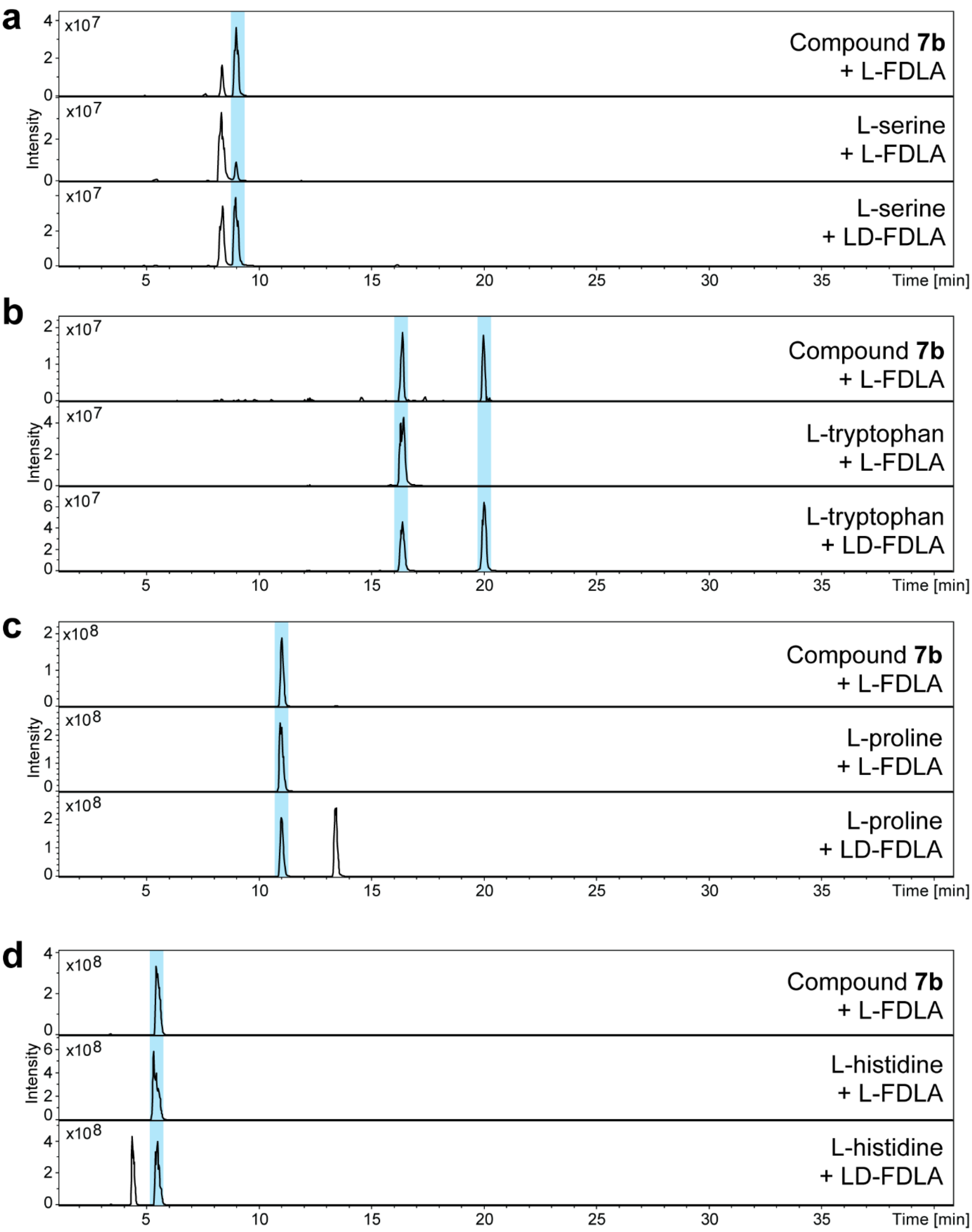

**Figure S56**. Configuration determination of amino acids in **7b** using the advanced Marfey's method. HPLC-MS analysis of hydrolyzed and L-FDLA derivatized **7b** along with derivatized amino acid standards with L-FDLA and LD-FDLA. Depicted are EIC traces for Marfey’s derivatives of L-serine (Ser, *m/z* 400.14 $[M+H]^+$)(**a**), L-tryptophan (Trp *m/z* 499.19 $[M+H]^+$)(**b**), L-proline (Pro, *m/z* 410.16 [M+H]+)(**c**) and L-histidine (His, *m/z* 450.17 $[M+H]^+$)(**d**).

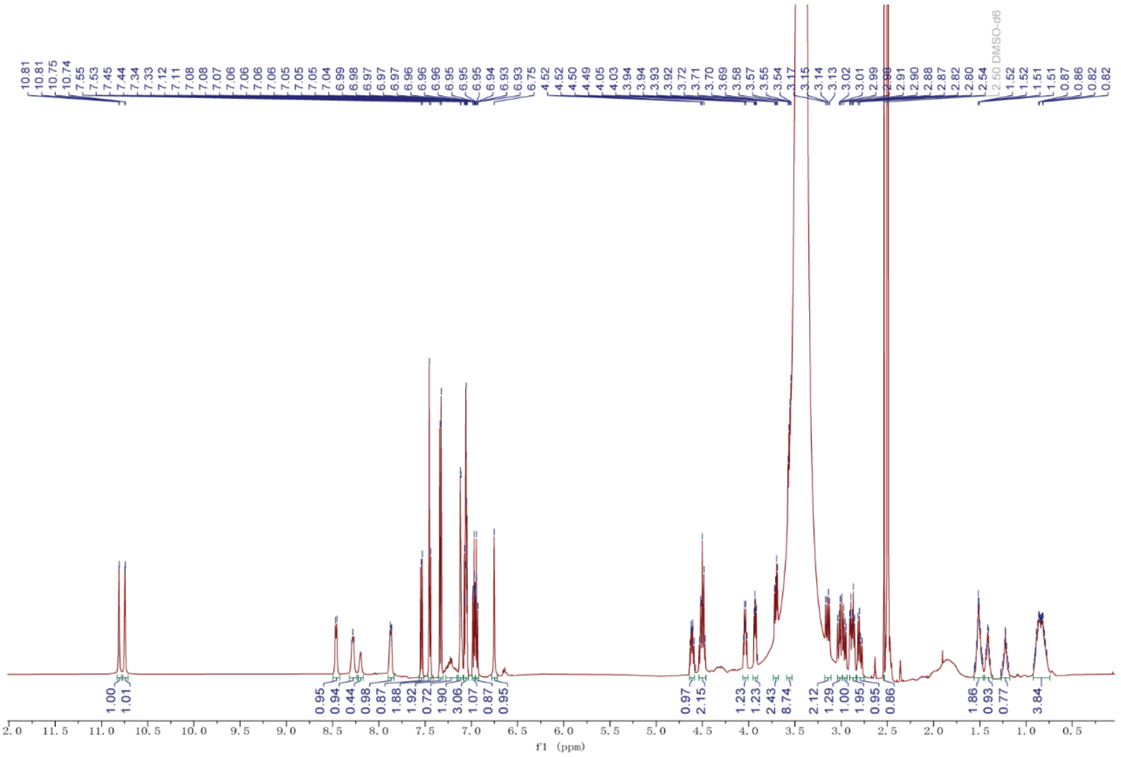

**Figure S57.** $^{1}H$ NMR spectrum (DMSO-$d_6$, 500 MHz) of **7a**.

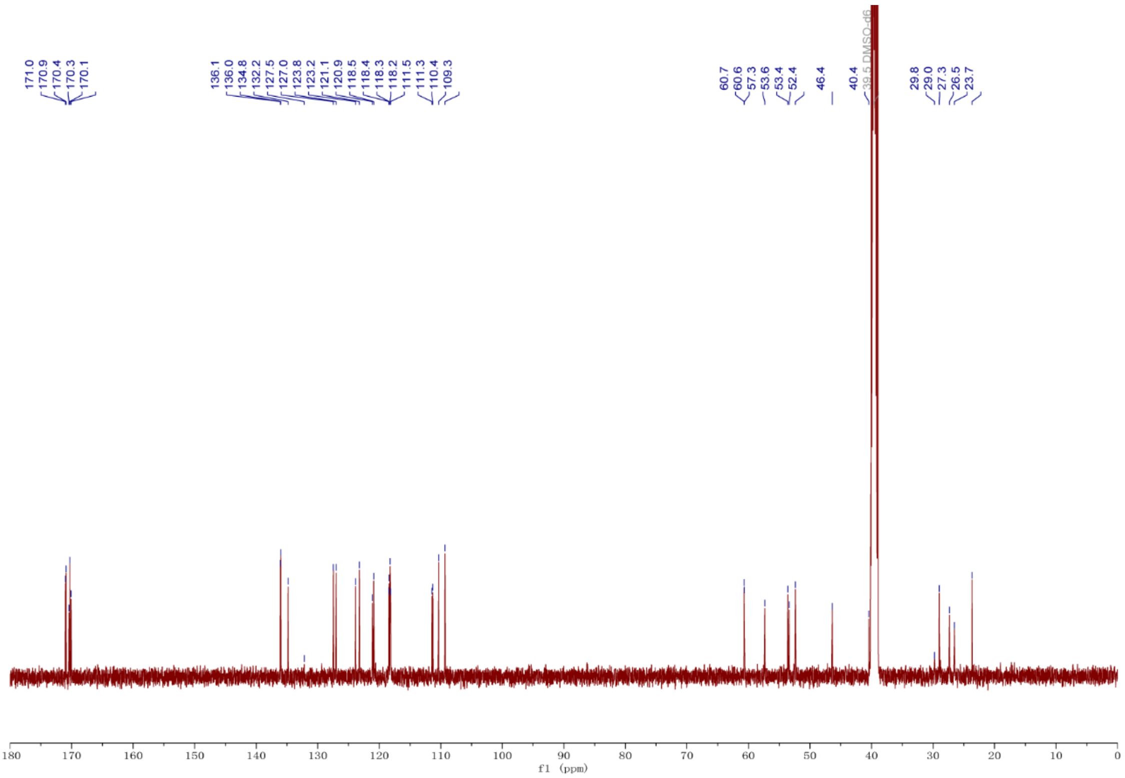

**Figure S58.** $^{13}C$ NMR spectrum (DMSO-$d_6$, 125 MHz) of **7a**.

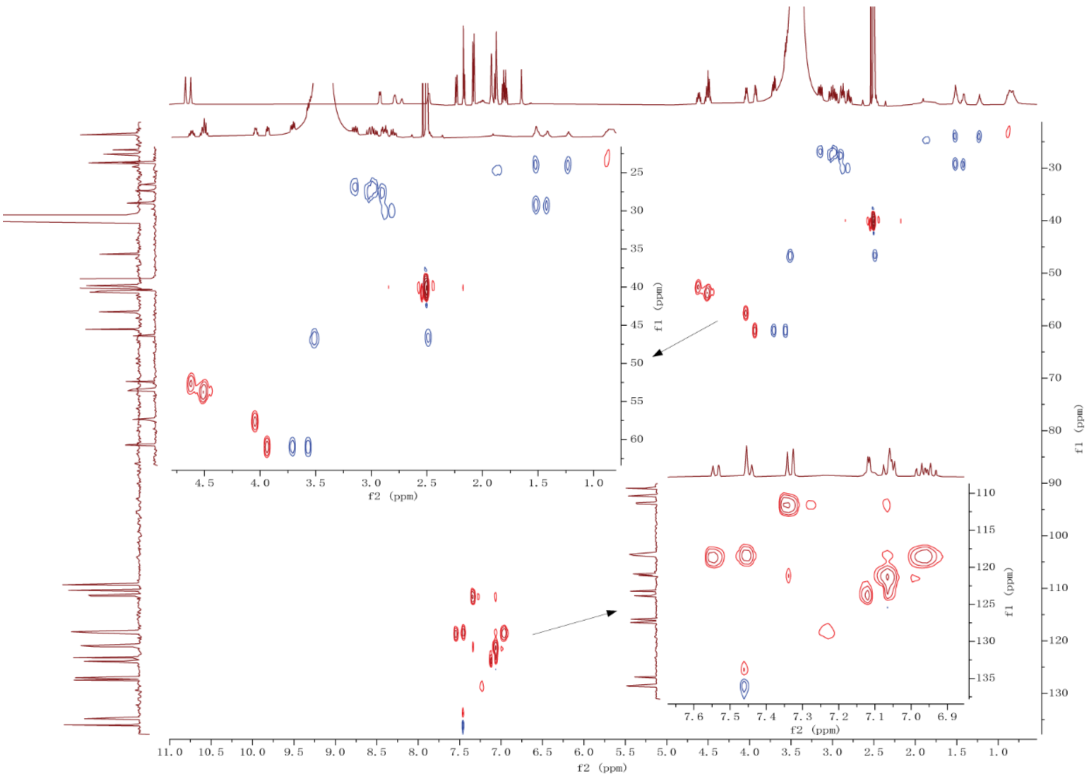

**Figure S59.** HSQC NMR spectrum (DMSO-$d_6$, 500 MHz) of **7a**.

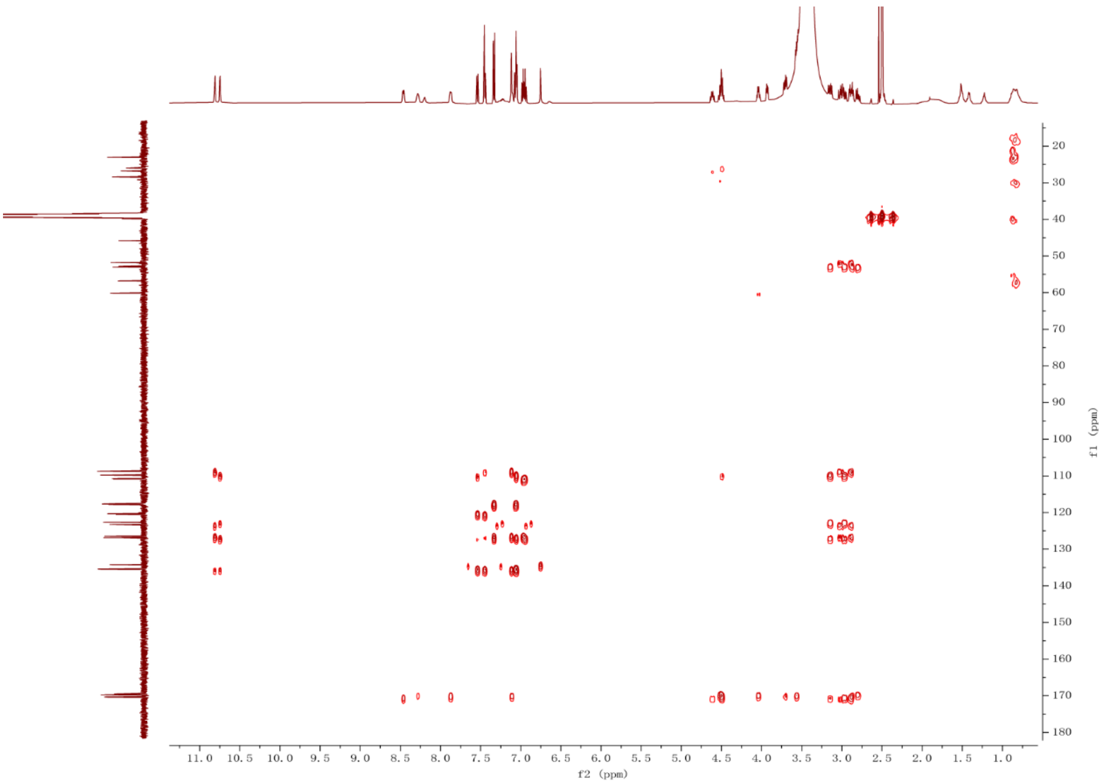

**Figure S60.** HMBC spectrum (DMSO-$d_6$, 500 MHz) of **7a**.

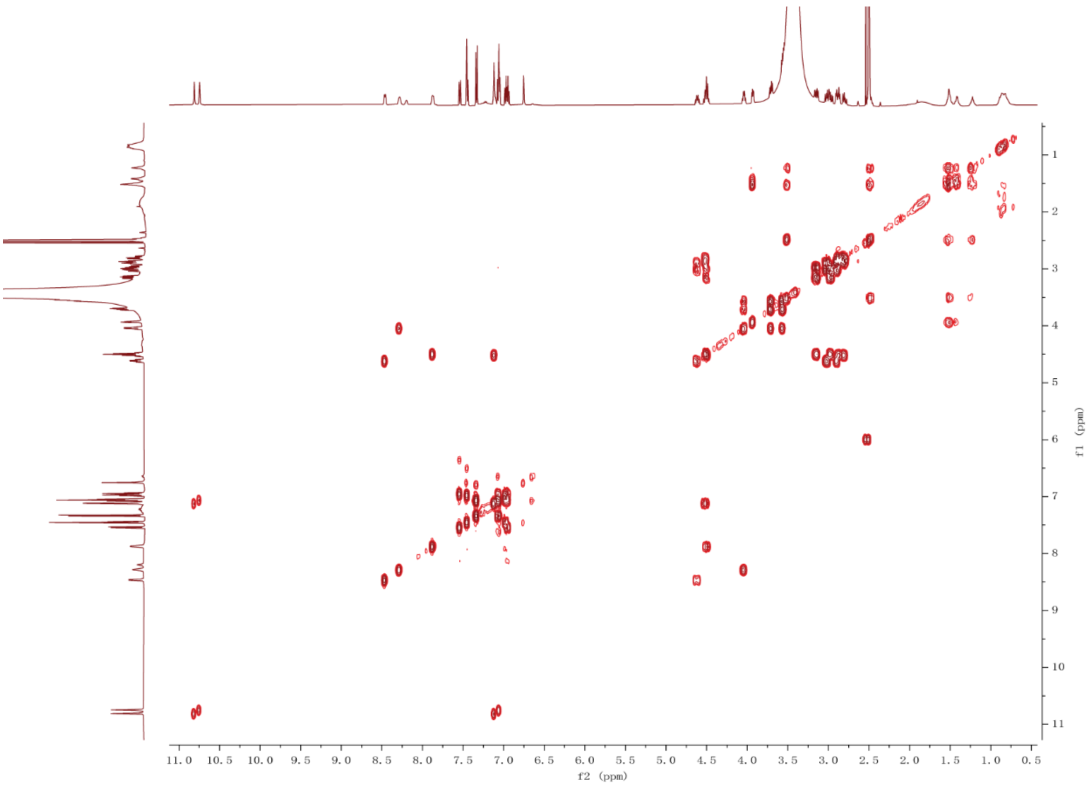

**Figure S61.** $^{1}H$-$^{1}H$ COSY spectrum (DMSO-$d_6$, 500 MHz) of **7a**.

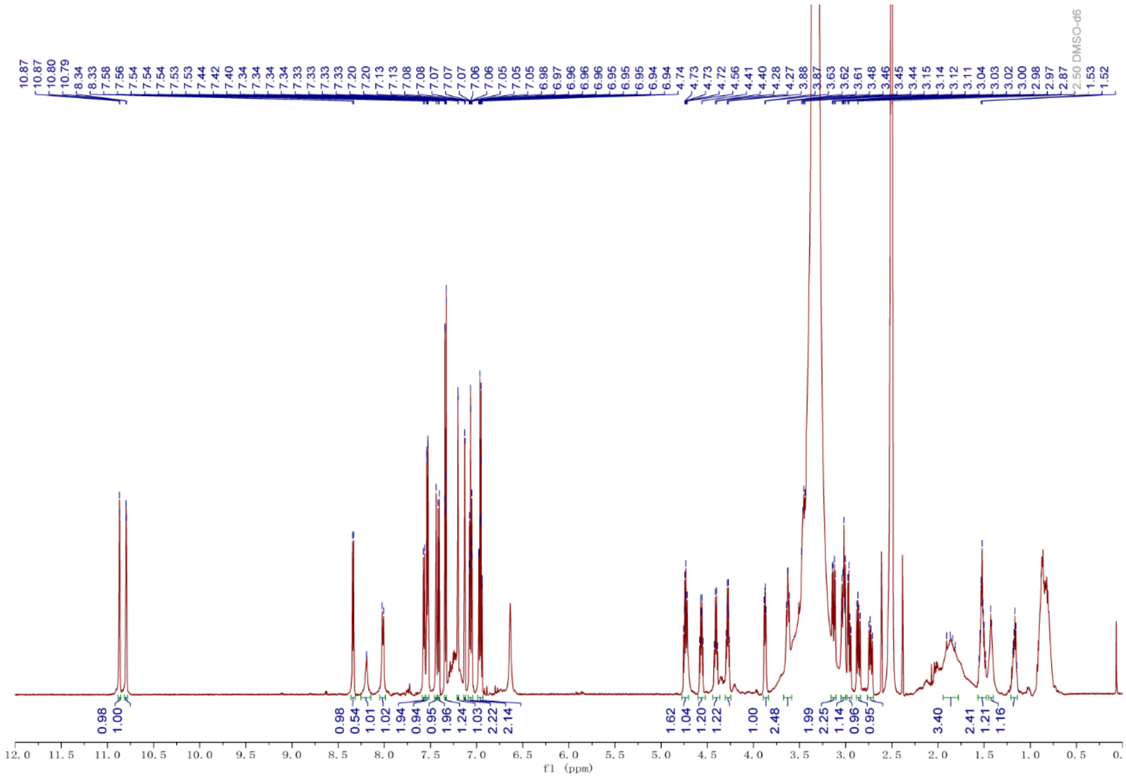

**Figure S62.** $^{1}H$ NMR spectrum (DMSO-$d_6$, 500 MHz) of **7b**.

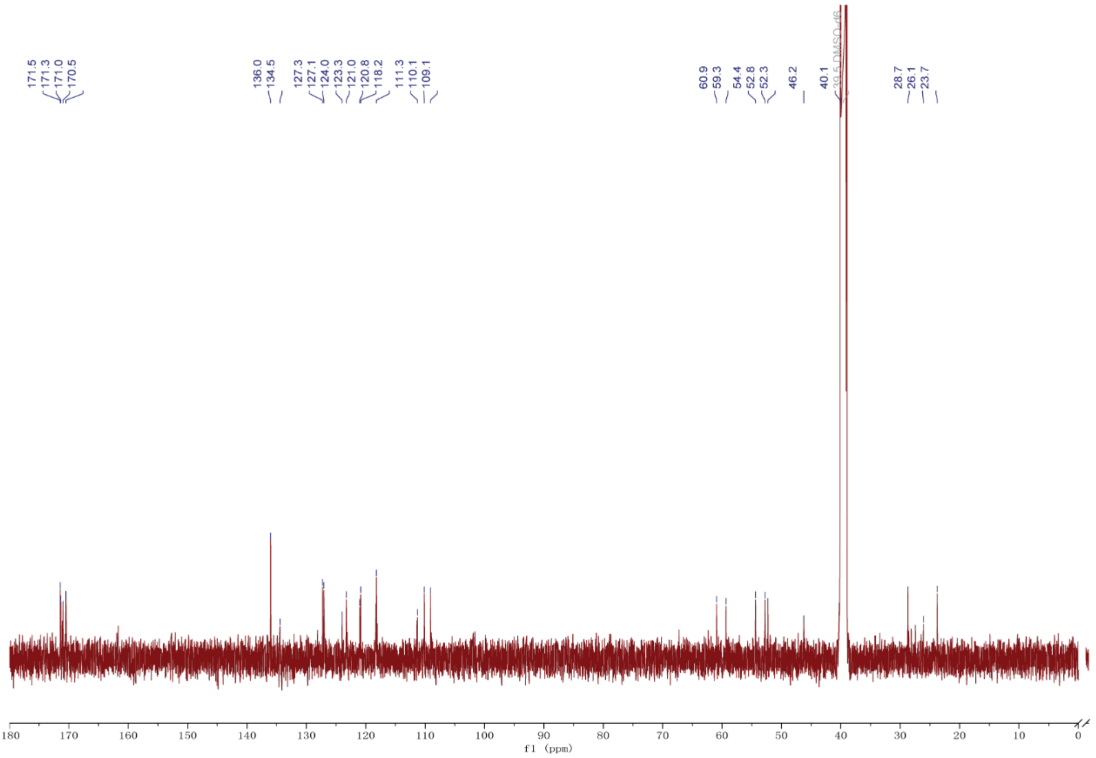

**Figure S63.** $^{13}C$ NMR spectrum (DMSO-$d_6$, 125 MHz) of **7b**.

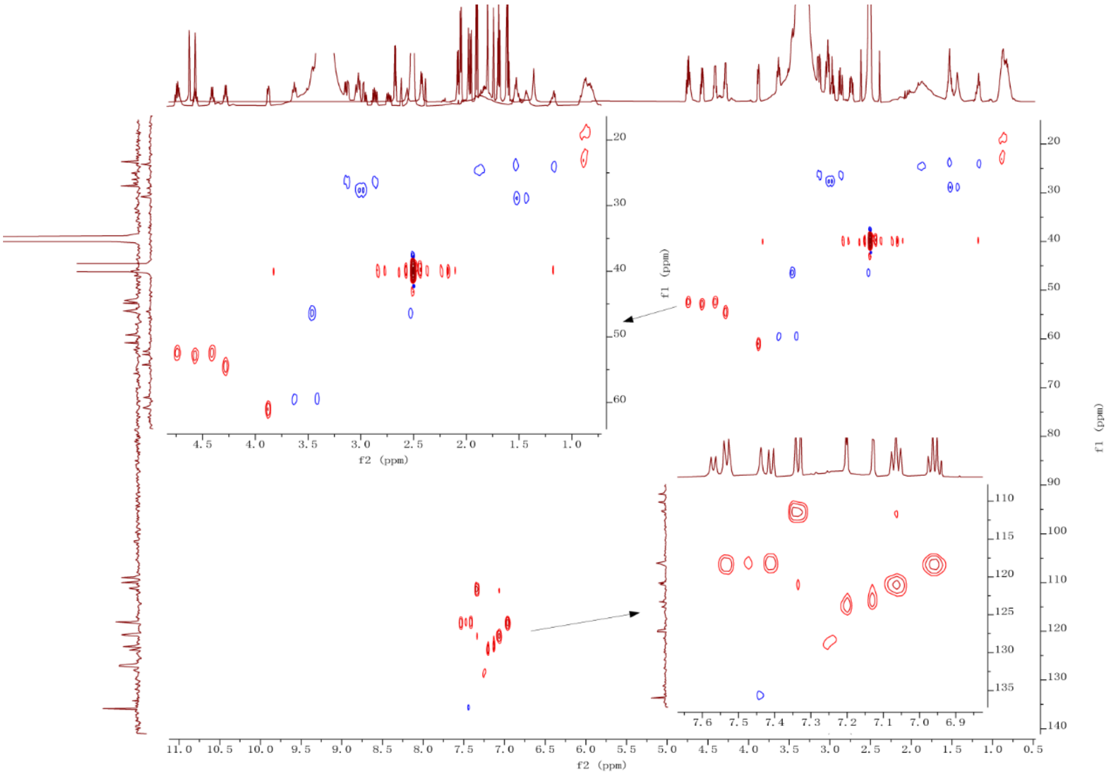

**Figure S64.** HSQC NMR spectrum (DMSO-$d_6$, 500 MHz) of **7b**.

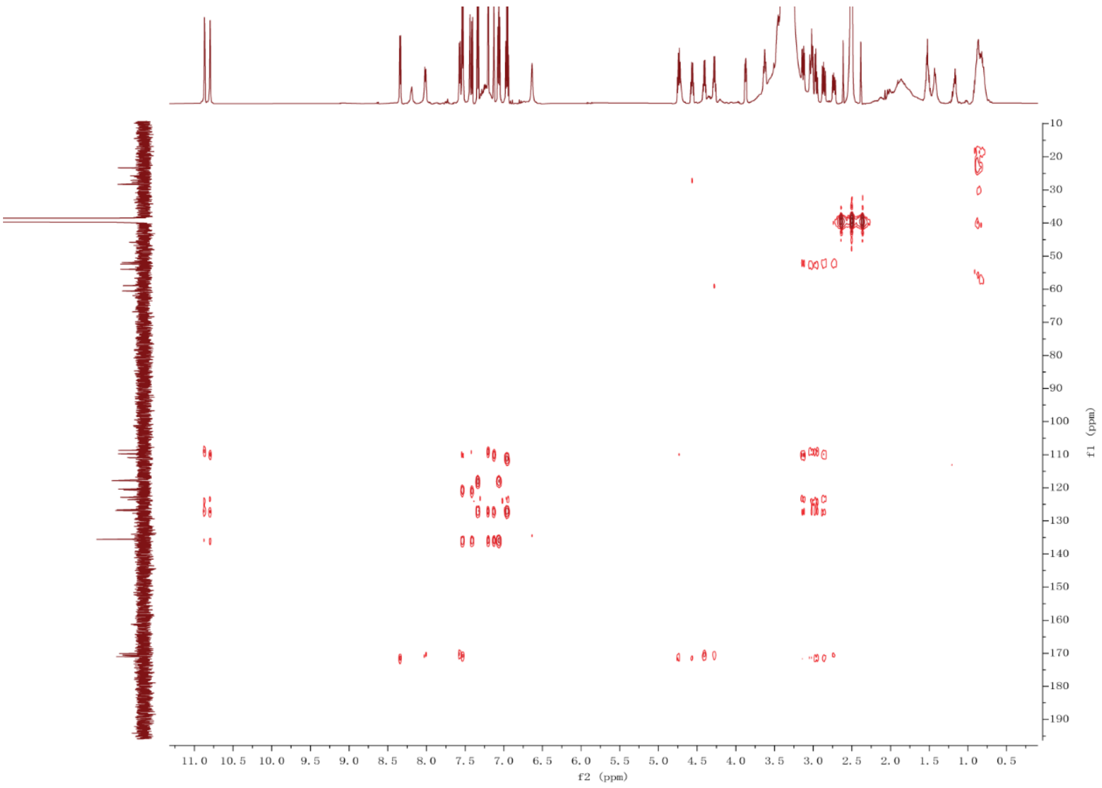

**Figure S65.** HMBC spectrum (DMSO-d$_6$, 500 MHz) of **7b**.

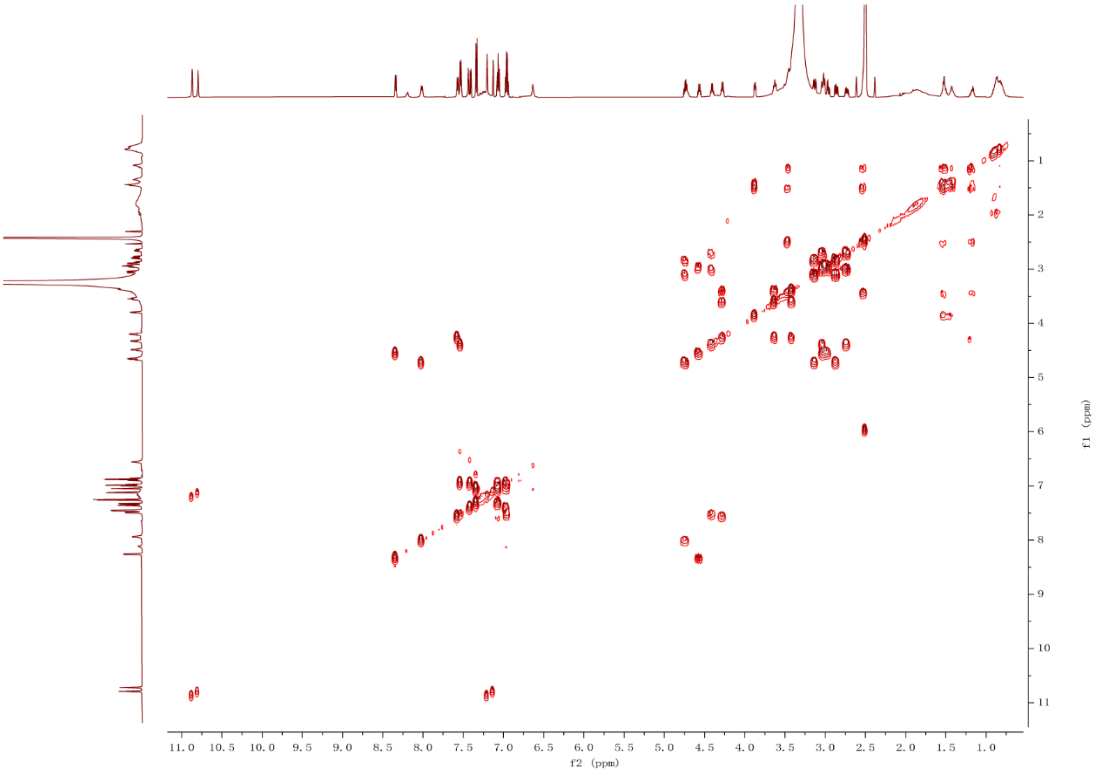

**Figure S66.** $^1H$-$^1H$ COSY spectrum (DMSO-d$_6$, 500 MHz) of **7b**.

a

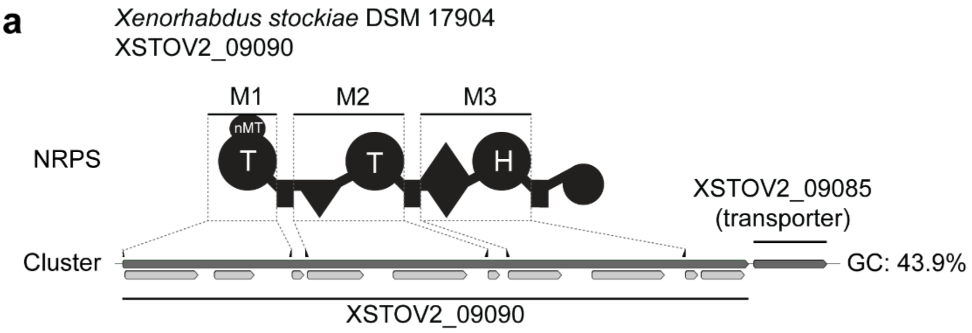

b

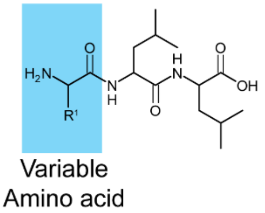

| Module | Detected mass [M+H]+ | Δppm | Sum formula | Estimated Amino Acid |
|---|---|---|---|---|
| M1 | 346.2331 | 1.4 | $C_{16}H_{32}N_3O_5$ | Thr |
| M1 | 360.2492 | 0.4 | $C_{17}H_{34}N_3O_5$ | $Thr^{Nme}$ |
| M2 | 346.2333 | 1.0 | $C_{16}H_{32}N_3O_5$ | Thr |
| M3 | 382.2449 | -0.1 | $C_{18}H_{32}N_5O_4$ | His |

c

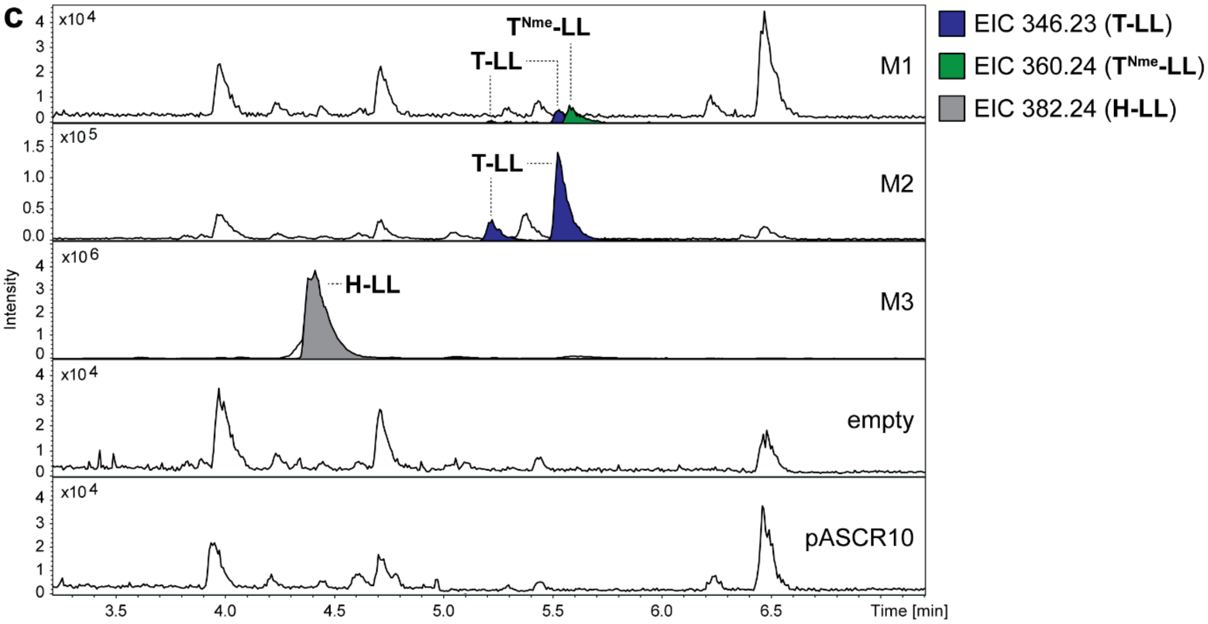

**Figure S67.** HPLC-MS/MS data referred to module screening of XSTOV2_09090 from *X. stockiae*. (**a**) Illustration of the gene cluster XSTOV2_09090 from *X. stockiae* with schematic illustration of the encoded NRPS and its adjacent encoded transporter XSTOV2_09085. Indication of the region of cloned modules M1 to M3 on gene and protein level. Utilized primers are shown above the cluster. A domain specitifity is indicated by the one-letter amino acid code. GC content is depicted next to the cluster. (**b**) Table of HPLC-MS/MS data indicating the targeted module cloned into pASCR8, the detected mass of the compound, the corresponding error, the sum formula and the incoporated amino acid introduced to the peptide at the highlighted position in the chemical structure (left). The amino acid is shown by the three-letter amino acid code ($T^{Nme}$ = N-methylated threonine). (**c**) HPLC-MS/MS data of *E. coli* DH10B::*mtaA* expressing M1 to M3 cloned into pASCR8. Expression of an empty vector and pASCR10 served as negative controls. Base Peak Chromatogram (BPC, top) with Extracted Ion Chromatogram (EIC, below with colors according to the depicted legend) of **T-LL** (*m/z* $[M+H]^+$ = 346.233), $\mathbf{T^{Nme}}$**-LL** (*m/z* $[M+H]^+$ = 60.249) and **H-LL** (*m/z* $[M+H]^+$ = 382.244). Indication of the targeted modules on the right of the chromatograms.

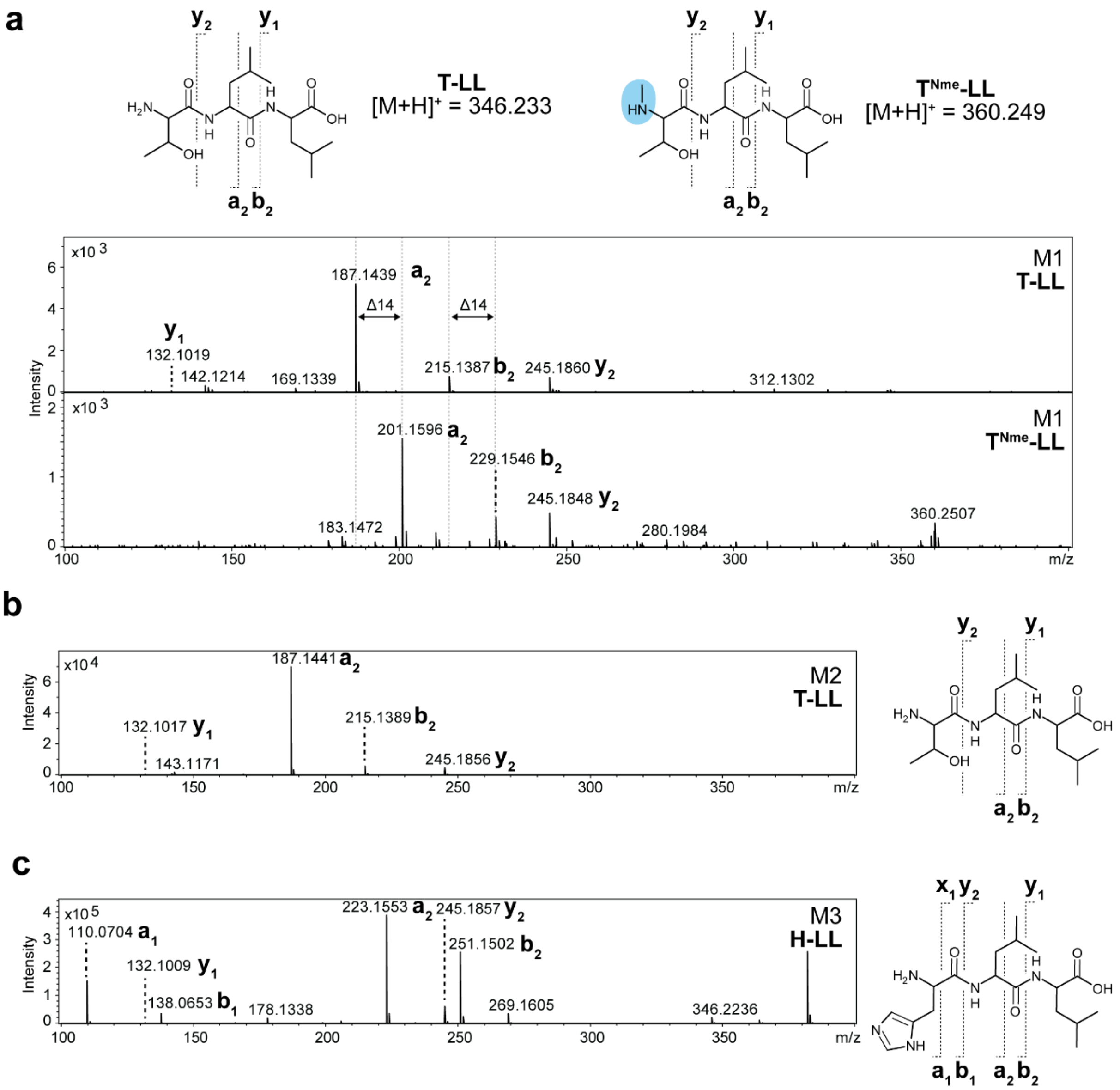

**Figure S68.** $MS^2$ spectra of HPLC-MS/MS data referred to figure S67. (**a-c**) The respective compounds are displayed, along with indications of the assigned fragments. The targeted module and peptide name is depicted within the spectra. (**a**) Comparison of the $MS^2$ spectra of **T-LL** and **T$^{Nme}$-LL**.

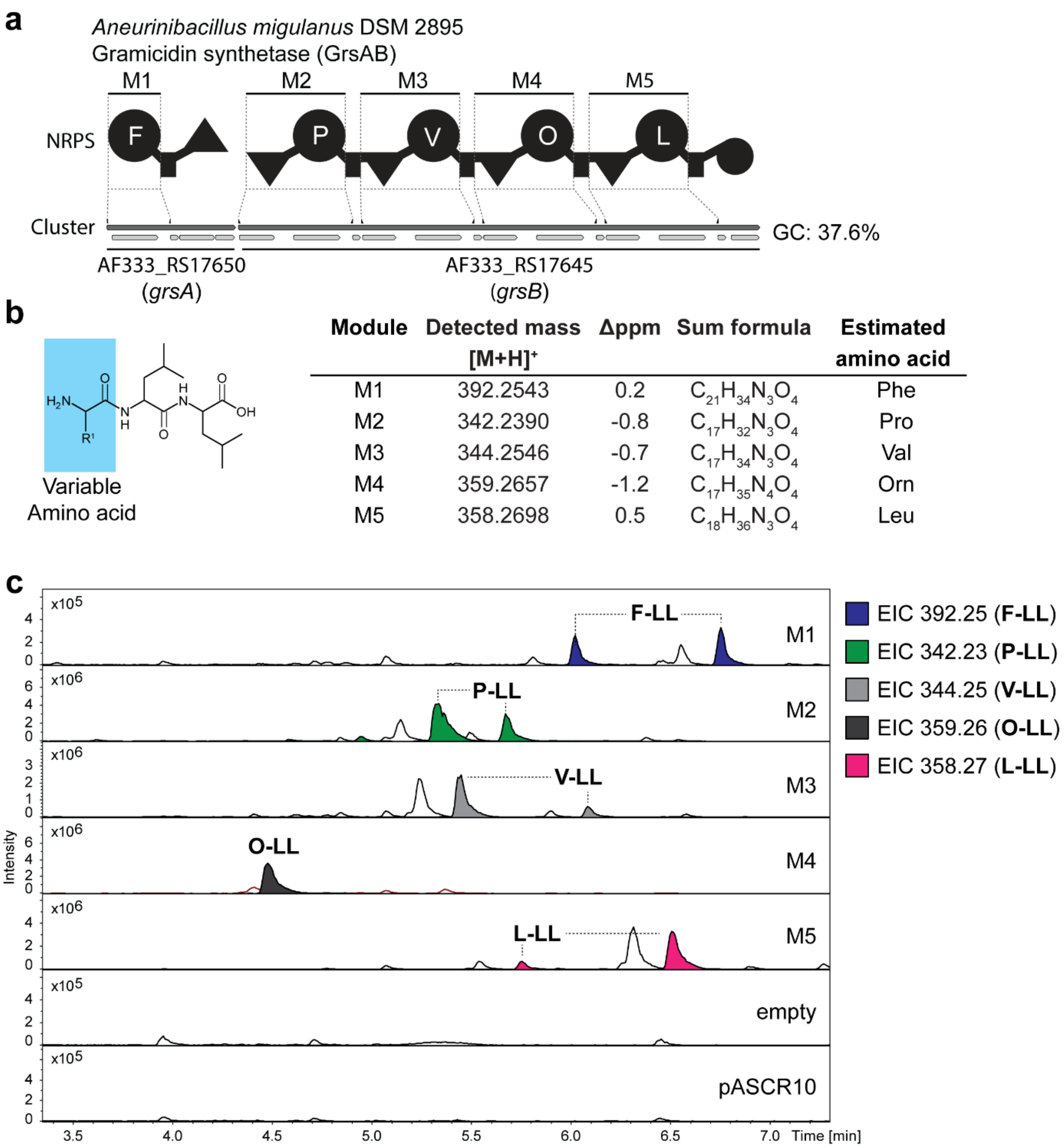

| Module | Detected mass [M+H]+ | Δppm | Sum formula | Estimated amino acid |
|---|---|---|---|---|
| M1 | 392.2543 | 0.2 | $C_{21}H_{34}N_3O_4$ | Phe |
| M2 | 342.2390 | -0.8 | $C_{17}H_{32}N_3O_4$ | Pro |
| M3 | 344.2546 | -0.7 | $C_{17}H_{34}N_3O_4$ | Val |
| M4 | 359.2657 | -1.2 | $C_{17}H_{35}N_4O_4$ | Orn |
| M5 | 358.2698 | 0.5 | $C_{18}H_{36}N_3O_4$ | Leu |

**Figure 69.** HPLC-MS/MS data referred to module screening of gramicidin synthetase from A. *migulanus*. (**a**) Illustration of the gene cluster *grsAB* from *A. migulanus* with schematic illustration of the encoded NRPS. Indication of the region of cloned modules M1 to M5 on gene and protein level. Utilized primers are shown above the cluster. A domain specitifity is indicated by the one-letter amino acid code (O = ornithine). GC content is depicted next to the cluster. (**b**) Table of HPLC-MS/MS data indicating the targeted module cloned into pASCR8, the detected mass of the compound, the corresponding error, the sum formula and the incoporated amino acid introduced to the peptide at the highlighted position in the chemical structure (left). The amino acid is shown by the three-letter amino acid code (O = ornithine). (**c**) HPLC-MS/MS data of *E. coli* DH10B::*mtaA* expressing M1 to M5 cloned into pASCR8. Expression of an empty vector and pASCR10 served as negative control. Base Peak Chromatogram (BPC, top) with Extracted Ion Chromatogram (EIC, below with colors according to the depicted legend) of **F-LL** (*m/z* $[M+H]^+$ = 392.254), **P-LL** (*m/z* $[M+H]^+$ = 342.254), **V-LL** (*m/z* $[M+H]^+$ = 344.254), **O-LL** (*m/z* $[M+H]^+$ = 359.265) and **L-LL** (*m/z* $[M+H]^+$ = 358.270). Indication of the targeted modules on the right of the chromatograms.

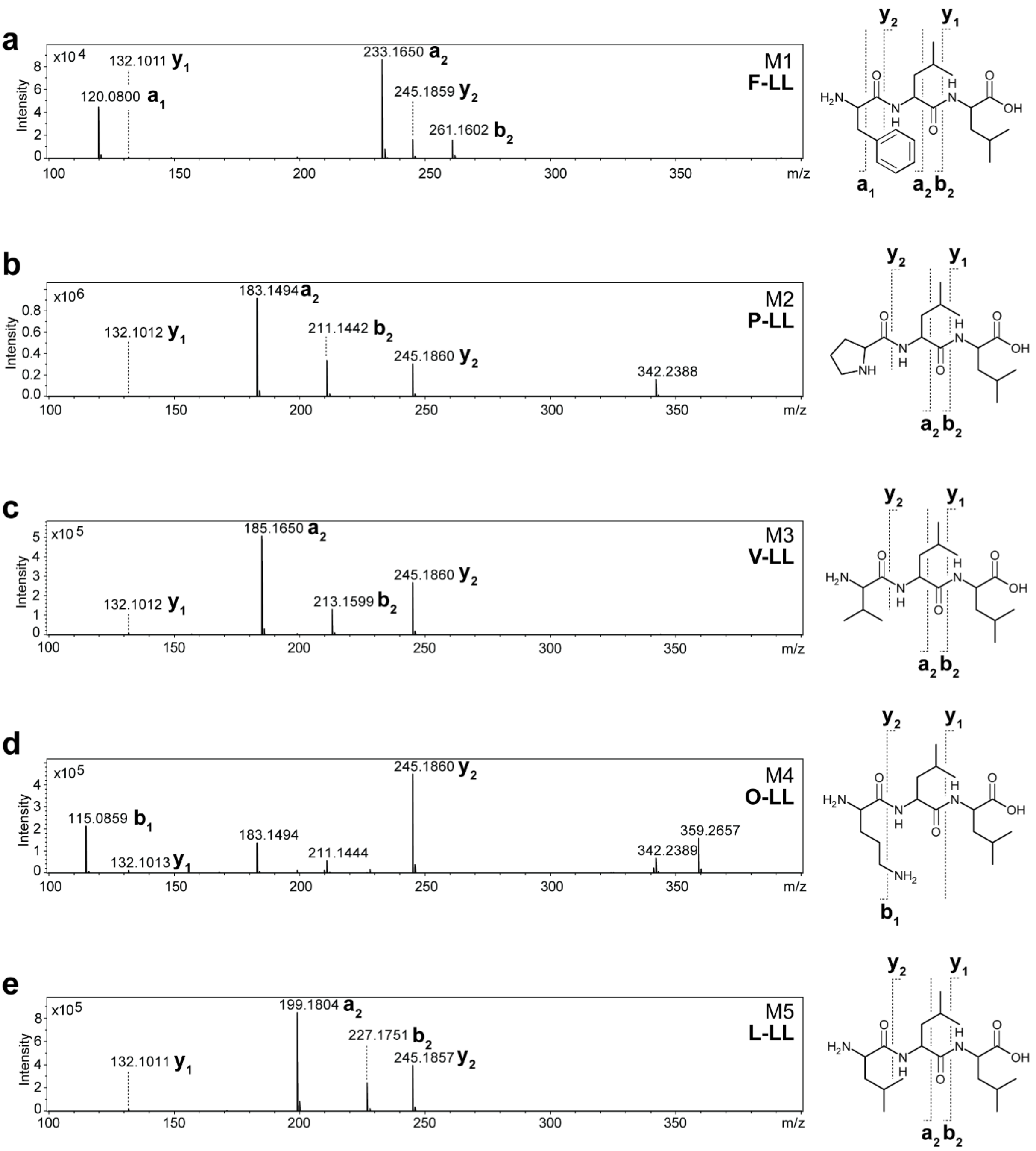

**Figure S70.** $MS^2$ spectra of HPLC-MS/MS data referred to figure S69. (**a-e**) The respective compounds are displayed, along with indications of the assigned fragments. The targeted module and peptide name is depicted within the spectra.

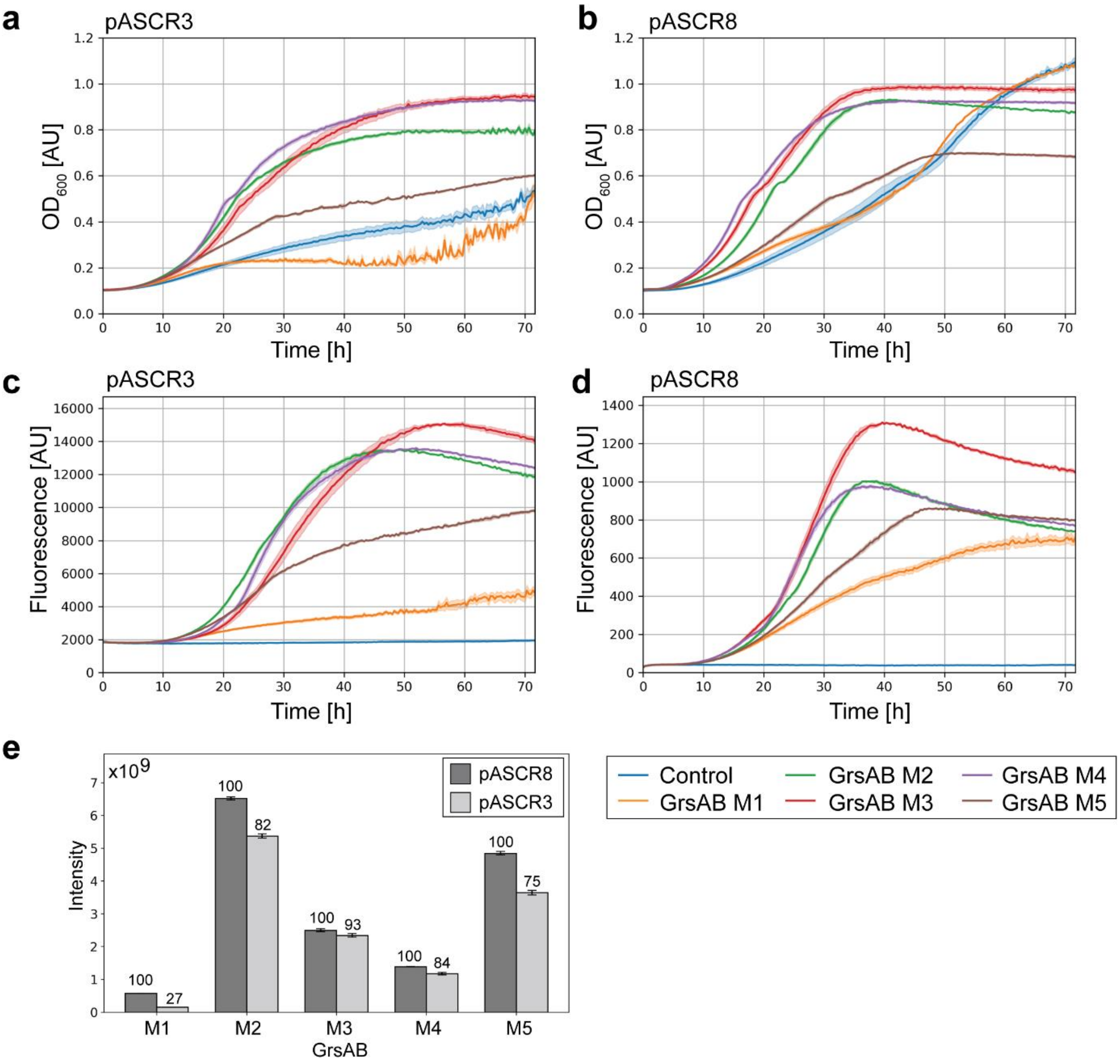

**Figure S71.** Comparison of expression, growth and peptide production for cloning vectors pASCR3 and pASCR8. The genes of the individual five modules of GrsAB were cloned onto pASCR3 and pASCR8 and were expressed in *E. coli* DH10B::*mtaA*. Expression of pASCR6 and pASCR11 served as controls for pASCR3 and pASCR8, respectively. The $OD_{600}$ with SEM was determined for 72 hours for cloning vectors pASCR3 (**a**) and pASCR8 (**b**). msfGFP fluorescence (Ex.: 480 nm; Em.: 525 nm, pASCR3) and mScarlett-I3 (Ex.: 560 nm; Em.: 605 nm, pASCR8) was determined for 72 hours for cloning vectors pASCR3 (**c**) or pASCR8 (**d**), respectively. The legend of the individual cloned modules is shown in the bottom right. (**e**) Barplot of the average of the mean with SEM of the determined peak areas of the respective EICs of **F-LL** (*m/z* $[M+H]^+$ = 392.254, GrsAB M1), **P-LL** (*m/z* $[M+H]^+$ = 342.254, GrsAB M2), **V-LL** (*m/z* $[M+H]^+$ = 344.254, GrsAB M3), **O-LL** (*m/z* $[M+H]^+$ = 359.265, GrsAB M4) and **L-LL** (*m/z* $[M+H]^+$ = 358.270, GrsAB M5). The numbers above the bars indicate the relative percentage production of a given module of GrsAB compared between the two cloning vector systems.

**a**

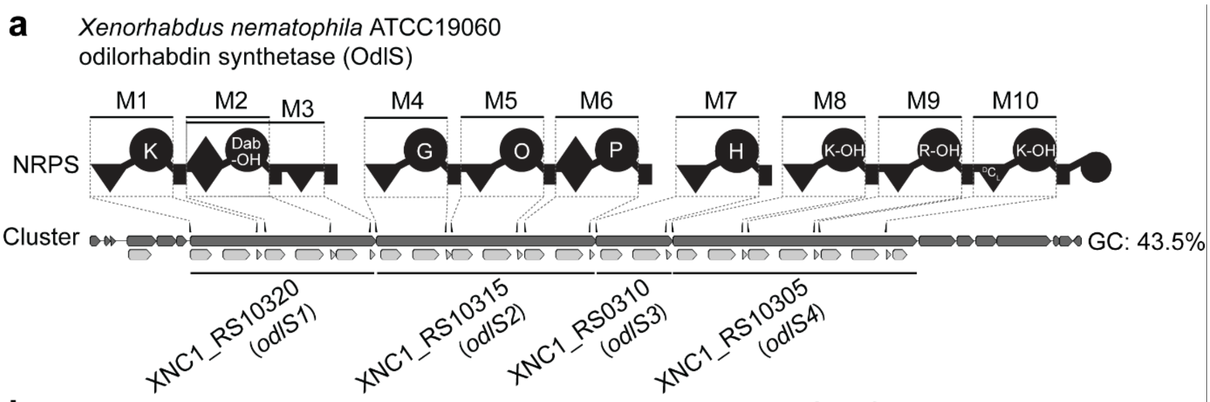

**b**

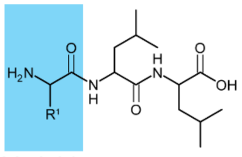

| Module | Detected mass $[M+H]^+$ | Δppm | Sum formula | Estimated amino acid |
|---|---|---|---|---|
| M1 | 373.2807 | 0.6 | $C_{18}H_{37}N_4O_4$ | Lys |
| M2 | 345.2489 | 2.0 | $C_{16}H_{33}N_4O_4$ | Dab |
| M4 | 302.2073 | 0.4 | $C_{14}H_{28}N_3O_4$ | Gly |
| M5 | 359.2657 | -1.3 | $C_{17}H_{35}N_4O_4$ | Orn |
| M6 | 342.2387 | 0.0 | $C_{17}H_{32}N_3O_4$ | Pro |
| M7 | 382.2453 | -1.1 | $C_{18}H_{32}N_5O_4$ | His |
| M8 | 373.2811 | -0.4 | $C_{18}H_{37}N_4O_4$ | Lys |
| M9 | 401.2867 | 0.8 | $C_{18}H_{37}N_6O_4$ | Arg |
| M10 | 373.2814 | -1.1 | $C_{18}H_{37}N_4O_4$ | Lys |

**c**

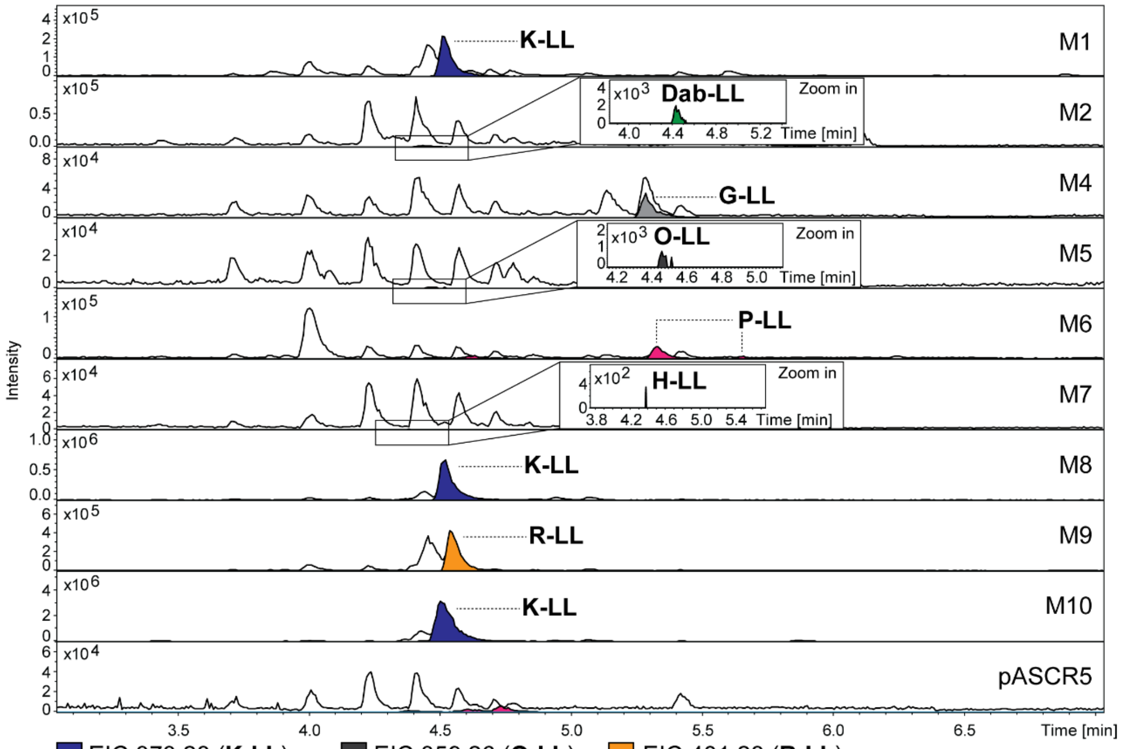

**Figure S72.** HPLC-MS/MS data referred to module screening of odilorhabdin synthetase (OdlS) from *X. nematophila*. (**a**) Illustration of the gene cluster of odilorhabdin synthetase (OdlS) from *X. nematophila* with schematic illustration of the encoded NRPS. Indication of the region of cloned modules M1 to M10 on gene and protein level. Utilized primers are shown above the cluster. A domain specitifity is indicated by the one-letter amino acid code (Dab-OH = β-hydroxy diamino butyric acid, O = ornithine, K-OH = δ-hydroxylysine, R-OH = β-hydroxy-arginine). GC content is depicted next to the cluster. (**b**) Table of HPLC-MS/MS data indicating the targeted module cloned into pASCR3, the detected mass of the compound, the corresponding error, the sum formula and the incoporated amino acid introduced to the peptide at the highlighted position in the chemical structure (left). The amino acid is shown by the three-letter amino acid code (Dab = diamino butyric acid, Orn = ornithine). (**c**) HPLC-MS/MS data of *E. coli* DH10B::*mtaA* expressing M1 to M10 cloned into pASCR3. Expression of pASCR5 served as negative control. Base Peak Chromatogram (BPC, top) with Extracted Ion Chromatogram (EIC, below with colors according to the depicted legend) of **K-LL** (*m/z* $[M+H]^+$ = 373.280), **Dab-LL** (*m/z* $[M+H]^+$ = 345.249), **G-LL** (*m/z* $[M+H]^+$ = 302.207), **O-LL** (*m/z* $[M+H]^+$ = 359.265), **P-LL** (*m/z* $[M+H]^+$ = 342.238), **H-LL** (*m/z* $[M+H]^+$ = 382.244) and **R-LL** (*m/z* $[M+H]^+$ = 401.287). Indication of the targeted modules on the right of the chromatograms.

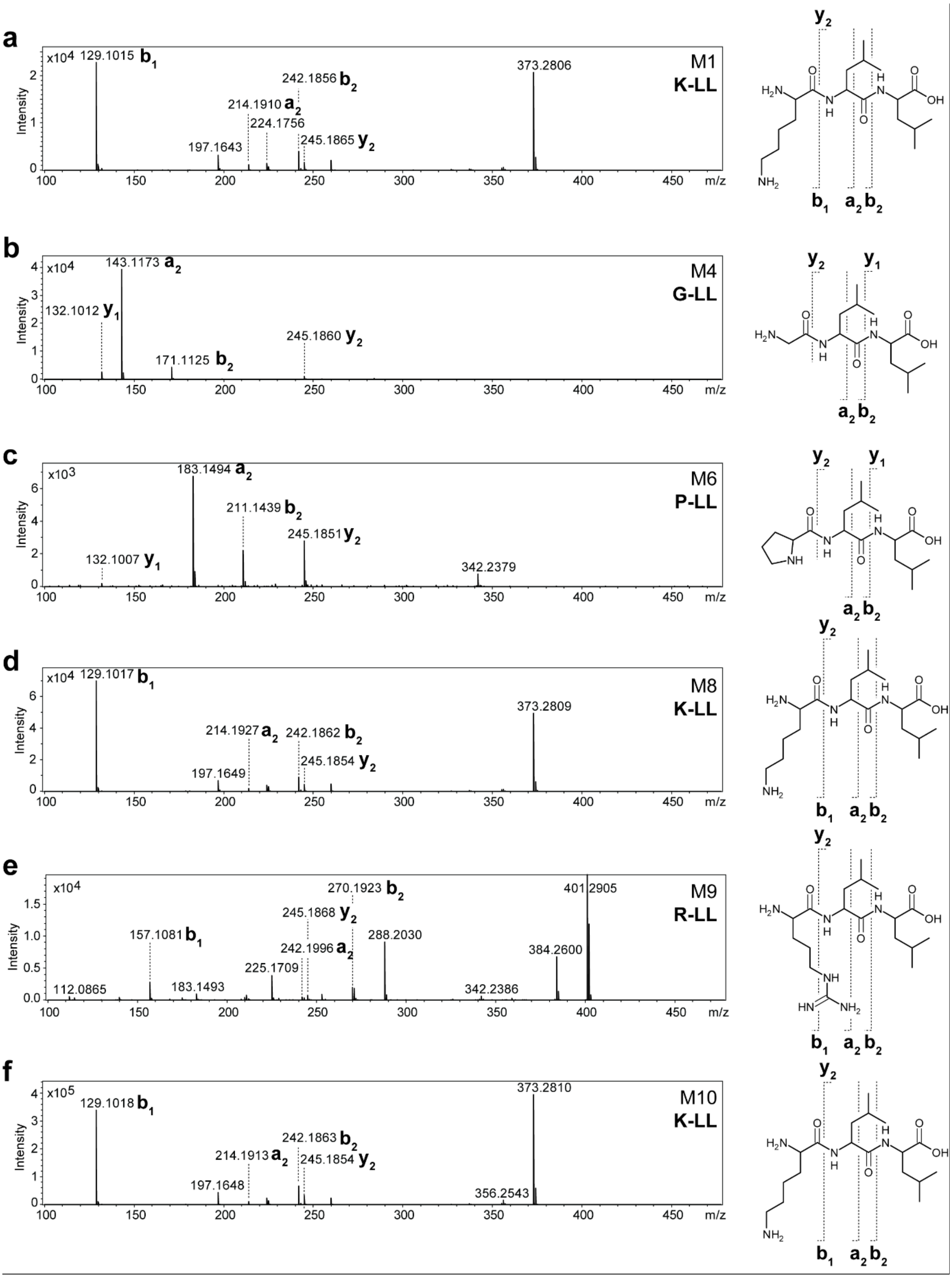

**Figure S73.** $MS^2$ spectra of HPLC-MS/MS data referred to figure S72. (**a-f**) The respective compounds are displayed, along with indications of the assigned fragments. The targeted module and peptide name is depicted within the spectra.

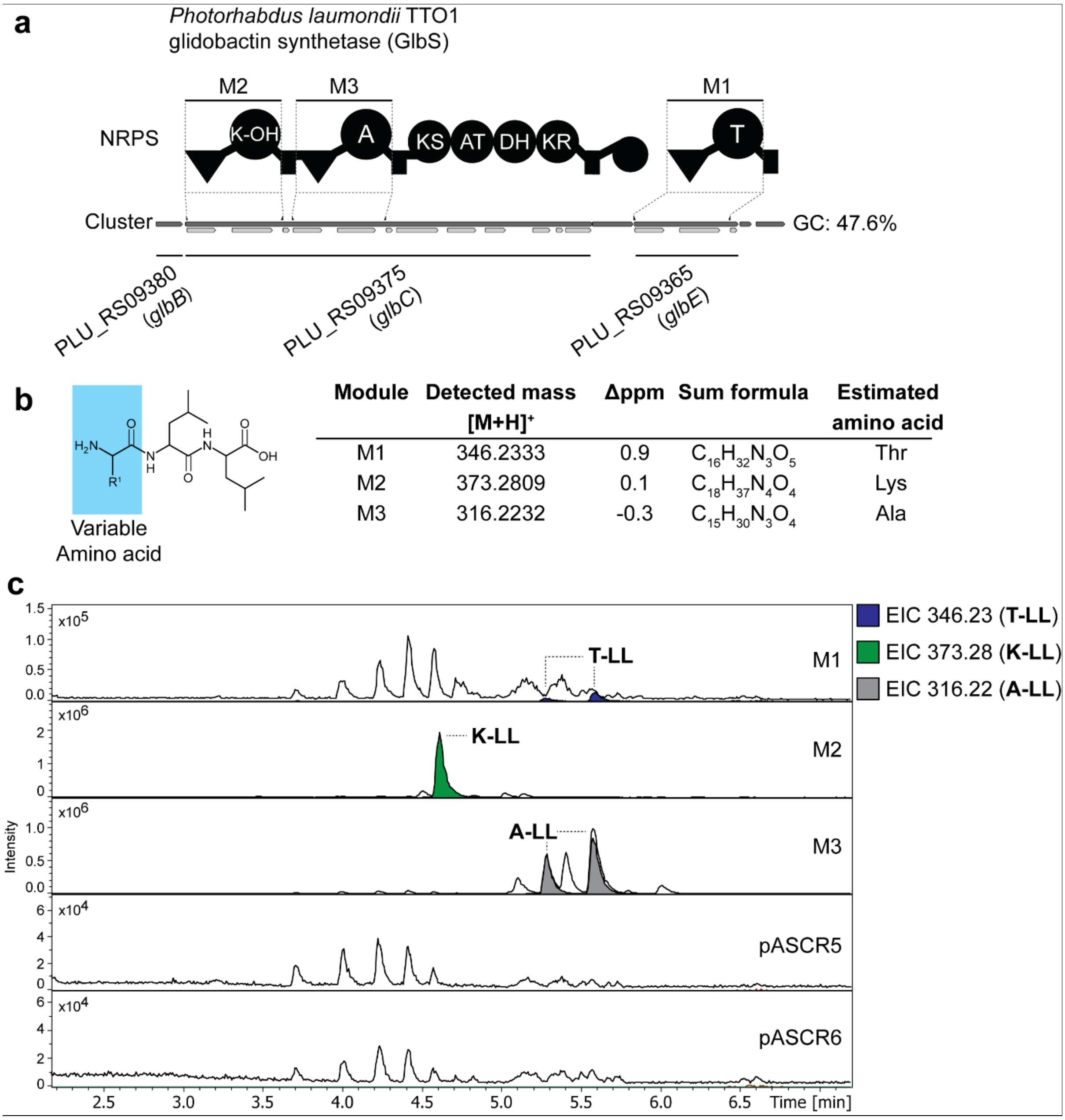

| Module | Detected mass $[M+H]^+$ | Δppm | Sum formula | Estimated amino acid |
|---|---|---|---|---|
| M1 | 346.2333 | 0.9 | $C_{16}H_{32}N_3O_5$ | Thr |
| M2 | 373.2809 | 0.1 | $C_{18}H_{37}N_4O_4$ | Lys |
| M3 | 316.2232 | -0.3 | $C_{15}H_{30}N_3O_4$ | Ala |

**Figure S74.** HPLC-MS/MS data referred to module screening of glidobactin synthetase (GlbS) from *P. laumondii* TTO1. (**a**) Illustration of the gene cluster of glidobactin synthetase from *P. laumondii* TTO1 with schematic illustration of the encoded NRPS and polyketide synthetase (PKS). Indication of the region of cloned modules M1 to M3 on gene and protein level. Utilized primers are shown above the cluster. A domain specitifity is indicated by the one-letter amino acid code (K-OH = γ-hydroxylysine). GC content is depicted next to the cluster. (**b**) Table of HPLC-MS/MS data indicating the targeted module cloned into pASCR3, the detected mass of the compound, the corresponding error, the sum formula and the incoporated amino acid introduced to the peptide at the highlighted position in the chemical structure (left). The amino acid is shown by the three-letter amino acid code (K-OH = γ-hydroxylysine). (**c**) HPLC-MS/MS data of *E. coli* DH10B::*mtaA* expressing M1 to M3 cloned into pASCR3. Expression of pASCR5 and pASCR6 served as negative control. Base Peak Chromatogram (BPC, top) with Extracted Ion Chromatogram (EIC, below with colors according to the depicted legend) of **T-LL** (*m/z* $[M+H]^+$ = 346.233), **K-LL** (*m/z* $[M+H]^+$ = 373.280) and **A-LL** (*m/z* $[M+H]^+$ = 316.223). Indication of the targeted modules on the right of the chromatograms.

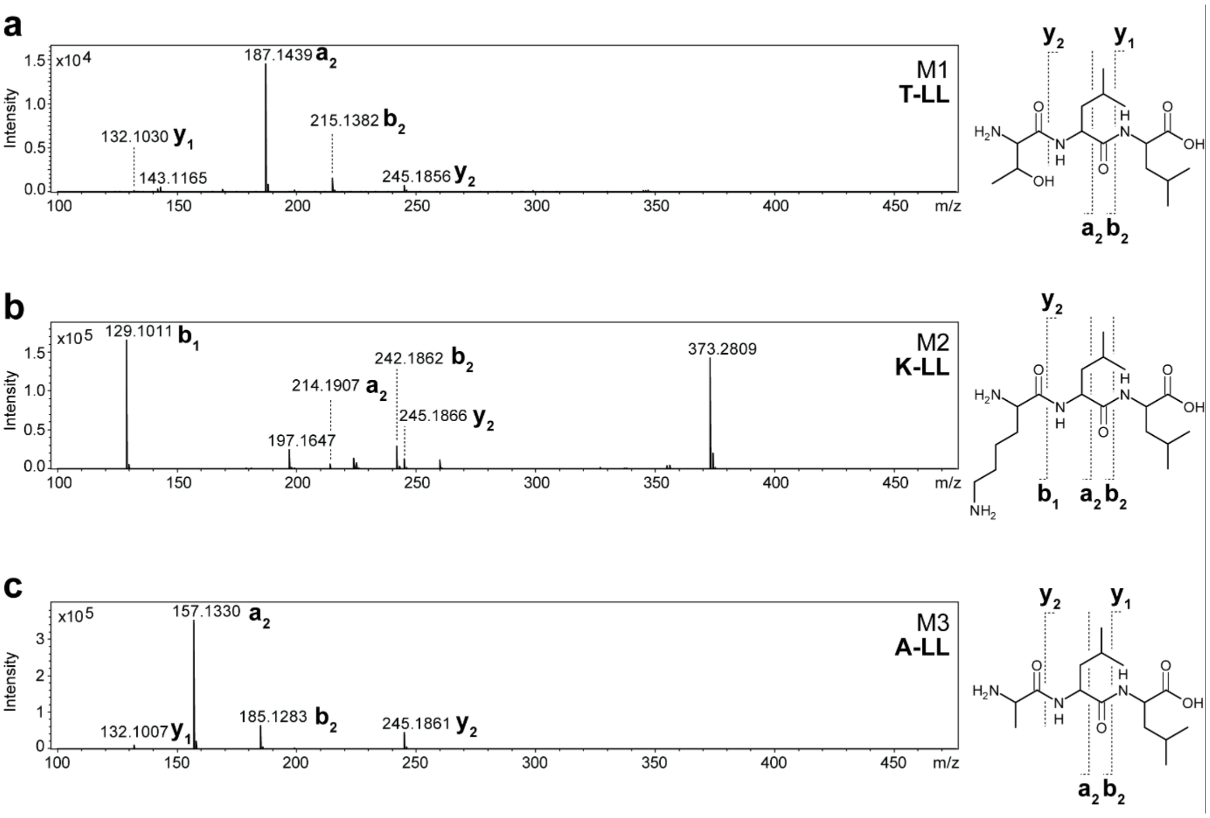

**Figure S75.** $MS^2$ spectra of HPLC-MS/MS data referred to figure S74. (**a-c**) The respective compounds are displayed, along with indications of the assigned fragments. The NRPS, targeted module and peptide name is depicted within the spectra.

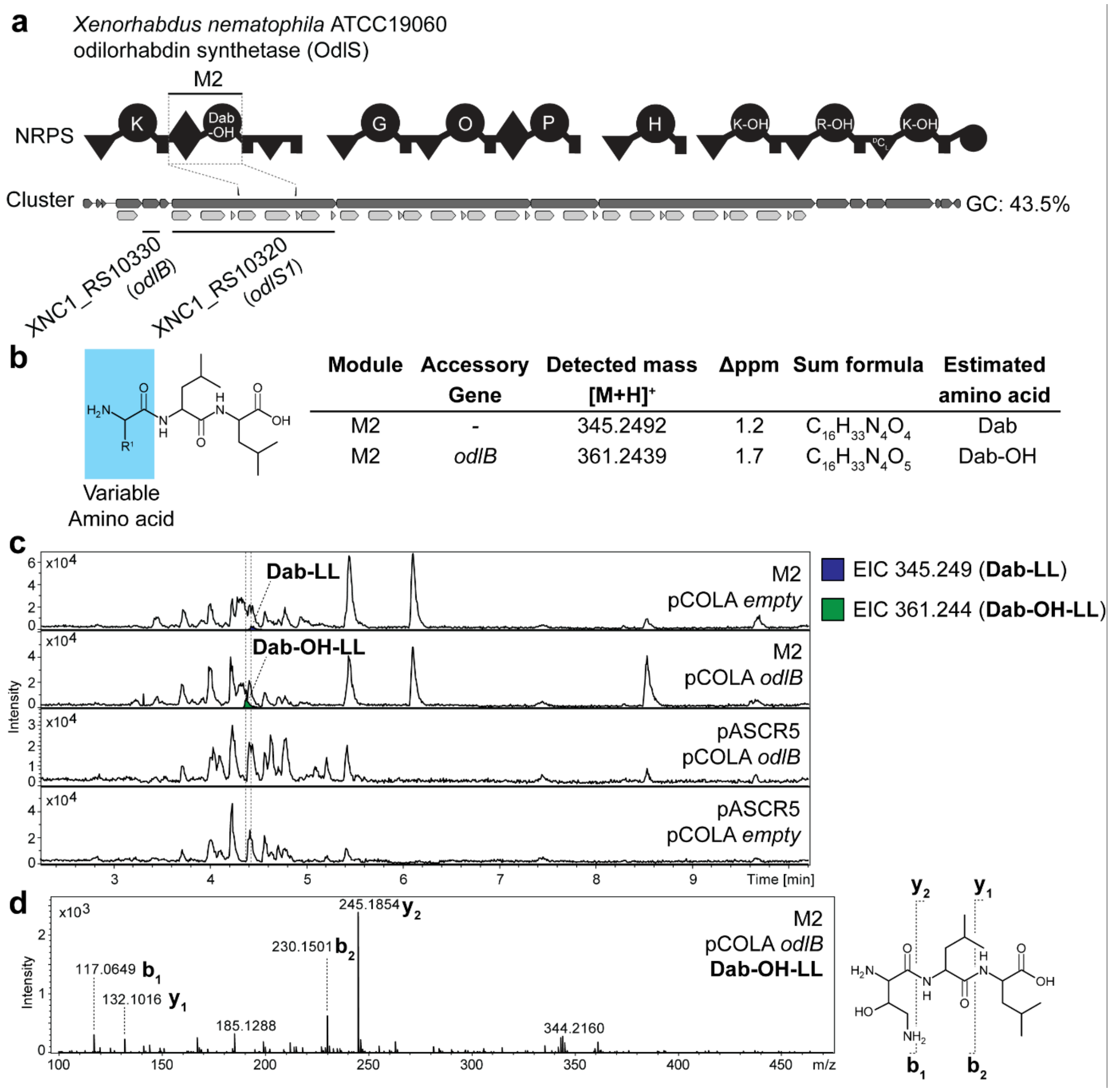

| Module | Accessory Gene | Detected mass [M+H]+ | Δppm | Sum formula | Estimated amino acid |
|---|---|---|---|---|---|
| M2 | - | 345.2492 | 1.2 | $C_{16}H_{33}N_4O_4$ | Dab |
| M2 | *odlB* | 361.2439 | 1.7 | $C_{16}H_{33}N_4O_5$ | Dab-OH |

**Figure S76.** HPLC-MS/MS data referred to accessory gene co-expression with odilorhabdin synthetase (OdlS) M2 from *X. nematophila*. (**a**) Illustration of the gene cluster of odilorhabdin synthetase (OdlS) from *X. nematophila* with schematic illustration of the encoded NRPS. Indication of the region of cloned modules M2 on gene and protein level. Utilized primers are shown above the cluster. A domain specitifity is indicated by the one-letter amino acid code (Dab-OH = β-hydroxy diamino butyric acid, O = ornithine, K-OH = δ-hydroxylysine, R-OH = β-hydroxy-arginine). GC content is depicted next to the cluster. (**b**) Table of HPLC-MS/MS data indicating the targeted module cloned into pASCR3, the co-expressed accessory gene, the detected mass of the compound, the corresponding error, the sum formula and the incoporated amino acid introduced to the peptide at the highlighted position in the chemical structure (left). The amino acid is shown by the three-letter amino acid code (Dab = diamino butyric acid, Dab-OH = β-hydroxy diamino butyric acid). (**c**) HPLC-MS/MS data of *E. coli* DH10B::*mtaA co-*expressing M2 cloned into pASCR3 and *odlB*. Expression of pASCR5 and pCOLA *empty* served as negative controls. Base Peak Chromatogram (BPC, top) with Extracted Ion Chromatogram (EIC, below with colors according to the depicted legend) of **Dab-LL** (*m/z* $[M+H]^+$ = 345.249) and **Dab-OH-LL** (*m/z* $[M+H]^+$ = 361.244). Indication of the targeted modules on the right of the chromatograms. (**d**) The $MS^2$ data for the respective compounds are displayed, along with indications of the assigned fragments.

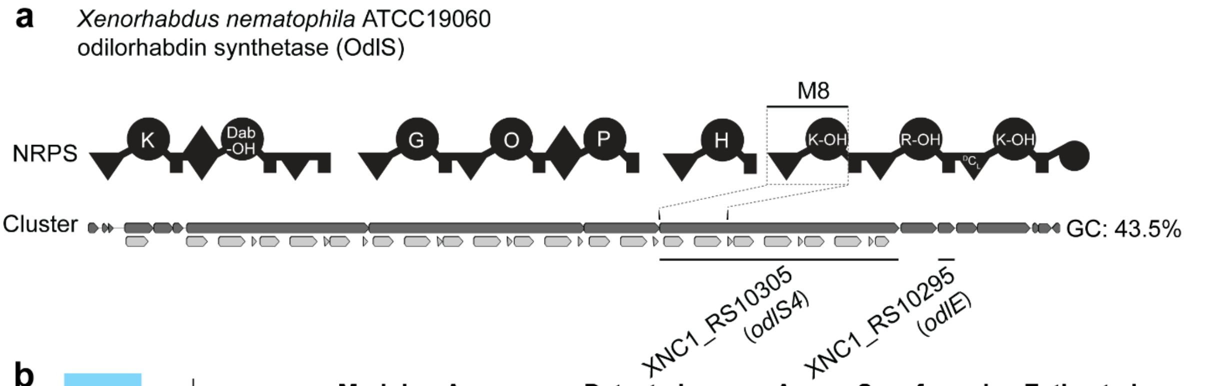

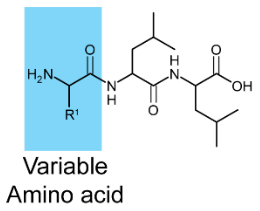

| Module | Accessory Gene | Detected mass [M+H]$^+$ | Δppm | Sum formula | Estimated amino acid |
|---|---|---|---|---|---|
| M8 | - | 373.2811 | -0.4 | $C_{18}H_{37}N_4O_4$ | Lys |
| M8 | *odlE* | 389.2760 | -0.3 | $C_{18}H_{37}N_4O_5$ | K-OH |

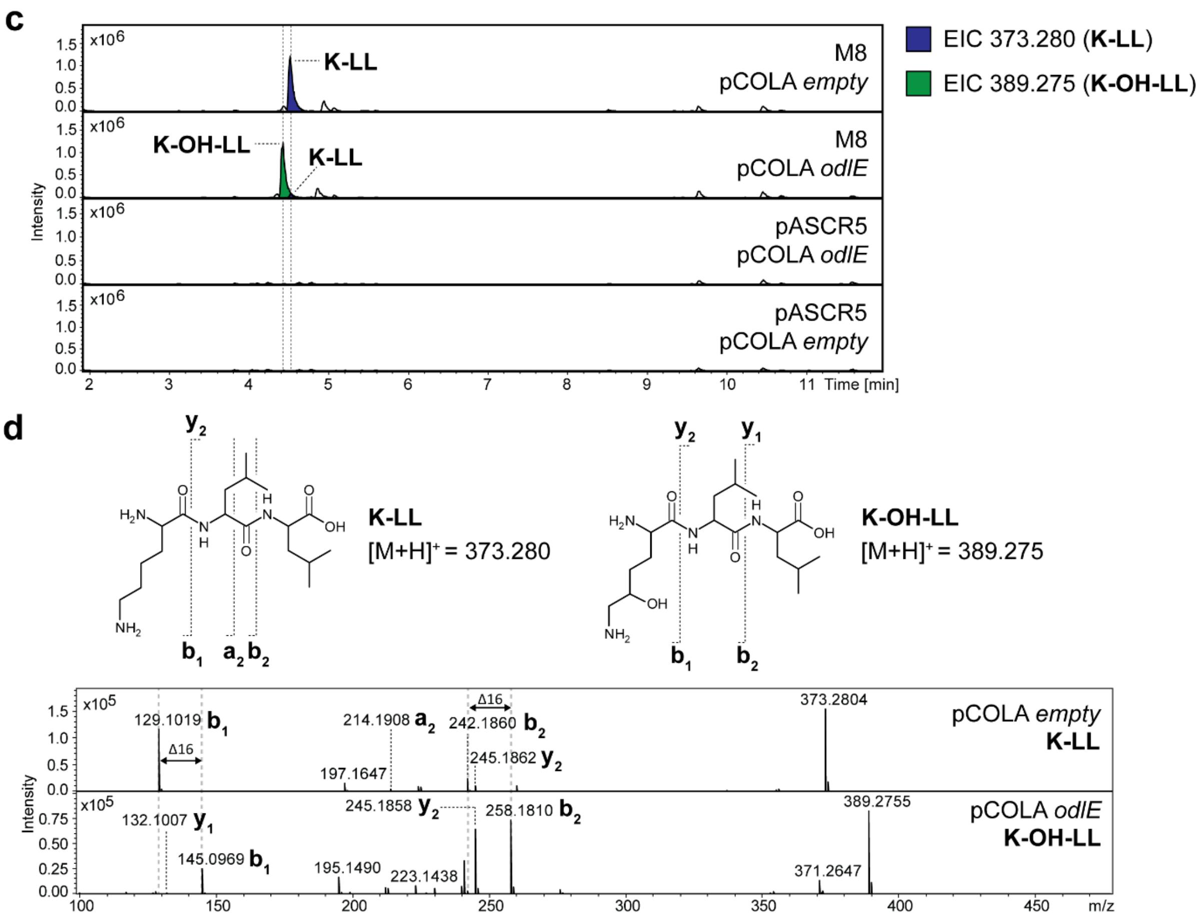

**Figure S77.** HPLC-MS/MS data referred to accessory gene co-expression with odilorhabdin synthetase (OdlS) M8 from *X. nematophila*. (**a**) Illustration of the gene cluster of odilorhabdin synthetase (OdlS) from *X. nematophila* with schematic illustration of the encoded NRPS. Indication of the region of cloned modules M8 on gene and protein level. Utilized primers are shown above the cluster. A domain specitifity is indicated by the one-letter amino acid code (Dab-OH = β-hydroxy diamino butyric acid, O = ornithine, K-OH = δ-hydroxylysine, R-OH = β-hydroxy-arginine). GC content is depicted next to the cluster. (**b**) Table of HPLC-MS/MS data indicating the targeted module cloned into pASCR3, the co-expressed accessory gene, the detected mass of the compound, the corresponding error, the sum formula and the incoporated amino acid introduced to the peptide at the highlighted position in the chemical structure (left). The amino acid is shown by the three-letter amino acid code (K-OH = δ-hydroxylysine). (**c**) HPLC-MS/MS data of *E. coli* DH10B::*mtaA co*-expressing M2 cloned into pASCR3 and *odlE*. Expression of pASCR5 and pCOLA *empty* served as negative controls. Base Peak Chromatogram (BPC, top) with Extracted Ion Chromatogram (EIC, below with colors according to the depicted legend) of **K-LL** (*m/z* $[M+H]^+$ = 373.280) and **K-OH-LL** (*m/z* $[M+H]^+$ = 389.275). Indication of the targeted modules on the right of the chromatograms. (**d**) The $MS^2$ data for the respective compounds are displayed, along with indications of the assigned fragments.

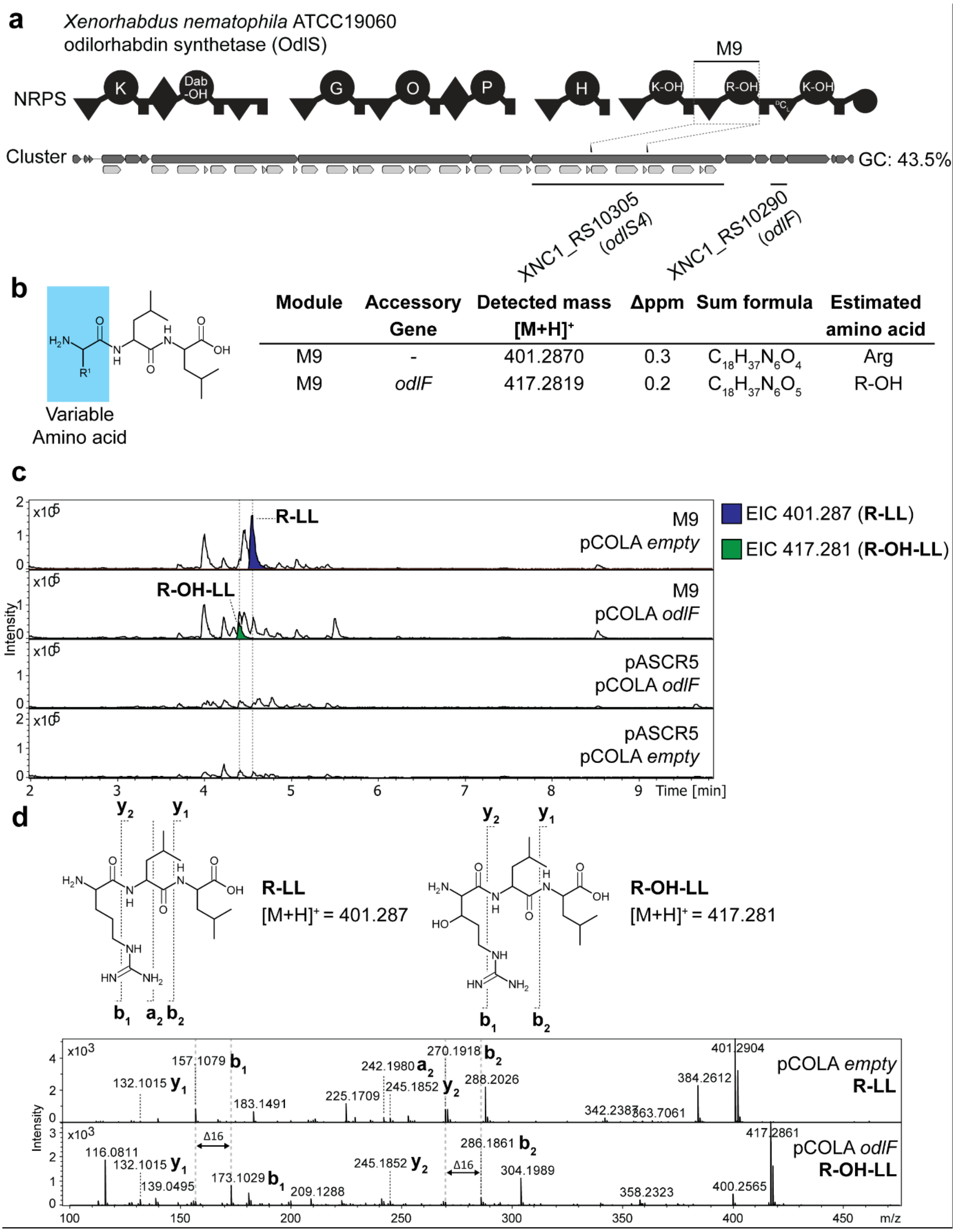

**Figure S78.** HPLC-MS/MS data referred to accessory gene co-expression with odilorhabdin synthetase (OdlS) M9 from *X. nematophila*. (**a**) Illustration of the gene cluster of odilorhabdin synthetase (OdlS) from *X. nematophila* with schematic illustration of the encoded NRPS. Indication of the region of cloned modules M9 on gene and protein level. Utilized primers are shown above the cluster. A domain specitifity is indicated by the one-letter amino acid code (Dab-OH = β-hydroxy diamino butyric acid, O = ornithine, K-OH = δ-hydroxylysine, R-OH = β-hydroxy-arginine). GC content is depicted next to the cluster. (**b**) Table of HPLC-MS/MS data indicating the targeted module cloned into pASCR3, the co-expressed accessory gene, the detected mass of the compound, the corresponding error, the sum formula and the incoporated amino acid introduced to the peptide at the highlighted position in the chemical structure (left). The amino acid is shown by the three-letter amino acid code (R-OH = β-hydroxy-arginine). (**c**) HPLC-MS/MS data of *E. coli* DH10B::*mtaA co*-expressing M9 cloned into pASCR3 and *odlF*. Expression of pASCR5 and pCOLA *empty* served as negative controls. Base Peak Chromatogram (BPC, top) with Extracted Ion Chromatogram (EIC, below with colors according to the depicted legend) of **R-LL** (*m/z* $[M+H]^+$ = 401.287) and **R-OH-LL** (*m/z* $[M+H]^+$ = 417.281). Indication of the targeted modules on the right of the chromatograms. (**d**) The $MS^2$ data for the respective compounds are displayed, along with indications of the assigned fragments.

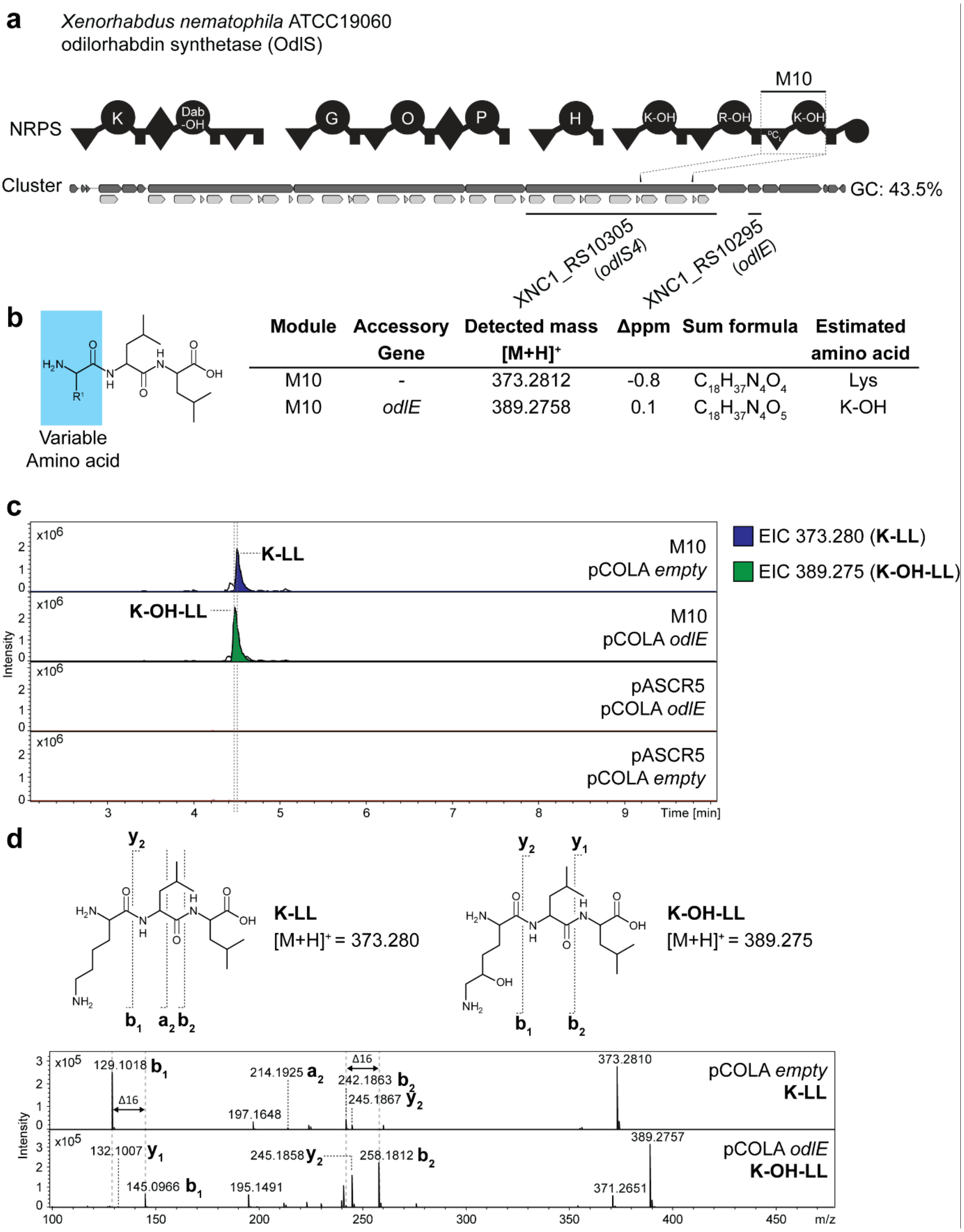

**Figure S79.** HPLC-MS/MS data referred to accessory gene co-expression with odilorhabdin synthetase (OdlS) M10 from *X. nematophila*. (**a**) Illustration of the gene cluster of a odilorhabdin synthetase (OdlS) from *X. nematophila* with schematic illustration of the encoded NRPS. Indication of the region of cloned modules M10 on gene and protein level. Utilized primers are shown above the cluster. A domain specitifity is indicated by the one-letter amino acid code (Dab-OH = β-hydroxy diamino butyric acid, O = ornithine, K-OH = δ-hydroxylysine, R-OH = β-hydroxy-arginine). GC content is depicted next to the cluster. (**b**) Table of HPLC-MS/MS data indicating the targeted module cloned into pASCR3, the co-expressed accessory gene, the detected mass of the compound, the corresponding error, the sum formula and the incoporated amino acid introduced to the peptide at the highlighted position in the chemical structure (left). The amino acid is shown by the three-letter amino acid code (K-OH = δ-hydroxylysine). (**c**) HPLC-MS/MS data of *E. coli* DH10B::*mtaA co*-expressing M10 cloned into pASCR3 and *odlE*. Expression of pASCR5 and pCOLA *empty* served as negative controls. Base Peak Chromatogram (BPC, top) with Extracted Ion Chromatogram (EIC, below with colors according to the depicted legend) of **K-LL** (*m/z* $[M+H]^+$ = 373.280) and **K-OH-LL** (*m/z* $[M+H]^+$ = 389.275). Indication of the targeted modules on the right of the chromatograms. (**d**) The $MS^2$ data for the respective compounds are displayed, along with indications of the assigned fragments.

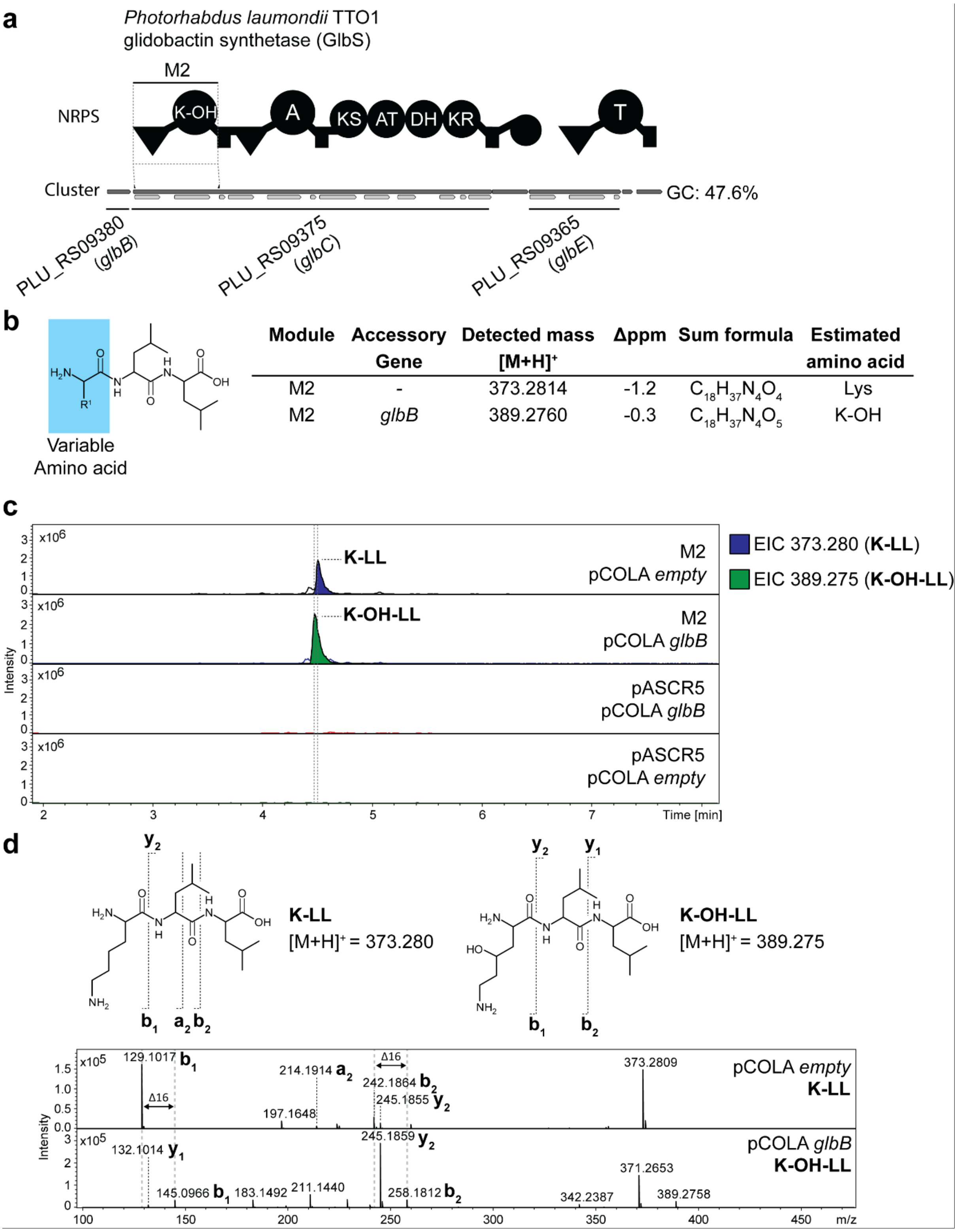

**Figure S80.** HPLC-MS/MS data referred to accessory gene co-expression with glidobactin synthetase (GlbS) M2 from *P. laumondii* TTO1. (**a**) Illustration of the gene cluster of a glidobactin synthetase (GlbS) from *P. laumondii* TTO1 with schematic illustration of the encoded NRPS. Indication of the region of cloned modules M2 on gene and protein level. Utilized primers are shown above the cluster. A domain specitifity is indicated by the one-letter amino acid code (K-OH = γ-hydroxylysine). GC content is depicted next to the cluster. (**b**) Table of HPLC-MS/MS data indicating the targeted module cloned into pASCR3, the co-expressed accessory gene, the detected mass of the compound, the corresponding error, the sum formula and the incoporated amino acid introduced to the peptide at the highlighted position in the chemical structure (left). The amino acid is shown by the three-letter amino acid code (K-OH = γ-hydroxylysine). (**c**) HPLC-MS/MS data of *E. coli* DH10B::*mtaA co*-expressing M2 cloned into pASCR3 and *glbB*. Expression of pASCR5 and pCOLA *empty* served as negative controls. Base Peak Chromatogram (BPC, top) with Extracted Ion Chromatogram (EIC, below with colors according to the depicted legend) of **K-LL** (*m/z* $[M+H]^+$ = 373.280) and **K-OH-LL** (*m/z* $[M+H]^+$ = 389.275). Indication of the targeted modules on the right of the chromatograms. (**d**) The $MS^2$ data for the respective compounds are displayed, along with indications of the assigned fragments.

**a**

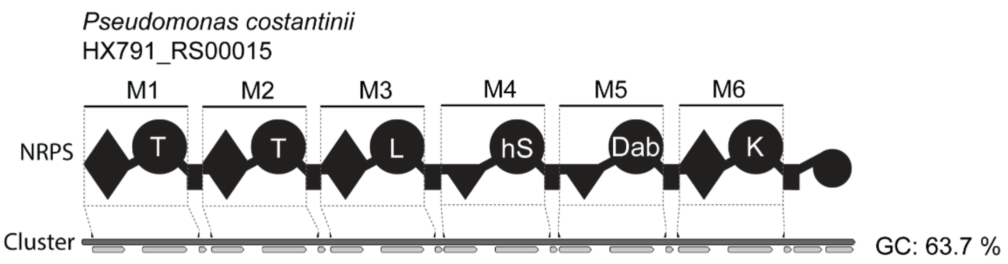

**b**

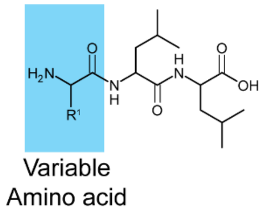

| Module | Detected mass $[M+H]^+$ | Δppm | Sum formula | Estimated amino acid |
|---|---|---|---|---|
| M1 | 346.2332 | 1.2 | $C_{16}H_{32}N_3O_5$ | Thr |
| M2 | 346.2339 | -0.8 | $C_{16}H_{32}N_3O_5$ | Thr |
| M3 | 358.2701 | -0.2 | $C_{18}H_{36}N_3O_4$ | Leu |
| M4 | 346.2334 | 0.6 | $C_{16}H_{32}N_3O_5$ | hS |
| M5 | 345.2496 | 0.1 | $C_{16}H_{33}N_4O_4$ | Dab |
| M6 | 373.2807 | 0.6 | $C_{18}H_{37}N_4O_4$ | Lys |

**c**

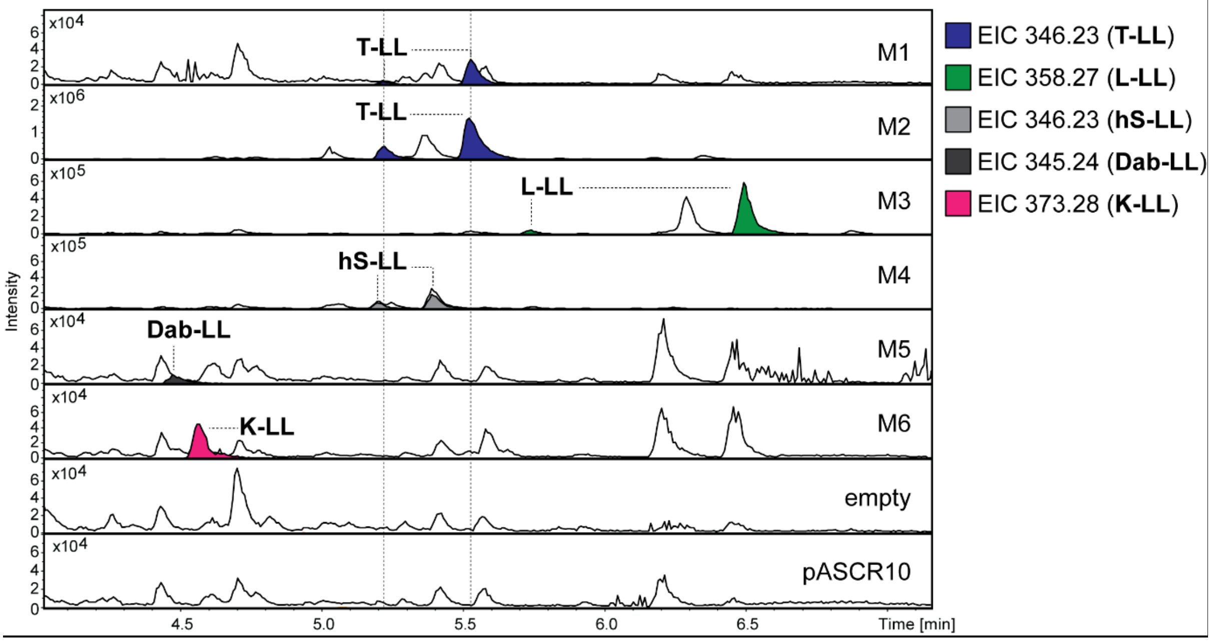

**Figure S81.** HPLC-MS/MS data referred to module screening of HX791_RS00015 from *P. costantinii*. (**a**) Illustration of the gene cluster HX791_RS00015 from *P. costantinii* with schematic illustration of the encoded NRPS. Indication of the region of cloned modules M1 to M6 on gene and protein level. Utilized primers are shown above the cluster. A domain specitifity is indicated by the one-letter amino acid code (hS = homoserine; Dab = diaminobuytric acid). GC content is depicted next to the cluster. (**b**) Table of HPLC-MS/MS data indicating the targeted module cloned into pASCR8, the detected mass of the compound, the corresponding error, the sum formula and the incoporated amino acid introduced to the peptide at the highlighted position in the chemical structure (left). The amino acid is shown by the three-letter amino acid code (hS = homoserine; Dab = diaminobuytric acid). (**c**) HPLC-MS/MS data of *E. coli* DH10B::*mtaA* expressing M1 to M6 cloned into pASCR8. Expression of an empty vector and pASCR10 served as negative controls. Base Peak Chromatogram (BPC, top) with Extracted Ion Chromatogram (EIC, below with colors according to the depicted legend) of **T-LL** (*m/z* $[M+H]^+$ = 346.233), **L-LL** (*m/z* $[M+H]^+$ = 358.270), **hS-LL** (*m/z* $[M+H]^+$ = 346.233), **Dab-LL** (*m/z* $[M+H]^+$ = 345.249) and **K-LL** (*m/z* $[M+H]^+$ = 373.280). Indication of the targeted modules on the right of the chromatograms.

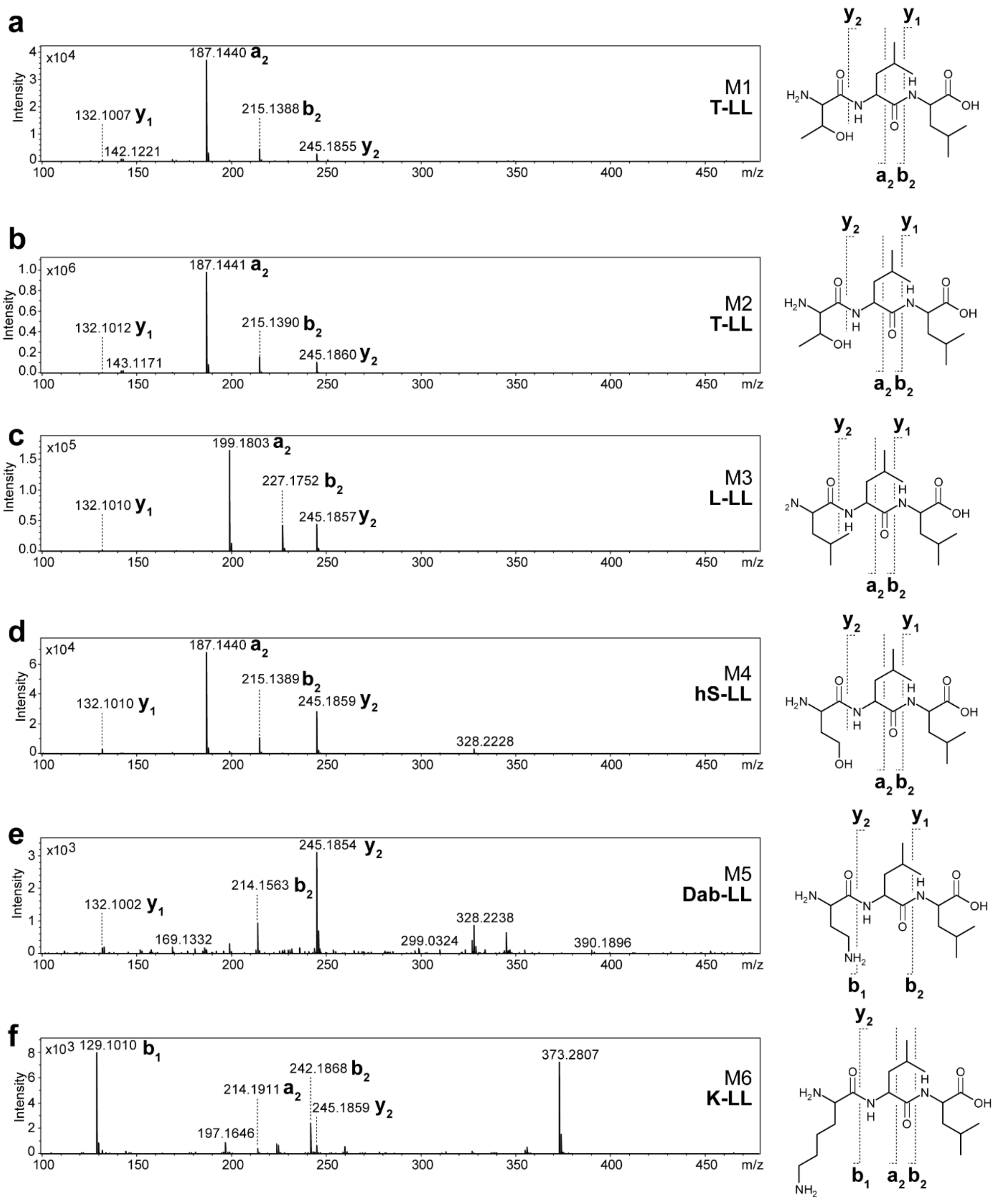

**Figure S82.** $MS^2$ spectra of HPLC-MS/MS data referred to figure S81. (**a-f**) The respective compounds are displayed, along with indications of the assigned fragments. The targeted module and peptide name is depicted within the spectra.

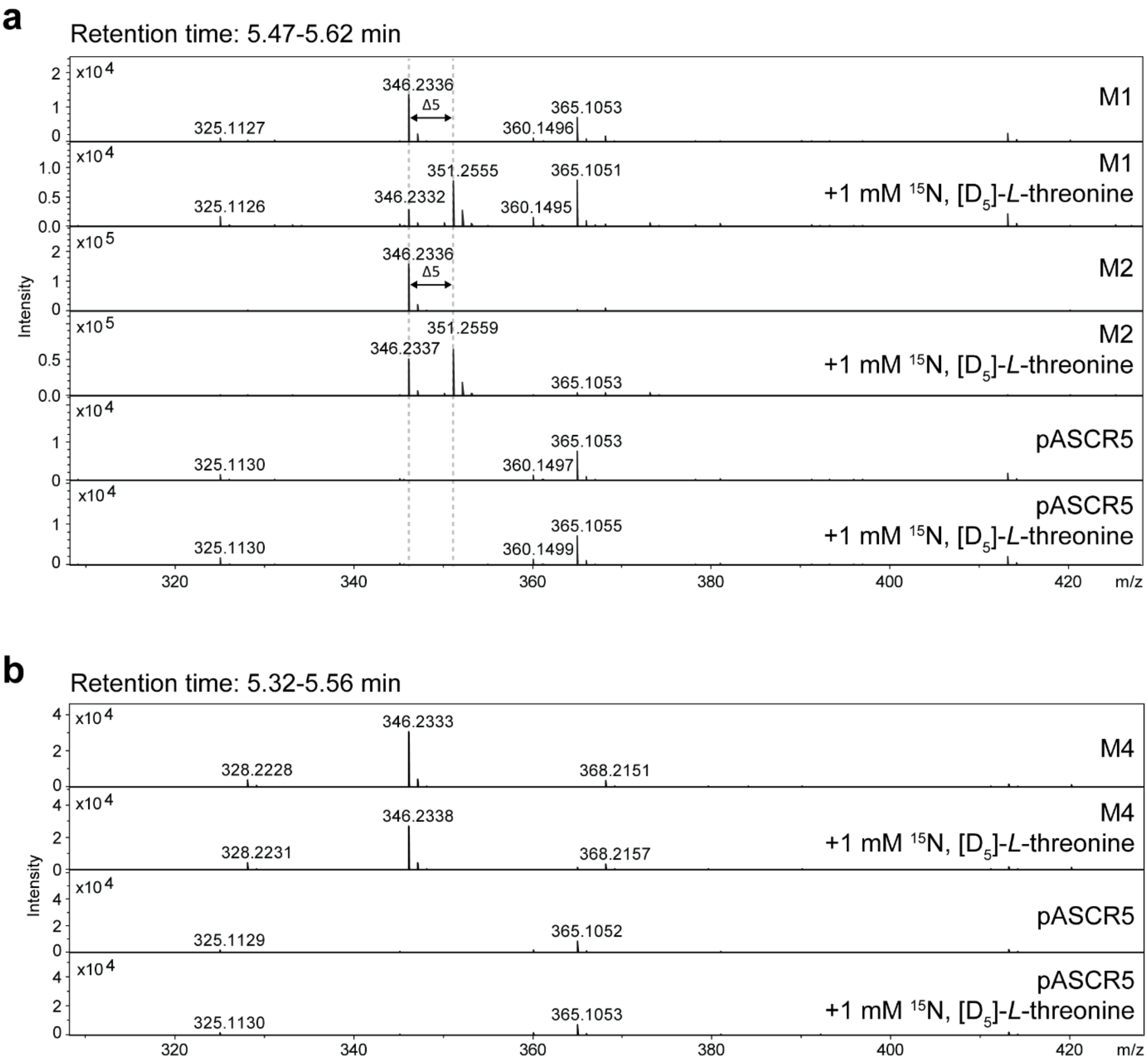

**Figure S83.** MS spectra of HPLC-MS/MS data referred to module screening of HX791_RS00015 from *P. costantinii* (see Fig. S80). Expression of M1, M2 and M4 cloned into pASCR3 in *E. coli* DH10B::*mtaA* cultivated in Bacto™ CD Supreme Fermentation production medium (0.5 g/L L-leucine and 10 g/L glycerol) with and without supplementation of 1 mM $^{15}N$, [$D_5$]-*L*-threonine. Expression of pASCR5 served as negative control. The MS spectra of retention time 5.47-5.62 min is displayed for M1 and M2 (**a**) while the retention time of 5.32-5.56 min is displayed for M4 (**b**).

a

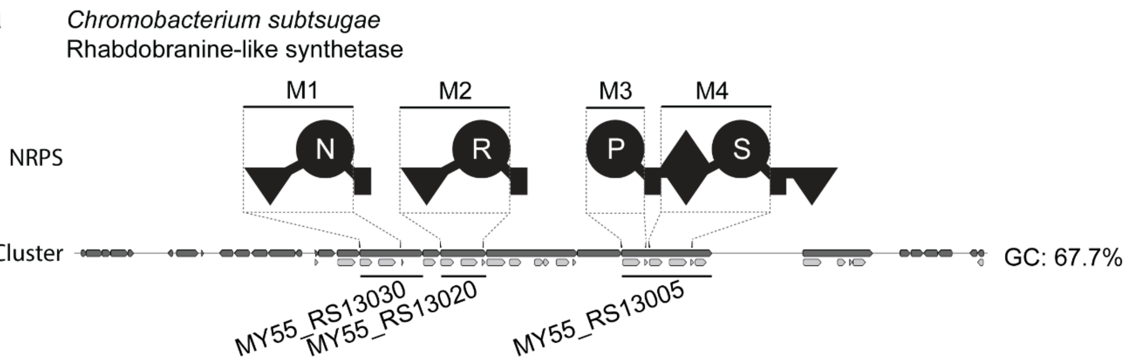

b

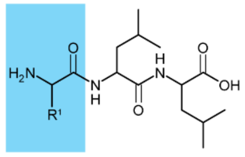

| Module | Detected mass $[M+H]^+$ | Δppm | Sum formula | Estimated amino acid |
|---|---|---|---|---|
| M1 | 359.2288 | 0.3 | $C_{16}H_{31}N_4O_5$ | Asn |
| M2 | 401.2872 | -0.4 | $C_{18}H_{37}N_6O_4$ | Arg |
| M3 | 342.2392 | -1.4 | $C_{17}H_{32}N_3O_4$ | Pro |
| M4 | 332.2179 | 0.2 | $C_{15}H_{30}N_3O_5$ | Ser |

c

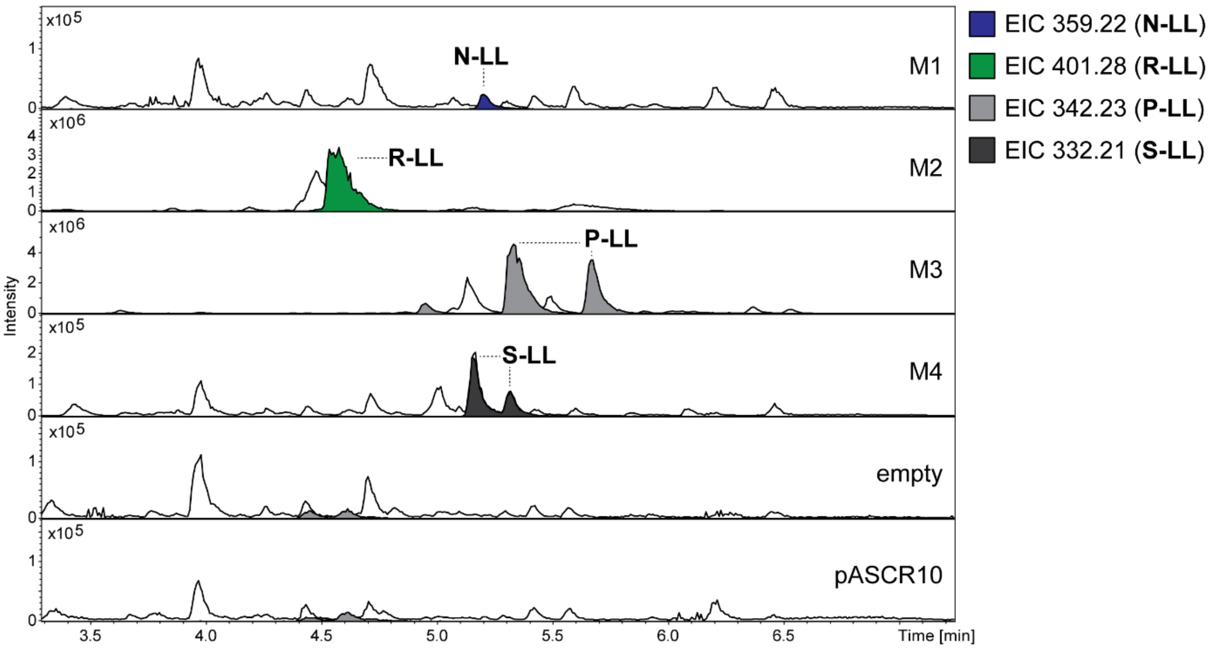

**Figure S84.** HPLC-MS/MS data referred to module screening of a rhabdobranine-like synthetase from *C. subtsugae*. (**a**) Illustration of the gene cluster of a rhabdobranine-like synthetase from *C. subtsugae* with schematic illustration of the encoded NRPS. Indication of the region of cloned modules M1 to M4 on gene and protein level. Utilized primers are shown above the cluster. A domain specitifity is indicated by the one-letter amino acid code. GC content is depicted next to the cluster. (**b**) Table of HPLC-MS/MS data indicating the targeted module cloned into pASCR8, the detected mass of the compound, the corresponding error, the sum formula and the incoporated amino acid introduced to the peptide at the highlighted position in the chemical structure (left). The amino acid is shown by the three-letter amino acid code. (**c**) HPLC-MS/MS data of *E. coli* DH10B::*mtaA* expressing M1 to M4 cloned into pASCR8. Expression of an empty vector and pASCR10 served as negative controls. Base Peak Chromatogram (BPC, top) with Extracted Ion Chromatogram (EIC, below with colors according to the depicted legend) of **N-LL** (*m/z* $[M+H]^+$ = 359.228), **R-LL** (*m/z* $[M+H]^+$ = 401.287), **P-LL** (*m/z* $[M+H]^+$ = 342.238) and **S-LL** (*m/z* $[M+H]^+$ = 332.217). Indication of the targeted modules on the right of the chromatograms.

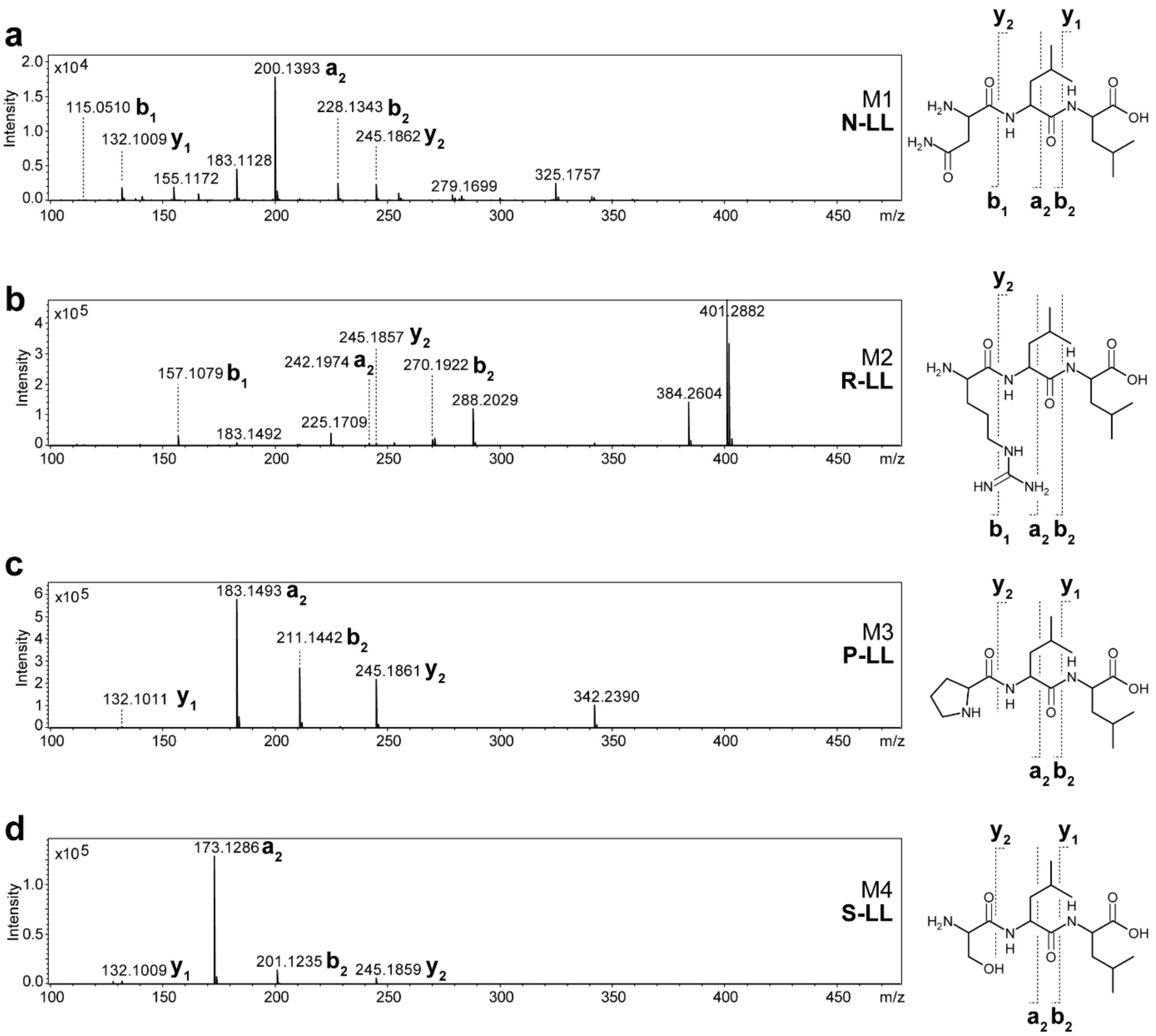

**Figure S85.** $MS^2$ spectra of HPLC-MS/MS data referred to figure S84. (**a-d**) The respective compounds are displayed, along with indications of the assigned fragments. The targeted module and peptide name is depicted within the chromatogram.

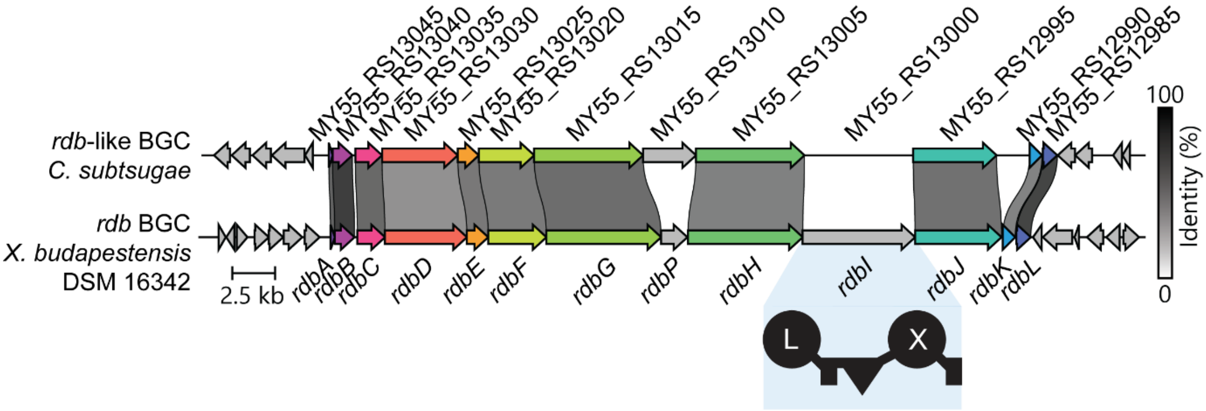

**Figure S86.** Comparison of *rdb*-like BGC from *C. subtsugae* and *rdb* BGC[25] from *X. budapestensis* using clinker[26]. Locus tag (*rdb*-like BGC) or gene names (*rdb* BGC) are indicated with percentage identity among genes. *rdbI* encoding for an NRPS is apparently not present in in the *rdb*-like BGC.

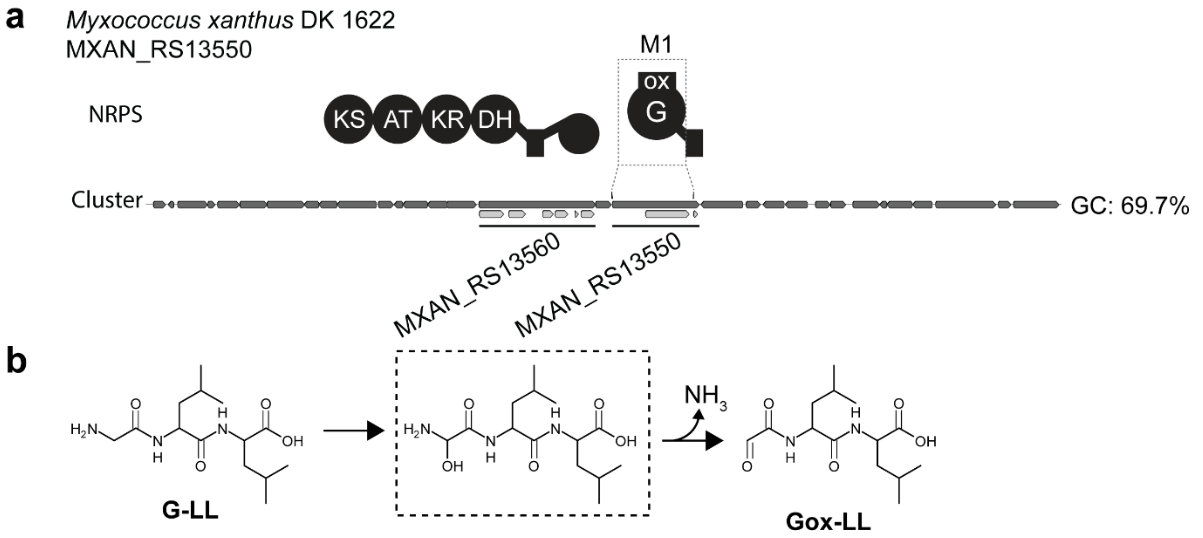

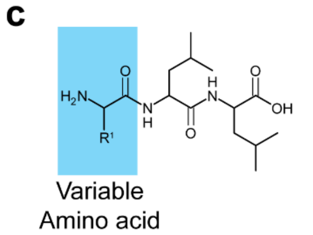

| Module | Detected mass [M+H]+ | Δppm | Sum formula | Estimated amino acid |
|---|---|---|---|---|
| M1 | 301.1758 | -0.1 | $C_{14}H_{25}N_2O_5$ | Gox |
| M1 | 302.2072 | 0.8 | $C_{14}H_{28}N_3O_4$ | Gly |
| M1 | 375.2122 | 0.9 | $C_{17}H_{31}N_2O_7$ | Gox |
| M1 | 357.2018 | 0.7 | $C_{17}H_{29}N_2O_6$ | ? |

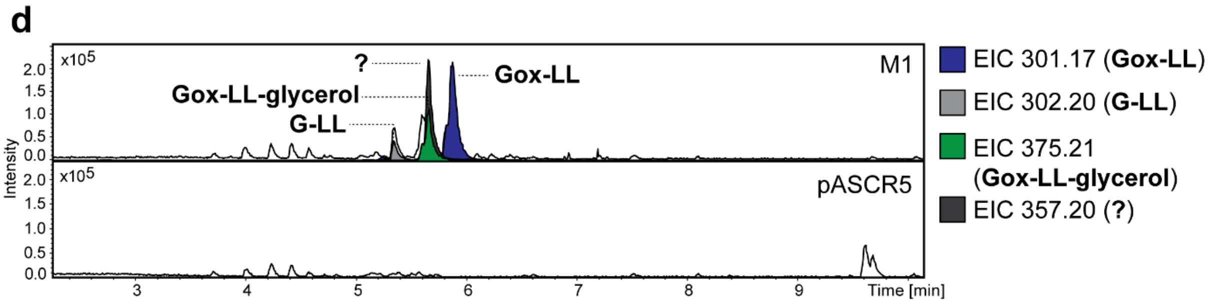

**Figure S87.** HPLC-MS/MS data referred to module screening of a MXAN_RS13550 from *M. xanthus*. (**a**) Illustration of the gene cluster of a XAN_RS13550 from *M. xanthus* with schematic illustration of the encoded NRPS and PKS. Indication of the region of cloned modules M1 on gene and protein level. Utilized primers are shown above the cluster. A domain specitifity is indicated by the one-letter amino acid code. GC content is depicted next to the cluster. (**b**) Table of HPLC-MS/MS data indicating the targeted module cloned into pASCR3, the detected mass of the compound, the corresponding error, the sum formula and the incoporated amino acid introduced to the peptide at the highlighted position in the chemical structure (left). The amino acid is shown by the three-letter amino acid code (Gox = glyoxal). (**c**) HPLC-MS/MS data of *E. coli* DH10B::*mtaA* expressing M1 cloned into pASCR3. Expression of pASCR5 served as negative control. Base Peak Chromatogram (BPC, top) with Extracted Ion Chromatogram (EIC, below with colors according to the depicted legend) of **Gox-LL** (*m/z* $[M+H]^+$ = 301.175), **G-LL** (*m/z* $[M+H]^+$ = 302.207), **Gox-LL-glycerol** (*m/z* $[M+H]^+$ = 375.211) and unkown compound **?** (*m/z* $[M+H]^+$ = 357.20). Indication of the targeted modules on the right of the chromatograms.

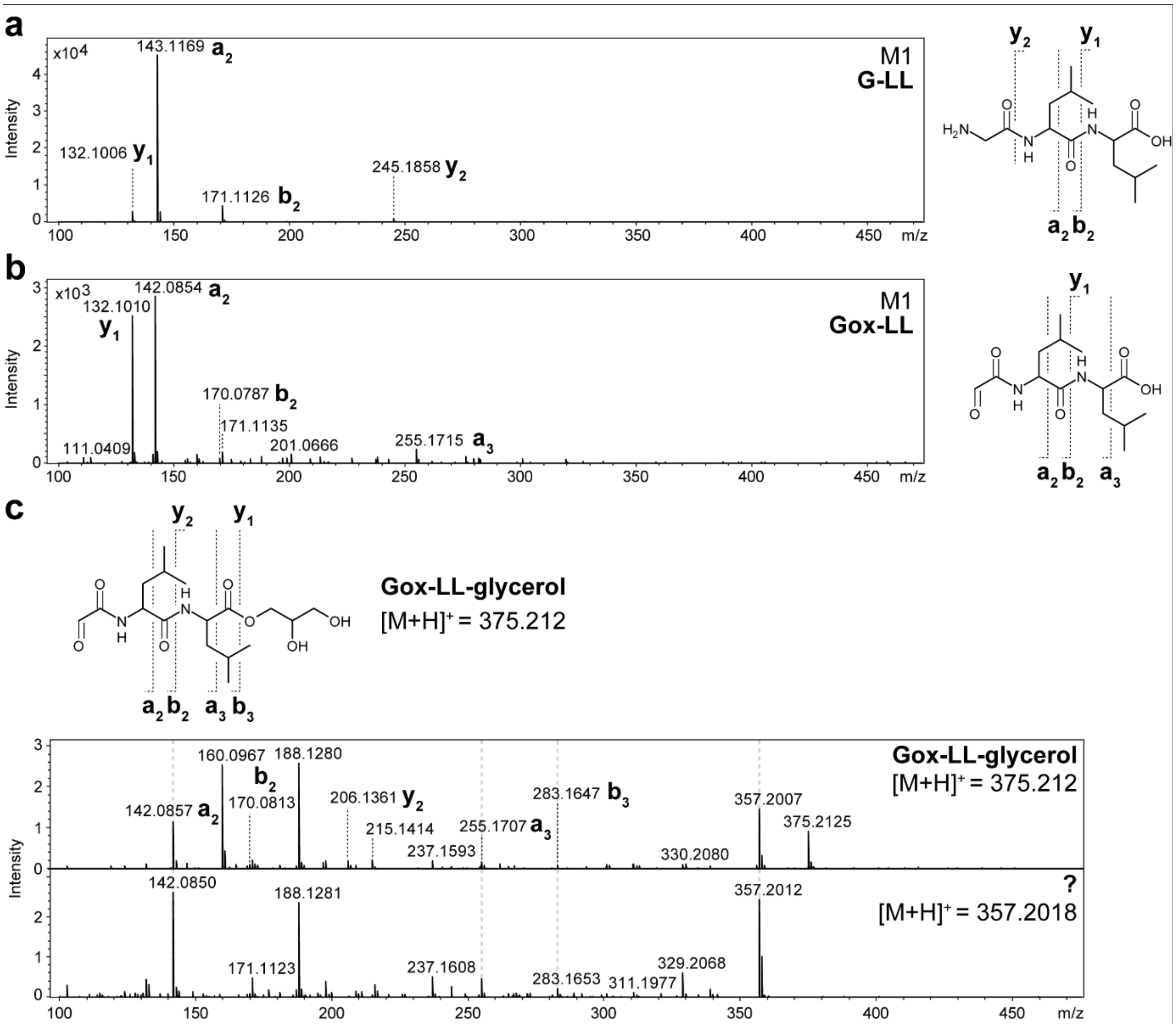

**Figure S88.** $MS^2$ spectra of HPLC-MS/MS data referred to figure S87. (**a-c**) The respective compounds are displayed, along with indications of the assigned fragments. The targeted module and peptide name is depicted within the spectra. (**c**) Comparison of $MS^2$ spectra of the **Gox-LL-glycerol** peptide and the unknown compound **?**.

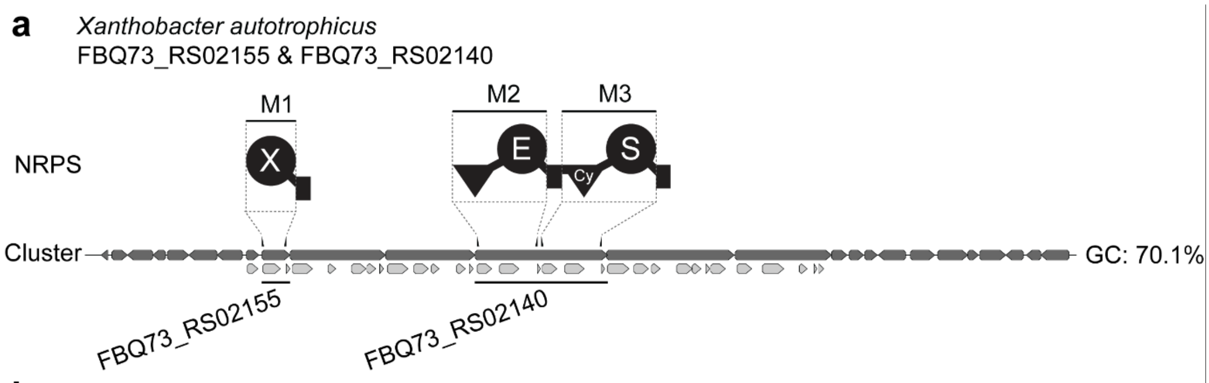

b

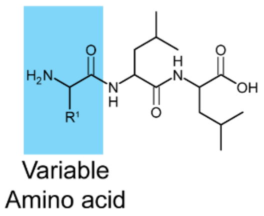

| Module | Detected mass [M+H]+ | Δppm | Sum formula | Estimated amino acid |
|---|---|---|---|---|
| M1 | - | - | - | - |
| M2 | 374.2289 | -0.9 | $C_{17}H_{32}N_3O_6$ | Glu |
| M3 | 332.2177 | 0.8 | $C_{15}H_{30}N_3O_5$ | Ser |

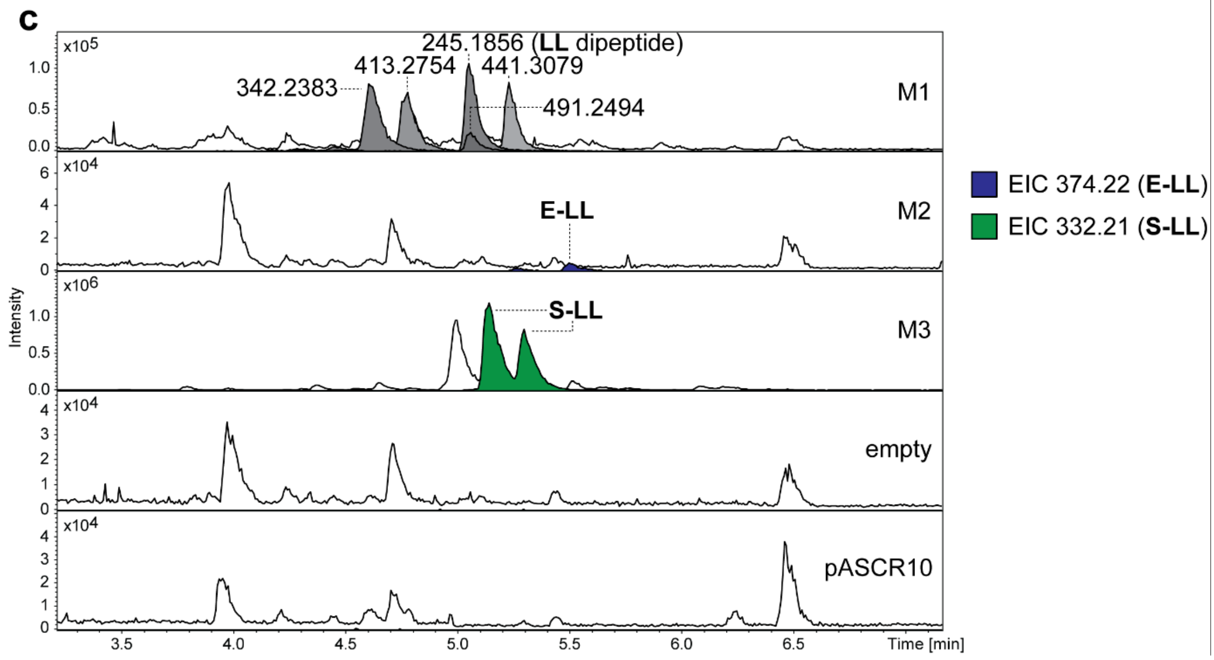

**Figure S89.** HPLC-MS/MS data referred to module screening of a FBQ73_RS02155 and FBQ73_RS02140 from *X. autotrophicus*. (**a**) Illustration of the gene cluster of a FBQ73_RS02155 and FBQ73_RS02140 from *X. autotrophicus* with schematic illustration of the encoded NRPS (Cy = heterocyclization domain). Indication of the region of cloned modules M1 to M3 on gene and protein level. Utilized primers are shown above the cluster. A domain specitifity is indicated by the one-letter amino acid code. GC content is depicted next to the cluster. (**b**) Table of HPLC-MS/MS data indicating the targeted module cloned into pASCR8, the detected mass of the compound, the corresponding error, the sum formula and the incoporated amino acid introduced to the peptide at the highlighted position in the chemical structure (left). The amino acid is shown by the three-letter amino acid code. (**c**) HPLC-MS/MS data of *E. coli* DH10B::*mtaA* expressing M1 to M3 cloned into pASCR8. Expression of an empty vector and pASCR10 served as negative control. Base Peak Chromatogram (BPC, top) with Extracted Ion Chromatogram (EIC, below with colors according to the depicted legend) of **E-LL** (*m/z* $[M+H]^+$ = 374.228) and **S-LL** (*m/z* $[M+H]^+$ = 332.217). Indication of the targeted modules on the right of the chromatograms.

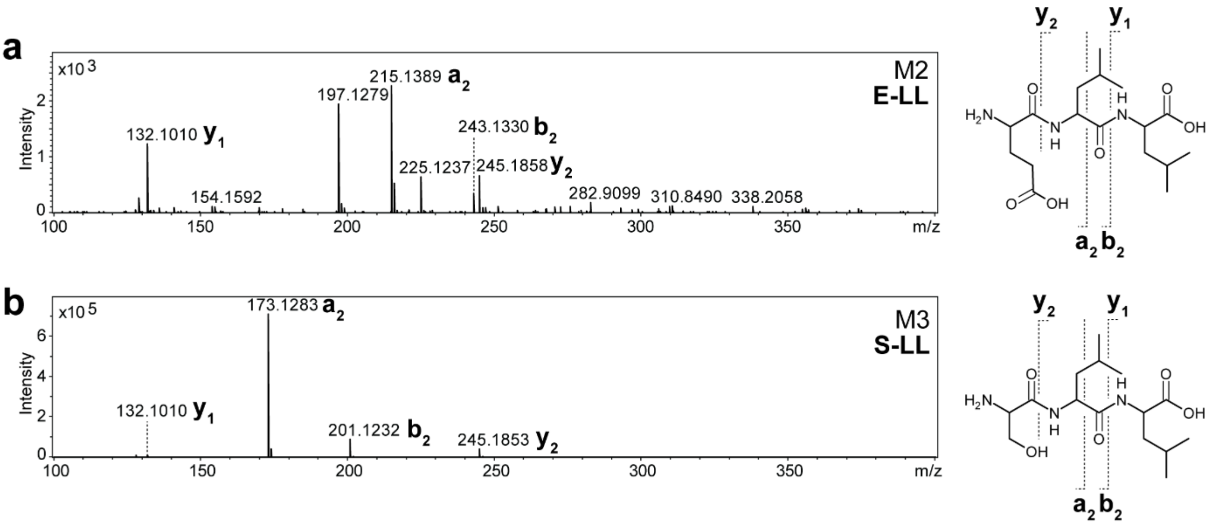

**Figure S90.** $MS^2$ spectra of HPLC-MS/MS data referred to figure S89. (**a** and **b**) The respective compounds are displayed, along with indications of the assigned fragments. The targeted module and peptide name is depicted within the spectra.

a

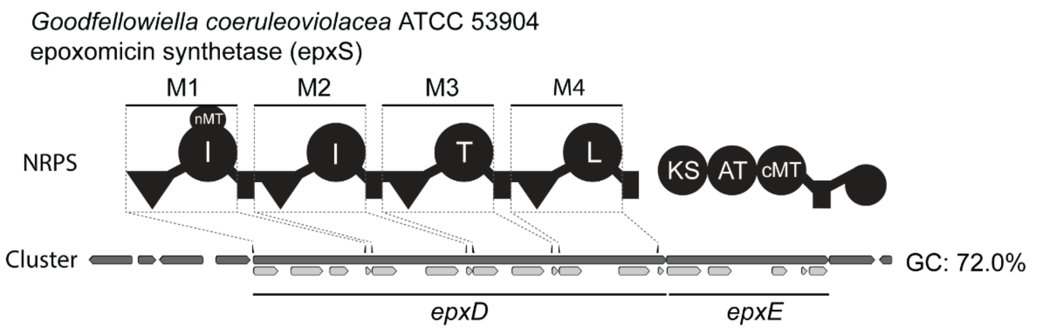

b

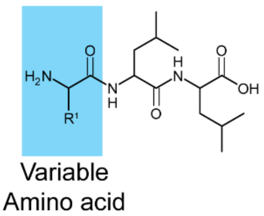

| Module | Detected mass $[M+H]^+$ | Δppm | Sum formula | Estimated amino acid |
|---|---|---|---|---|
| M1 | - | - | - | - |
| M2 | 358.2698 | 0.6 | $C_{18}H_{36}N_3O_4$ | Ile |
| M2 | 346.2338 | -0.4 | $C_{16}H_{32}N_3O_5$ | Thr |
| M3 | 358.2699 | 0.4 | $C_{18}H_{36}N_3O_4$ | Leu |

c

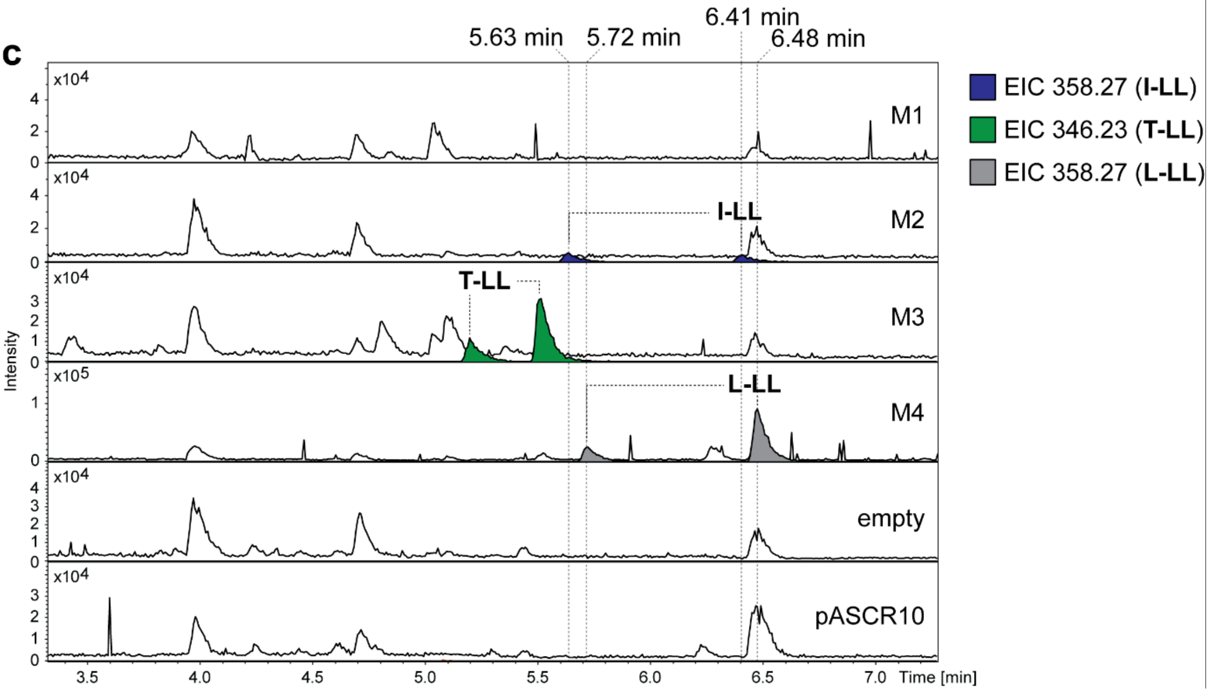

**Figure S91.** HPLC-MS/MS data referred to module screening of epoxomicin synthetase (EpxS) from *G. coeruleoviolacea*. (**a**) Illustration of the gene cluster of EpxS from *G. coeruleoviolacea* with schematic illustration of the encoded NRPS and polyketide synthetase (PKS). Indication of the region of cloned modules M1 to M4 on gene and protein level. Utilized primers are shown above the cluster. A domain specitifity is indicated by the one-letter amino acid code. GC content is depicted next to the cluster. (**b**) Table of HPLC-MS/MS data indicating the targeted module cloned into pASCR8, the detected mass of the compound, the corresponding error, the sum formula and the incoporated amino acid introduced to the peptide at the highlighted position in the chemical structure (left). The amino acid is shown by the three-letter amino acid code. (**c**) HPLC-MS/MS data of *E. coli* DH10B::*mtaA* expressing M1 to M4 cloned into pASCR8. Expression of an empty vector and pASCR10 served as negative control. Base Peak Chromatogram (BPC, top) with Extracted Ion Chromatogram (EIC, below with colors according to the depicted legend) of **I-LL** or **L-LL** (*m/z* $[M+H]^+$ = 358.270) and **T-LL** (*m/z* $[M+H]^+$ = 346.233). Indication of the targeted modules on the right of the chromatograms. Indications of retention times for **L-LL** and **I-LL**.

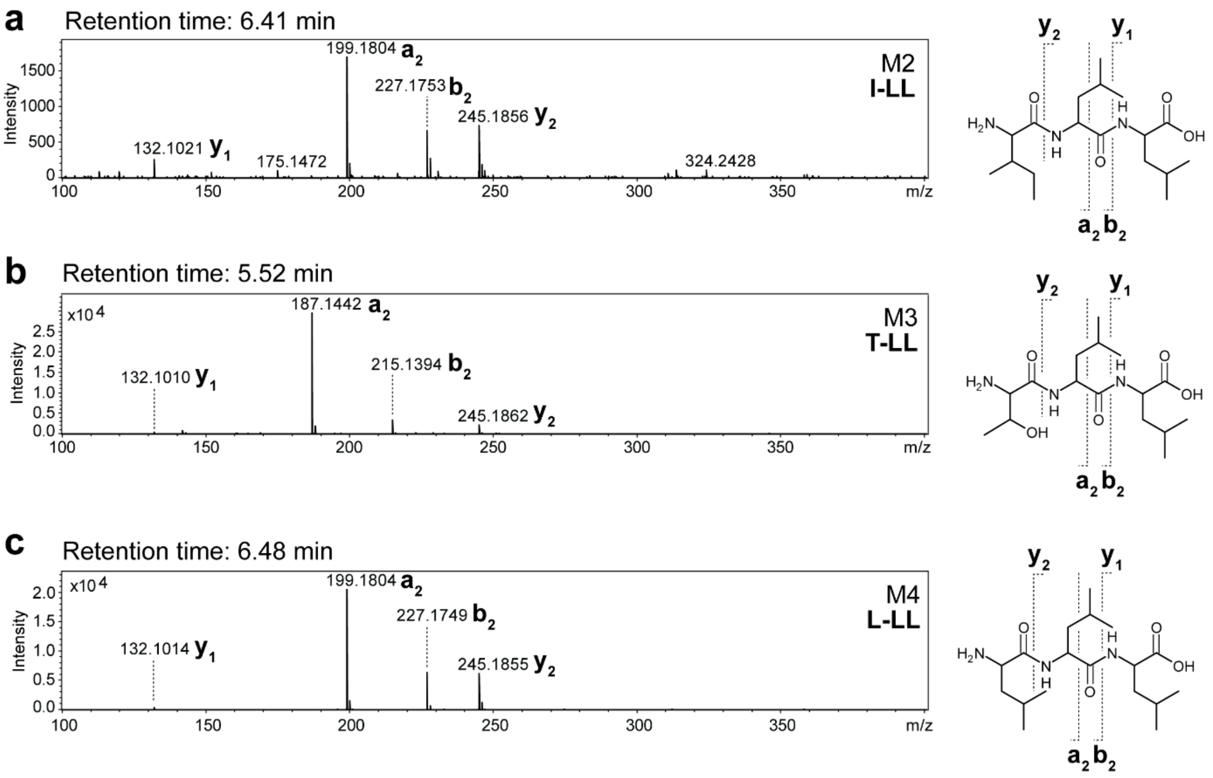

**Figure S92.** $MS^2$ spectra of HPLC-MS/MS data referred to figure S91. (**a-c**) The respective compounds are displayed, along with indications of the assigned fragments. The targeted module and peptide name is depicted within the spectra. The retention time is shown above the spectra.

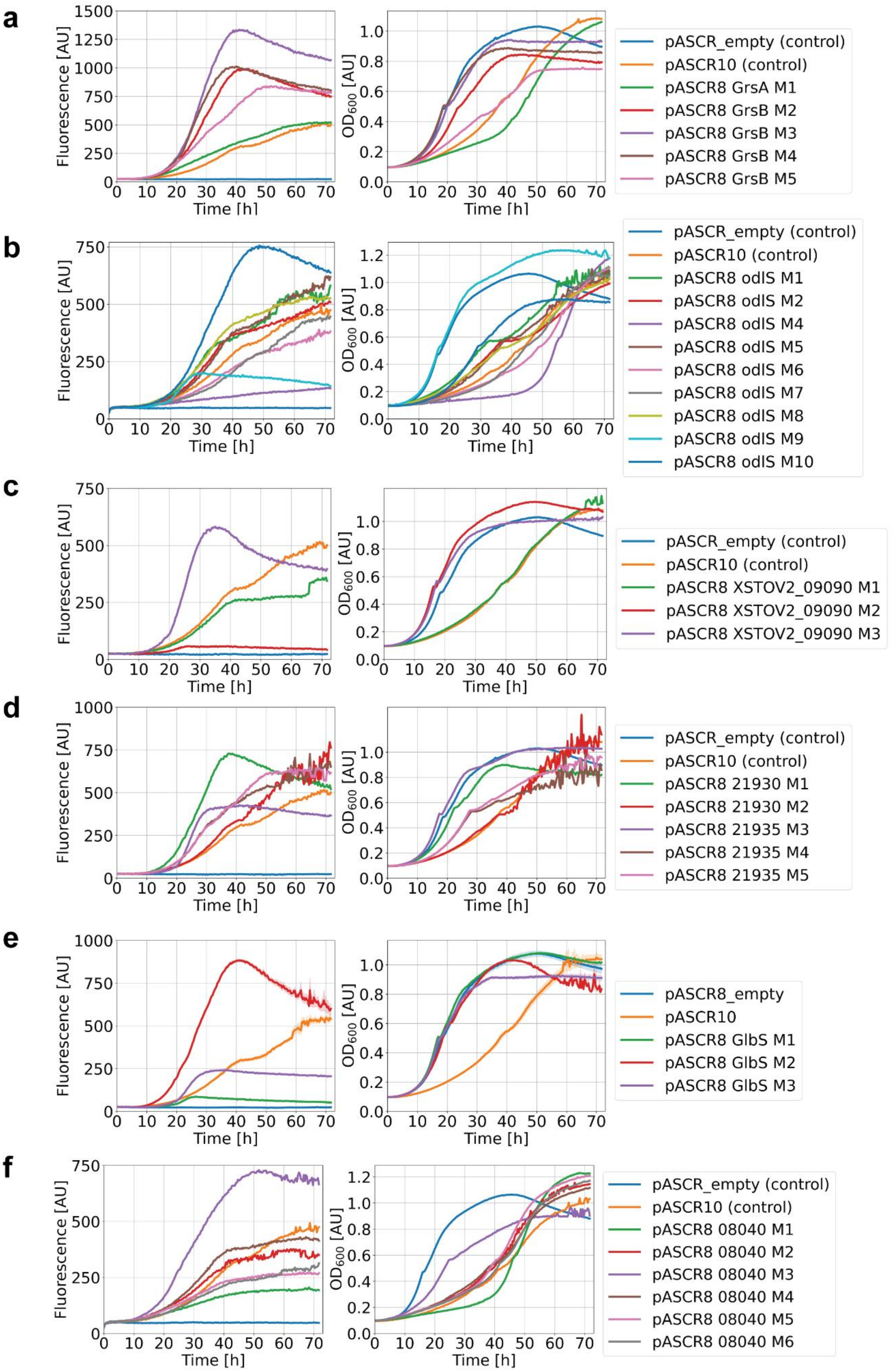

**Figure S93.** Expression and growth determination for the A domain specificity screening assay using pASCR8. *E. coli* DH10B::*mtaA* containing pASCR8 with gene insertion encoding for the individual C-A didomains from *grsAB* (**a**), *odlS* (**b**), XSTOV2_09090 (**c**), PTEMK122_21930/PTEMK122_21935 (**d**), *glbS* (**e**) and PTEMK12_08040 (**f**) were expressed and cultivated for 72 hours in a 96-micro titer plate. The fluorescence of mScarlet-I3 (Ex.: 560 nm; Em.: 605 nm, left) and the $OD_{600}$ (right) was measured for 72 hours at 22 °C. Expression of pASCR8_empty and pASCR10 served as negative control.

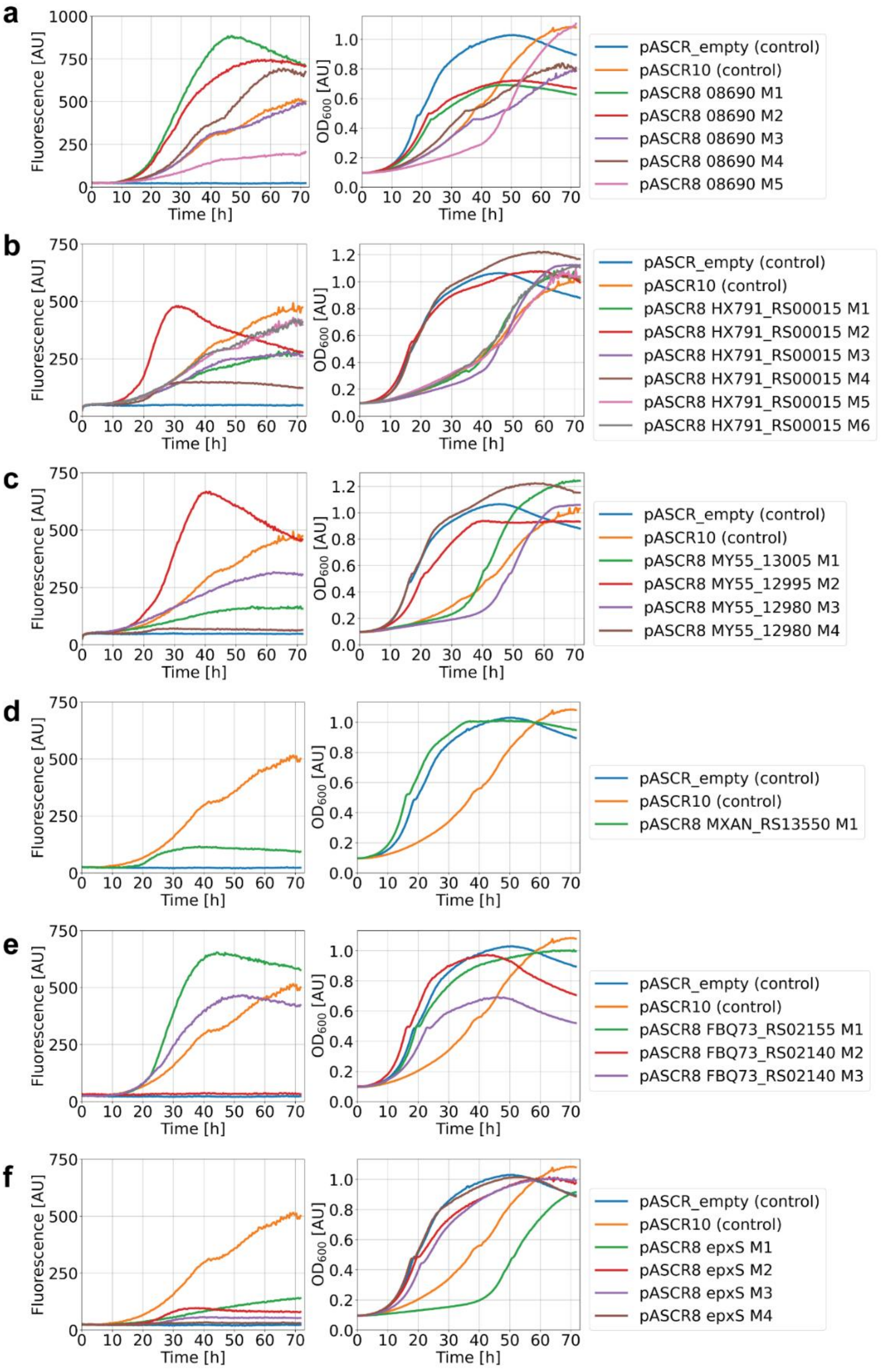

**Figure S94.** Expression and growth determination for the A domain specificity screening assay using pASCR8. *E. coli* DH10B::*mtaA* containing pASCR8 with gene insertion encoding for the individual C-A didomains from PTEMK12_08690 (**a**), HX791_RS00015 (**b**), MY55_13005, MY55_12995, MY55_12980 (*rdbS*-like synthetase) (**c**), MXAN_RS13550 (**d**), FBQ73_RS02155, FBQ73_RS02150 (**e**) and *epxS* (**f**) were expressed and cultivated for 72 hours in a 96-micro titer plate. The fluorescence of mScarlet-I3 (Ex.: 560 nm; Em.: 605 nm, left) and the $OD_{600}$ (right) was measured for 72 hours at 22 °C. Expression of pASCR8_empty and pASCR10 served as negative control.

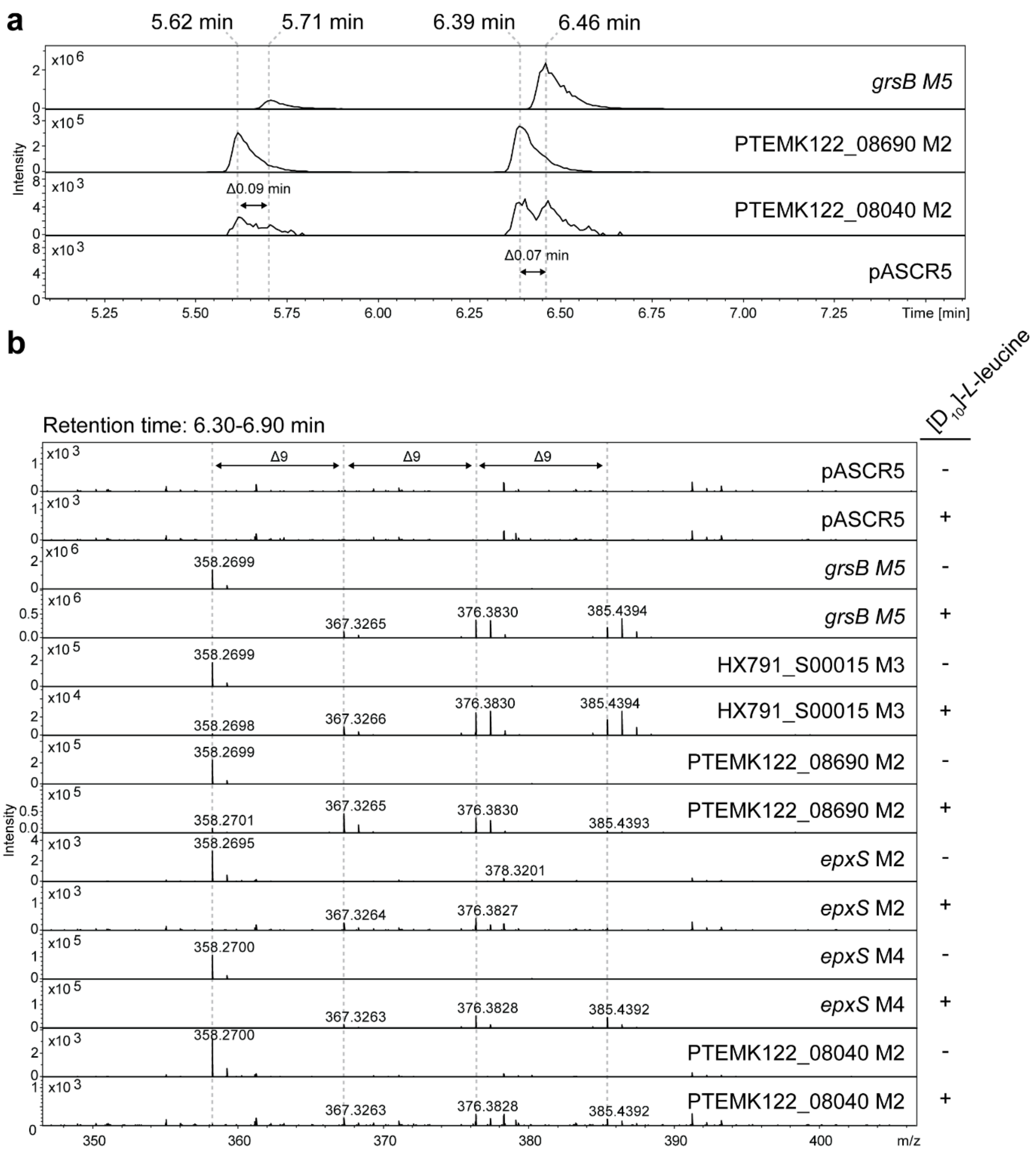

**Figure S95.** HPLC-MS/MS data of isotope feeding experiment referred to module screening of leucine or isoleucine specific modules. (**a**) HPLC-MS/MS data of *E. coli* DH10B::*mtaA* expressing C-A didomains (M) cloned into pASCR3 to compare the retention times of **I-LL** (*grsB* M5), **L-LL** (PTEMK122_08690 M2) or both (PTEMK122_08040 M2). Extracted Ion Chromatogram (EIC) of **I/L-LL** (*m/z* $[M+H]^+$ = 358.270) with indication of the cloned C-A didoamin (M). Expression of pASCR5 served as negative control. (**b**) HPLC-MS/MS data of *E. coli* DH10B::*mtaA* expressing C-A didomains cloned into pASCR3 with and without supplementation of 2 mM $[D_{10}]$-*L*-leucine to the medium. MS spectra of the retention time between 6.3 and 6.9 minutes with indication of the cloned C-A didoamin. Expression of pASCR5 served as negative control.

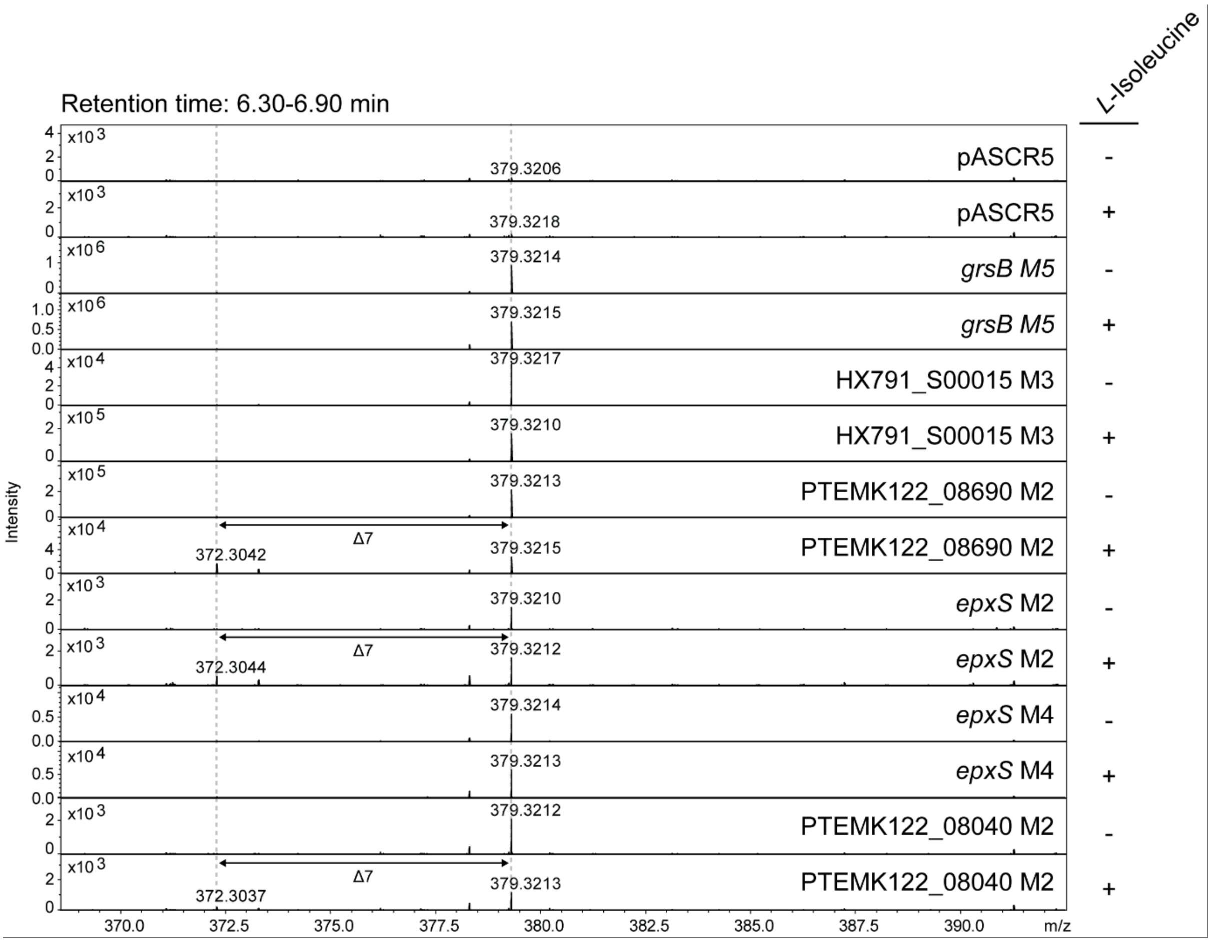

**Figure S96.** HPLC-MS/MS data of an inverse isotope feeding experiment referred to module screening of leucine or isoleucine. (**a**) HPLC-MS/MS data of *E. coli* DH10B::*mtaA* expressing C-A didomains (M) cloned into pASCR3 in ISOGRO$^{TM}$ $^{13}C$, $^{15}N$ labelled medium (Sigma-Aldrich) with and without supplementation of 1 mM unlabelled *L*-isoleucine to the medium. MS spectra of the retention time between 6.3 and 6.9 minutes with indication of the cloned C-A didoamin. Expression of pASCR5 served as negative control.

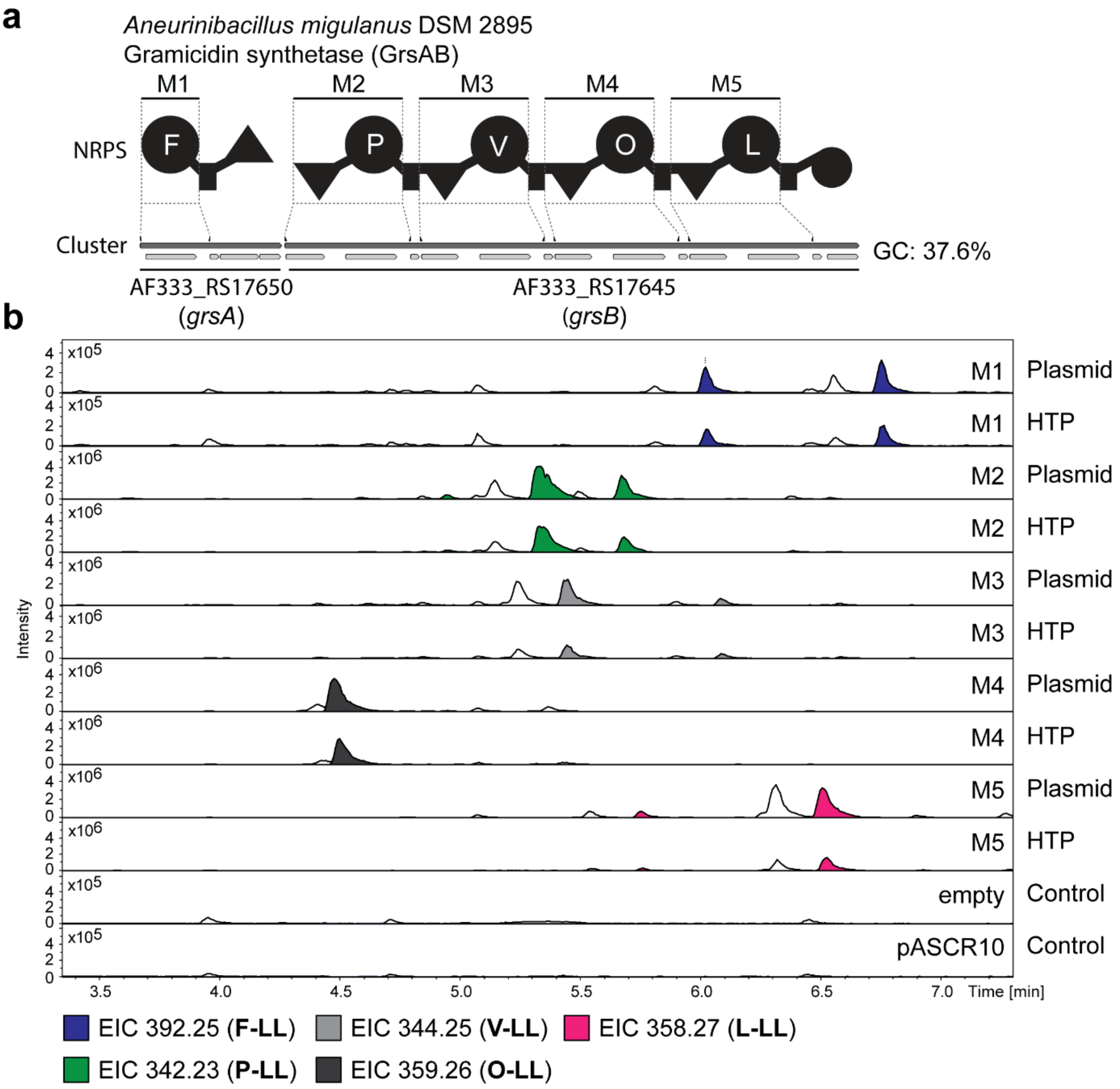

**Figure S97.** HTP workflow evaluation. (**a**) Illustration of the gene cluster *grsAB* from *A. migulanus* with schematic illustration of the encoded NRPS. Indication of the region of cloned modules M1 to M5 on gene and protein level. Utilized primers are shown above the cluster. A domain specitifity is indicated by the one-letter amino acid code (O = ornithine). GC content is depicted next to the cluster. (**b**) HPLC-MS data of *E. coli* DH10B::*mtaA* expressing M1 to M5 cloned into pASCR8 either from a verified plasmid (Plasmid) or by the HTP workflow (HTP). Expression of an empty vector and pASCR10 served as negative control. Base Peak Chromatogram (BPC, top) with Extracted Ion Chromatogram (EIC, below with colors according to the depicted legend) of **F-LL** (*m/z* $[M+H]^+$ = 392.254), **P-LL** (*m/z* $[M+H]^+$ = 342.254), **V-LL** (*m/z* $[M+H]^+$ = 344.254), **O-LL** (*m/z* $[M+H]^+$ = 359.265) and **L-LL** (*m/z* $[M+H]^+$ = 358.270). Indication of the targeted modules on the right of the chromatograms

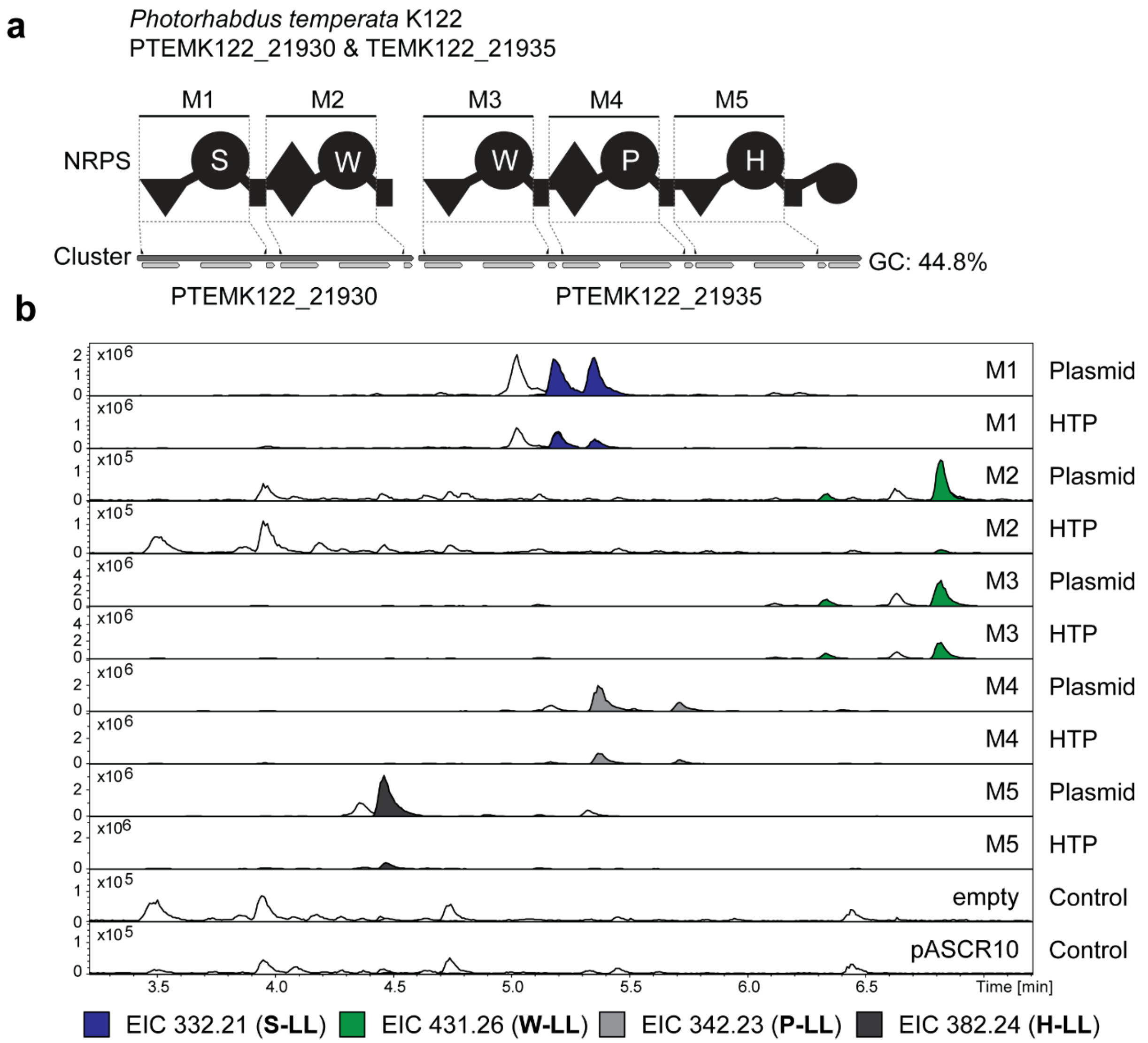

**Figure S98.** HTP workflow evaluation. (**a**) Illustration of the gene cluster PTEMK122_21930 and PTEMK122_21935 from *P. temperata* K122 with schematic illustration of the encoded NRPS. Indication of the region of cloned modules M1 to M5 on gene and protein level. Utilized primers are shown above the cluster. A domain specitifity is indicated by the one-letter amino acid code. GC content is depicted next to the cluster. (**b**) HPLC-MS data of *E. coli* DH10B::*mtaA* expressing M1 to M5 cloned into pASCR8 either from a verified plasmid (Plasmid) or by the HTP workflow (HTP). Expression of an empty vector and pASCR10 served as negative control. Base Peak Chromatogram (BPC, top) with Extracted Ion Chromatogram (EIC, below with colors according to the depicted legend) of **S-LL** (*m/z* $[M+H]^+$ = 332.217), **W-LL** (*m/z* $[M+H]^+$ = 431.265), **P-LL** (*m/z* $[M+H]^+$ = 342.254) and **H-LL** (*m/z* $[M+H]^+$ = 382.244). Indication of the targeted modules on the right of the chromatograms.

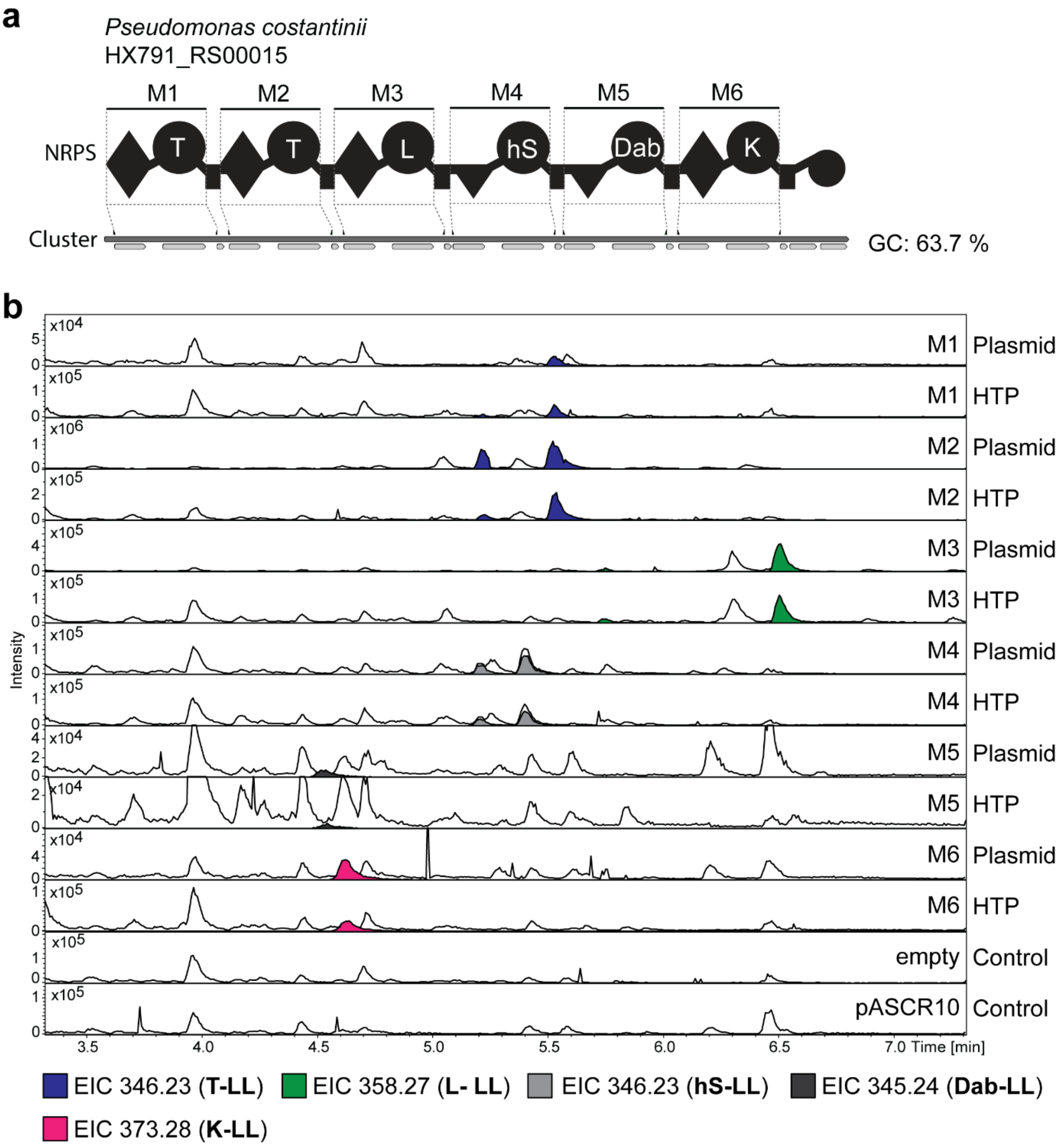

**Figure S99.** HTP workflow evaluation. (**a**) Illustration of the gene cluster HX791_RS00015 from *P. costantinii* with schematic illustration of the encoded NRPS. Indication of the region of cloned modules M1 to M6 on gene and protein level. Utilized primers are shown above the cluster. A domain specitifity is indicated by the one-letter amino acid code (hS = homoserine; Dab = diaminobuytric acid). GC content is depicted next to the cluster. HPLC-MS data of *E. coli* DH10B::*mtaA* expressing M1 to M5 cloned into pASCR8 either from a verified plasmid (Plasmid) or by the HTP workflow (HTP). Expression of an empty vector and pASCR10 served as negative control. Base Peak Chromatogram (BPC, top) with Extracted Ion Chromatogram (EIC, below with colors according to the depicted legend) of **T-LL** (*m/z* $[M+H]^+$ = 346.233), **L-LL** (*m/z* $[M+H]^+$ = 358.270), **hS-LL** (*m/z* $[M+H]^+$ = 346.233), **Dab-LL** (*m/z* $[M+H]^+$ = 345.249) and **K-LL** (*m/z* $[M+H]^+$ = 373.280). Indication of the targeted modules on the right of the chromatograms.

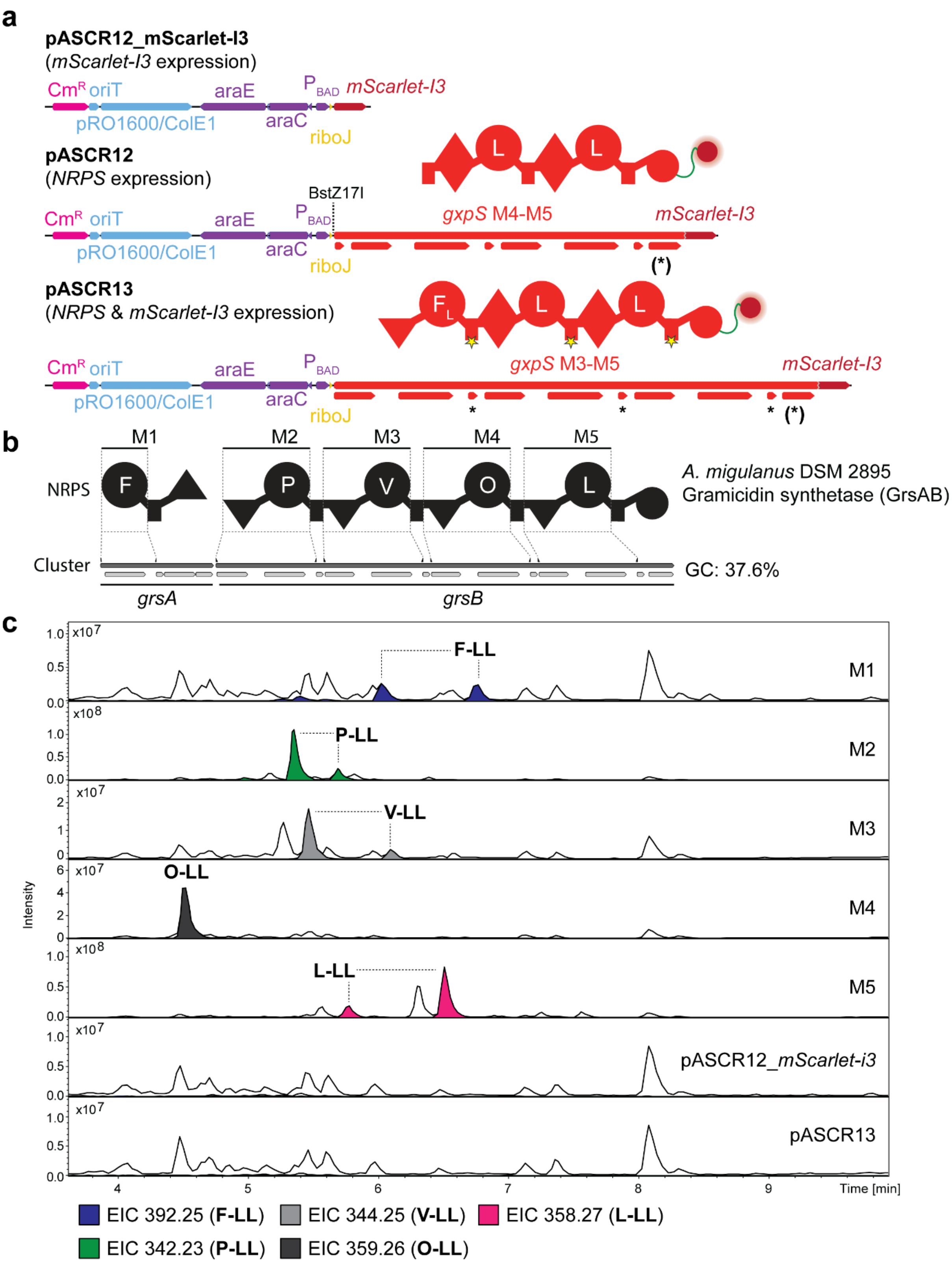

**Figure S100.** *Vibrio natriegens* ATCC14048 Δ*dns* as fast expression system. (**a**) Linearized illustration of the plasmid maps of pASCR12, pASCR12_mScarlet-I3 and pASCR13. Indication of the resistance genes, origin of replications, promoter regions, NRPS-encoding regions and added tags. An asterisk (*) indicates the introduced mutations within the T domains to inactivate the NRPS. An asterisk within brackets indicates a silent mutation at G2315 (GGT→GGG) to ensure single cleavage by the restriction enzyme. For pASCR13 the point mutation is present at G3274 (GGT→GGG). Schematic illustration of the encoded protein above. (**b**) Illustration of the gene cluster *grsAB* from *A. migulanus* with schematic illustration of the encoded NRPS. Indication of the region of cloned modules M1 to M5 on gene and protein level. Utilized primers are shown above the cluster. A domain specitifity is indicated by the one-letter amino acid code (O = ornithine). GC content is depicted next to the cluster. (**c**) HPLC-MS data of *Vibrio natriegens* ATCC14048 Δ*dns* expressing M1 to M5 cloned into pASCR12. Expression of *mScarlet-I3* and pASCR13 served as negative control. Base Peak Chromatogram (BPC, top) with Extracted Ion Chromatogram (EIC, below with colors according to the depicted legend) of **F-LL** (*m/z* $[M+H]^+$ = 392.254), **P-LL** (*m/z* $[M+H]^+$ = 342.254), **V-LL** (*m/z* $[M+H]^+$ = 344.254), **O-LL** (*m/z* $[M+H]^+$ = 359.265) and **L-LL** (*m/z* $[M+H]^+$ = 358.270). Indication of the targeted modules on the right of the chromatograms

#### 4 Supplementary Data

The attached supplementary data provides gene sequences in the fasta format for the gene locus tags listed in table S1 of *Photorhabdus temperata* K122 (PTEMK122_08040, PTEMK122_08690, PTEMK122_21930 & PTEMK122_21935) and *Xenorhabdus stockiae* (XSTOV2_09090).
